# Standing genetic variation fuels rapid evolution of herbicide resistance in blackgrass

**DOI:** 10.1101/2021.12.14.472587

**Authors:** Sonja Kersten, Jiyang Chang, Christian D. Huber, Yoav Voichek, Christa Lanz, Timo Hagmaier, Patricia Lang, Ulrich Lutz, Insa Hirschberg, Jens Lerchl, Aimone Porri, Yves Van de Peer, Karl Schmid, Detlef Weigel, Fernando A. Rabanal

## Abstract

Repeated herbicide applications exert enormous selection on blackgrass (*Alopecurus myosuroides*), a major weed in cereal crops of the temperate climate zone including Europe. This inadvertent large-scale experiment gives us the opportunity to look into the underlying genetic mechanisms and evolutionary processes of rapid adaptation, which can occur both through mutations in the direct targets of herbicides and through changes in other, often metabolic, pathways, known as non-target-site resistance. How much either type of adaptation relies on *de novo* mutations versus pre-existing standing variation is important for developing strategies to manage herbicide resistance. We generated a chromosome-level reference genome for *A. myosuroides* for population genomic studies of herbicide resistance and genome-wide diversity across Europe in this species. Bulked-segregant analysis evidenced that non-target-site resistance has a complex genetic architecture. Through empirical data and simulations, we showed that, despite its simple genetics, target-site resistance mainly results from standing genetic variation, with only a minor role for *de novo* mutations.

## Introduction

Unintendedly, humans have been carrying out a series of – often alarming – evolutionary “experiments” that have selected for resistance to herbicides in many agricultural weeds over the past several decades. Among these, blackgrass (*Alopecurus myosuroides*) has become the most economically damaging herbicide resistant weed in Europe^1, 2^. In England alone, the annual cost of resistance was estimated to be £0.4 billion (€0.47 billion) in lost gross profit^3^.

We distinguish two resistance mechanisms. First, there is target-site resistance (TSR), which is caused by mutations in the genes encoding the proteins targeted by herbicides^4–7^. Second, there is non-target-site resistance (NTSR), which is associated with enhanced metabolic processes as herbicide detoxification or translocation^6, 8^.

To better understand how either type of resistance arises and comes to dominate *A. myosuroides* populations, we need to learn more about the population structure and genetic diversity of the species across Europe. Previous regional studies have only found weak, if any, population structure, suggesting a very rapid and recent spread of the species^9, 10^.

Two important drivers of the modes of evolution of herbicide resistance are the genetic architecture of the trait and the types of mutations that can give rise to it. TSR is conferred by mutations in single genes, with a very small number of coding sequence changes allowing for herbicide resistance without eliminating activity of the targeted protein. As in many other weeds, TSR in *A. myosuroides* has increased rapidly^11, 12^, and as a consequence, herbicides that inhibit the action of acetolactate synthase (ALS) and acetyl-CoA carboxylase (ACCase) have widely lost their efficacy as weed control agents.

In contrast, several different gene families, which encode detoxifying enzymes and transporters such as cytochrome P450 monooxygenases, glutathione S-transferases, ATP-binding cassette Transporters, MFS-type transporters and glycosyltransferases, have been found to contribute to NTSR (reviewed in ref. ^13^). NTSR now accounts for a substantial proportion of resistance in agricultural fields and is becoming a major focus of herbicide resistance research^11^. Since the genetic basis of NTSR often appears to be oligogenic, convergent evolution should be less common than for TSR. This is supported by recent findings in glyphosate resistant weeds in North America^14, 15^, but whether this extends to other species and other herbicides is not known.

Finally, the rapid speed with which herbicide resistance spreads in individual weed species raises the question whether this is primarily due to repeated selection for rare *de novo* mutations, or more commonly arising from standing genetic variation, with herbicide resistant alleles segregating in the population already before the widespread adoption of herbicide application. An optimal framework to distinguish between these hypotheses is provided by forward-in-time genetic simulations^16^.

To enable a better understanding of herbicide resistance evolution in *A. myosuroides*, we have generated a high-quality reference genome with PacBio long reads. Genotyping with double-digest restriction-site associated DNA (ddRAD) sequencing markers in 47 European field populations revealed considerable geographical population structure along with high effective population sizes. To begin to shed light on NTSR mechanisms in the species, we used bulked-segregant analysis of natural field populations to identify promising candidate genes. To characterise TSR haplotype diversity at the field level, we generated PacBio long-read amplicons for the known TSR genes ACCase and ALS, and compared our empirical data with the results from probabilistic models of adaptation via selective sweeps and forward simulations. We infer that standing genetic variation is the most likely mechanism behind the TSR mutations of independent origin, with only a minor role for *de novo* mutations.

## Results & Discussion

### Genome assembly and annotation

For genome sequencing, we selected a single plant from a herbicide sensitive population (Appels Wilde Samen GmbH, Darmstadt) from Germany and ascertained that it did not carry known TSR mutations at the ACCase, ALS and psbA loci (see Methods). Previous genome size estimates of *A. myosuroides* based on Feulgen photometry ranged from 4.2 Gb^17^ to 4.7 Gb^18^. To estimate genome size of the selected individual more accurately, we performed flow cytometry using rye (*Secale cereale*) as a reference^19^. We estimated the haploid genome size of *A. myosuroides* to be 3.56 Gb (Figure 1a). Next, we generated ∼90x genome coverage of PacBio continuous long reads, ∼44x genome coverage of Illumina PCR-free short reads and ∼66x genome coverage of Hi-C chromatin contact data. We *de novo* assembled the genome with FALCON-Unzip^20^, deduplicated primary contigs with purge_dups^21^ and scaffolded contigs with HiRise^22^ (Table 1). The size of the final assembly was 3.53 Gb and consisted of seven super-scaffolds (Figure 1b, Supplementary Figure 1a), in agreement with the known karyotype of the species with seven chromosomes ^18^.

**Figure 1.**
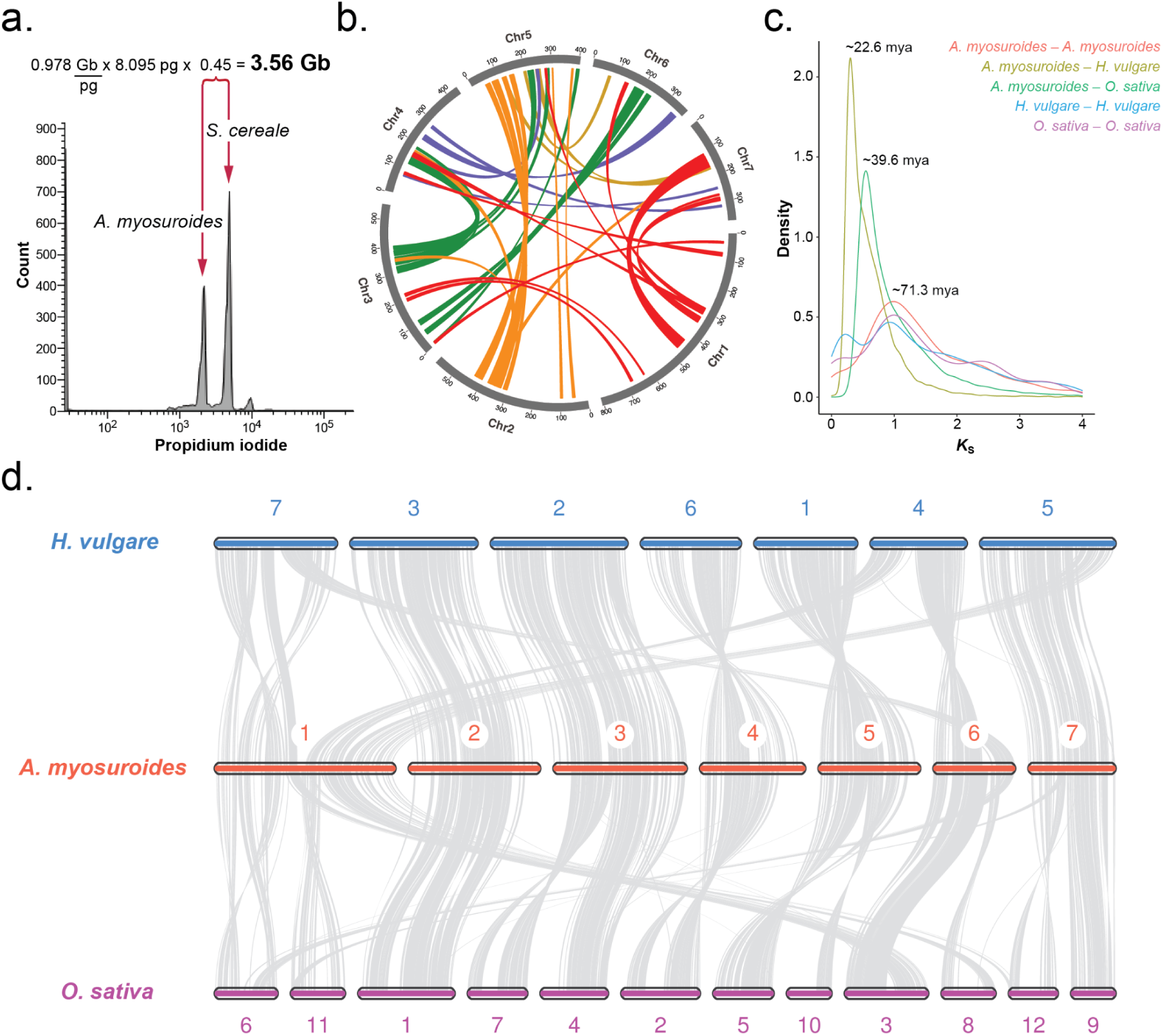
A reference genome of an *Alopecurus myosuroides* individual from a German herbicide-sensitive population. **a,** Histogram of relative DNA content obtained by flow cytometry analysis of propidium iodide stained nuclei of *A. myosuroides* and the reference standard *Secale cereale* cv. Daňkovské (diploid genome size = 16.9 pg). **b,** Circos plot of the *A. myosuroides* genome, with coloured lines connecting anchor pairs (genes in the collinear regions) with synonymous substitution rates (*K*_S_) > 0.5. Numbers represent megabases. **c,** *K*_S_ distributions for paralogs within the *A. myosuroides*, *Hordeum vulgare*^23^ and *Oryza sativa*^24^ genomes, and for orthologs between *A. myosuroides* and each of those species. Divergence time, expressed as million years ago (mya), was estimated based on 7.0 x 10^-9^ as the substitution rate in grasses^25^. **d,** Syntenic relationships between the chromosomes of *A. myosuroides* and other sequenced grasses, including *H. vulgare* (top) and *O. sativa* (bottom).

**Table 1.** Genome assembly metrics.

| Descriptor | Value |
| --- | --- |
| Total assembly size | 5,218,837,661 bp |
| Number of contigs | 6,215 |
| Contig N50 | 1,783,999 bp |
| Largest contig | 11,174,859 bp |
| Chromosome-level assembly size | 3,529,081,863 bp |
| Chromosome N50 | 554,019,051 bp |
| Largest chromosome | 807,086,175 bp |
| Number of protein-coding genes | 50,029 |
| Mean gene length | 2,789 bp |
| BUSCO score | C:94.6%[S:82.0%,D:12.6%],F:0.9%,M:4.5% |
| TE content | Class I [LTR:63.80%,non-LTR:0.07%]<br>Class II [TIR:10.92%,Helitron:8.21%]<br>Other repeated region:2.15% |
| Contig metrics are shown before deduplication. Benchmarking Universal Single-Copy Orthologs (BUSCO)<sup>27</sup> scores were obtained with the 'embryophyta_odb10' set (n=1,614). Complete (C), single copy (S), duplicated (D), fragmented (F) and missing (M) genes are indicated. Transposable element (TE) content was determined with the Extensive *de-novo* TE Annotator (EDTA)<sup>28</sup>. |  |

Given that in plants, transposable elements are a major driver of genome size, it is not surprising that repetitive sequences account for 85.16% of the *A. myosuroides* genome, with 63.8% classified as long terminal repeats (LTR) retrotransposons (Table 1). We annotated 50,029 protein-coding genes based on a combination of both RNA-seq and PacBio Iso-Seq long transcripts from five different tissues (i.e., leaves, roots, flowers, anthers and pollen), *ab initio* prediction and protein homology. Transcriptome data supported 87.45% of the annotated genes, and 95% of all genes could be assigned functions with InterProScan^26^. In fact, 94.6% of the BUSCO v4.0.4^27^ predicted protein sequences are found as complete genes (82% as single copy, 12.6% as duplicated genes, n=1,614). On average, protein-coding genes in *A. myosuroides* are 2,789 bp long and contain 3.71 exons (Table 1).

Plant genomes typically contain both whole-genome and segmental duplications. We therefore investigated collinear regions indicative of recent duplications. When we analyzed the divergence of closely related paralogs present in these regions based on synonymous substitution rates (*K*_S_), we noticed two main peaks, one at *K*_S_ ∼0.16 and another one at *K*_S_ ∼1.2 (Supplementary Figure 1b). The *K*_S_ of the first peak is unusually low, and would normally indicate very recent duplicates. To explore the nature of the gene pairs with low *K*_S_, we extracted all gene pairs in these regions with *K*_S_ <= 0.5 and asked how they are distributed in the genome. Collinear blocks containing these pairs are generally very close and always within the same chromosome (Supplementary Figure 1c), while pairs with *K*_S_ > 0.5 are located in different chromosomes (Figure 1b). One explanation would be that these blocks are the products of recent duplication events, although there is not much evidence for large-scale local duplications in plant genomes. Alternatively, they could be an artifact of the assembly process, as in highly heterozygous genomes, different alleles can be assembled independently into different contigs. If these duplicates are not properly purged, which is particularly difficult if alleles are very dissimilar, then during scaffolding they are placed close to each other on the same chromosome. With the data at hand, it is difficult to distinguish between these two possibilities, but based on the close paralogs being almost always present close to each other, we favor the second explanation. The second peak (*K*_S_ ∼1.2), mostly representing paralogs in different chromosomes (Figure 1b, Supplementary Figure 1b), coincides with a known whole-genome duplication (WGD) event common in all grasses^29, 30^ that occurred ∼70 million years ago (mya). The *K*_S_ distributions for orthologous gene pairs between blackgrass and other grasses indicate a more recent divergence time with rice (∼39.6 mya) and barley (∼22.6 mya) (Figure 1c).

Chromosome level synteny with other grasses was high, particularly with the more closely related barley genome (Figure 1d). For instance, chromosomes 2, 3, 4, 5 and 7 in blackgrass have a near 1:1 relationship with chromosomes 3, 2, 6, 1 and 5 in barley, respectively. An exception is chromosome 1 in blackgrass (807 Mb), which contains sequences that are syntenic with chromosomes 4, 5 and 7 in barley. In conclusion, this first chromosome-level genome assembly of blackgrass displays the expected characteristics typically found in grass genomes.

### Population structure

Our new reference genome then allowed us to easily assess the distribution of genome-wide diversity across Europe. To this end, we performed ddRAD-Seq in 1,123 individuals (47 populations from nine European countries with 22-24 individual plants each; Figure 2a), and defined 109,924 single nucleotide polymorphism (SNP) markers with an average sequencing depth of 22.6x (Supplementary Figure 2a). A clear phylogeny per country was not discernible from the maximum likelihood tree (Figure 2a), but a Treemix tree per population captured the geographic distribution at the country scale (Supplementary Figure 3).

**Figure 2.**
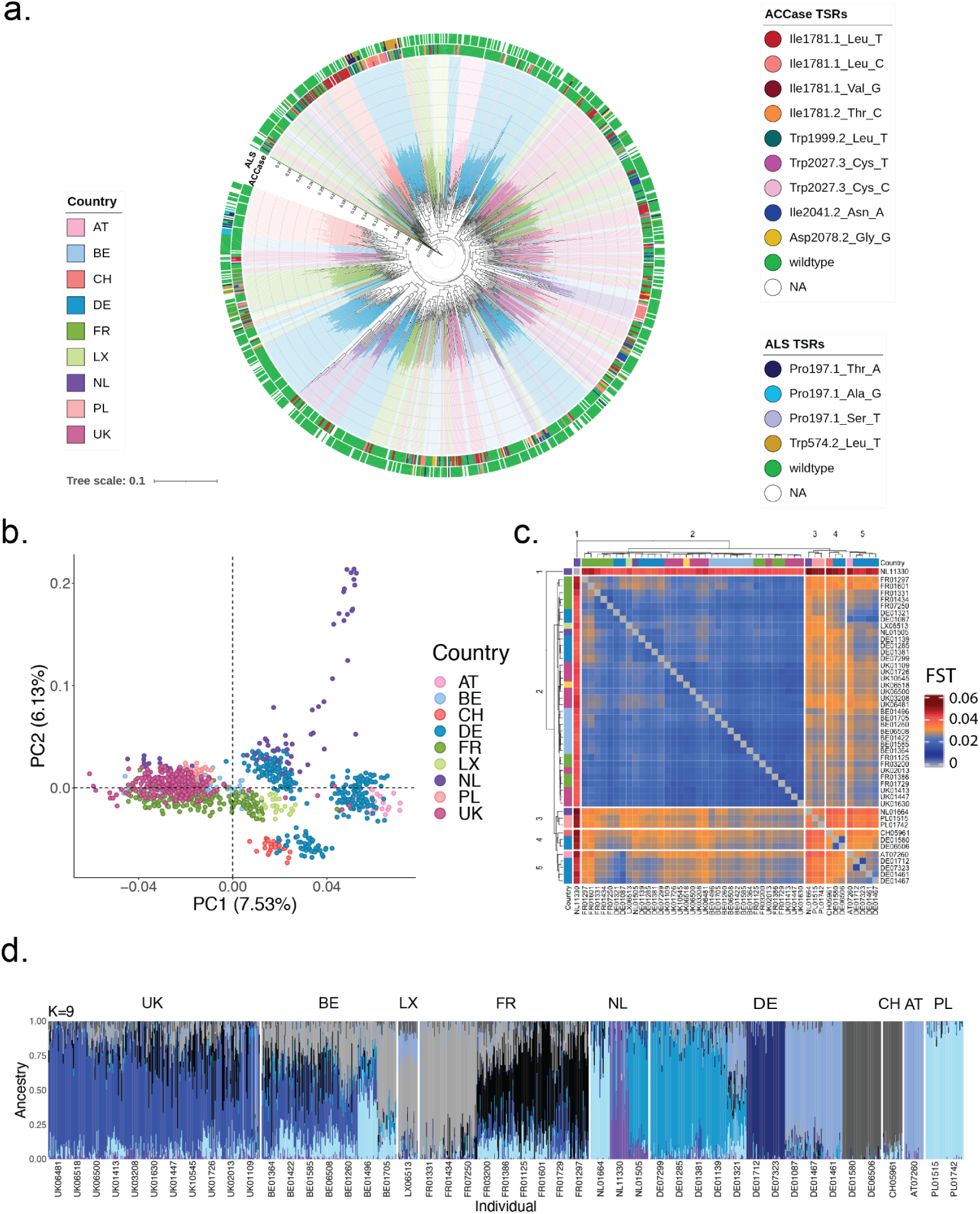
Population structure analysis of 47 European *A. myosuroides* populations with 109,924 genome-wide ddRAD-seq markers. **a,** Maximum likelihood phylogenetic tree. Branch ends are marked in country colours. TSR mutations for ALS and ACCase in each individual are indicated in the outer and inner rings, respectively. **b,** Principal component analysis (PCA) showing the first two eigenvectors. The genetic variance explained by the two principal components is shown in parenthesis. Colours reflect country-specific origin of the populations. **c**, Fixation index (F_ST_) heatmap displaying the F_ST_ values between the different populations from 0 - 0.02 (blue) to 0.04 - 0.06 (red). Annotation colours of the branch tips reflect country-specific origins of the population. **d,** Admixture proportions with ancestry groups of K=9. Admixture proportions with ancestry groups of K=7 and K=8 can be found in Supplementary Figure 4. Every bar is an individual, and these are grouped by population. Austria (AT), Belgium (BE), Switzerland (CH), Germany (DE), France (FR), Luxembourg (LX], Netherlands (NL), Poland (PL), United Kingdom (UK).

Overall genetic differentiation between populations was low (F_ST_ range: 0.014 to 0.052, n=47; Figure 2c), consistent with other studies of *A. myosuroides*^9, 10^ and other wild grasses such as *Panicum virgatum*^31^. The relatedness of individuals within populations was high (F_IS_ = 0.11; range 0.063 to 0.124). In the admixture analysis, we could identify between 7 and 9 ancestry groups (Figure 2d, Supplementary Figure 4c) that were consistent with the clusters formed in a principal component analysis (PCA; Figure 2b, 2d, Supplementary Figure 4a). Populations from Belgium (BE), the United Kingdom (UK), Luxemburg (LX) and France (FR) clustered together and had high genetic similarity. The population from the Netherlands (NL) NL11330 was most differentiated from all others with F_ST_-values up to 0.052 (Figure 2c). Germany (DE) was divided into three subclusters, one having common ancestry with the population from Switzerland (CH), one with the Austrian (AT) and the third cluster being highly admixed. The populations from Poland (PL) shared common ancestry with the Netherland population NL01664. In summary, while population differentiation is low, there is clear geographical population structure across Europe.

The mean observed SNP heterozygosity was 0.11, with no significant difference between populations that were under herbicide selection and those that were not (Supplementary Figure 2b). This agrees with previous suggestions that genome-wide genetic variation in *A. myosuroides* was not affected by herbicide selection^9^.

We estimated Watterson theta θ_W_ on the 1.1% sequenced fraction of our genome. Our θ_W_ (mean = 0.0047) are within the range of other outcrossing plant species^32^. With these θ_W_ estimates and the mutation rate of 3.0 x 10^-8^ from maize^33^, we determined effective population sizes ranging from 30,366 to 41,941 individuals (Supplementary Figure 2c). Among countries for which we had more than six populations, Germany had significantly smaller (p-value range: 0.01 - 0.03) effective population sizes than France, Belgium and the United Kingdom (Supplementary Figure 2d). This observation is reminiscent of the westward range expansion of the species detected within the United Kingdom^10^, and suggests that this trend might be true at a European scale; however, more extensive sampling would be needed to critically examine this hypothesis. Given that we have estimated our effective population sizes in *A. myosuroides* mostly from populations under selection that have already experienced a decrease in population sizes, it is very likely that we rather underestimate the long-term effective population sizes of our *A. myosuroides* field populations^34^. Messer and Petrov (2013) noted that temporal fluctuations in population size can strongly influence estimates of effective population size, especially in recent bottlenecks, as would be the case with adaptation processes to herbicides^35^. Adaptation to strong selection pressure is a rapid process, and the probability of adaptive mutations is higher for larger population sizes ^36, 37^.

### Genetics of non-target site resistance

Herbicide resistance may originate from target-site mutations or from less specific mechanisms known as non-target site resistance (NTSR). We phenotyped half of our collection for resistance to the ACCase inhibitor pinoxaden (Figure 3a), which is particularly problematic^2, 38^. We not only found differences in the degree of resistance, but subsequent targeted analysis of ACCase (see below) revealed that almost half of the resistance to ACCase inhibitors could not be explained by known TSRs at the ACCase locus, and must be due to NTSR (Figure 3b). This is consistent with previous studies that reported that NTSR, which most likely has a more complex genetic architecture than TSR genes, accounts for a significant proportion of total herbicide resistance in *A. myosuroides*^11^. To begin to understand the genetics and possible loci underlying NTSR, we performed bulked-segregant analysis based on phenotypic classification of resistance of the ACCase inhibitor Axial*®*. For this purpose, we selected a pool of 61 Axial*®* resistant individuals in which we had not found TSR mutations by amplicon analysis, and matched these with a pool of 61 individuals that were highly sensitive to Axial*®* (Supplementary Figure 5).

**Figure 3.**
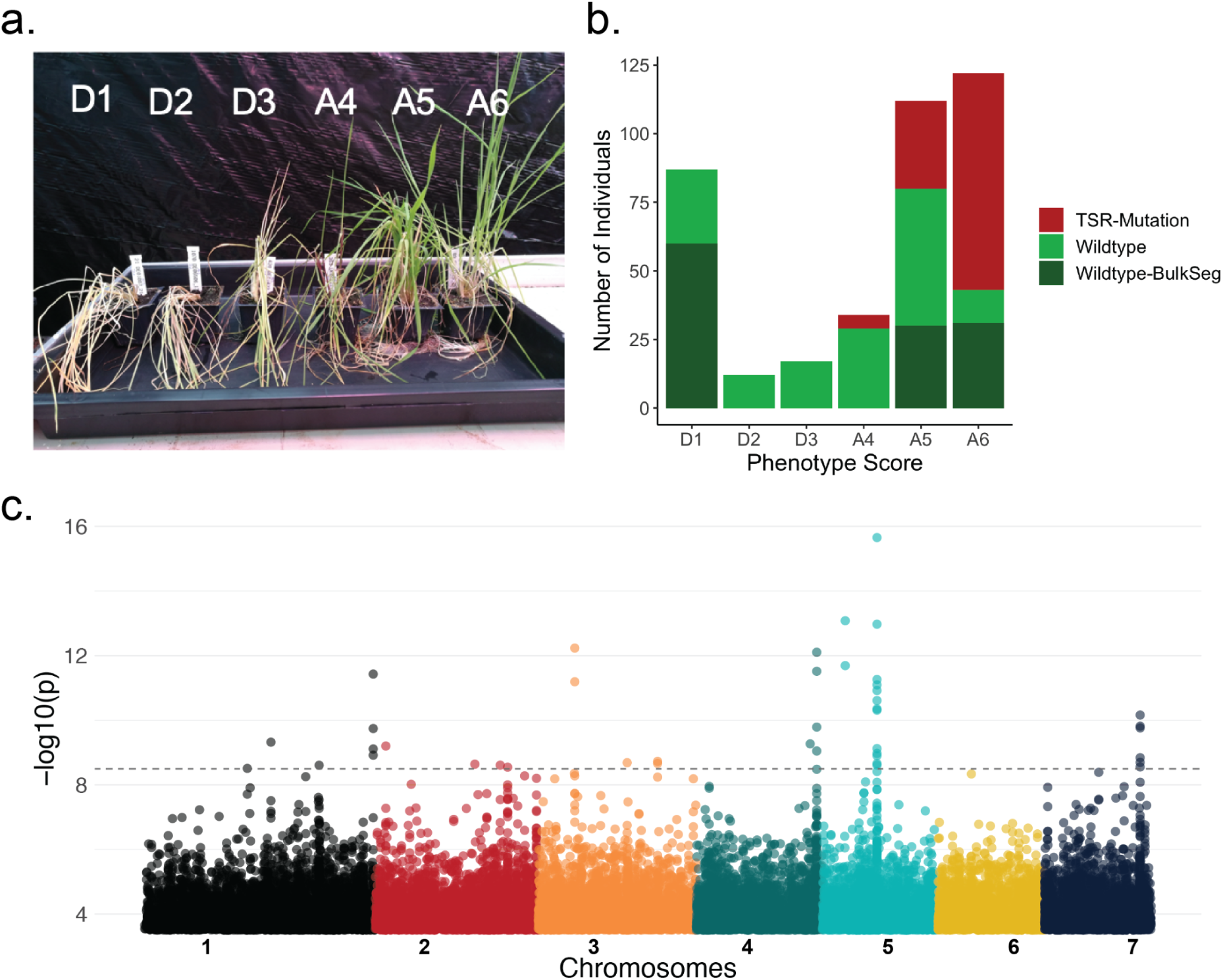
Bulked-segregant analysis. **a,** Phenotyping with ACCase inhibitor pinoxaden (Axial® 50). Gradient score starting from D1=Dead (no more green material) to A6 =Alive (no difference from untreated control) **b,** Phenotyping combined with genetic information of the ACCase genotyping. In red, the number of individuals that carry a TSR mutation. In green, wildtype individuals that do not carry known TSR mutations. In dark-green, wildtype individuals that were selected for the two extreme bulks of the bulked-segregant analysis (D1= sensitive bulk, A5 & A6 = resistant bulk) **c,** Manhattan plot, colour code represents the 7 chromosomes, 0.05 Bonferroni p-value threshold for 16 million SNPs: p= -log_10_(3.1e^-09^) = 8.5.

We separately pooled DNA libraries from the 61 individuals in each cohort, sequenced these and used BayPass^39, 40^ to find regions of the genome where the two pools were associated with contrasting allelic patterns (Figure 3c). While we found several candidate genes in the vicinity of highly associated SNP markers (Extended data File 1), we also noticed that most of these had unusually high read coverage (Supplementary Figure 6a), increasing statistical power to detect differences between the pools in these regions (Supplementary Figure 6b). It is therefore likely that we missed additional associations, which would be consistent with complex genetics and multiple loci with modest individual effect sizes contributing to the trait.

### Haplotype networks of herbicide target genes ACCase and ALS

Most of the work on the molecular mechanisms underlying herbicide resistance has focused on mutations in the genes that encode the enzymes inhibited by herbicides. Two prominent herbicide targets are ALS and ACCase, with mutations at multiple codons at these loci known to confer inhibitor resistance^4–7^. To understand the diversity not only of specific mutations, but also of entire haplotypes on which these mutations arose, we aimed to characterize extended linked sequences surrounding the ALS and ACCase loci. To preserve haplotype information, we amplified the entire coding sequences including introns, ∼13.2 kb for ACCase and ∼3.6 kb for ALS, by PCR and analyzed complete amplicons with PacBio Circular Consensus Sequencing (CCS) for all individuals in our European collection. We applied very stringent criteria to call haplotype sequences in our dataset – requiring high accuracy (>99% or q20) and a minimal CCS read depth per sample of 25X. This enabled us to characterize entire haplotypes for 1,046 individuals for ACCase and 842 individuals for ALS. We were able to recover two haplotypes for the vast majority of our samples that passed quality control filters, 84.9% for ACCase and 59.8% for ALS. We assume that the remaining individuals are homozygous for the same haplotype.

Some TSR mutations were less common than others. For example, Trp2027Cys and Asp2078Gly were underrepresented, consistent with these mutations reducing fitness in the absence of herbicide selection in other species^41,42,43,44^. The most common mutation was Ile1781Leu, consistent with pleiotropic effects of this substitution that increase fitness also in the absence of selection^41, 42^ (Figure 3a-c, Supplementary extended data Figure 1). Finally, TSR mutations typically act in a dominant fashion, and the majority of TSR mutations (71.4%) at ACCase in our data occurred as heterozygotes.

Principal component analysis (PCA) of ACCase haplotypes clusters the European populations in three distinct groups (Supplementary Figure 7). Since each group included representatives from all countries (including those countries for which we only analysed single populations), and since alleles without obvious TSR mutations were present in all clusters, the major haplotype groups likely arose before the geographical spread of the species in Europe. TSR substitutions Ile1781Leu, Ile1781Thr, Trp2027Cys and Ile2041Asn were the most wide-spread and found in all European groups, while Asp2078Gly and Ile1781Val were found in two, and Trp1999Leu in one group. This pattern of the same TSR mutation appearing independently in separate geographic locations across Europe (Figure 2a) extends previous observations made at local or country scales using small samples of short amplicons that included only a limited number of variable sites for haplotype detection^45–47^.

To better characterise ACCase haplotype diversity, we inferred haplotype trees and networks at the level of single fields (Figure 4, Supplementary extended data Figure 1). We observe haplotype networks of varying complexity (Figure 4a-c), likely reflecting the selection pressure to which each population was subjected. If the allele frequency of a single mutation – on a single haplotype – increases rapidly in a population, this is called a ‘hard sweep’. If, on the other hand, there are several different haplotypes in a population that confer resistance – whether they all carry the same beneficial mutation or different ones – and increase in frequency at the same time, this is referred to as a ‘soft sweep’^34^. In our collection, only four out of the 27 populations with recorded TSRs contain a single TSR haplotype – and in these four cases, the TSR haplotypes were found at low frequency, with fewer than 10% of sequences having a TSR mutation. In principle, this pattern may well reflect an early state of a hard sweep. That the other 24 populations contain at least two haplotypes with TSR mutations indicates that soft sweeps are the norm (Supplementary Table 1). In 14 of these populations, we found different haplotypes with the same TSR mutation resulting from multiple independent mutation events, as opposed to the same TSR mutation being transferred to other haplotypes by recombination (Figure 4d, Supplementary extended data Figure 1). This observation confirms in an unbiased manner inferences from earlier explorative studies^45–47^. We also found seven instances in which two or three different TSR mutations had arisen in a single field, from the same haplotype (Figure 4b,c,e, Supplementary extended data Figure 1). The maximum number of independent (non-recombinant) ACCase TSR haplotypes within a field population was 10 (Supplementary Table 2).

**Figure 4.**
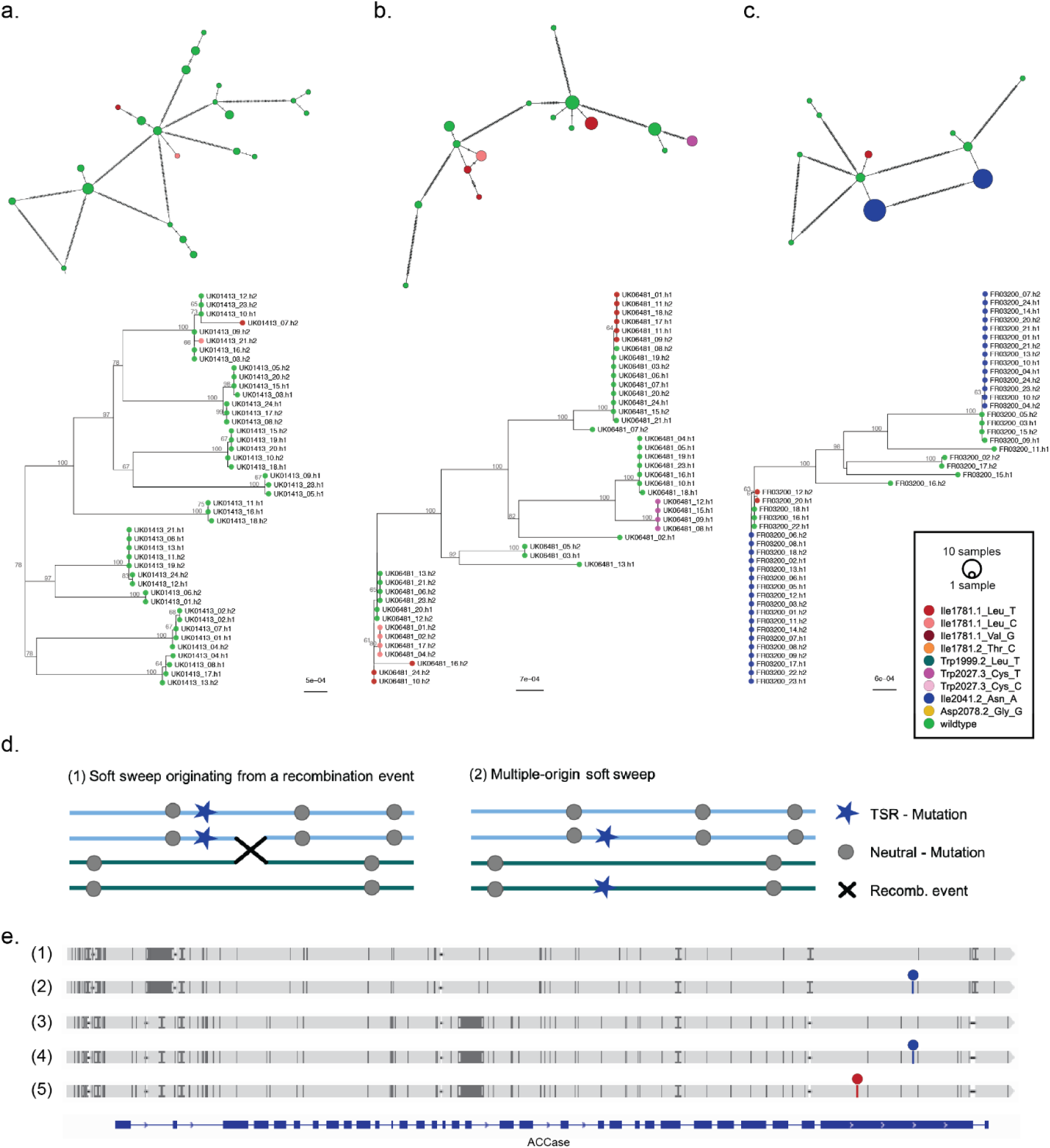
Haplotype analysis of the complete ACCase gene (13.2 kb). **a,** Network and maximum likelihood (ML)-tree of 44 haplotypes from the sensitive reference population UK01413 (HerbiSeed standard), which has not been under herbicide selection. The colour code in all networks and trees indicates different target-site resistance (TSR) mutations, with haplotypes that have wild-type sequence at known TSR positions in green. **b,** Network and ML-tree of 44 haplotypes from the British population UK06481, which shows a selection pattern characteristic of an emerging soft sweep for the TSR mutations. **c,** Network and ML-tree of 46 haplotypes from the French population FR03200. A predominant soft sweep pattern for the TSR mutation Ile2041.2_Asn_A is apparent. **d,** Schematic representation of alternative origins of soft sweep patterns: recombination *vs* independent mutation events. **e,** Two distinct wild-type haplotypes in the FR03200 population (1 and 3) that have given rise to identical TSRs, haplotypes (2) and (4). In addition, wild-type haplotype (3) has given rise to a second TSR, haplotype (5). Positions of TSR Ile1781.1_Leu_T and TSR Ile2041.2_Asn_A mutations are marked with red and blue lollipops, respectively.

In the complete assembly of the *A. myosuroides* genome we discovered at least two copies of the ALS gene in chromosome 1 (see Methods). These copies encode functional open reading frames, and high quality Iso-Seq reads (>99.9% or q30) span full-length transcripts (Supplementary Figure 8). The copy most similar to the GenBank sequence AJ437300.2 (ref. ^48^) was designated ALS1, and we selectively amplified ALS1 with primers that should not target the other ALS loci (or locus). The existence of multiple ALS copies in *A. myosuroides* may have confounded previous studies, which relied on primers in the coding region to genotype ALS TSR mutations. This strategy was used in a pyrosequencing assay^49^ commonly used for this type of study, which, differently from our work, did not lead to the identification of homozygous Pro197Thr mutations^50, 51^.

ALS1 haplotypes fell into three major Europe-wide groups (Supplementary Figure 9). In our collection, TSR mutations for this gene were only present in Germany, France, United Kingdom and Poland. TSR mutations Pro197Thr and Trp574Leu were found in two of these groups, Pro197Ala and Pro197Ser only in one, and no obvious TSR mutation was found in the third group. Similar to ACCase, although less often, two or more TSR haplotypes of independent origin could be detected within single fields, in six of the nine populations with recorded TSRs (Supplementary Table 1; Supplementary extended data Figure 2).

### Simulations of standing genetic variation vs. *de novo* mutations

Strong selection pressure exerted by herbicides leads to very rapid adaptation, but a major question is whether herbicide resistance evolves predominantly from standing genetic variation that was present already before the onset of herbicide selection or from *de novo* mutations that arose after herbicide selection began. In other words, are spontaneous TSR mutations sufficiently frequent for rapid resistance evolution and are the typical population dynamics in terms of effective population size and drift compatible with a reservoir of TSR mutations available for herbicides to act on, or not?

To answer this question, we first used equations from Hermisson & Pennings (2005)^36^ to derive expectations for the probability of adaptation (i.e., evolution of herbicide resistance via TSR mutations) and the likelihood that this adaptation is due to standing genetic variation. First, we calculated the probability of adaptation, assuming a mutation rate of 3.0 x 10^-8^ (ref. ^33^), a mutational target size of seven nucleotides, corresponding to the TSR mutations investigated here, and an onset of herbicide selection 30 generations ago. As mentioned above, diversity estimates of N_e_ integrate over a long period of time and past bottlenecks will reduce it, leading to estimates that are lower than the actual N_e_ before the bottlenecks^35^. Therefore, we considered both N_e_=42,000 (Figure 5a./c./e.-f., Supplementary Figure 11 a./c.-d.), which is the highest estimate from our field populations, and N_e_=84,000 (Figure 5 b./d./g.-h., Supplementary Figure 11 b./d.-e.).

**Figure 5.**
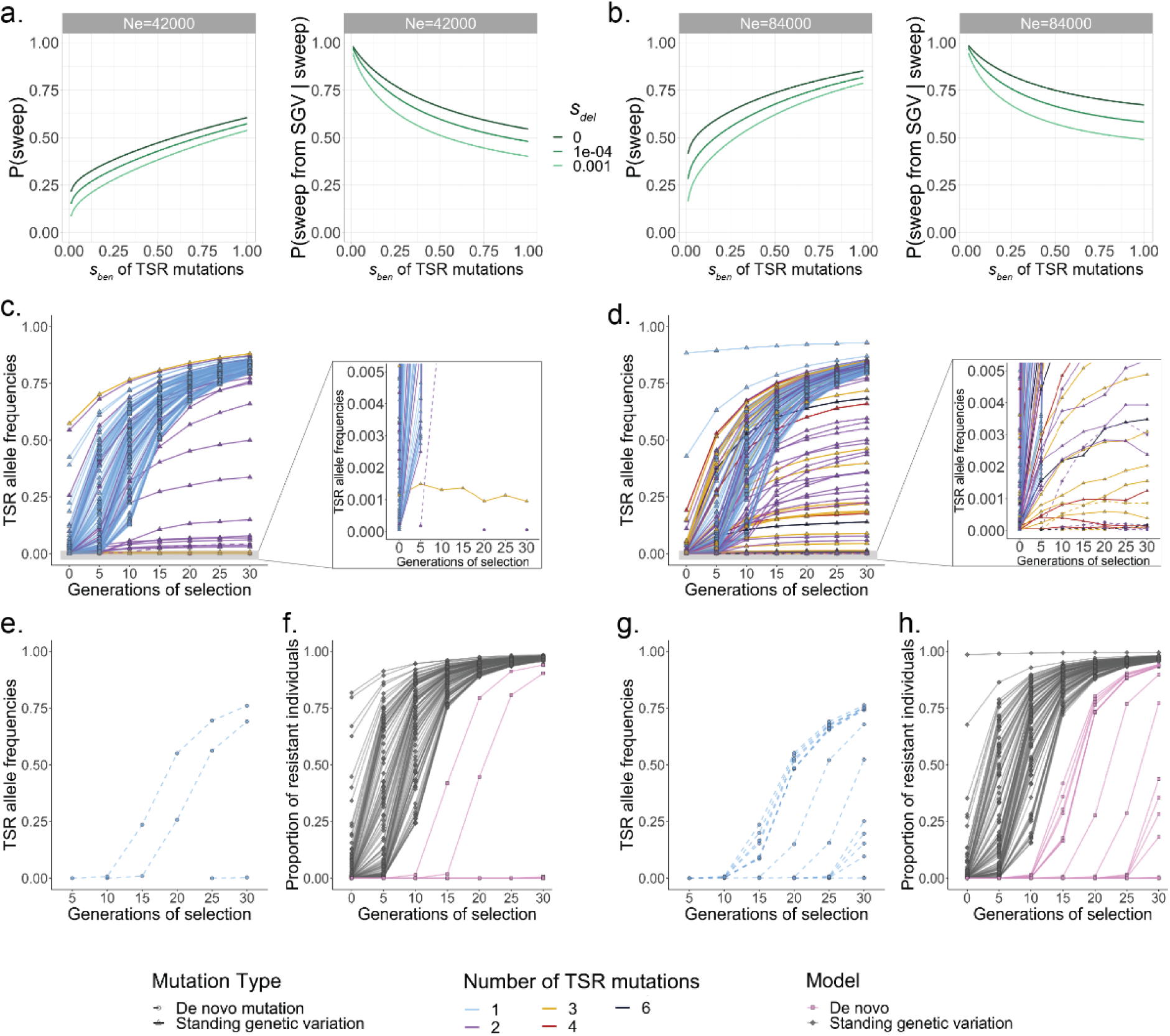
Simulations of adaptation through standing genetic variation vs *de novo* mutation. Simulations of adaptation probabilities via a selective sweep from beneficial target-site resistance (TSR) mutations in general (left panel) and originating from standing genetic variation (right panel) for effective population sizes of **a,** 42,000 and **b,** 84,000 individuals. The colours indicate different deleterious selection coefficients for the TSR mutations before the start of selection. **c-g,** Simulations of allele frequencies for mutations conferring TSR. From these simulations, the proportions of resistant individuals per field population can be inferred. The distribution of mutations is generated with a generic gene model that has the same number of exons and introns and ratio between coding and non-coding sequences as the ACCase gene. One hundred runs per model are shown for an effective population size of 42,000 individuals (**c, e, f**) and 84,000 individuals (**d, g, h**). Continuous lines represent mutations originating from standing genetic variation, *de novo* TSR mutations are shown with dashed lines. The colours indicate the total number of TSR mutations per population. **c, d,** Standing genetic variation model, with TSR mutations pre-existing in the populations before herbicide selection. Shown is the increase in TSR allele frequencies under herbicide selection of up to 30 generations, with one herbicide application per generation. The right panel shows a truncated y-axis at 0.005 TSR allele frequencies. **e, g,** *De novo* mutation model. Any TSR mutation that might have arisen before the start of selection has been lost again, so that no TSR mutations are present when the simulation starts. **f, h,** Proportion of resistant individuals across the hundred simulated populations under each of the two different models.

With N_e_=42,000, we observed that the probability of adaptation strongly depends on the beneficial selection coefficient of the TSR mutation during the herbicide selection phase (*s_ben_*). For strong positive selection (*s_ben_* >= 1), which we expect for herbicide application, the probability of adaptation is high (> 50%). Only for weakly beneficial mutations (*s_ben_* <= 0.01) it decreases below 20% (Figure 5a, left panel). With N_e_ = 84,000, the probability of adaptation increases to up to 80% due to both higher levels of standing genetic variation and a larger rate of *de novo* mutations (Figure 5b, left panel). The deleterious selection coefficient of TSR mutations before the onset of herbicide selection has only a minor influence on the probability of adaptation.

Next, we asked if adaptation to herbicide selection pressure predominantly occurs via standing genetic variation or *de novo* mutation^36^. We observe that fixation from standing genetic variation is more probable (>50%) for neutral or almost neutral mutations (*s_del_* < 1e^-4^) and has a probability larger than 40% even for deleterious mutations (*s_del_* = 1e^-3^) and the smaller population size (Figure 5a-b, right panels). The probability of adaptation from standing genetic variation generally increases with smaller *s_ben_* or larger *N_e_* because of the decreasing fixation probability of *de novo* mutations and the increasing levels of standing genetic variation, respectively^36^. These results suggest that herbicide resistance should occur predominantly, although not exclusively, via standing genetic variation.

However, the remarkable per-field diversity of TSR haplotypes of independent origin observed (Figure 4) prompted us to examine the speed of adaptation and the expected level of haplotype diversity in more detail through forward-in-time simulations with the software SLiM^16^. We simulated two different models: one in which resistant alleles were already present in the population before the start of herbicide selection (standing genetic variation model), and another one in which they emerged only after selection pressure was imposed (*de novo* mutation model) (Supplementary Figure 10). Under standing genetic variation, additional TSR mutations could of course arise after herbicide selection began. Both models assume that individuals with at least one TSR mutation have a 20 times higher chance of surviving the herbicide treatment than individuals without any TSR mutation. We applied a gamma distribution of deleterious mutations in exons^52^, although a model with exclusively neutral mutations gave similar results (Figure 5, Supplementary Figure 11). We analyzed the changes in allele frequencies of TSR mutations, the number of independent TSR mutations per population, as well as the proportion of resistant individuals under two different N_e_ scenarios, as explained above.

With N_e_ = 42,000, we found a higher number of independent TSR mutations per population for the standing genetic variation model, up to three mutations, than for the *de novo* model, not more than 1 mutation (Figure 5c, 5e, Supplementary Figure 11a, 11c). Since we had observed in most of our populations with TSR, namely 17 out of 27, three and more independent TSR mutations, the simulations support standing genetic variation as a main driver of herbicide resistance evolution in our system. Similarly, the standing genetic variation model showed a more rapid increase in the proportion of resistant individuals per population under selection than the *de novo* model (Figure 5f, 5h, Supplementary Figure 11d, 11f). This can be attributed to the fact that under the model of standing genetic variation including TSR mutations, TSR mutations could already have risen in frequency in the population before the onset of herbicide selection due to drift or because some of them are slightly beneficial even in the absence of herbicide application^41, 42^. Under selection pressure by herbicides, TSR mutations suddenly become highly beneficial and therefore increase very rapidly in frequency.

High effective population sizes favor rapid adaptation processes, because advantageous alleles are more likely to be immediately available and at higher frequencies^34, 36^. With N_e_ = 84,000, we found up to six independent TSR haplotypes in the standing genetic variation model (Figure 5d, Supplementary Figure 11b). In the single simulation that led to six independent TSR mutations, five were already present at the onset of selection, and a sixth arose later on (Figure 5d). Higher effective population sizes are more likely for our populations and are consistent with reports from farmers of heavy infestations in fields due to resistance problems. Large effective population sizes likely result from large census population sizes, thus maintaining genetic variation under herbicide selection and providing a large genetic pool for accumulation of resistance mutations. Factors that promote high census population sizes in specific years include climatic variables that lead to poor weed control conditions, seed dormancy, reduced tillage efficiency, large seed banks and crop rotations with high amounts of winter cereals^1, 53, 54^.

Simulations have shown that in the time it takes for a particular allele to become fixed in a population starting from standing genetic variation, the same mutation can arise *de novo*^36^. In the case of *de novo* mutations, our simulations reveal that there is a considerable risk that they are directly lost again through drift, since they are on average initially much rarer than mutations that are part of standing genetic variation (Figure 5c-d). And although we cannot exclude *de novo* mutations as a source of TSR alleles, they are characterized by a slow initial phase of adaptation (after 10-15 generations under selection) in our simulations, thus cannot compete with pre-existing mutations from standing genetic variation (Figure 5e, 5g, Supplementary Figure 11c, 11e). Therefore, the standing genetic variation model, with the presence of multiple alleles, as is typical for soft sweeps, is closer to what we observed in our experimental data. Furthermore, to estimate how many TSR alleles per generation are present as standing genetic variation in a field, we ran 100 simulations under neutrality. This revealed the emergence and loss due to random genetic drift in field populations before the start of herbicide selection by farmers. We could detect up to four TSR alleles at the same time (Supplementary Figure 12).

Previous studies documented rapid adaptation of *A. myosuroides* to herbicide applications within a few generations, sometimes as quickly as three or four generations^55, 56^, which is in agreement with anecdotal reports from farmers. The degree of herbicide resistance in a field is also closely correlated with the frequency of application^57^. This would be consistent with TSR mutations having been present already at low frequency before herbicides came into use, as shown through herbicide treatment of naive populations of *Lolium rigidum*^58^ or the analysis of herbarium samples of *A. myosuroides* collected before the advent of modern herbicides^59^. In the case of *L. rigidum*, the frequency of sulfometuron-methyl resistance in previously untreated populations was around 10^-4^ (ref. ^58^) while among 685 *A. myosuroides* herbarium specimens, one individual collected nearly hundred years before the introduction of herbicides carried the ACCase Ile-1781-Leu mutation^59^. We found TSR mutations in two of our three sensitive reference populations by deep amplicon sequencing (HerbiSeed standard, WHBM72 greenhouse standard APR/HA from Sep. 2014), although it is unknown when these populations were collected with respect to the relevant herbicides coming into broad use. On the other hand, in a study originally set out to empirically determine the *de novo* mutation rate of TSRs in a natural population of grain amaranth (*Amaranthus hypochondriacus*), not a single spontaneous resistant genotype was found among 70 million screened plants^60^. This would give 1.4 × 10^-8^ as an approximate upper bound of spontaneous mutations conferring resistance to a specific herbicide, which is in the range of spontaneous mutation rates that have been empirically measured in plants^61, 62^.

## Conclusions

Plants have evolved a remarkable number of mechanisms to protect themselves against damage and extinction from changing environmental conditions, including ones due to human activity. In particular, the outsized selection imposed by repeated application of herbicides had led to extraordinarily rapid evolutionary adaptation in many weed species.

While several TSR mutations incur fitness penalties in the absence of herbicide applications^43, 44^, some do not, and there is even at least one report of a TSR mutation being favorable independently of herbicide application^41, 42^. The extent of fitness costs before herbicide application began in a population will in turn affect whether TSR mutations can accumulate in a population that is not under herbicide selection. Our study demonstrates that rapid evolution of resistance in *A. myosuroides* populations primarily results from standing genetic variation, suggesting that the TSR mutations that are the focus of our investigation have limited fitness costs in the absence of herbicide treatment. In many cases, evolutionary adaptation in response to a change in the environment occurs via soft selective sweeps, as this allows a greater proportion of ancestral genetic diversity to be maintained^63^. The preservation of genetic diversity is particularly important in agricultural fields with highly variable conditions in terms of crop rotation, pest management and other field management measures, and can be crucial for weed populations to thrive under a range of different environmental conditions. Resistance apparently evolves in parallel in numerous fields with weed populations, occurring through different mechanisms, depending on which resistance pathways have pre-existing mutations.

Our examination of different scenarios for adaptation to herbicides indicates that, with the diversity of resistance mechanisms available, a large fraction of *A. myosuroides* populations is likely to have the genetic prerequisites not only for rapid evolution of resistance to currently used herbicide modes of action, but also to potential new future modes of action.

## Methods

### Reference genome sequencing, assembly and annotation

#### Plant selection and flow cytometry

A single plant from a sensitive German reference population provided by BASF was selected. All required tissues for all described reference related sequencing methods were collected from the same plant. We excluded the incidence of known TSR mutations on the Accase, ALS and psbA gene loci using Illumina amplicon sequencing. PCRs of the three target genes were performed (Extended data file 1), pooled and sequencing libraries were generated with a purified *Tn5* transposase as described in a previous study^64^. The library was spiked into an Illumina HiSeq 3000 lane. The resulting reads were checked for known TSR mutations causing herbicide resistance^4, 5^.

Leaf tissue from both the selected *A. myosuroides* plant and the reference standard *Secale cereale* cv. Daňkovské^19^ were simultaneously chopped with a razor blade in 250 μl of nuclei extraction buffer (CyStain PI Absolute P kit; P/N 05-5022). After the addition of 1 ml of staining solution (including 6 μl of propidium iodide (PI) and 3 μl of RNase from the same kit) the suspension was filtered through a 30 μm filter (CellTrics®; P/N 04-0042-2316). Five replicates of these samples were stored in darkness for 4 h at 4°C prior to flow cytometry analysis. PI-area was detected with a BD FACSMelody^TM^ Cell Sorter (BD Biosciences) equipped with a yellow-green laser (561 nm) and 613/18BP filtering. A total of 25,000 events were recorded per replicate, and the ratio of the mean PI-area values of each target sample and reference standard 2C peaks was used to estimate DNA content according to ref. ^65^ (mean = 3.53 Gb; sd = 0.0052 Gb; n = 5).

#### High-molecular-weight (HMW) DNA extraction, SMRTbell library preparation and PacBio sequencing

Prior to the HMW extraction the reference plant was kept for 48 hours in the dark to reduce the starch accumulation. We harvested ca. 30 g of young leaf material and ground it in liquid nitrogen. Nuclei isolation was performed according to the protocol of Workman et al. 2018 (ref. ^66^) with the following modifications: we used 16 reactions, each with 1 g input material in a 20 ml nuclear isolation buffer. The filtered cellular homogenate was centrifuged at 3500 x g, followed by 3x washes in nuclear isolation buffer. The isolated plant cell nuclei were resuspended in 60 μl Proteinase K (#19131, Qiagen). For HMW-DNA recovery, the Nanobind Plant Nuclei Big DNA Kit (SKU NB-900-801-01, Circulomics) was used. In total, we got approximately 80 μg of HMW-DNA, which was subjected to needle shearing once (FINE-JECT® 26Gx1’’ 0.45×25mm, LOT 14-13651). A 75-kb template library was prepared with the SMRTbell® Express Template Preparation Kit 2.0, and size-selected with the BluePippin system (SageScience) with 15-kb cutoff and a 0.75% agarose, 1-50kb cassette (BLF7510, Biozym) according to the manufacturer’s instructions (P/N 101-693-800-01, Pacific Biosciences, California, USA). The library was sequenced on a Sequel I system (Pacific Biosciences) using the Binding Kit 3.0. and MagBead loading. In total, we sequenced 18 SMRT cells of 10 hours and 8 SMRT cells of 20 hours movie time.

#### PCR-free library preparation and sequencing

The genomic DNA was fragmented to 350 bp size using a Covaris S2 Focused Ultrasonicator (Covaris) with the following settings: duty cycle 10%, intensity 5, 200 cycles and 45s treatment time. The library prep was performed according to the manufacturer’s instructions for the NxSeq® AmpFREE Low DNA Library Kit from Lucigen® (Cat No. 14000-2) with the addition of a large-cutoff bead-cleanup (0.6 : 1, bead:library ratio) after the adapter ligation, followed by the recommended standard bead-cleanup at the final purification step. The library was quantified with the Qubit Fluorometer (Invitrogen) and quality checked on a Bioanalyzer High Sensitivity Chip on an Agilent Bioanalyzer 2100 (Kit #5067-4626, Agilent Technologies). The library was sequenced on two lanes of a Illumina HiSeq 3000 system in paired end mode and 150bp read length.

#### RNA extraction and library preparation for short-read RNA Illumina sequencing

RNA was extracted from five different tissues (leaf, flower, anther, pollen, root) following the RNA extraction protocol of Yaffe *et al.* (2012; ref. ^67^). The remaining DNA was removed with DnaseI (#EN0521, Thermo Scientific) following manufacturer’s recommendations. The quality was checked with an RNA 6000 Nano Chip on an Agilent Bioanalyzer 2100 (Kit #5067-1511, Agilent Technologies). All RIN-scores were higher than 5.4.

For the library preparation, the NEBNext Ultra II Directional RNA Library Prep Kit for Illumina in combination with the Poly(A) mRNA Magnetic Isolation Module (#E7760, #E7490, NEB) was used. The heat fragmentation was performed for a duration of 9 min resulting in final library sizes of around 545 bp. All 5 libraries were equally pooled and sequenced on one lane of an Illumina HiSeq 3000 system in paired-end mode and 150 bp read-length.

#### RNA extraction and library preparation for long-read PacBio Iso-sequencing

We used the same five tissue samples as for the short-read sequencing. To ensure a high RNA quality for long-read sequencing, we performed the protocol of Acosta-Maspons *et al.* (2019, ref. ^68^), which is a CTAB based method for high-quality total RNA applications from different plant tissues. The remaining DNA was removed with the TURBO DNA-free Kit (Invitrogen), designed for optimal preservation of RNA during the DNase treatment. The quality check on the Agilent Bioanalyzer 2100 (Agilent Technologies) with a RNA Nano 6000 Chip resulted in RIN Scores higher than 7.6 for all tissues.

The IsoSeq libraries were prepared following the PacBio protocol for ‘Iso-Seq™ Express Template Preparation for Sequel® and Sequel II Systems’ (P/N 101-763-800 Version 02; October 2019, Pacific Biosciences, California, USA). The cDNA was amplified in 12 cycles and purified using the ‘standard’ workflow for samples primarily composed of transcripts centered ∼2 kb.

#### Hi-C library preparation

Hi-C libraries were prepared in a similar manner as described previously^69^ by Dovetail Genomics, LLC. Briefly, for each library, chromatin was fixed in place with formaldehyde in the nucleus and then extracted. Fixed chromatin was digested with DpnII, the 5’ overhangs filled in with biotinylated nucleotides, and then free blunt ends were ligated. After ligation, crosslinks were reversed and the DNA purified from protein. Purified DNA was treated to remove biotin that was not internal to ligated fragments. The DNA was then sheared to ∼350 bp mean fragment size and sequencing libraries were generated using NEBNextUltra enzymes and Illumina-compatible adapters. Biotin-containing fragments were isolated using streptavidin beads before PCR enrichment of each library. The libraries were sequenced on an Illumina HiSeq X.

#### Genome assembly

Genome assembly was done with the FALCON and FALCON-Unzip toolkit^20^ distributed with the ‘PacBio Assembly Tool Suite’ (falcon-kit 1.3.0; pypeflow 2.2.0; https://github.com/PacificBiosciences/pb-assembly). For the pre-assembly step, in which CLR subreads are aligned to each other for error correction, we opted for auto-calculating our own seed read length (’length_cutoff = -1’) with ‘genome_size = 3530000000’ and ‘seed_coverage = 40’ . Details of the FALCON assembly parameters used in this study are provided in the dedicated GitHub for this study (https://github.com/SonjaKersten/Herbicide_resistance_evolution_in_blackgrass_2022). Primary contigs were subjected to deduplication with purge_dups v1.0.0 (ref. ^21^) using cutoffs (5, 36, 60, 72, 120, 216). For scaffolding, deduplicated primary contigs and Hi-C library reads were used as input data for HiRise, a software pipeline designed specifically for using proximity ligation data to scaffold genome assemblies^22^. Dovetail Hi-C library sequences were aligned to the draft input assembly using bwa^70^. The separations of Dovetail Hi-C pairs mapped within draft scaffolds were analyzed by HiRise to produce a likelihood model for genomic distance between read pairs, and the model was used to identify and break putative misjoins, to score prospective joins, and make joins above a threshold.

#### Genome annotation

Transposable elements were annotated with the tool Extensive *de-novo* TE Annotator (EDTA) v1.9.7 (ref. ^28^). The protein-coding gene annotation pipeline involved merging three independent approaches: RNA-aided annotation, *ab initio* prediction and protein homology search. The first approach is based on both RNA-seq and Iso-seq data from five tissues (anther, flower, leaf, pollen and root). Pre-processing of Iso-seq data was carried out with PacBio® tools (https://github.com/PacificBiosciences/pbbioconda) that included in a first step the generation of CCS reads (minimum predicted accuracy 0.99 or q20) with ccs v5.0.0 and demultiplexing with lima v2.0.0. In a second step, poly-A trimming and concatemer removal were done at the sample level (i.e., separately for each tissue) while clustering was carried out for all tissues combined with functions from isoseq3 v3.4.0. Unique isoforms had a mean length of 2,210 bp. Iso-seq clusters were aligned to the *A. myosuroides* genome using GMAP v2017-11-15 using default parameters^71^, whereas RNA-seq datasets were first mapped to the *A. myosuroides* genome using Hisat2 (ref. ^72^) and subsequently assembled into transcripts by StringTie2 (ref. ^73^). All transcripts from Iso-seq and RNA-seq were combined using Cuffcompare^74^. Transdecoder v5.0.2 (https://github.com/TransDecoder) was then used to find potential open reading frames (ORFs) and to predict protein sequences. To further maximize sensitivity for capturing ORFs that may have functional significance, BLASTP^75^ (v2.6.0+, arguments -max_target_seqs 1 -evalue 1e-5) was used to compare potential protein sequences with the Uniprot database^76^. In the second approach, *ab initio* prediction was performed by BRAKER2 (ref. ^77^) using a model trained with RNA-seq data from *A. myosuroides*. For the third approach, consisting of homology prediction, the protein sequences from five closely related species (*Brachypodium distachyon*, *Oryza sativa*, *Setaria italica*, *Sorghum bicolor* and *Hordeum vulgare*) that belong to the same family were used as query sequences to search the reference genome using TBLASTN (-evalue 1e-5). These databases were downloaded from Plaza v4.5 (ref. ^78^) (https://bioinformatics.psb.ugent.be/plaza/). Regions mapped by these query sequences were subjected to Exonerate^79^ to generate putative transcripts. Finally, EvidenceModeler v1.1.1 (ref. ^80^) was used to integrate all of the above sources of evidence, and the Benchmarking Universal Single-Copy Orthologs (BUSCO; v4.0.4; embryophyta_odb10) set to assess the quality of annotation results^27^. Putative gene functions were identified using InterProScan^81^ with different databases, including PFAM, Gene3D, PANTHER, CDD, SUPERFAMILY, ProSite, GO. Meanwhile, functional annotation of these predicted genes was obtained by aligning the protein sequences of these genes against the sequences in public protein databases and the UniProt database using BLASTP (E-value <1 × 10^−5^).

#### Comparative genomics analyses

Analyses related to synonymous substitution rates (*K*_S_) were performed using the wgd package^82^. First, the paranome (entire collection of duplicated genes) was obtained with ‘wgd mcl’ using all-against-all BLASTP and MCL clustering. Then, the *K*_S_ distribution of *A. myosuroides* was calculated using ‘wgd ksd’ with default settings, MAFFT V7.453 (ref. ^83^) for multiple sequence alignment, and codeml from PAML package v4.4c (ref. ^84^) for maximum likelihood estimation of pairwise synonymous distances. Anchors or anchor pairs (duplicates located in collinear or syntenic regions of the genome) were obtained using i-ADHoRe^85^, employing the default settings in ‘wgd syn’. The list of anchors and their *K*_S_ values is available in Extended data File 1.

MCscan JCVI^86^ was used to do the analysis of syntenic relationships and depth ratio by providing the coding DNA sequences (CDS) and annotation file in gff3 format. TBtools was used to visualize the results via a circos plot^87^.

### Population study

#### Sample collection and DNA extraction

Seeds from 44 *A. myosuroides* populations from nine European countries were provided by BASF. The seeds were collected from farmers with suspected herbicide resistance in their fields against ACCase – and/or ALS-inhibiting herbicides. In addition, we included three sensitive reference populations (HerbiSeed standard, Broadbalk long-term experiment Rothamsted 2013, WHBM72 greenhouse standard APR/HA from Sep. 2014).

The seeds of all 47 populations were sown in vermiculite substrate and stratified in a 4°C climatic chamber for one week, and subsequently placed in the greenhouse at 23°C / 8 h daytime, 18°C / 16 h nighttime regime. After one week in the greenhouse, one plant per pot was transferred to standard substrate (Pikiererde Typ CL P, Cat.No EN12580, Einheitserde) for a total of 27 plants per population. We aimed to collect 8-weeks-old leaf tissue from 24 individuals per population, but due to insufficient germination in two populations, we were unable to collect material from two individuals and therefore finally obtained 1,126 samples for further processing. 300 mg of plant material was collected into a 2 ml screw cap tube filled with 4-5 porcelain beads and ground with a FastPrep tissue disruptor (MP Biomedicals). For DNA extraction, we used a lysis buffer consisting of 100 mM Tris (ph 8.0), 50 mM EDTA (ph 8.0), 500 mM NaCl, 1,3% SDS and 0.01 mg/ml RNase A. The DNA was precipitated with 5M potassium acetate, followed by two bead-cleanups for DNA purification (see detailed hands-on protocol: https://github.com/SonjaKersten/Herbicide_resistance_evolution_in_blackgrass_2022).

#### ddRAD library preparation and sequencing

The ddRAD libraries were prepared according to the published method for fresh samples of Lang *et al.* (2020; ref. ^88^). 200 ng input DNA per sample were digested with the two restriction enzymes EcoRI (#FD0274, Thermo Fisher Scientific) and Mph1103I (FD0734, Thermo Fisher Scientific), followed by double-stranded custom-adapter ligation. The custom-adapters contain different amounts of base-shifts to shift the sequencing of the restriction enzyme sites and prevent the sequencer from causing an error due to unique signalling. After the restriction enzyme digestion step and the adapter ligation, large cutoff bead-cleanups (0.6:1, bead:library ratio) with homemade magnetic beads (Sera-Mag SpeedBeads™, #65152105050450, GE Healthcare Life Sciences) in PEG/NaCL buffer^89^ were used to clean the samples from the buffers and remove large fragments above ∼600 bp length. We used a dual-indexing PCR to be able to multiplex up to six 96-well plates of samples. Thus, two pools of libraries were sufficient for all our samples. Since it is challenging to determine exact library concentrations, our strategy consisted of pooling all samples to the best of our abilities with the concentrations at hand and spike them into a Illumina HiSeq 3000 lane for about 5% of the total coverage. Afterwards, the library concentrations were re-calculated from the reads coverage output and re-pooled accordingly to achieve a more even coverage. Size selection was performed using a BluePippin system (SageScience) with a 1.5% agarose cassette, 250bp-1.5kb (#BDF1510, Biozym) for a size range of 300–500 bp. The library pools were quantified with the Qubit Fluorometer (Invitrogen) and quality checked on a Bioanalyzer High Sensitivity Chip on an Agilent Bioanalyzer 2100 (Kit #5067-4626, Agilent Technologies). A detailed hands-on protocol can be found here: https://github.com/SonjaKersten/Herbicide_resistance_evolution_in_blackgrass_2022.

First, each library pool was sequenced in-house on an Illumina HiSeq 3000 lane in paired-end mode and 150 bp read length to assess the performance and quality. Afterwards, both pools were submitted to CeGaT GmbH, Paul-Ehrlich-Straße 23, D-72076 Tübingen, and sequenced with an Illumina NovaSeq 6000 system on a S2 FlowCell with XP Lane Loading in paired-end mode and 150 bp read length. Total data output was 1.4 Tb, representing an average coverage of 22.6x read depth.

#### Alignment, SNP calling and SNP filtering

Demultiplexed raw reads were first trimmed for the base-shifts of the homemade adapters in the 5’ and 3’ fragment ends. Afterwards, all remaining adapter sequences and low quality bases were removed and only reads with a minimum read-length of 75 bp were kept using cutadapt v2.4 (ref. ^90^). The read quality was checked before and after trimming with FastQC v0.11.5 (Andrews S. (2010). FastQC: A quality control tool for high throughput sequence data: http://www.bioinformatics.babraham.ac.uk/projects/fastqc). Paired-end reads were first merged using Flash v1.2.11 (ref. ^91^), then the extended and the unmerged reads were independently aligned to the reference genome using bwa-mem v0.7.17-r1194-dirty (ref. ^70^). We used samtools v1.9 (ref. ^92^) to sort and index the bam-files and to finally combine the bamfiles of the extended and unmerged aligned reads per sample.

Variant calling was performed with the HaplotypeCaller function of GATK v4.1.3.0 (ref. ^93^). For joint genotyping, we broke the reference at N-stretches and generated an interval list with Picard’s v2.2.1 function ‘ScatterIntervalsByNs’ (http://broadinstitute.github.io/picard/). Next we generated a genomic database by using GATK v4.1.3.0 ‘GenomicsDBImport’, followed by joint genotyping with ‘GenotypeGVCFs’. A first missing data filter (--max-missing 0.3) was applied with VCFtools v0.1.15 (ref. ^94^) to the VCF outputs of all intervals to reduce the amount of not usable variants. Afterwards all interval VCFs were merged with Picard v2.2.1 ‘MergeVcfs’. The combined VCF was filtered following the recommendations of the RAD-Seq variant-calling pipeline ‘dDocent’ of Puritz *et al.* (2014; ref. ^95^). First, basic filters were applied with VCFtools v0.1.15 (--max-missing 0.5 --mac 3 --minQ 30 --minDP 3 --max-meanDP 35), followed by advanced filter options for RAD-Seq data with ‘vcffilter’ (ABHet > 0.25 & ABHet < 0.75 | ABHet < 0.01 & QD > 5 & MQ > 40 & MQRankSum > (0 - 5) & MQRankSum < 5 & ExcessHet < 30 & BaseQRankSum > (0 - 5) & BaseQRankSum < 5) (2012 Erik Garrison: https://github.com/vcflib/vcflib). We also filtered individuals with missing data more than 0.5, which removed four individuals from our dataset and we ended up with a total of 1,122 individuals. Lastly, we used a population specific variant filter, which allowed for 30% missing data, but every variant had to be called in at least 10 populations. Our final VCF for further analysis contained 109,924 informative SNPs.

#### Population statistics

A maximum likelihood (ML) phylogenetic tree was inferred with RAXML-NG v0.9.0 (ref. ^96^) to display the genetic relationship between the samples of our European dataset. We inferred a single ML-tree without bootstrapping using the model GTR+G+ASC_LEWIS of nucleotide evolution with ascertainment bias correction since we inferred it on RADSeq data. The annotation of the tree for the known TSR mutations was done based on the ALS and ACCase amplicons described below. For visualisation, we used the interactive Tree Of Life (iTOL) online tool^97^.

To assess the population structure of our European collection we ran a principal component analysis (PCA) with the R-package SNPrelate^98^ on 101,114 biallelic informative SNPs. To perform the admixture analysis on shared ancestry, we first pruned the dataset with PLINK v1.90b4.1 (ref. ^99^) for only biallelic SNPs. Admixture v1.3.0 (ref. ^100^) was run for up to 10 k groups, using a 10-fold cross-validation procedure to infer the right amount of k groups. TreeMix v1-13 (ref. ^101^) was run on the VCF filtered with PLINK as previously described in the admixture analysis. The transformation into the right input file format was done with STACKS v1.48 (--treemix)^102^. The tree was rooted with the most divergent outgroup population NL11330 (-root NL11330) and inferred in windows of 50 SNPs (-k 50) with 5 bootstrap replicates (-bootstrap 5). Since the treemix F3 statistic did not show significant migration, no migration events were added to the tree. FSTs were calculated with STACKS v1.48 (--fstats)^102^ and visualised with the R package ComplexHeatmap 2.0.0 (ref. ^103^).

Since we only covered about 1.1% of the entire genome with our ddRAD-Seq reads, we calculated the Watterson thetas θ_W_ and effective population sizes exclusively from the sequenced portion of our genome. Therefore, we used ANGSD v0.930 (ref. ^104^) on our previously generated bam-files and applied some basic filters (-uniqueOnly 1 -remove_bads 1 -only_proper_pairs 0 -trim 0 -C 50 -baq 1 -minMapQ 20), followed by calculation of the site-frequency spectra (SFS) (-doCounts 1 -GL 1 -doSaf 1) and Watterson thetas θ_W_ in sliding windows of 50,000 bp with a step size of 10,000 bp. The effective population size was calculated after the formula *N_e_ = θ_W_ / 4*μ* for a diploid organism. The mutation rate *μ* for the calculation was taken from the *Zea mays* literature^33^ as a genome-wide average of 3.0 x 10^-8^.

VCFtools v0.1.15 (ref. ^94^) was used to calculate the coverage (--depth) of the SNP markers and the observed homozygosity O(HOM) (--het). Using the number of sites N_SITES, the proportion of observed heterozygous sites can be calculated according to the formula (N_SITES - O(HOM)) / N_SITES.

### Bulked-segregant analysis for NTSR

#### Phenotyping

For phenotyping, 27 plants per population described in a preceding section were divided into two treatment and two control groups, following a specific tray design to minimize spatial growth effects. Treatment 1: Atlantis WG® (Bayer Crop Science) + Synergist Atlantis WG® (10 plants per population). Control 1: Only Synergist Atlantis WG® (3 plants per population). Treatment 2: Axial® 50 (Syngenta) + Synergist Hasten (10 plants per population). Control 2: Only Synergist Hasten (4 plants per population). All plants were sprayed 11 weeks after transplanting. Herbicides and synergists were applied with a lab sprayer (Schachtner), nozzle Teejet 8001 EVS and an air pressure of between 200 - 225 kPa. The sprayer was calibrated for a field application rate of 400 l/ha in 4 rounds of 3 independent replicates each (M = 396.4, SD = 7.53). Axial® 50 (50 g l^-1^ of pinoxaden + 12.5 g l^-1^ Cloquintocet-mexyl) was applied in combination with the synergist Hasten (716 g l^-1^ rapeseed oil ethyl and methyl esters, 179 g l^-1^ nonionic surfactants, ADAMA Deutschland GmbH). Atlantis WG® (29.2 g kg^-1^ of mesosulfuron and 5.6 g kg^-1^ of iodosulfuron) was used with the provided synergist (276,5 g l^-1^ sodium salt, fatty alcohol ether sulfate, Bayer Crop Science). Control plants were sprayed only with the synergists. Axial® 50 was applied at the recommended field rate of 1.2 l ha^-1^, Atlantis WG® at 800 g ha^-1^ and both synergists at 1 l ha^-1^. After 4 weeks all plants were scored according to the scheme in (Figure 3a). The score D1 represents completely dead plants and the A6 score represents plants without any growth reductions compared to the control plants of the respective population.

#### Bulked-segregant sample selection and library preparation

For the bulked-segregant experiment, only individuals phenotyped with the ACCase inhibitor Axial® 50 and that do not carry any of the kown TSR mutations at the ACCase locus were selected. Two bulks were formed, one sensitive bulk composed of individuals scored with D1 and one resistant bulk consisting of individuals with scores A5 and A6. Each bulk included a total of 61 individuals from different populations. A maximum of 4 individuals per population were chosen per bulk to minimize population structure effects (Figure 3a-b, Supplementary Figure 5).

For the selected individuals, previously extracted DNA was used and libraries were generated with a purified *Tn5* transposase as described in a previous study^64^. All libraries were pooled equally and sequenced on two lanes on an Illumina HiSeq 3000 system in paired-end mode and 150 bp read length. The mean coverage was 21.4x for the resistant pool and 21.5x for the sensitive pool.

#### Bulked-segregant alignment, SNP calling and analysis

We trimmed the Nextera adapters of the demultiplexed raw reads, removed low quality bases and kept only reads with a minimum read-length of 50 bp using cutadapt v2.4 (ref. ^90^). The quality was checked before and after trimming with FastQC v0.11.5 (Andrews S. (2010). FastQC: A quality control tool for high throughput sequence data: http://www.bioinformatics.babraham.ac.uk/projects/fastqc). We aligned the reads to the reference using bwa-mem v0.7.17-r1194-dirty (ref.^70^). We used samtools v1.9 (ref. ^92^) to sort and index the bam-files and to merge the files of each respective pool (sensitive/resistant). Samtools ‘mpileup’, combined with bcftools v1.9-15-g7afcbc9 function ‘bcftools call’ (--consensus-caller --variants-only) were used for variant calling on both pools^92, 105^. Basic SNP filters were applied with ‘bcftools filter’ (-g 10 -G 10 (DP4[0]+DP4[1])>1 & (DP4[2]+DP4[3])>1 & FORMAT/DP[]>5 MQ>40 INFO/DP<200) and ‘bcftools view’ for biallelic SNPs (-m 2 -M 2). In addition, we filtered SNPs overlapping TE intervals with the function ‘VariantFiltration’ from GATK v4.1.3.0 (ref. ^93^), providing a list of intervals with TEs to exclude. This resulted in 16 million informative SNPs. ‘Bcftools query’ was used to extract information for further downstream analysis (-f ‘%CHROM\t%POS\t%REF\t%ALT{0}\t%DP[\t%AD\t%DP]\n’).

The genome-wide scan for NTSR was performed with BayPass v2.2 (refs. ^39, 40^). The genotype file specified the reference and alternative allele counts of each pool, the haploid poolsize was 122 per pool, the initial delta of the y_ij_ proposal distribution was set to 24 (-d=yij 24) and both the contrastfile and the covariate file contained 1 and -1 for the resistant and sensitive pool, respectively. The summary file of the contrast statistic was loaded in R (ref. ^106^) and the Manhattan plot was generated with a custom script. A Bonferroni-corrected significance threshold (0.05) of *-log_10_(3.125e^-09^) = 8.5* for 16 million SNPs was used. The annotation of genes within 50 kb downstream or 50 kb upstream of the highest 100 SNP associations is reported in Extended data File 1.

### TSR amplicons

#### ALS and ACCase PacBio amplicon preparation and sequencing

To generate ALS and ACCase amplicons for long-read PacBio sequencing we used the same DNA from the European collection described in a preceding section. Before PCR amplification DNA was normalized to 10 ng/μl. Then, 30 ng (ALS) and 50 ng (ACCase) total input DNA was used for the PCR Master Mix reaction (1 μl P5 indexing primer (5 μM), 1 μl P7 indexing primer (5 μM), 4 μl of 5x Prime STAR buffer, 1.6 μl dNTPs, 0.4 μl Prime STAR polymerase (Takara, R050B), filled up to 20 μl with water). The indexing PCR program for ALS was a 2-step PCR with 10 seconds of denaturation at 98°C and 210 seconds of annealing and extension at 68°C for 28 cycles, followed by a final extension for 10 min at 72°C. For ACCase, the annealing and extension step was elongated to 660 seconds. Amplicons were then pooled equally per gene and bead-cleaned. In the case of the 13.2 kb amplicon from ACCase, we added a BluePippin (SageScience) size selection to remove any remaining fragments below 10 kb. PacBio libraries were created according to the following PacBio amplicon protocol (part number 101-791-800 version 02 (April 2020)) and SMRT cells were loaded on a PacBio Sequel® I system with Binding Kit and Internal Ctrl Kit 3.0 (part number 101-461-600 version 10; October 2019). An extended hands-on protocol can be found online: https://github.com/SonjaKersten/Herbicide_resistance_evolution_in_blackgrass_2022.

#### PacBio amplicons analysis

Most steps were carried out with tools developed by PacBio® (https://github.com/PacificBiosciences/pbbioconda). First, CCS reads were generated with ccs v6.0.0 (minimum predicted accuracy 0.99 or q20). Then, demultiplexing was carried out with lima v1.11.0 (with parameters ‘--ccs --different --peek-guess --guess 80 --min-ref-span 0.875 --min-scoring-regions 2 --min-length 13000 --max-input-length 14000’ for ACCase while for ALS similar parameters were used except for ‘--min-length 3200 --max-input-length 4200’). Next, pbaa cluster (v1.0.0) was run with default parameters followed by a series of amplicon-specific filtering steps.

For ACCase, a minimum of 25 CCS reads per sample were required, and only “passed clusters” were further considered for analysis. Samples with either 0 or more than 2 clusters were discarded. In samples in which a single cluster was identified (i.e., homozygous individuals for this locus), both haplotypes were assigned the same cluster sequence. In samples in which two different clusters (haplotypes) were identified, the difference between their respective frequencies had to be <= 0.50, otherwise the sample was discarded.

In the case of ALS, PCR amplification was suboptimal. That is, our primers seem to preferentially amplify certain haplotypes over others in individuals heterozygous for this locus. We presumed this was due to various structural variations downstream of the gene between major haplotypes (Supplementary Figure 9). Therefore, to be able to analyse haplotype diversity of this locus, we employed less strict filtering steps than for ACCase. For ALS, a minimum of 25 CCS reads per sample were required, while both ‘passed clusters’ and originally ‘failed clusters’ (mostly due to low frequency) were re-evaluated. First, only samples with cluster diversity <= 0.40 and cluster quality >= 0.7 were kept. In samples in which a single cluster was identified (i.e., homozygous individuals for this locus), its frequency had to be >= 0.98 to then assign the same cluster sequence to both haplotypes. In samples in which two different clusters (haplotypes) were identified, the difference between their respective frequencies was allowed to be <= 0.85, otherwise the sample was discarded. In the few samples in which three or more different clusters (haplotypes) were identified, the sum of the frequencies of the two main clusters had to be >= 0.96, and their difference <= 0.85 to be considered for downstream analyses.

#### Haplotype networks, haplotype trees and Haplotype PCA

To annotate the clusters generated with pbaa with TSR metadata information, the single cluster fasta files representing two alleles per individual were first converted to fastq files using ‘Fasta_to_fastq’ (https://github.com/ekg/fasta-to-fastq). The resulting fastq files were aligned to the ACCase reference using minimap2 v2.15-r913-dirty (ref. ^107^), followed by sorting and indexing of the output bam files with samtools v1.9 (ref. ^92^). Read groups were assigned with the Picard function ‘AddOrReplaceReadGroups’ (RGID=$SAMPLE RGLB=ccs RGPL=pacbio RGPU=unit1 RGSM=$SAMPLE) (http://broadinstitute.github.io/picard/), followed by variant calling using GATK v4.1.3.0 (ref. ^93^) with functions ‘HaplotypeCaller’ (-R $REF --min-pruning 0 -ERC GVCF) and ‘GenotypeGVCFs’ with default settings. Variant annotation in the resulting VCF was performed with SnpEff v4.3t (ref. ^108^). The VCF was loaded in R to extract the TSR information and annotate the haplotype networks, trees and PCA with custom R scripts.

For the multiple alignments per population, we first combined all respective individual fasta files of the pbaa clusters into a single fasta file and then aligned them using MAFFT v7.407 (--thread 20 --threadtb 10 --threadit 10 --reorder --maxiterate 1000 --retree 1 --genafpair)^83^. We used PGDSpider v2.1.1.5 (ref. ^109^) to transfer the multiple alignment fasta file into a Nexus-formatted file. Minimum spanning networks were inferred and visualized with POPART v.1.7 (ref. ^110^). Per population haplotype trees were inferred with RAXML-NG v0.9.0 (ref. ^96^) from the multiple sequence alignment files. ‘Tree search’ was performed with 20 distinct starting trees and bootstrapping analysis with the model GTR+G and 10,000 bootstrap replicates. Tree visualisation was done in R with ggtree v1.16.6 (ref. ^111^). The packages treeio v1.8.2 (ref. ^112^) and tibble v3.0.4 (https://github.com/tidyverse/tibble/) were used to add the TSR metadata information to the tree object. The branch length and node support values were extracted from Felsenstein’s bootstrap proportions (FBP) output files. The haplotype PCAs were performed using the R package SNPrelate^98^ on the previously generated VCFs for ALS and ACCase and visualized using ggplot2 (ref. ^113^).

#### Identification of multiple copies of ALS

Using the ALS GenBank sequence of *A. myosuroides* AJ437300.2 (ref. ^48^) as a query, BLASTN v2.2.29+ (ref. ^114^) retrieved three hits in chromosome 1 of our assembly. These loci corresponded to three gene models annotated as the largest subunit of ALS: model.Chr1.12329 (identity = 1921/1923 bp; 99.8%; hereafter ALS1), model.Chr1.11275 (identity = 1820/1915 bp; 95.0%; hereafter ALS2) and model.Chr1.11288 (identity = 1818/1915; 94.9%; hereafter ALS3).

To better characterize the relationship between these putative copies of the ALS gene, we analyzed synonymous substitution rates (*K*_S_) and Iso-seq full-transcripts. *K*_S_ values between paralogs ALS1-ALS2 and paralogs ALS1-ALS3 were 0.1526 and 0.1645, respectively, while between paralogs ALS2-ALS3 was 0.0281. Although all *K*_S_ values between these paralogs were low (< 0.5), they are not present in our list of anchor pairs from the comparative genomics analysis (Extended data file 1) for not being located among the collinear regions identified by i-ADHoRe^85^.

For the analysis of Iso-seq data, we first generated very high quality reads (this time with a minimum predicted accuracy 0.999 or q30) per tissue up until the poly-A trimming and concatemer removal step with isoseq3 v3.4.0 as described before for genome annotation. Next, we combined the q30 Iso-seq transcripts from all tissues, and extracted only those that matched the following internal ALS sequences conserved among the three loci: ‘CGCGCTACCTGCCCGCCTC’, ‘GTCTCCGCGCTCGCCGATGCT, ‘GTCCAAGATTGTGCACAT’ and ‘GAGTGAAGTCCGTGCAGCAATC’. We obtained 343 Iso-seq q30 full-length transcripts, and it is worth mentioning that different internal ALS sequences yield near identical numbers of transcripts. Since Iso-seq q30 reads have heterogeneous lengths, we used cutadapt v3.6.8 (ref. ^90^) to trim all reads at the 5’ and 3’ borders (-a CTTATTAATCA -g CCACAGCCGTCGC) of the CDS to make them all the same length. Finally, clustering with pbaa v1.0.0 (--min-read-qv 30) resulted in only three clusters with 143 reads corresponding to ALS1, 100 reads to ALS2 and 100 reads to ALS3. Representative full-length Iso-seq reads (average read quality of q93) from each cluster were used for Supplementary Figure 8. Therefore, all ALS gene models express full-length transcripts. Taking together *K*_S_ values and Iso-seq data, we could only conclude that ALS1 is clearly distinct from ALS2 and ALS3, but we could not distinguish whether ALS2 and ALS3 are two distinct loci or two alleles of the same locus.

### Model simulations

Using equations 8, 11, 14, 18 and 20 from Hermisson and Pennings (2005; ref. ^34^), we first modeled the general probability of adaptation through a sweep and then specifically from standing genetic variation. We set the population size to 42,000 individuals, which is the highest possible N_e_ from the populations characterized with RAD-Seq data. Since diversity estimates of N_e_ integrate over a long period of time and past bottlenecks will reduce it, leading to estimates that are lower than the actual N_e_ before the bottlenecks^35^, we additionally simulated the doubled effective population size of 84,000 individuals. As maize is a diploid grass with a similar genome size to *A. myosuroides*, we adopted the mutation rate 3.0 x 10^-8^ (ref. ^33^). Both target site resistance genes in our study contain seven well described SNP positions that cause resistance^4–7^, therefore we set the mutational target size to seven. Before selection, we assumed three different selection coefficients for those mutations: 0, 1e-04, 0.001. Under selection, those TSR positions were beneficial in a range from 0 to 1 (Figure 5a.-b., x-axes). The number of generations of selection was 30.

#### Standing genetic variation model vs. *de novo* model

Forward simulations were executed on a computing cluster with SLiM v3.4 (ref. ^16^) using SLiMGui v3.4 for model development. We used the ACCase locus as a template for all our simulations. Since we sequenced 585 bp upstream and 364 bp downstream of the gene, we defined the length of our simulated genomic element as 13,199 bp with TSR mutations at the following positions: 11052 (Ile1781), 11706 (Trp1999), 11790 (Trp2027), 11832 (Ile2041), 11943 (Asp2078), 11973 (Cys2088), 11997 (Gly2096). Since *A. myosuroides* is an annual grass, all models were built as Wright-Fisher models with non-overlapping generations and standard Wright-Fisher model assumptions (http://benhaller.com/slim/SLiM_Manual.pdf, p.35/36). As described above, we set the population size to 42,000 and 48,000 individuals. Both the mutation rate (3.0 x 10^-8^) (ref. ^33^) and genome-wide average recombination rate (7.4 x 10^-9^) (ref. ^115^) were adopted from maize. We implemented a burn-in period of 10 x N_e_ generations to generate the initial genetic diversity and, since this is a computationally intensive process, saved that state of the population for runs that did not satisfy the conditions at the checkpoint 10,000 generations after (see below). For the same reason, we scaled our models down by a factor of 5.

For the standing-genetic-variation model, we ran the model until generation 10 x N_e_ + 10,000 generations. This generation was a checkpoint for the presence of at least one individual in the total population carrying at least one of the TSR mutations in a heterozygous state. If this was not the case, the simulation returned to the saved state at 10 x N_e_ generations and generated random mutations until the checkpoint was reached again. This loop ran until the conditions of the checkpoint were satisfied. We defined three genomic element types: exon, intron and non-coding region. For introns and non-coding regions, all mutations were considered to be neutral with a selection coefficient of 0 and a dominant coefficient of 0.5. In exons, a ratio of 0.25/0.75 (neutral/deleterious) mutations was assumed, with selection coefficients (s) for deleterious mutations drawn from a gamma distribution with E[s] = -0.000154 and a shape parameter of 0.245 ^52^. After the herbicide selection started, mutations at the specified TSR positions became highly beneficial and dominant, with a selection coefficient of 1.0 and a dominance coefficient of 1.0. In practice, a herbicide is usually applied in the field once or twice each year. Since in *A. myosuroides* one generation time corresponds to about one year, we simulated one selection event per generation.

Foster et al. 1993 (ref. ^116^) specifically reported an ACCase inhibiting herbicide efficiency rate of 95-97%. However, it is likely that some *A. myosuroides* plants without TSR mutations will later emerge and thus escape the lethal effect of herbicide treatment contributing to the genetic diversity in the field. Therefore, in our simulations, we assume a fitness of 10% for individuals that do not carry a TSR mutation to account for plants that escaped herbicide treatment or germinated at a later time point. The selection pressure was applied at the end of every generation for a total of 30 generations. Only survivor individuals could reproduce and contribute to the next generation. To assess the robustness of our simulation, we also ran the model without exons and introns, thus considering all mutations to be neutral with a selection coefficient of 0 and a dominance coefficient of 0.5 until the time point of selection (Supplementary Figure 11).

In the *de novo* mutation model, we assumed that none of the TSR mutations were present at the starting point of selection. Therefore, the only difference in the model is the condition for the loop at the checkpoint, which we reversed: no TSR mutation must be present; if there is, the simulation reverts to the saved state at 10 x N_e_ generations. A schematic representation of both models can be found in Supplementary Figure 10. For each model, 100 independent runs were performed. Allele frequencies and proportion of resistant individuals for 7 different time points (before selection, 5, 10, 15, 20, 25 and 30 generations after start of selection) were written to a log file and plotted in R with the ggplot2 R-package^113^.

#### TSR occurrence

We created a model to examine how often a TSR mutation occurs on our simulated ACCase locus and how long it remains in the population before either being lost due to genetic drift or increase in frequency toward fixation. This allows us to quantify how often resistance mutations are present as standing genetic variation in a field population before herbicide selection starts. To this end, we built a neutral model and ran it for 1,000 generations. The general parameters are the same as described above: 42,000 and 84,000 individuals, mutation rate 3.0 x 10^-8^ (maize)^33^, recombination rate 7.4 x 10^-9^ (maize)^115^. Mutations were modeled using an intron/exon gene model for the ACCase locus as described in the previous section. After each generation, we output the number of TSR mutations at the predetermined TSR positions in the population. We performed 100 independent simulation runs per N_e_. Detailed scripts for all simulations can be found at https://github.com/SonjaKersten/Herbicide_resistance_evolution_in_blackgrass_2022.

### Data manipulation and plotting

The visualisation of our data was done with R version v3.6.1 (ref. ^106^) and RStudio v1.1.453 (RStudio Team (2018). RStudio: Integrated Development for R. RStudio, Inc., Boston, MA. http://www.rstudio.com). All R packages and versions used for general data manipulation and visualisation can be found in Supplementary Table 2.

## Data availability

Raw data for the genome assembly of *Alopecurus myosuroides* such as PacBio CLRs, Illumina PCR-free, Hi-C, Iso-Seq and Illumina RNA raw sequencing data can be accessed in the European Nucleotide Archive (ENA; https://www.ebi.ac.uk/ena/browser/home) under the project accession number PRJEB49257. Raw ddRAD-seq data for the population study, Illumina DNA-seq data for the bulked-segregant analysis, and PacBio CCS q20 reads can be downloaded from the ENA project accession number PRJEB49288. Genome assembly, annotation files, the SNP matrices for both the ddRAD-seq and bulked-segregant experiments, and the fasta files with the haplotypes of ACCase and ALS can be found on: https://keeper.mpdl.mpg.de/d/520501d08acc4bd887f7

## Acknowledgments

We thank Andreas Landes (BASF) for the European blackgrass accessions, Daniel Hewitt and John Cussans (NIAB) for the sensitive reference seeds of the Broadbalk long-term experiment (Rothamsted, 2013), Wei Yuan for recommendations for bulked-segregant analysis, Johannes Herrmann for helpful discussions on population structure and haplotype networks, Derek Lundberg for advice on amplicon sequencing, Frank Chan for facilitating the use of the flow cytometer, Marek Kučka for providing purified *Tn5* transposase, Dovetail Genomics for Hi-C library preparation and genome scaffolding, and Rudi Antonise (KeyGene, Wageningen) for the genotyping-by-sequencing service of herbicide sensitive populations that allowed selection of the reference population. S.K. was supported by a stipend from the Landesgraduiertenförderung (LGFG) of the State of Baden-Württemberg. F.A.R. was supported by a Human Frontiers Science Program (HFSP) Long-Term Fellowship (LT000819/2018-L). The majority of funding was provided by BASF and the Max Planck Society.

## Author contributions

S.K., D.W. and F.A.R. conceived and designed the project. F.A.R. assembled the genome. J.C. annotated the genome and performed comparative genomic analysis with the supervision of Y.V.d.P. S.K. and C.D.H. performed genetic simulations. S.K. and F.A.R. performed amplicons analyses. Y.V. contributed to GWAS. C.L. and S.K. carried out genome sequencing. T.H. helped with greenhouse experiments. P.L. helped with ddRAD-seq. U.L. genotyped the reference individual for known TSRs. F.A.R. and I.H. performed flow cytometry. S.K. carried out all other experiments and most of the other data analyses with support from F.A.R. J.L. and A.P. provided genetic material. S.K. wrote the first draft of the manuscript. S.K., D.W. and F.A.R. edited the final manuscript with input from all authors. Y.V.d.P, K.S. and D.W. supervised research. F.A.R. administered the project.

## Supplementary Figures

**Supplementary Figure 1.**
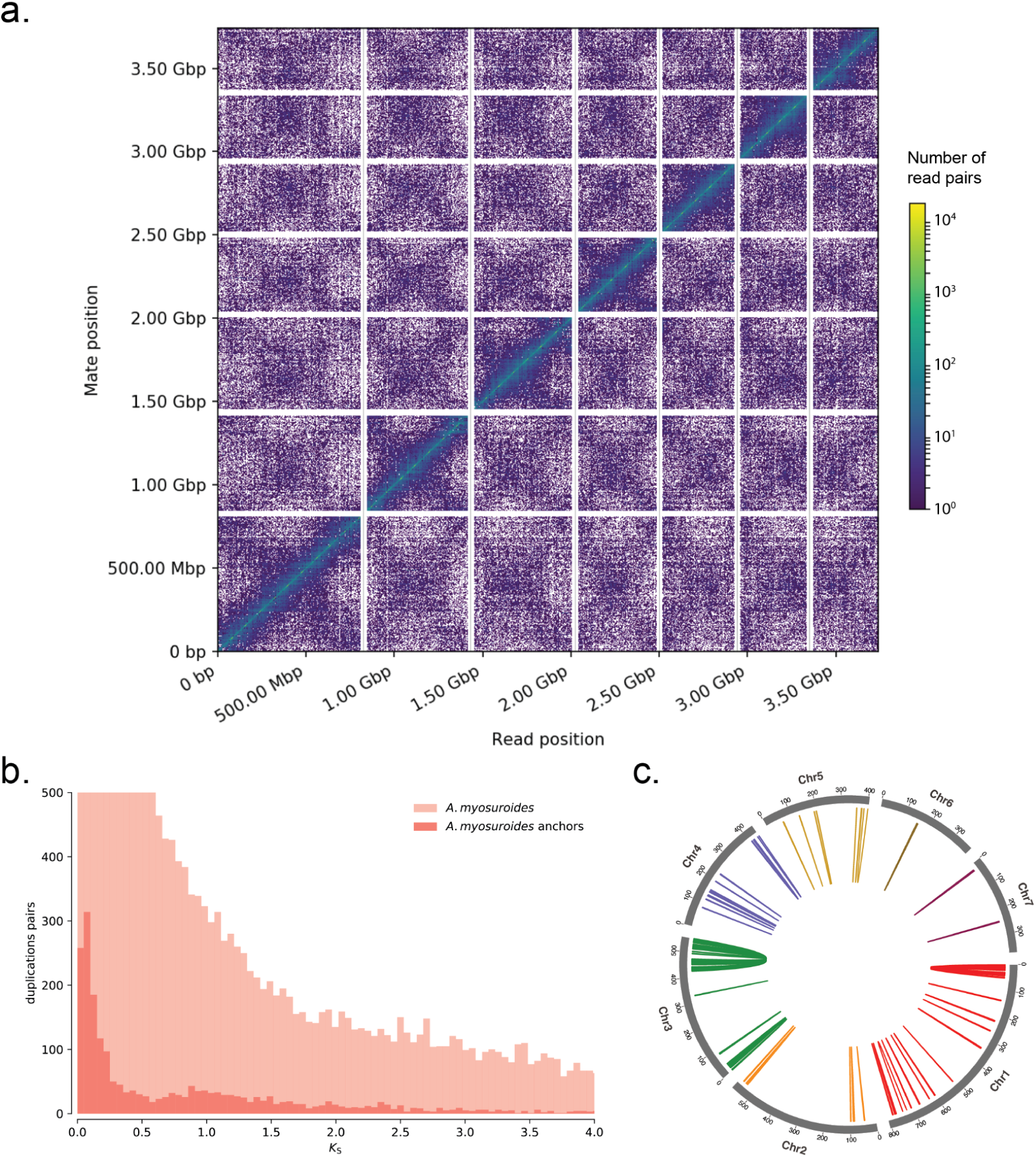
Genome scaffolding and analysis of anchors. **a,** Link density plot of the mapping positions of the first (x-axis) and second read (y-read) in the read pair, grouped into bins. The colour of each square indicates the number of read pairs in that bin. Scaffolds < 1 Mb are excluded. **b,** *K*_S_ distributions for all paralogs within the *A. myosuroides* (light colour) and for the paralogs retained in collinear regions, also known as anchors (dark colour). **c,** Circos plot of the *A. myosuroides* genome, with coloured lines connecting anchor pairs (genes in the collinear regions) with *K*_S_ < 0.5 (Extended data File 1).

**Supplementary Figure 2.**
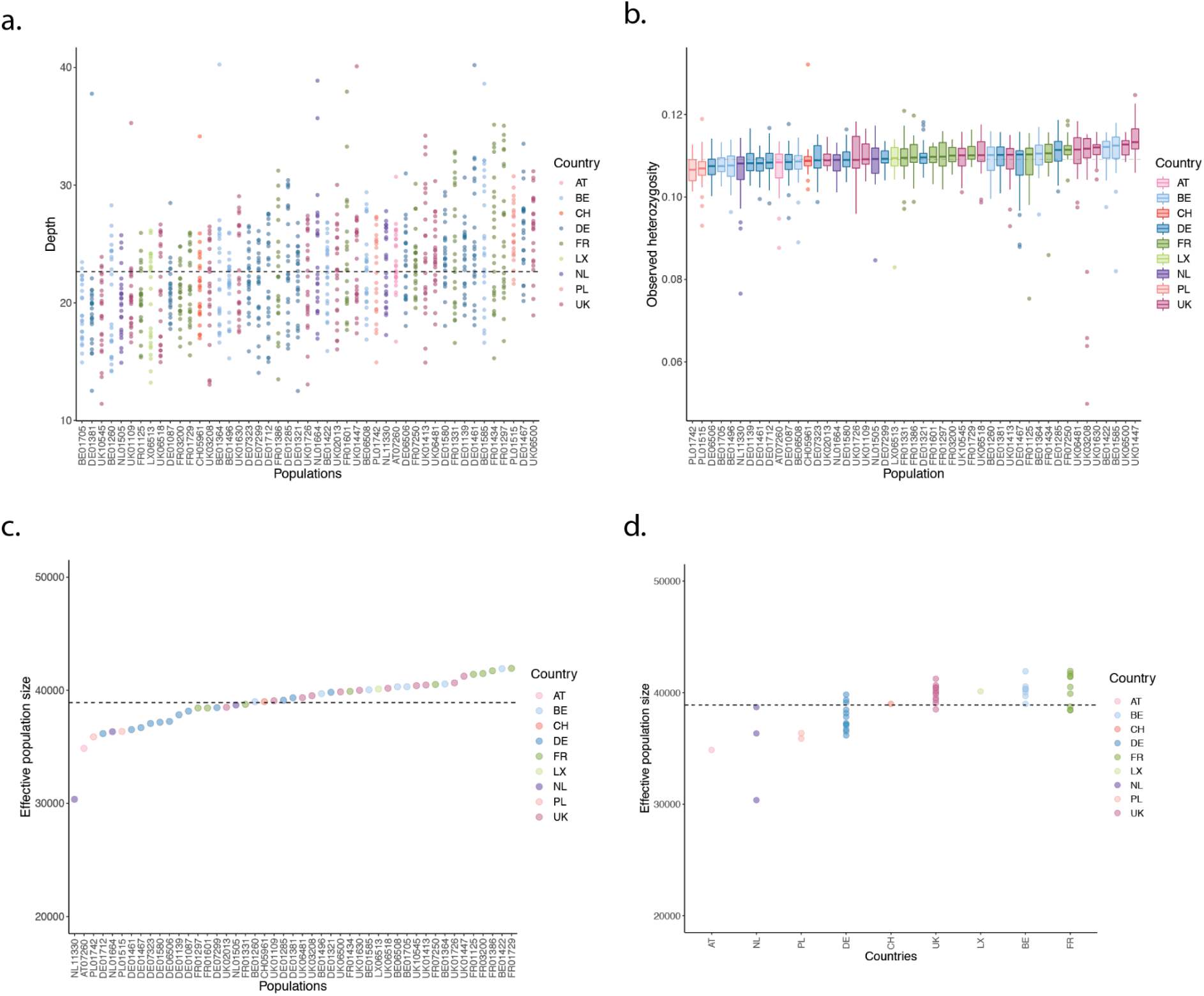
Basic statistics of the ddRAD-Seq dataset and diversity metrics. Colours reflect country-specific origin of the populations. **a,** Sequencing depth. **b,** Observed SNP heterozygosity. **c,** Effective population sizes. Mean= 38,912 individuals (dashed line) **d,** Effective population sizes ordered by countries. The Tukey HSD test showed a significant difference between the mean effective population size of DE and UK (p<0.03), DE and BE (p<0.03), DE and FR (p<0.01). Austria[AT], Belgium [BE], Switzerland [CH], Germany [DE], France [FR], Luxembourg [LX], Netherlands [NL], Poland [PL], United Kingdom [UK].

**Supplementary Figure 3.**
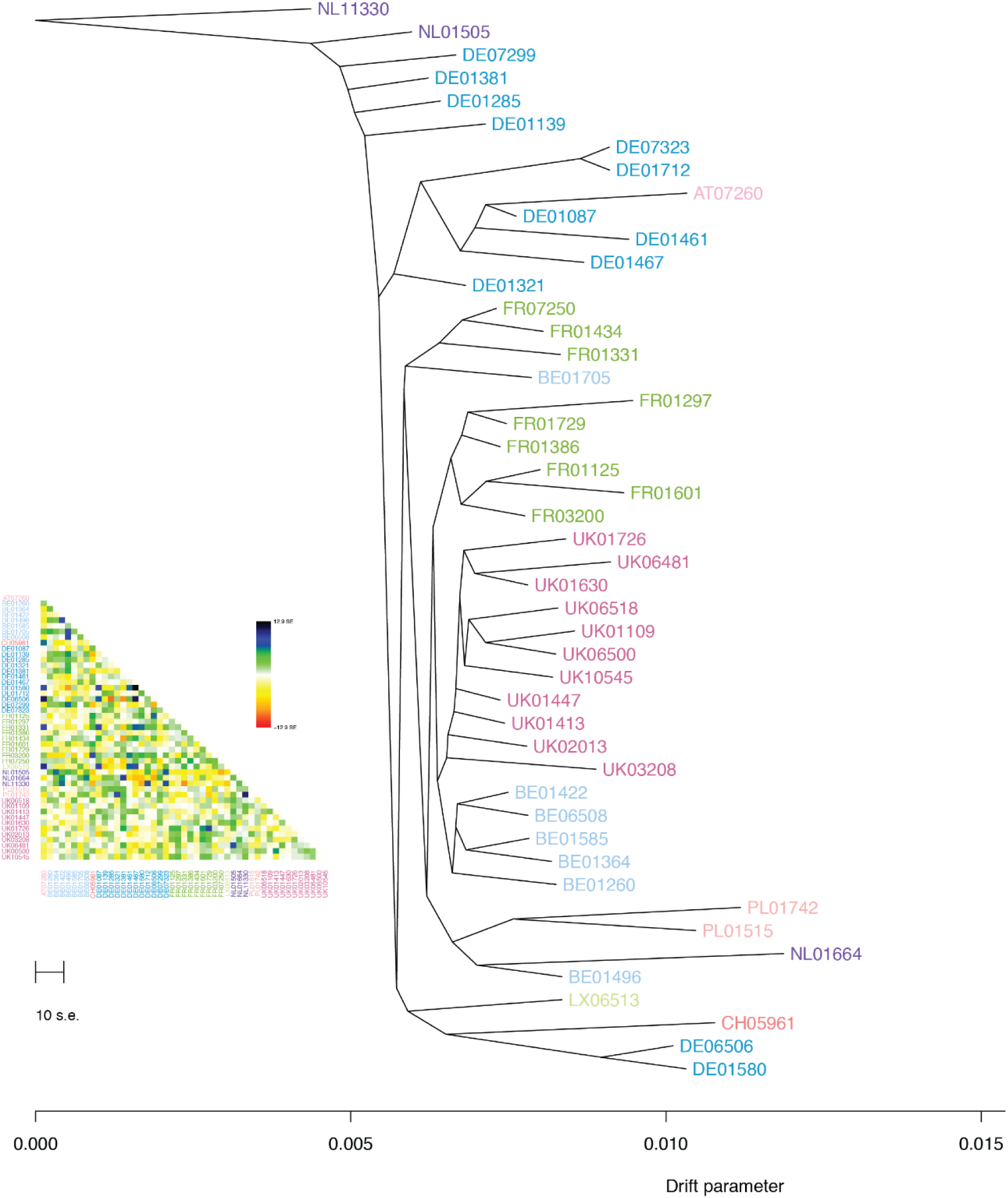
Phylogenetic population tree with residual plot. Colours reflect the country-specific origin of the populations. Austria (AT), Belgium (BE), Switzerland (CH), Germany (DE), France (FR), Luxembourg (LX), Netherlands (NL), Poland (PL), United Kingdom (UK).

**Supplementary Figure 4.**
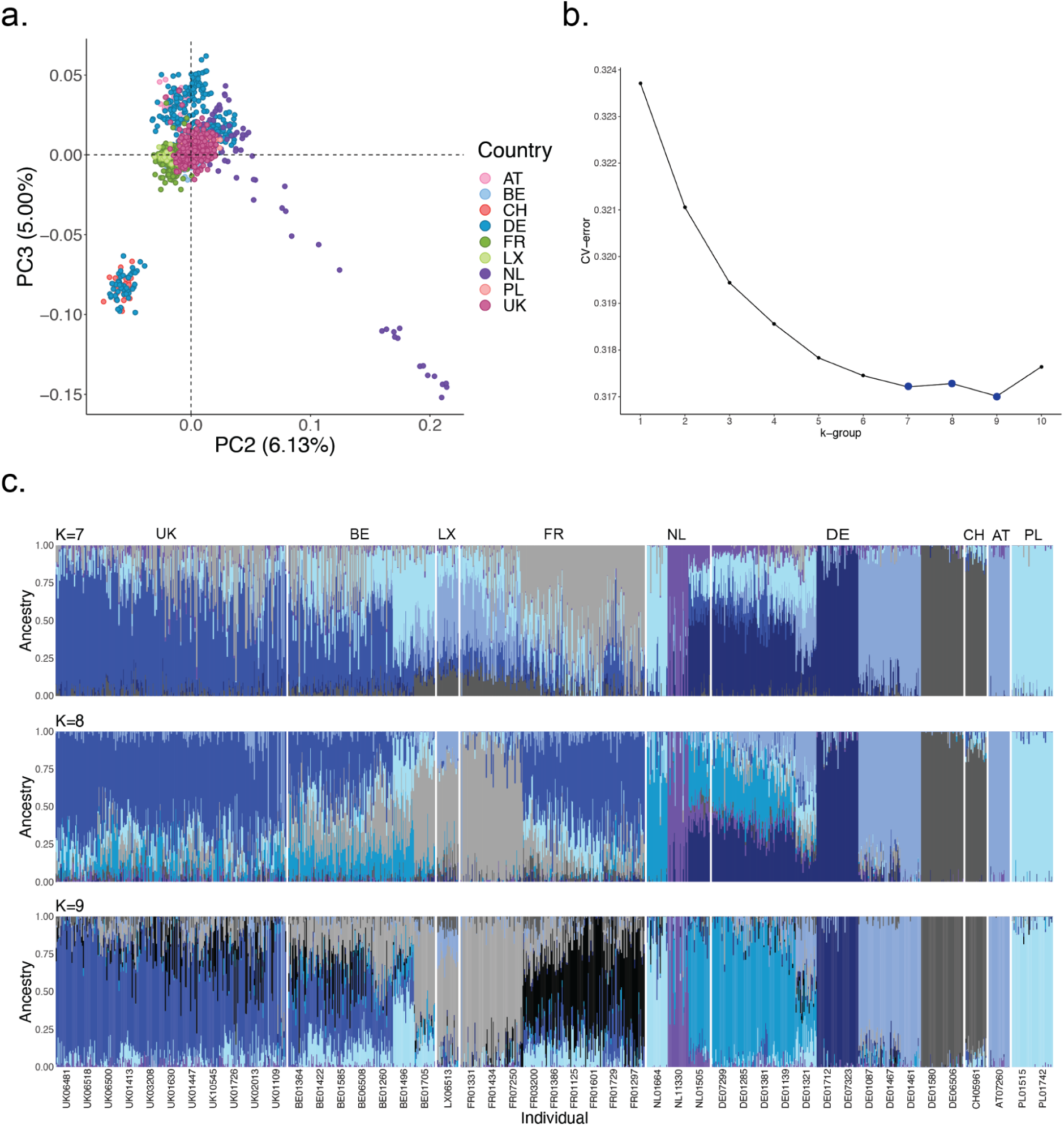
Population structure analysis. **a,** Second and third eigenvectors of the principal component analysis (PCA). The genetic variance of the second and third component is shown in brackets. Colours reflect country-specific origin of the populations. **b,** Cross validation error as a function of K of the admixture analysis. c, Admixture proportions with ancestry groups of K=9, K=7 and K=8. Austria (AT), Belgium (BE), Switzerland (CH), Germany (DE), France (FR), Luxembourg (LX), Netherlands (NL), Poland (PL), United Kingdom (UK).

**Supplementary Figure 5.**
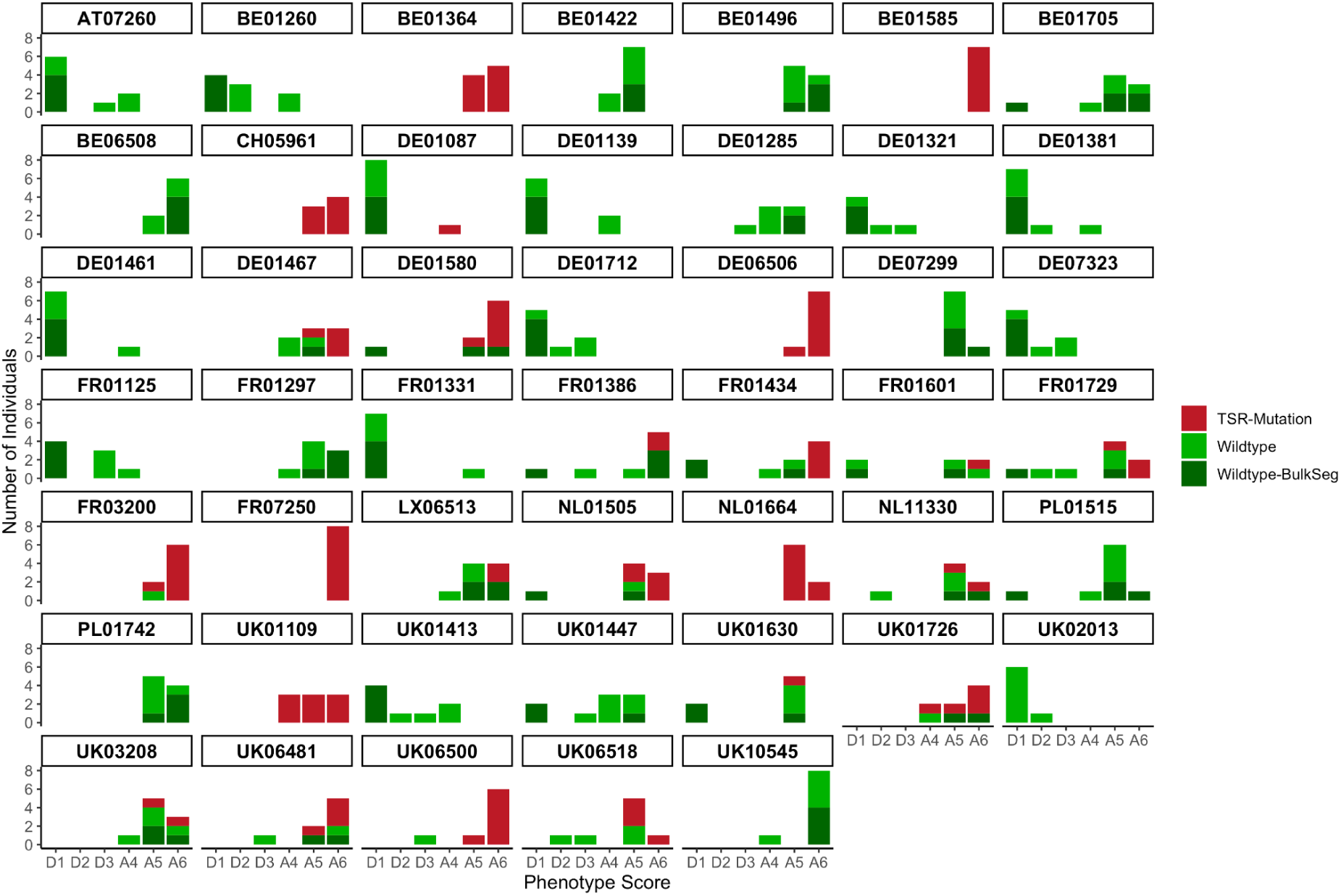
Bulked-segregant experiment. Phenotyping combined with genetic information of the ACCase genotyping per population. In red the number of individuals that carry a TSR mutation. In green wildtype individuals. In dark-green, wildtype individuals were selected for the two extreme bulks of the bulked-segregant analysis (D1= sensitive bulk, A5 & A6 = resistant bulk). Phenotyping was done with ACCase inhibitor Axial® 50. Gradient score starting from D1=Dead (no more green material) to A6 =Alive (no difference from untreated control).

**Supplementary Figure 6.**
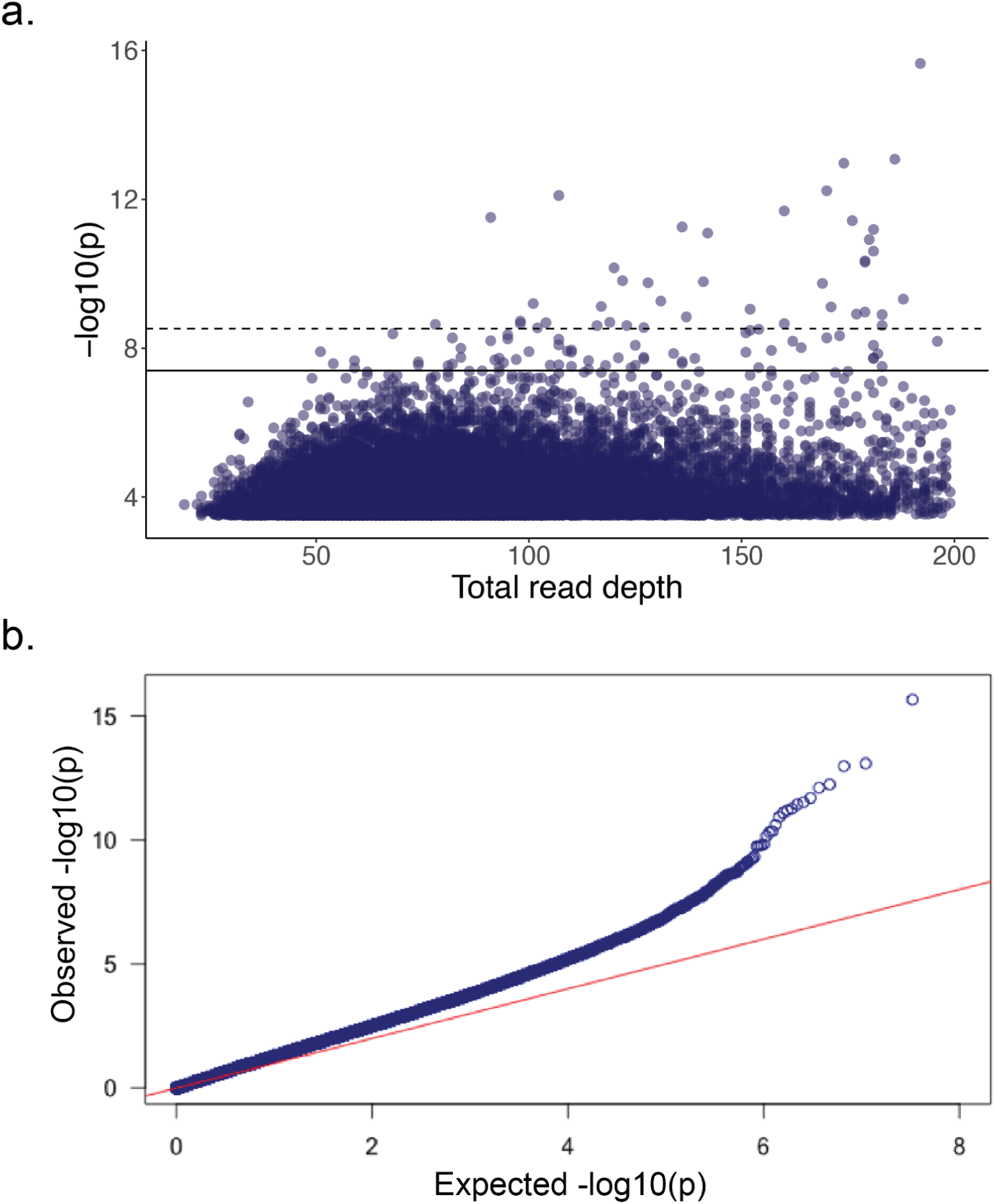
Bulked-segregant analysis. **a,** Read depth of -log_10_(p-values) higher than 3.5. The 100 SNPs with the highest p-values are plotted above the solid line (7.4), and the dashed line represents the 0.05 Bonferroni p-value threshold for 16 million SNPs: p= -log_10_(3.1e^-09^) = 8.5. **b,** qqplot of Manhattan plot in Figure 3c. The analysis was performed with BayPass using the IS covariate mode for pools^39, 40^.

**Supplementary Figure 7.**
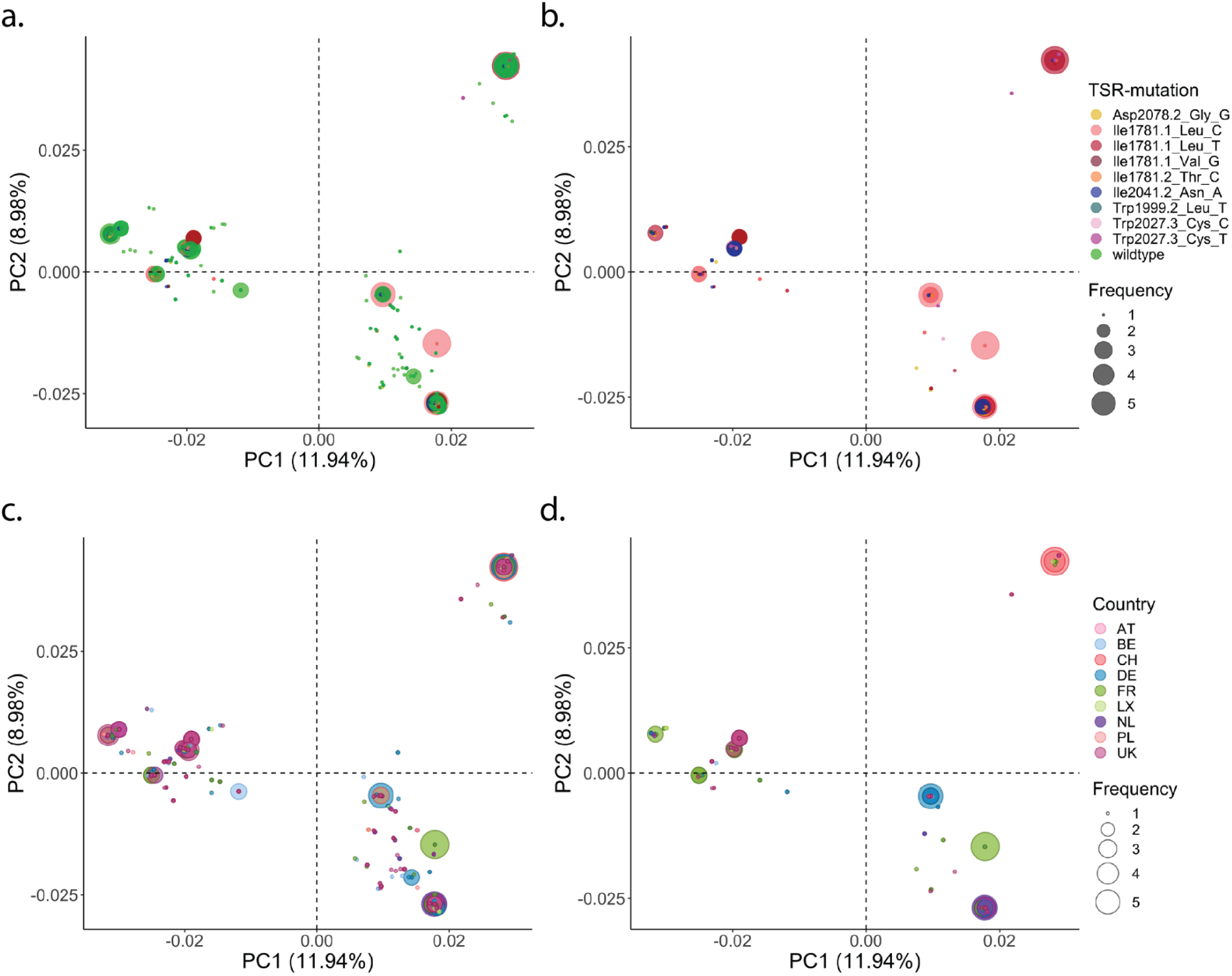
ACCase haplotype principal component analysis (PCA). Eigenvectors of the first two components are shown. **a,** Target-site-resistance (TSR) annotation of all existing haplotypes including wildtype haplotypes. **b,** Only TSR haplotypes. **c,** Country-specific colouring of all existing haplotypes. **d,** Country-specific colouring of exclusively TSR haplotypes. The values in brackets show the explained variance.

**Supplementary Figure 8.**
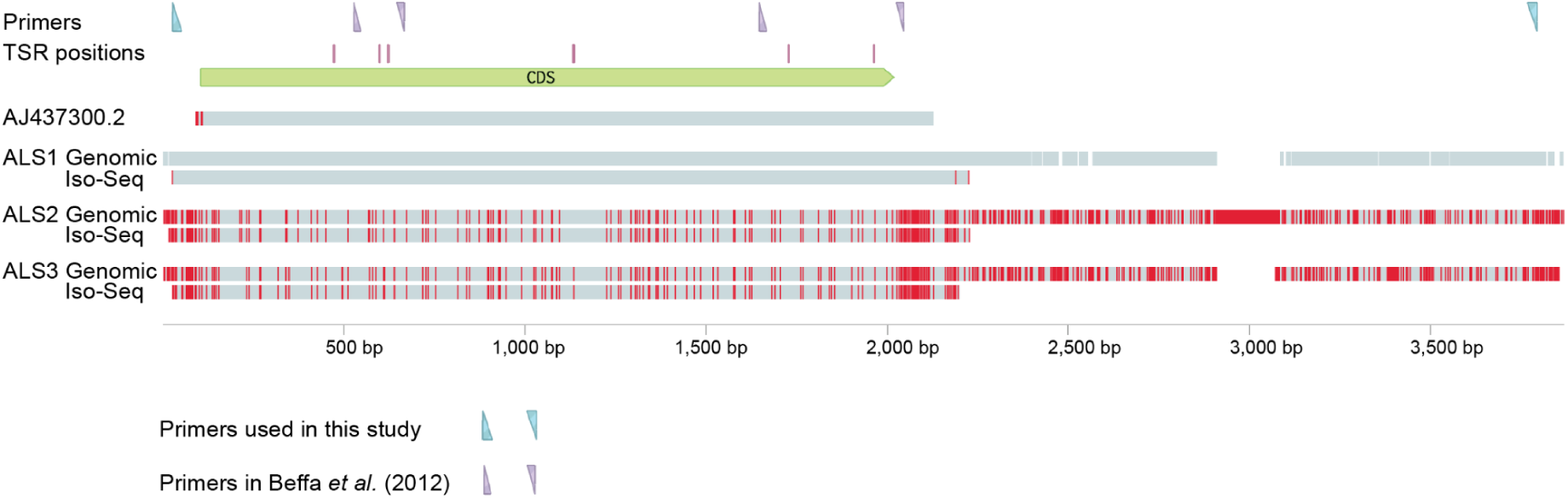
The multiple copies of ALS. Multiple alignment performed with Clustal Omega^117^ between the widely studied ALS GenBank entry of *A. myosuroides* AJ437300.2 (ref. ^48^), three genomic loci encoding ALS genes, and three representative Iso-Seq reads (each with an average read quality of q93) corresponding to each of the three Iso-Seq clusters determined by pbaa (https://github.com/PacificBiosciences/pbAA) with data from all five tissues combined. Indicated are also the positions of the seven known TSR mutations in ALS, the primers used in this study to selectively amplify ALS1, and the two pairs of primers commonly used to genotype TSRs Pro197 and Ala205 (first pair), and Trp574 and Ser653 (second pair)^49^.

**Supplementary Figure 9.**
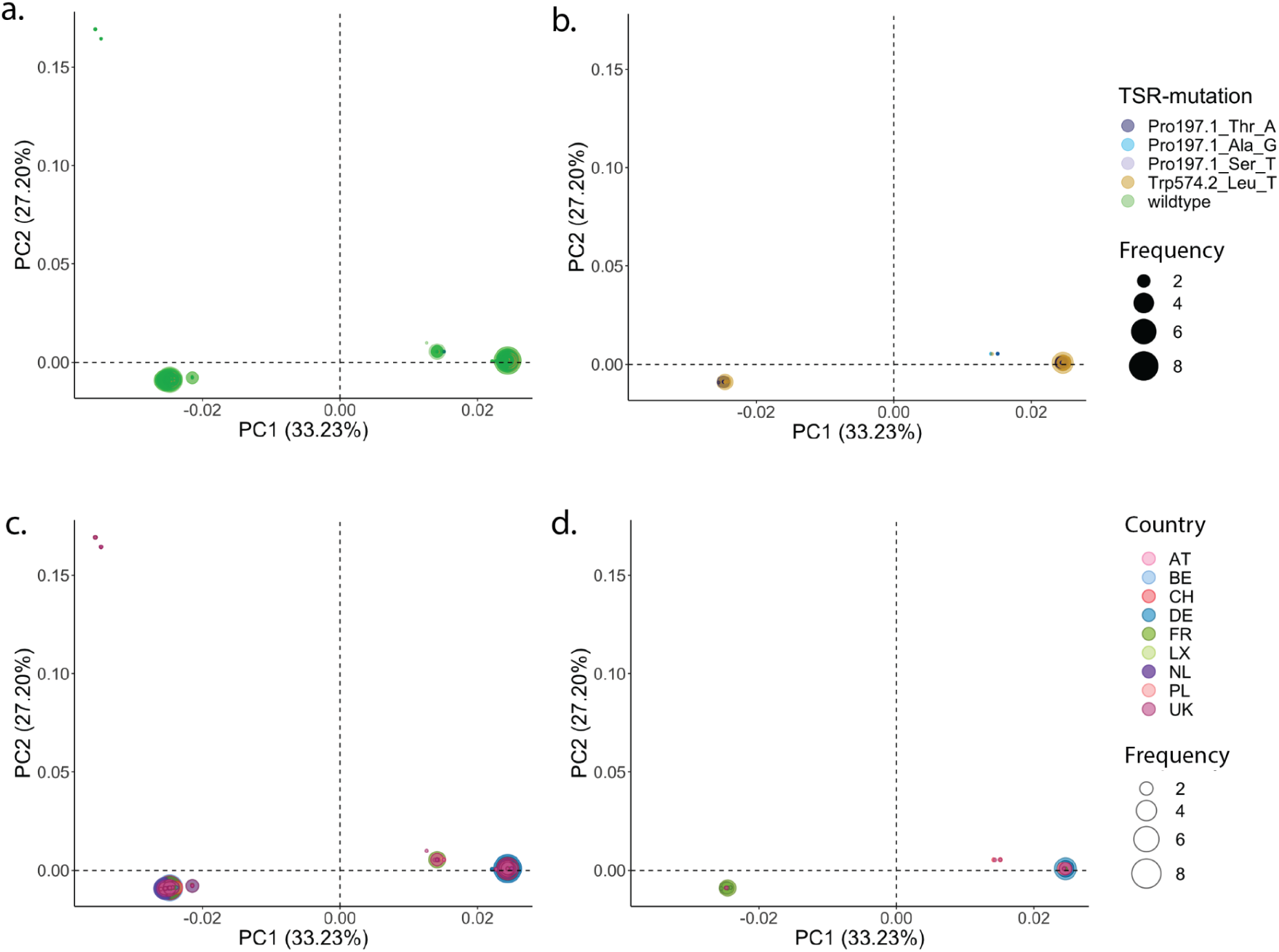
ALS haplotype principal component analysis (PCA). Eigenvectors of the first two components are shown. **a,** Target-site-resistance (TSR) annotation of all existing haplotypes including wildtype haplotypes. **b,** Only TSR haplotypes. **c,** Country-specific colouring of all existing haplotypes. **d,** Country-specific colouring of exclusively TSR haplotypes. The values in brackets show the explained variance.

**Supplementary Figure 10.**
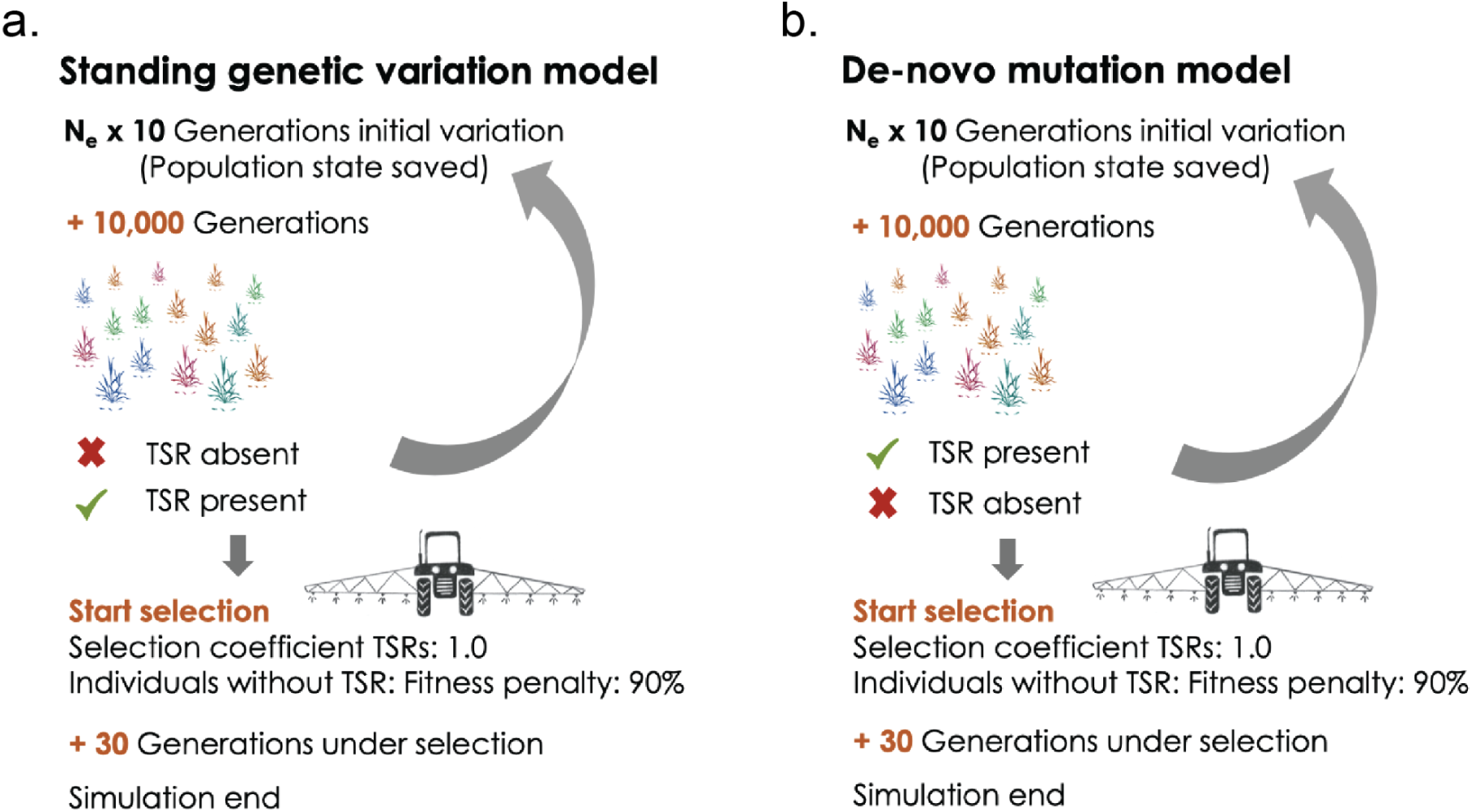
Visualisation of the SLiM simulation models. **a,** Standing genetic variation model. **b,** *De novo* mutation model.

**Supplementary Figure 11.**
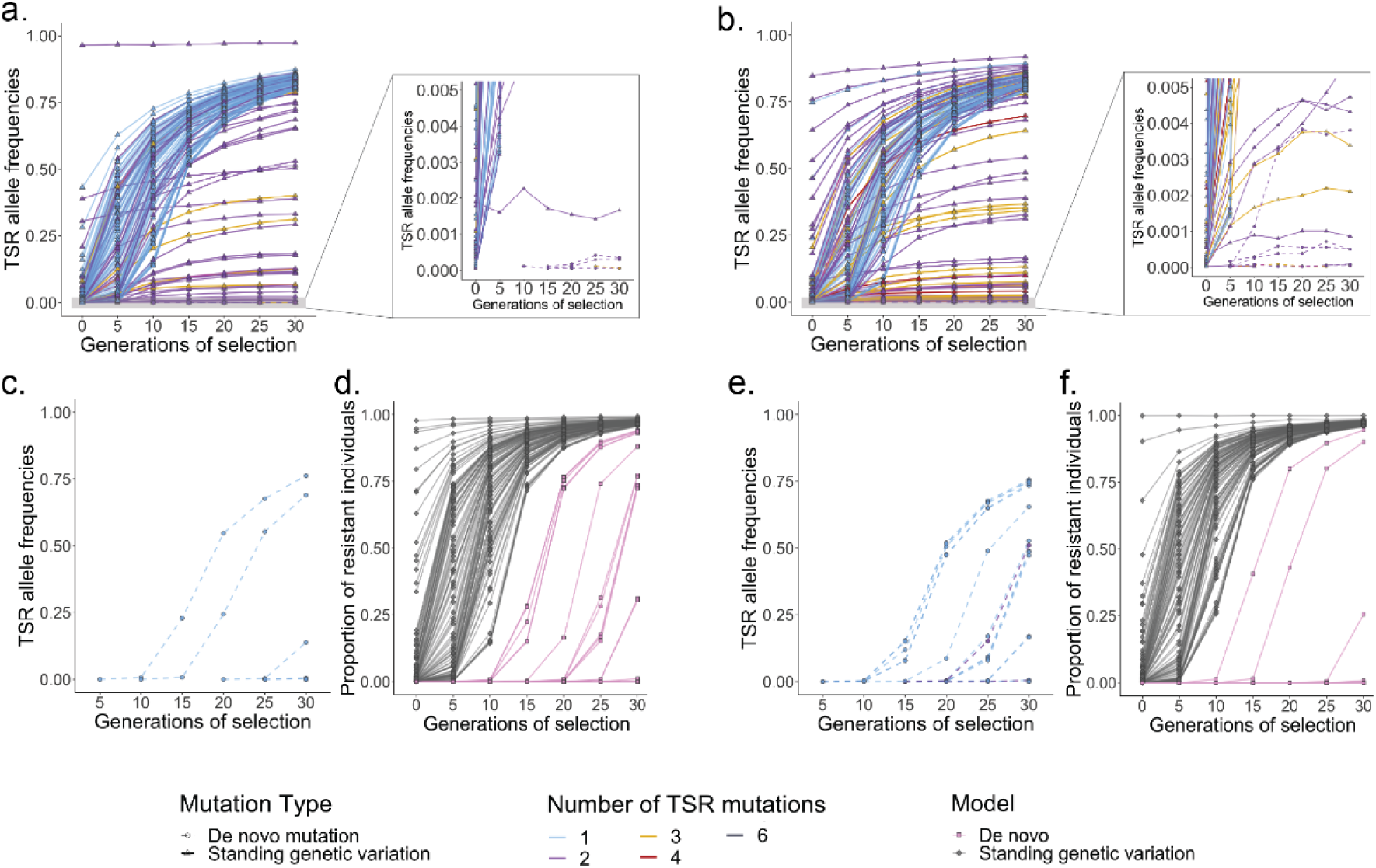
Simulations of allele frequencies for mutations conferring target-site resistance (TSR) (no Intron/Exon structure). From these simulations, the proportions of resistant individuals per field population can be inferred. Mutations other than TSRs after the start of selection pressure are considered as neutral. One hundred runs per model are shown for an effective population size of 42,000 individuals (**a, c, d**) and 84,000 individuals (**b, e, f**). Continuous lines represent mutations originating from standing genetic variation, *de novo* TSR mutations are shown with dashed lines. The colours indicate the total number of TSR mutations per population. **a, b,** Standing genetic variation model, with TSR mutations pre-existing in the populations before herbicide selection. Shown is the increase in TSR allele frequencies under herbicide selection of up to 30 generations, with one herbicide application per generation. The right panel shows a truncated y-axis at 0.005 TSR allele frequencies. **c, e,** *De novo* mutation model. Any TSR mutation that might have arisen before the start of selection has been lost again, so that no TSR mutations are present when the simulation starts. **d, f,** Proportion of resistant individuals across the hundred simulated populations under each of the two different models.

**Supplementary Figure 12.**
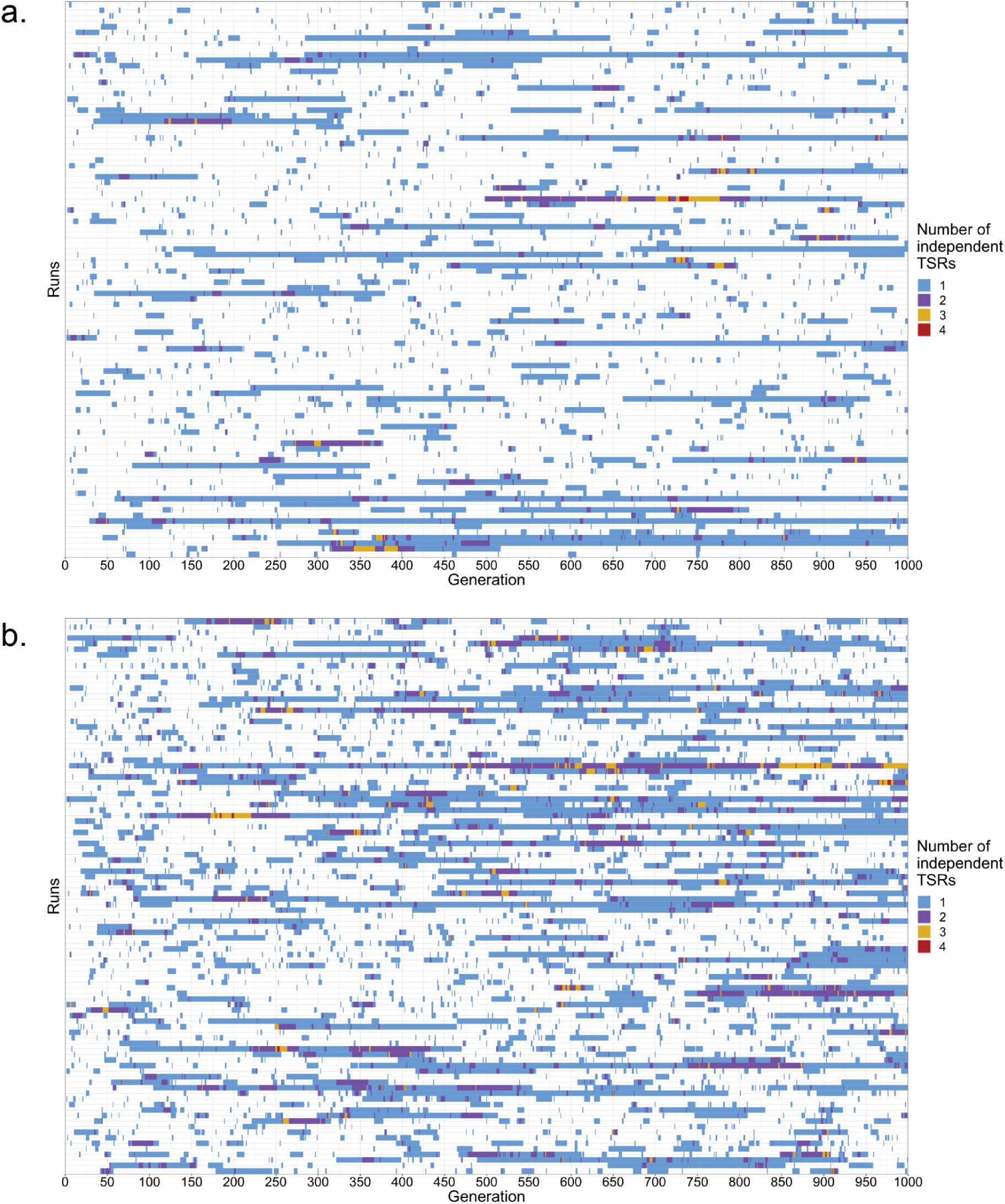
Simulations of TSR abundance under neutrality. SLiM simulations representing 100 independent runs of population evolution for **a,** 42,000 individuals and **b,** 84,000 individuals over 1,000 generations under neutrality. The occurrence and loss of TSRs due to genetic drift can be observed. Colours indicate the number of TSRs present in each generation in a given run.

## Extended data

**Extended data File 1**

**[Extended data File 1.](https://keeper.mpdl.mpg.de/f/7a0477d313d847cfa2a6/?dl=1) Sheet1,** Top 100 SNP associations in the bulked-segregant analysis, and the annotation of genes within 50 Kb downstream or upstream of the hit. **Sheet2,** List of paralogs retained in collinear regions (anchors), their *K*_S_ values, and whether they are part of Supplementary Figure 1c. **Sheet3,** List of primers used in this study.

**Supplementary extended data Figure 1**

**[Extended data Figure 1.](https://keeper.mpdl.mpg.de/f/21b2d051e0c94c36bebc/?dl=1)** ACCase networks and trees for 47 European populations. Haplotype network and maximum likelihood (ML)-tree per population. The colour code in all networks and trees shows target-site resistances (TSRs) and wildtype haplotypes in green.

**Supplementary extended data Figure 2**

**[Extended data Figure 2.](https://keeper.mpdl.mpg.de/f/9f8db9dafb744c03970f/?dl=1)** ALS networks and trees for 47 European populations. Haplotype network and maximum likelihood (ML)-tree per population. The colour code in all networks and trees shows target-site resistances (TSRs) and wildtype haplotypes in green.

## Supplementary Tables

**Supplementary Table 1.**
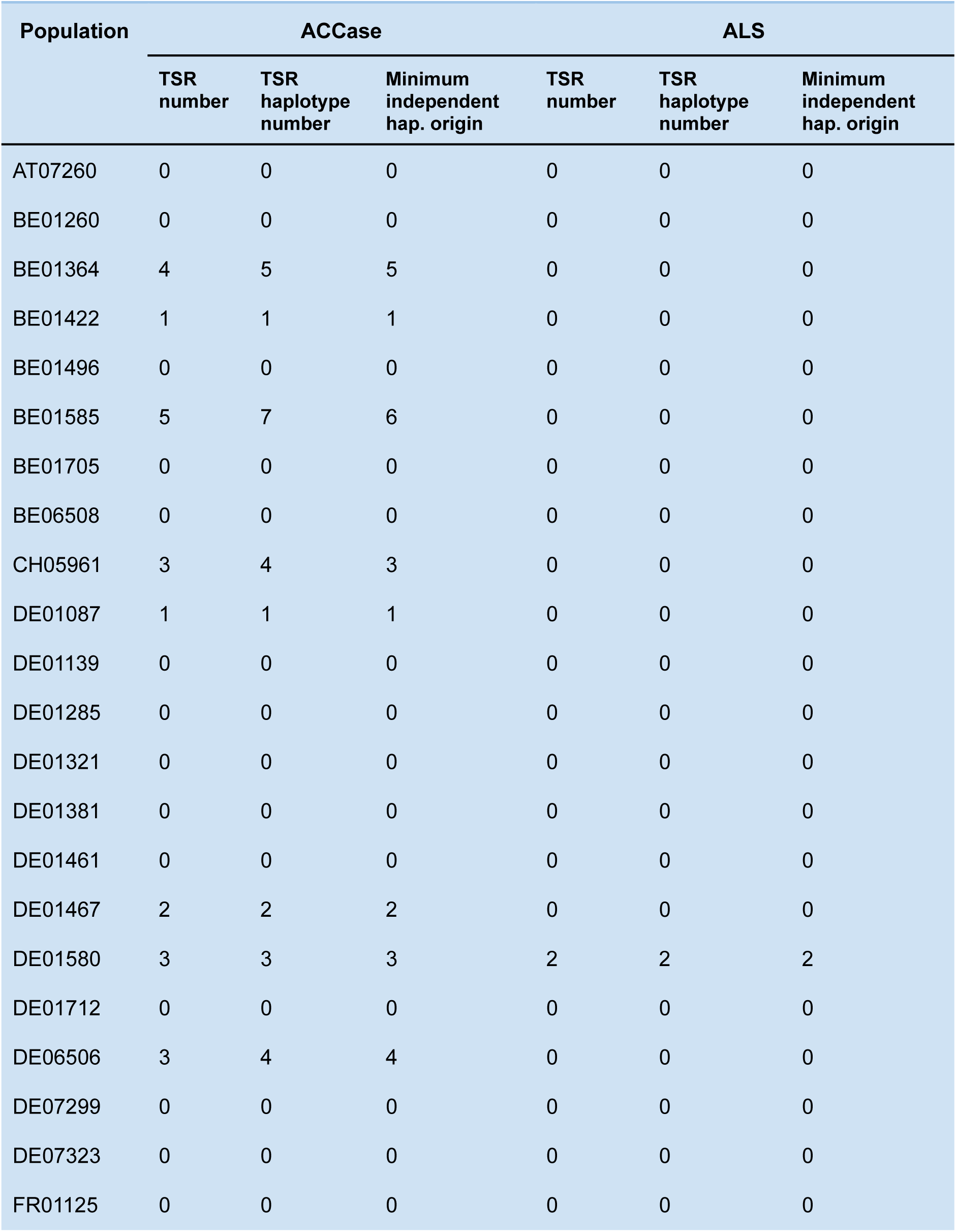
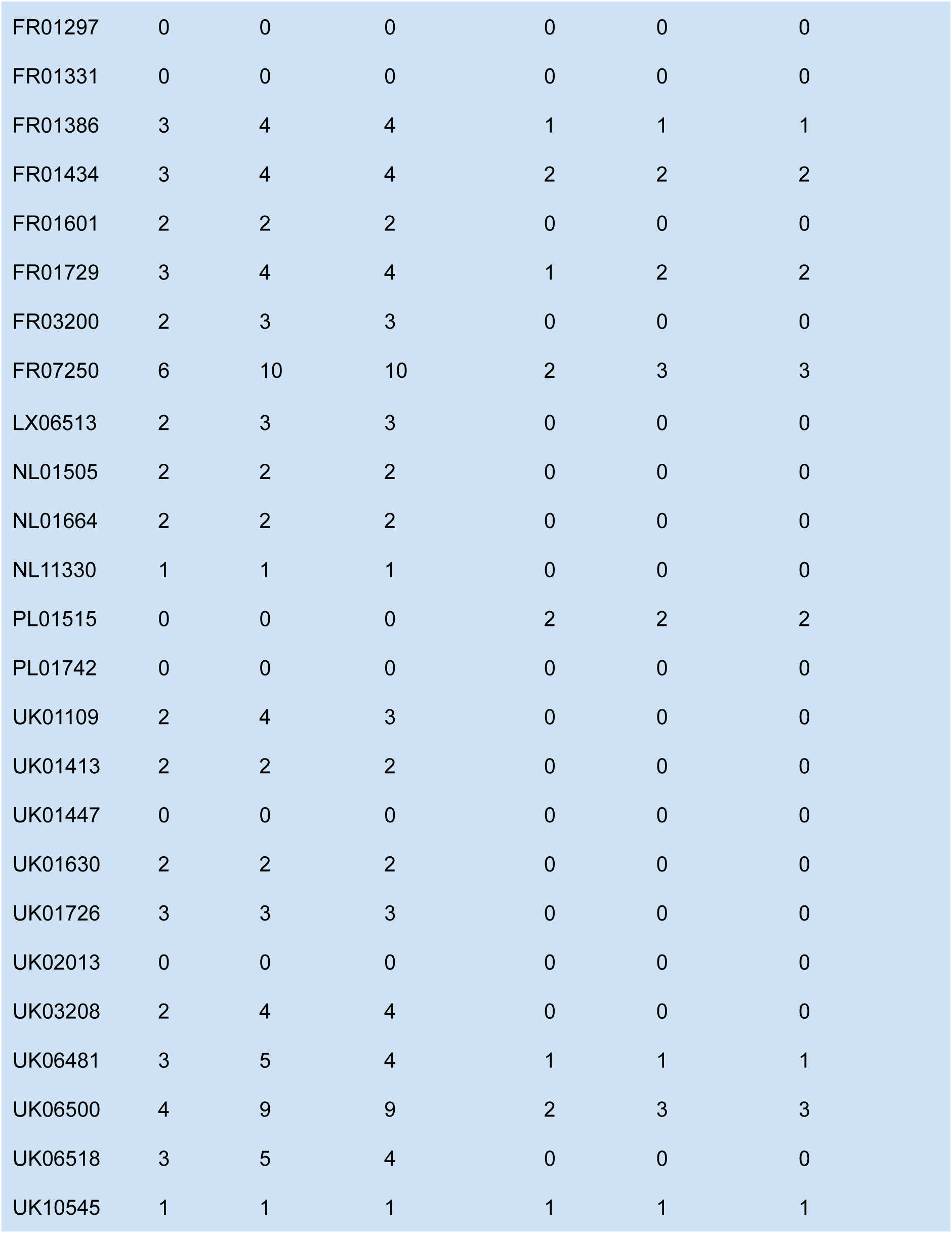
TSR and TSR haplotype number per population.

| Population | ACCase |  |  | ALS |  |  |
| --- | --- | --- | --- | --- | --- | --- |
|  | TSR number | TSR haplotype number | Minimum independent hap. origin | TSR number | TSR haplotype number | Minimum independent hap. origin |
| AT07260 | 0 | 0 | 0 | 0 | 0 | 0 |
| BE01260 | 0 | 0 | 0 | 0 | 0 | 0 |
| BE01364 | 4 | 5 | 5 | 0 | 0 | 0 |
| BE01422 | 1 | 1 | 1 | 0 | 0 | 0 |
| BE01496 | 0 | 0 | 0 | 0 | 0 | 0 |
| BE01585 | 5 | 7 | 6 | 0 | 0 | 0 |
| BE01705 | 0 | 0 | 0 | 0 | 0 | 0 |
| BE06508 | 0 | 0 | 0 | 0 | 0 | 0 |
| CH05961 | 3 | 4 | 3 | 0 | 0 | 0 |
| DE01087 | 1 | 1 | 1 | 0 | 0 | 0 |
| DE01139 | 0 | 0 | 0 | 0 | 0 | 0 |
| DE01285 | 0 | 0 | 0 | 0 | 0 | 0 |
| DE01321 | 0 | 0 | 0 | 0 | 0 | 0 |
| DE01381 | 0 | 0 | 0 | 0 | 0 | 0 |
| DE01461 | 0 | 0 | 0 | 0 | 0 | 0 |
| DE01467 | 2 | 2 | 2 | 0 | 0 | 0 |
| DE01580 | 3 | 3 | 3 | 2 | 2 | 2 |
| DE01712 | 0 | 0 | 0 | 0 | 0 | 0 |
| DE06506 | 3 | 4 | 4 | 0 | 0 | 0 |
| DE07299 | 0 | 0 | 0 | 0 | 0 | 0 |
| DE07323 | 0 | 0 | 0 | 0 | 0 | 0 |
| FR01125 | 0 | 0 | 0 | 0 | 0 | 0 |
| FR01297 | 0 | 0 | 0 | 0 | 0 | 0 |
| FR01331 | 0 | 0 | 0 | 0 | 0 | 0 |
| FR01386 | 3 | 4 | 4 | 1 | 1 | 1 |
| FR01434 | 3 | 4 | 4 | 2 | 2 | 2 |
| FR01601 | 2 | 2 | 2 | 0 | 0 | 0 |
| FR01729 | 3 | 4 | 4 | 1 | 2 | 2 |
| FR03200 | 2 | 3 | 3 | 0 | 0 | 0 |
| FR07250 | 6 | 10 | 10 | 2 | 3 | 3 |
| LX06513 | 2 | 3 | 3 | 0 | 0 | 0 |
| NL01505 | 2 | 2 | 2 | 0 | 0 | 0 |
| NL01664 | 2 | 2 | 2 | 0 | 0 | 0 |
| NL11330 | 1 | 1 | 1 | 0 | 0 | 0 |
| PL01515 | 0 | 0 | 0 | 2 | 2 | 2 |
| PL01742 | 0 | 0 | 0 | 0 | 0 | 0 |
| UK01109 | 2 | 4 | 3 | 0 | 0 | 0 |
| UK01413 | 2 | 2 | 2 | 0 | 0 | 0 |
| UK01447 | 0 | 0 | 0 | 0 | 0 | 0 |
| UK01630 | 2 | 2 | 2 | 0 | 0 | 0 |
| UK01726 | 3 | 3 | 3 | 0 | 0 | 0 |
| UK02013 | 0 | 0 | 0 | 0 | 0 | 0 |
| UK03208 | 2 | 4 | 4 | 0 | 0 | 0 |
| UK06481 | 3 | 5 | 4 | 1 | 1 | 1 |
| UK06500 | 4 | 9 | 9 | 2 | 3 | 3 |
| UK06518 | 3 | 5 | 4 | 0 | 0 | 0 |
| UK10545 | 1 | 1 | 1 | 1 | 1 | 1 |

**Supplementary Table 2.**
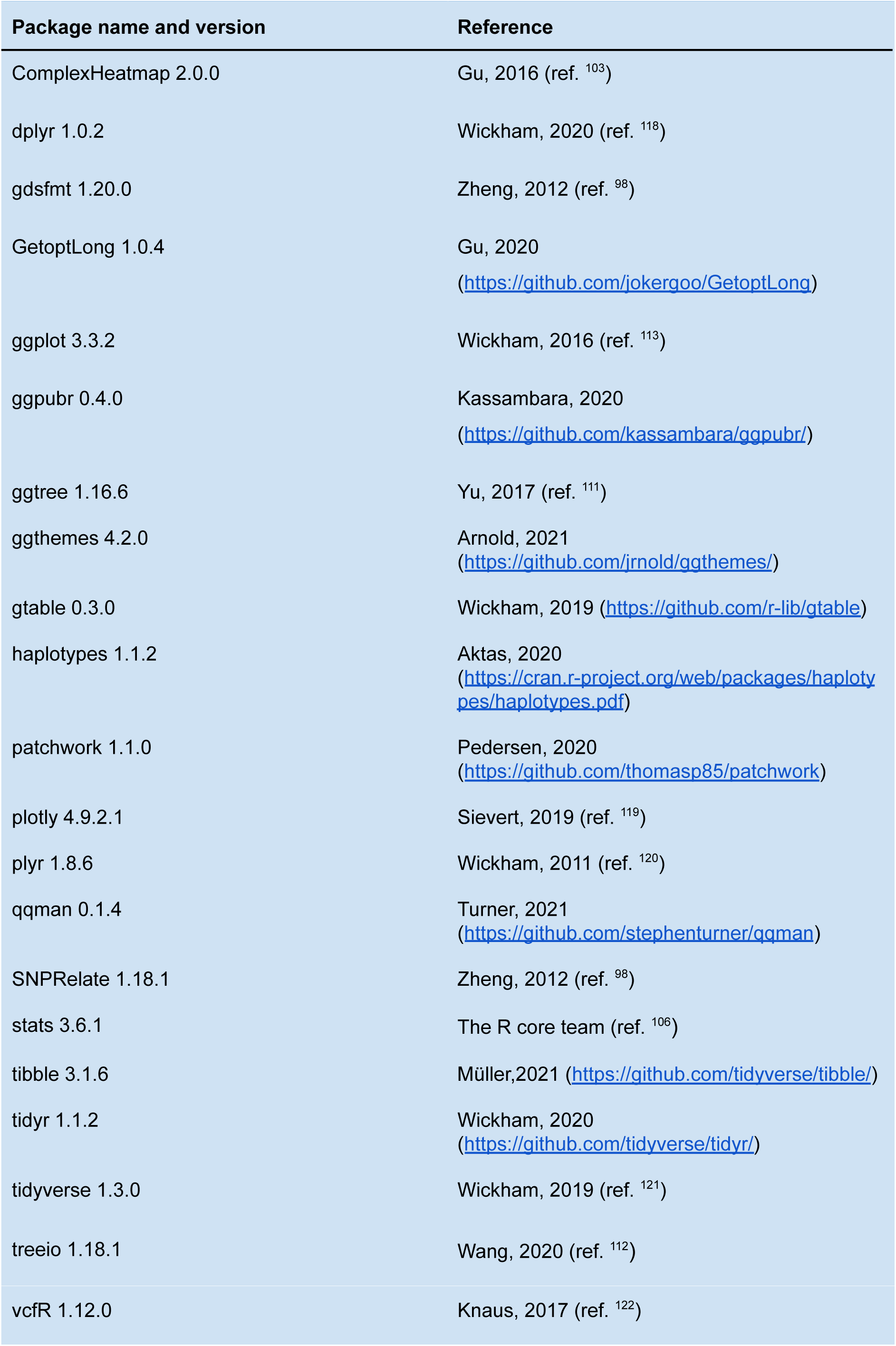
R-packages used for data manipulation and visualisation.

