## Supplementary extended data Figure 2 for "Standing genetic variation fuels rapid evolution of herbicide resistance in blackgrass"

# AT07260

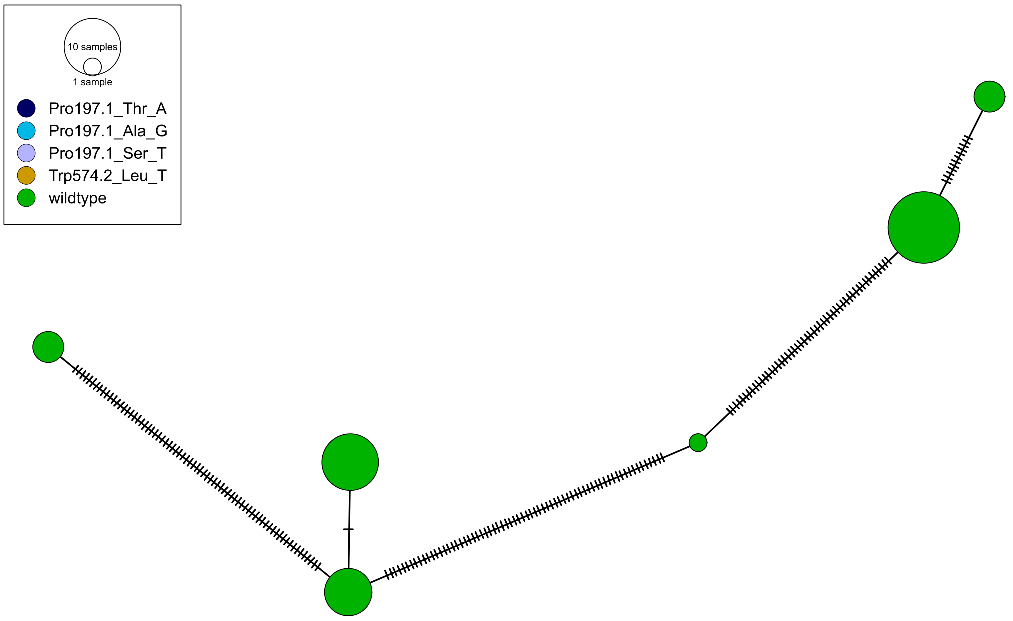
10 samples
1 sample
Pro197.1_Thr_A
Pro197.1_Ala_G
Pro197.1_Ser_T
Trp574.2_Leu_T
wildtype

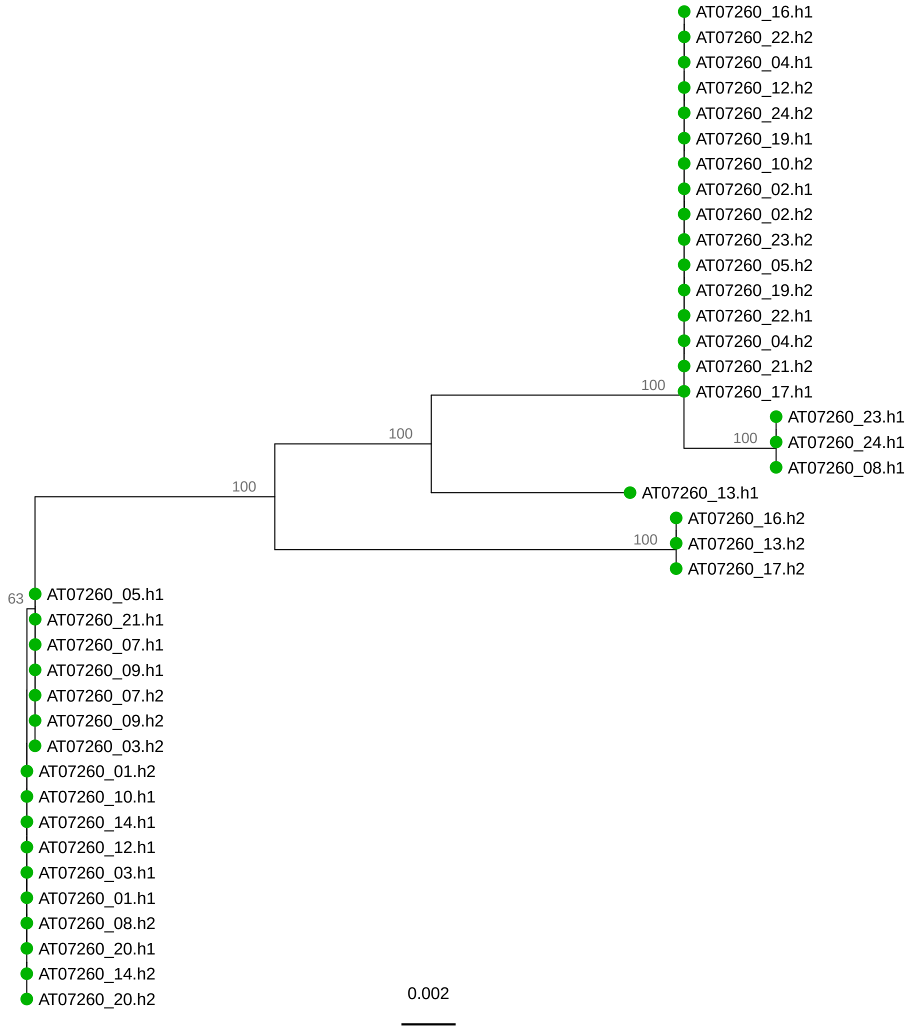
AT07260_16.h1
AT07260_22.h2
AT07260_04.h1
AT07260_12.h2
AT07260_24.h2
AT07260_19.h1
AT07260_10.h2
AT07260_02.h1
AT07260_02.h2
AT07260_23.h2
AT07260_05.h2
AT07260_19.h2
AT07260_22.h1
AT07260_04.h2
AT07260_21.h2
100
AT07260_17.h1
AT07260_23.h1
100
AT07260_24.h1
100
AT07260_08.h1
100
AT07260_13.h1
AT07260_16.h2
100
AT07260_13.h2
AT07260_17.h2
63
AT07260_05.h1
AT07260_21.h1
AT07260_07.h1
AT07260_09.h1
AT07260_07.h2
AT07260_09.h2
AT07260_03.h2
AT07260_01.h2
AT07260_10.h1
AT07260_14.h1
AT07260_12.h1
AT07260_03.h1
AT07260_01.h1
AT07260_08.h2
AT07260_20.h1
AT07260_14.h2
AT07260_20.h2
0.002

# BE01260

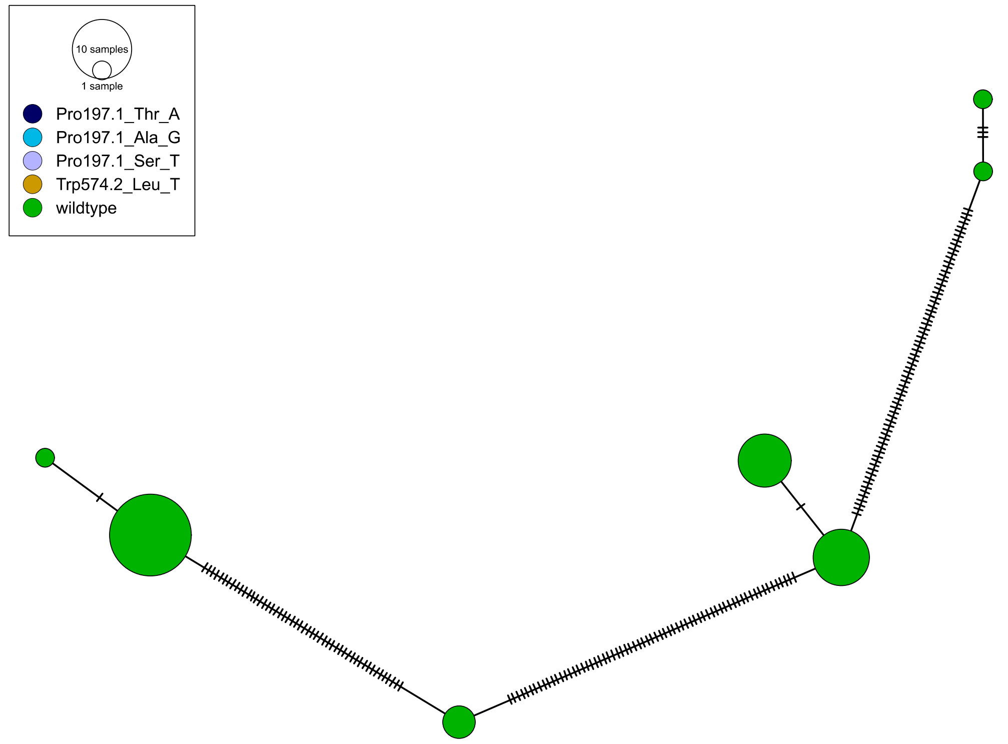
10 samples
1 sample
Pro197.1_Thr_A
Pro197.1_Ala_G
Pro197.1_Ser_T
Trp574.2_Leu_T
wildtype

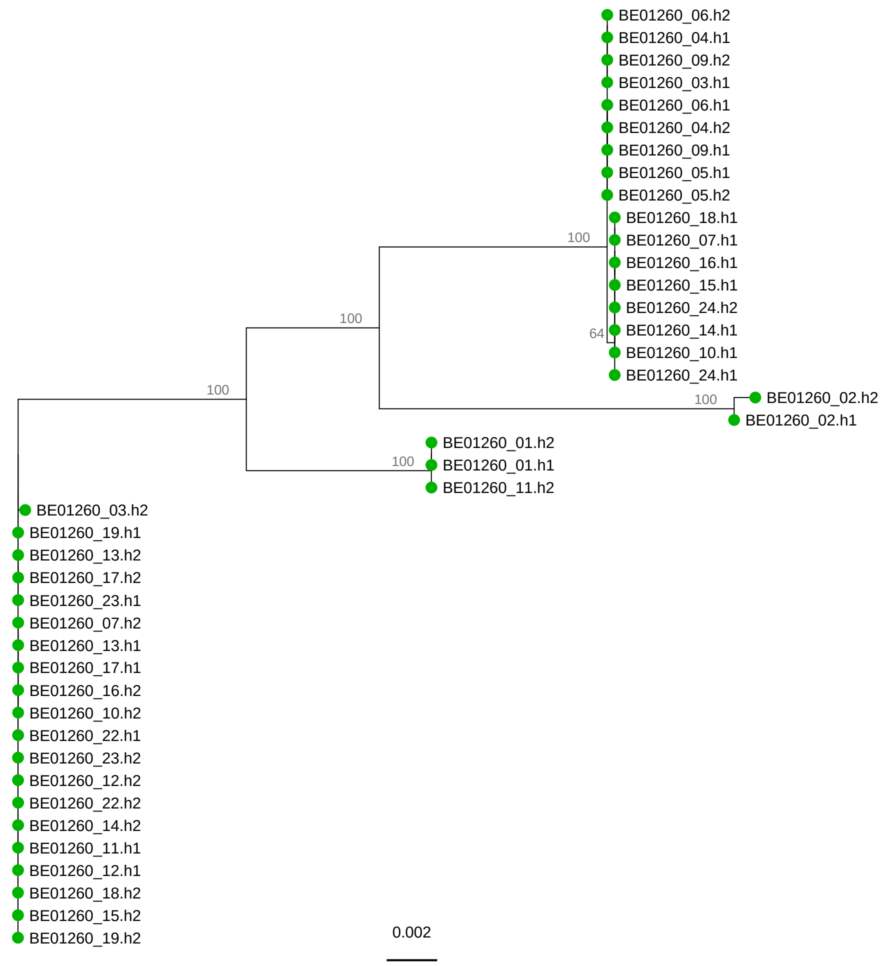
BE01260_06.h2
BE01260_04.h1
BE01260_09.h2
BE01260_03.h1
BE01260_06.h1
BE01260_04.h2
BE01260_09.h1
BE01260_05.h1
BE01260_05.h2
BE01260_18.h1
BE01260_07.h1
BE01260_16.h1
BE01260_15.h1
BE01260_24.h2
BE01260_14.h1
BE01260_10.h1
BE01260_24.h1
100
100
100
64
100
BE01260_02.h2
BE01260_02.h1
100
BE01260_01.h2
BE01260_01.h1
BE01260_11.h2
BE01260_03.h2
BE01260_19.h1
BE01260_13.h2
BE01260_17.h2
BE01260_23.h1
BE01260_07.h2
BE01260_13.h1
BE01260_17.h1
BE01260_16.h2
BE01260_10.h2
BE01260_22.h1
BE01260_23.h2
BE01260_12.h2
BE01260_22.h2
BE01260_14.h2
BE01260_11.h1
BE01260_12.h1
BE01260_18.h2
BE01260_15.h2
BE01260_19.h2
0.002

BE01364

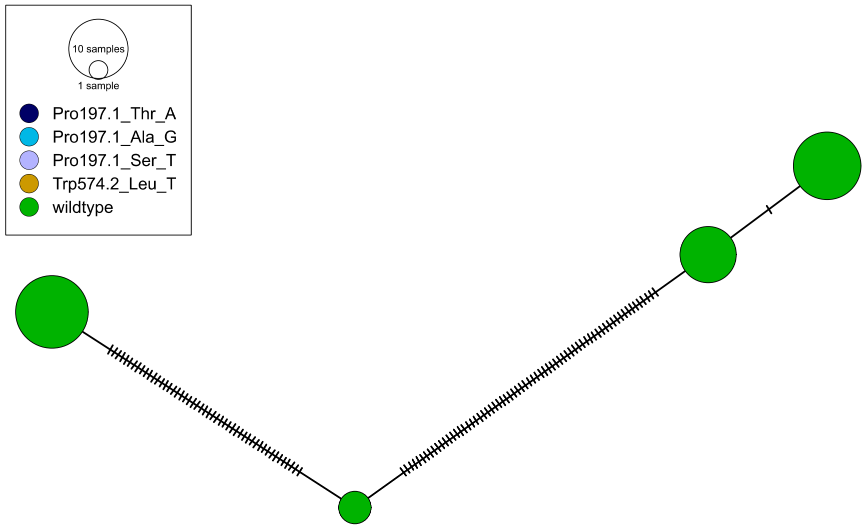
10 samples
1 sample
Pro197.1_Thr_A
Pro197.1_Ala_G
Pro197.1_Ser_T
Trp574.2_Leu_T
wildtype

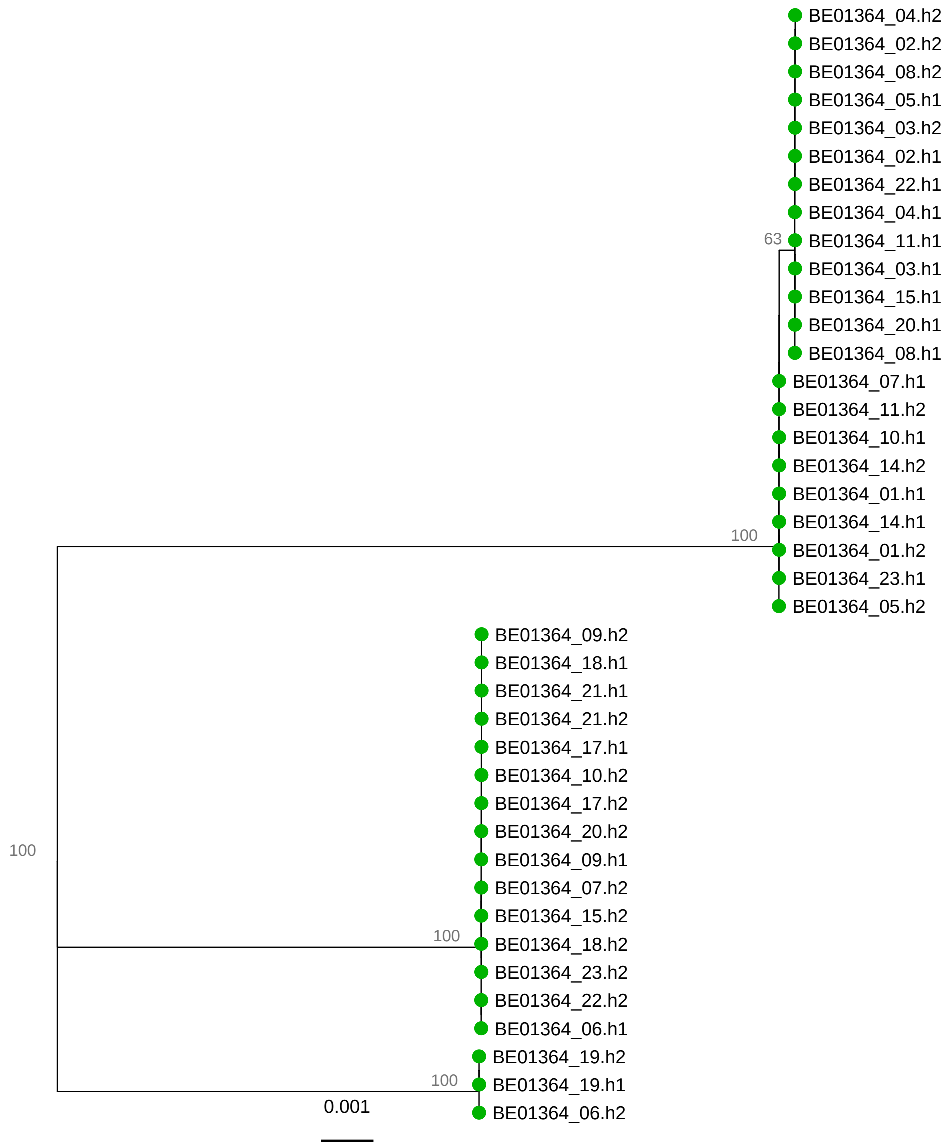
BE01364_04.h2
BE01364_02.h2
BE01364_08.h2
BE01364_05.h1
BE01364_03.h2
BE01364_02.h1
BE01364_22.h1
BE01364_04.h1
63
BE01364_11.h1
BE01364_03.h1
BE01364_15.h1
BE01364_20.h1
BE01364_08.h1
BE01364_07.h1
BE01364_11.h2
BE01364_10.h1
BE01364_14.h2
BE01364_01.h1
BE01364_14.h1
100
BE01364_01.h2
BE01364_23.h1
BE01364_05.h2
BE01364_09.h2
BE01364_18.h1
BE01364_21.h1
BE01364_21.h2
BE01364_17.h1
BE01364_10.h2
BE01364_17.h2
BE01364_20.h2
100
BE01364_09.h1
BE01364_07.h2
BE01364_15.h2
100
BE01364_18.h2
BE01364_23.h2
BE01364_22.h2
BE01364_06.h1
BE01364_19.h2
100
BE01364_19.h1
0.001
BE01364_06.h2

BE01422

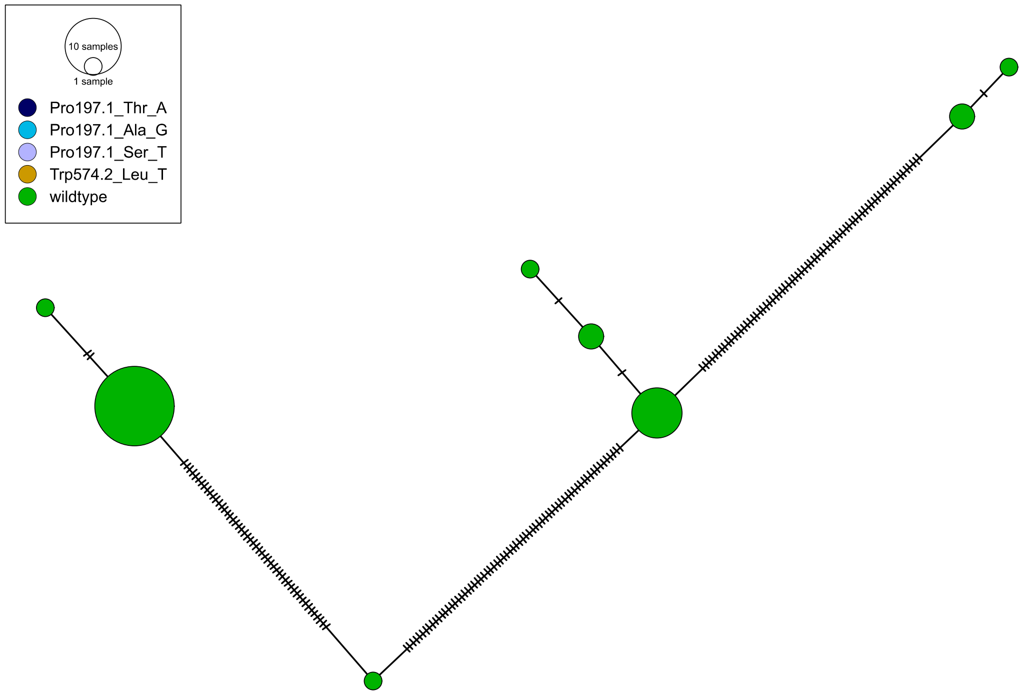

10 samples
1 sample
Pro197.1_Thr_A
Pro197.1_Ala_G
Pro197.1_Ser_T
Trp574.2_Leu_T
wildtype


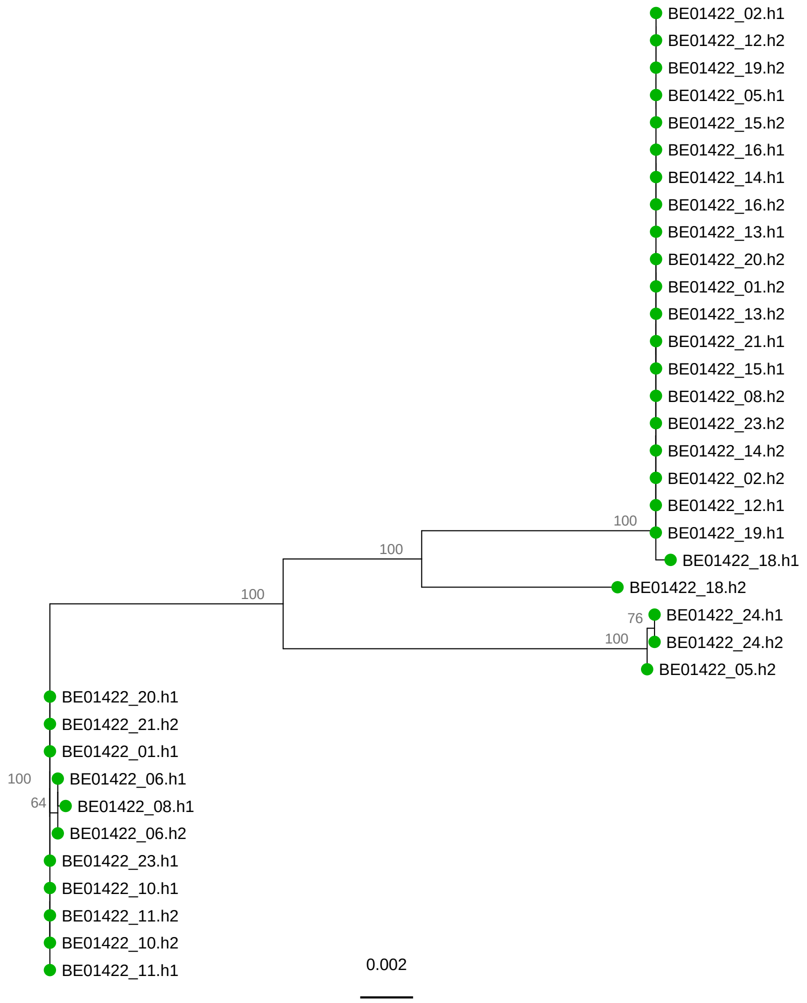

BE01422_02.h1
BE01422_12.h2
BE01422_19.h2
BE01422_05.h1
BE01422_15.h2
BE01422_16.h1
BE01422_14.h1
BE01422_16.h2
BE01422_13.h1
BE01422_20.h2
BE01422_01.h2
BE01422_13.h2
BE01422_21.h1
BE01422_15.h1
BE01422_08.h2
BE01422_23.h2
BE01422_14.h2
BE01422_02.h2
BE01422_12.h1
100
BE01422_19.h1
100
BE01422_18.h1
100
BE01422_18.h2
76
BE01422_24.h1
100
BE01422_24.h2
BE01422_05.h2
BE01422_20.h1
BE01422_21.h2
BE01422_01.h1
100
BE01422_06.h1
64
BE01422_08.h1
BE01422_06.h2
BE01422_23.h1
BE01422_10.h1
BE01422_11.h2
BE01422_10.h2
BE01422_11.h1
0.002

# BE01496

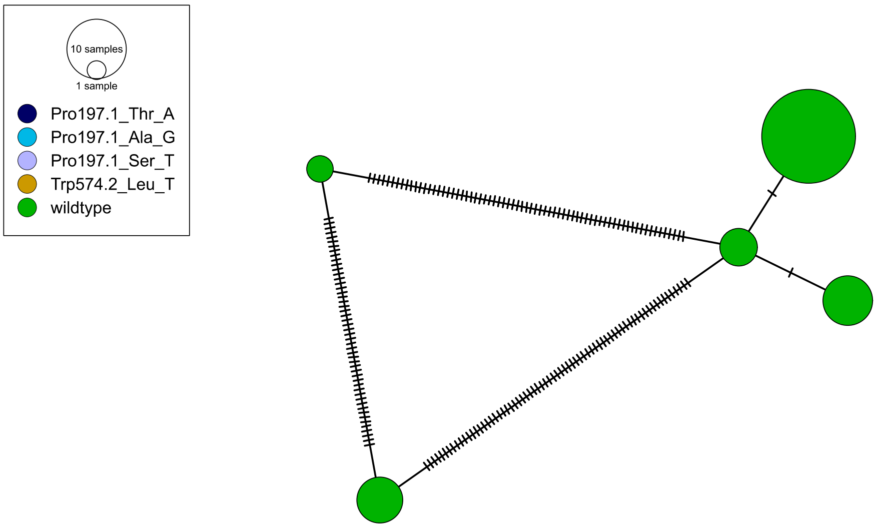

10 samples
1 sample
Pro197.1_Thr_A
Pro197.1_Ala_G
Pro197.1_Ser_T
Trp574.2_Leu_T
wildtype


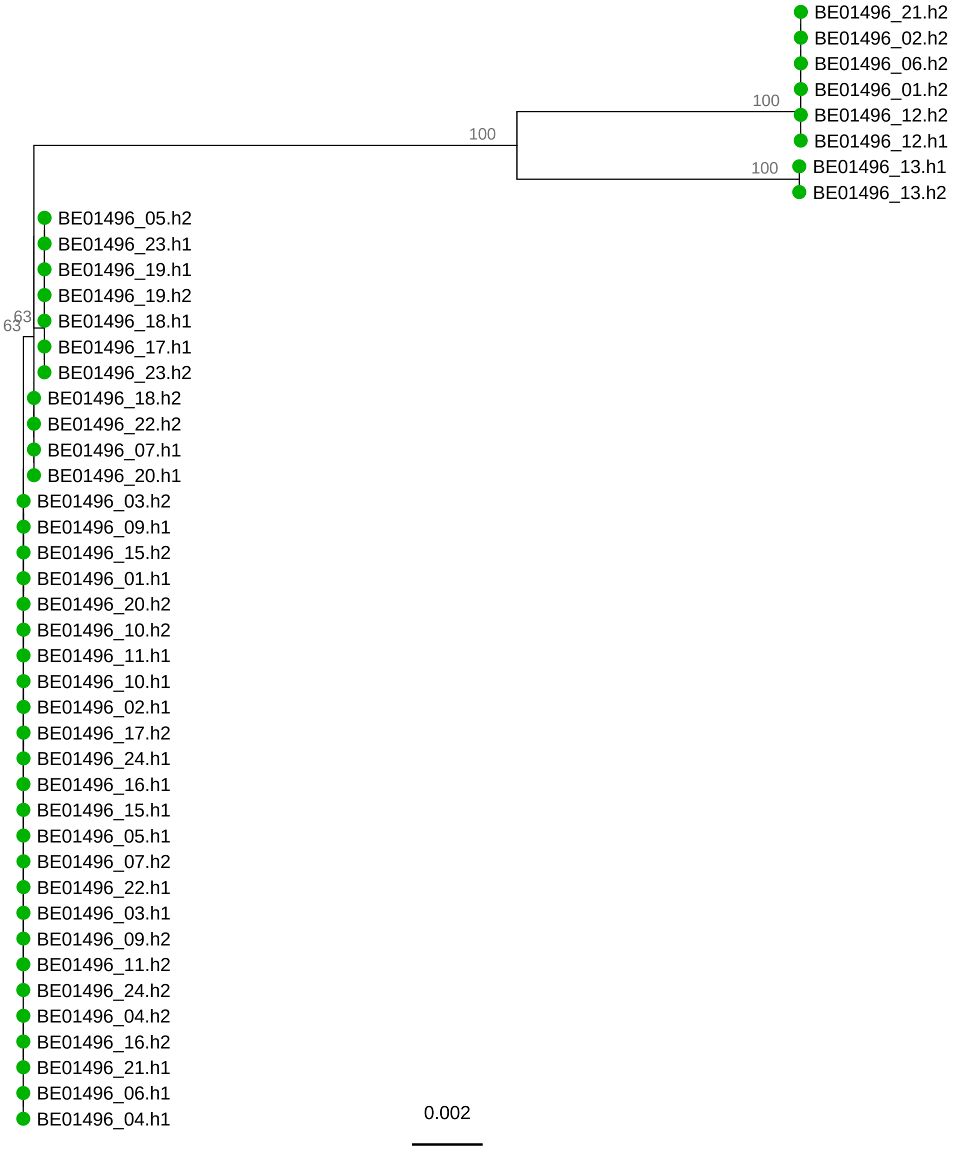

BE01496_21.h2
BE01496_02.h2
BE01496_06.h2
BE01496_01.h2
BE01496_12.h2
BE01496_12.h1
BE01496_13.h1
BE01496_13.h2
100
100
100
63
63
BE01496_05.h2
BE01496_23.h1
BE01496_19.h1
BE01496_19.h2
BE01496_18.h1
BE01496_17.h1
BE01496_23.h2
BE01496_18.h2
BE01496_22.h2
BE01496_07.h1
BE01496_20.h1
BE01496_03.h2
BE01496_09.h1
BE01496_15.h2
BE01496_01.h1
BE01496_20.h2
BE01496_10.h2
BE01496_11.h1
BE01496_10.h1
BE01496_02.h1
BE01496_17.h2
BE01496_24.h1
BE01496_16.h1
BE01496_15.h1
BE01496_05.h1
BE01496_07.h2
BE01496_22.h1
BE01496_03.h1
BE01496_09.h2
BE01496_11.h2
BE01496_24.h2
BE01496_04.h2
BE01496_16.h2
BE01496_21.h1
BE01496_06.h1
BE01496_04.h1
0.002

BE01585

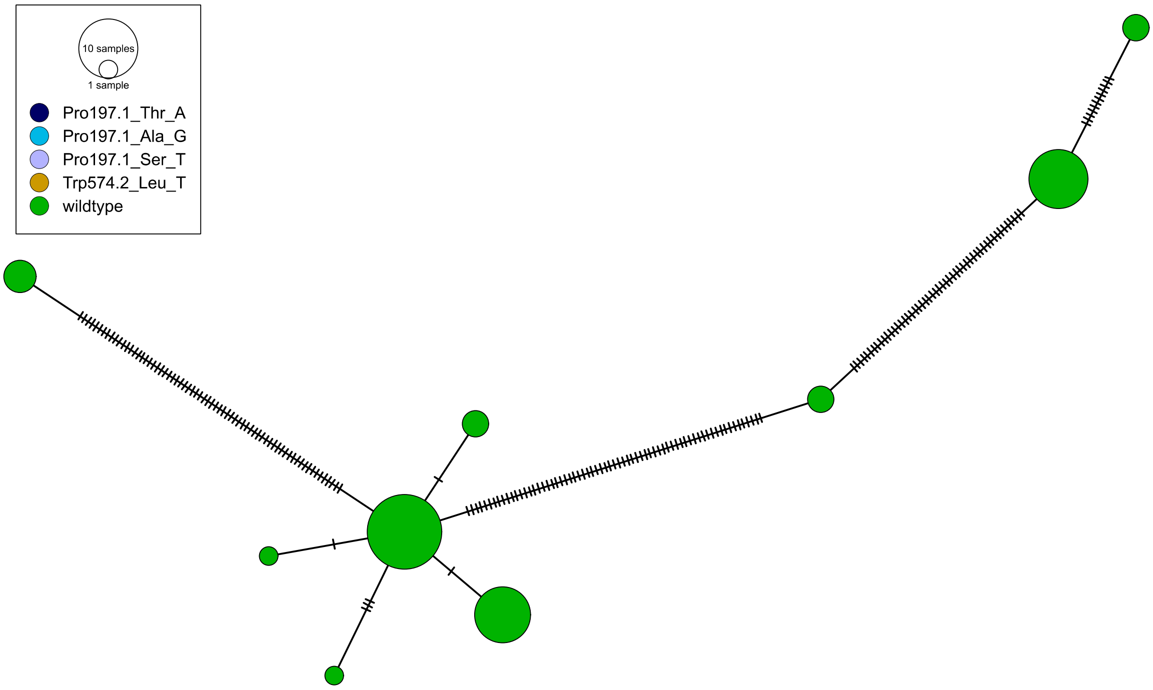
10 samples
1 sample
Pro197.1_Thr_A
Pro197.1_Ala_G
Pro197.1_Ser_T
Trp574.2_Leu_T
wildtype

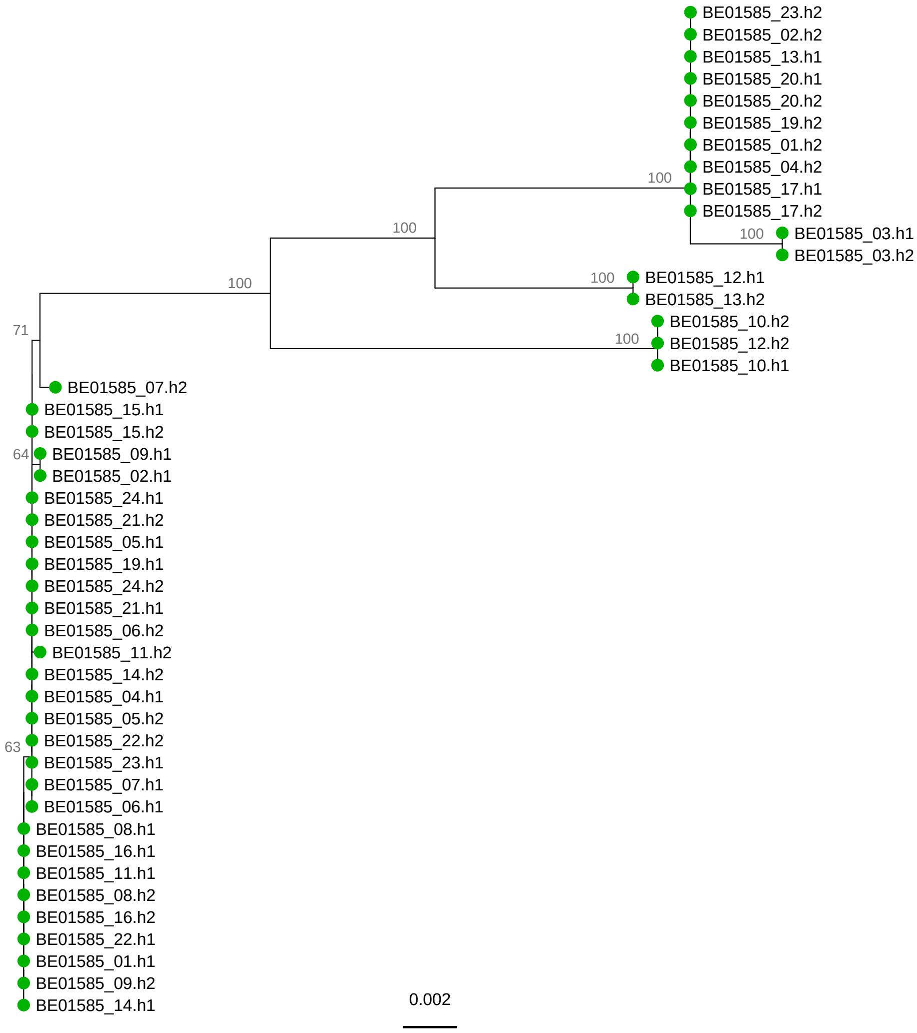
BE01585_23.h2
BE01585_02.h2
BE01585_13.h1
BE01585_20.h1
BE01585_20.h2
BE01585_19.h2
BE01585_01.h2
BE01585_04.h2
100
BE01585_17.h1
BE01585_17.h2
100
100
BE01585_03.h1
BE01585_03.h2
100
100
BE01585_12.h1
BE01585_13.h2
BE01585_10.h2
100
BE01585_12.h2
BE01585_10.h1
71
BE01585_07.h2
BE01585_15.h1
BE01585_15.h2
64
BE01585_09.h1
BE01585_02.h1
BE01585_24.h1
BE01585_21.h2
BE01585_05.h1
BE01585_19.h1
BE01585_24.h2
BE01585_21.h1
BE01585_06.h2
BE01585_11.h2
BE01585_14.h2
BE01585_04.h1
BE01585_05.h2
63
BE01585_22.h2
BE01585_23.h1
BE01585_07.h1
BE01585_06.h1
BE01585_08.h1
BE01585_16.h1
BE01585_11.h1
BE01585_08.h2
BE01585_16.h2
BE01585_22.h1
BE01585_01.h1
BE01585_09.h2
BE01585_14.h1
0.002

BE01705

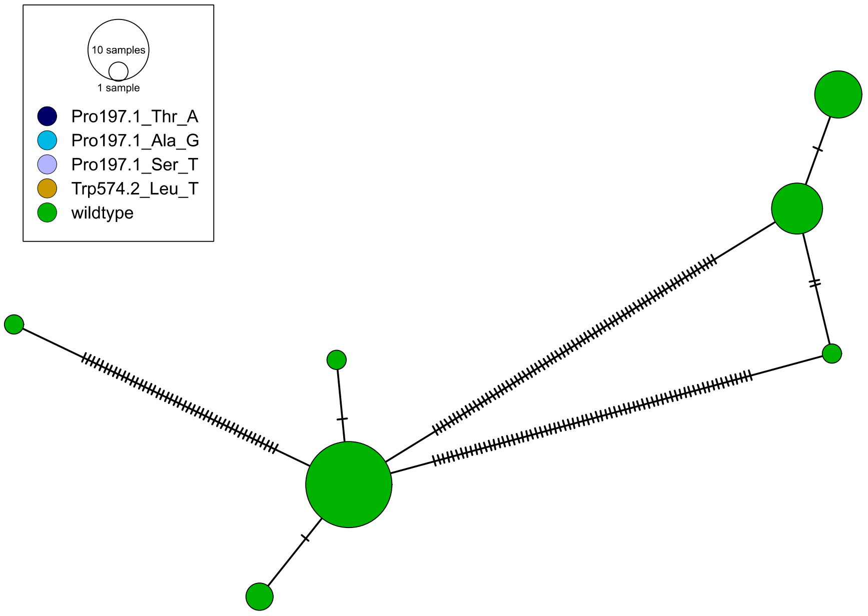
10 samples
1 sample
Pro197.1_Thr_A
Pro197.1_Ala_G
Pro197.1_Ser_T
Trp574.2_Leu_T
wildtype

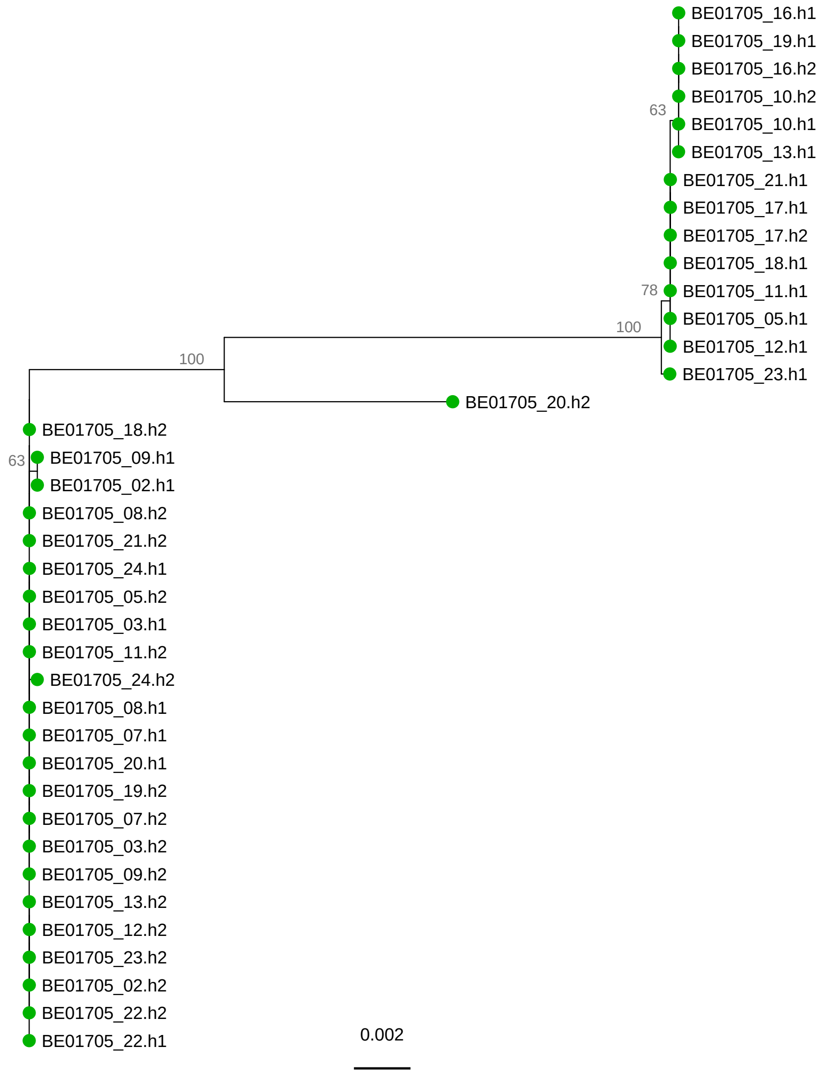
BE01705_16.h1
BE01705_19.h1
BE01705_16.h2
BE01705_10.h2
63
BE01705_10.h1
BE01705_13.h1
BE01705_21.h1
BE01705_17.h1
BE01705_17.h2
BE01705_18.h1
78
BE01705_11.h1
BE01705_05.h1
100
BE01705_12.h1
BE01705_23.h1
100
BE01705_20.h2
BE01705_18.h2
63
BE01705_09.h1
BE01705_02.h1
BE01705_08.h2
BE01705_21.h2
BE01705_24.h1
BE01705_05.h2
BE01705_03.h1
BE01705_11.h2
BE01705_24.h2
BE01705_08.h1
BE01705_07.h1
BE01705_20.h1
BE01705_19.h2
BE01705_07.h2
BE01705_03.h2
BE01705_09.h2
BE01705_13.h2
BE01705_12.h2
BE01705_23.h2
BE01705_02.h2
BE01705_22.h2
BE01705_22.h1
0.002

# BE06508

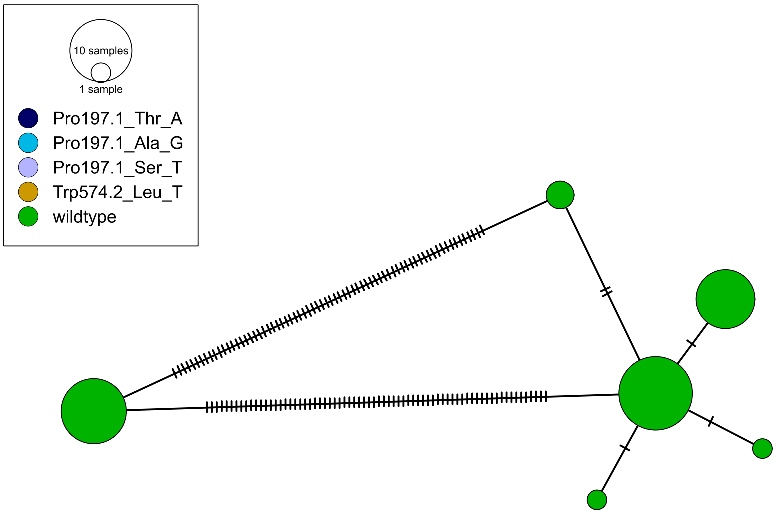

10 samples
1 sample
Pro197.1_Thr_A
Pro197.1_Ala_G
Pro197.1_Ser_T
Trp574.2_Leu_T
wildtype


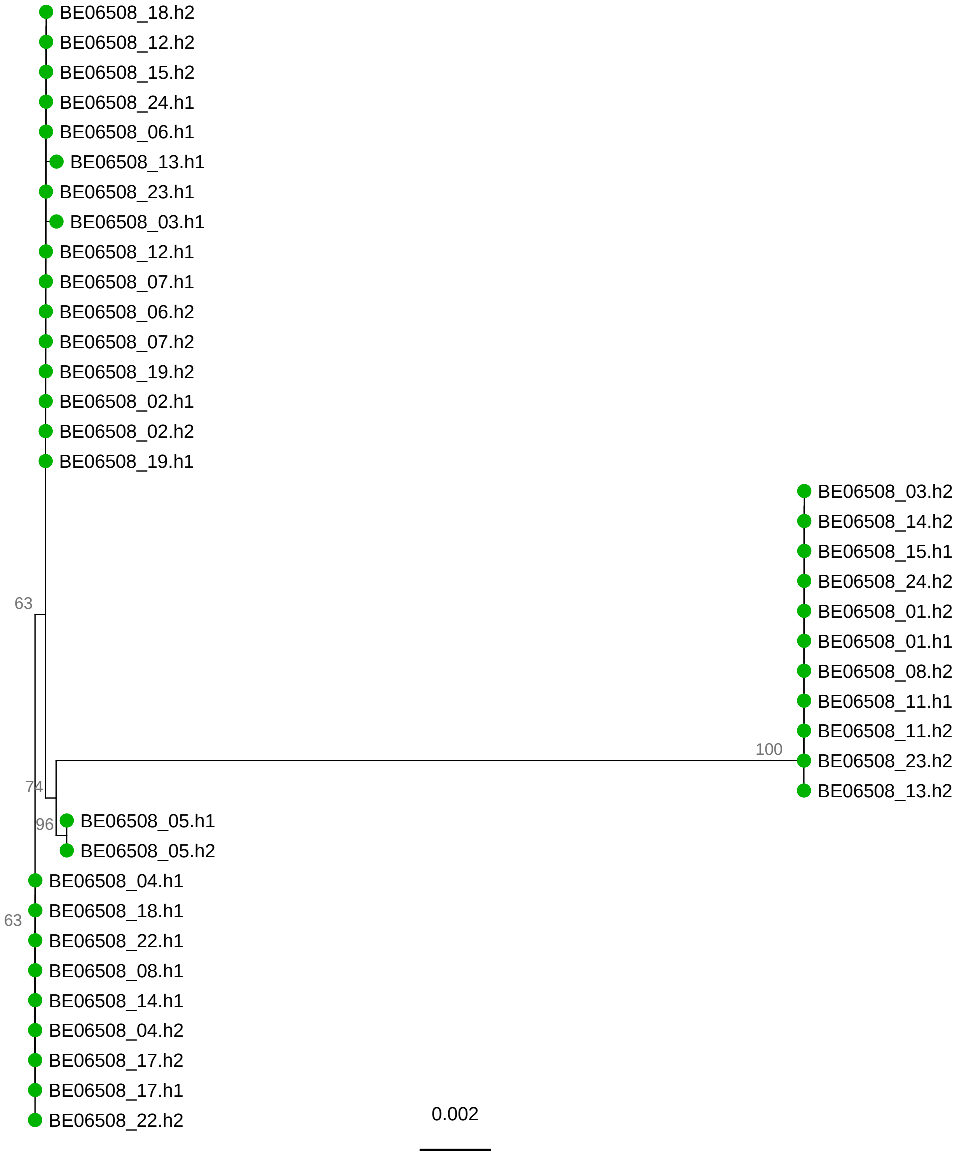

BE06508_18.h2
BE06508_12.h2
BE06508_15.h2
BE06508_24.h1
BE06508_06.h1
BE06508_13.h1
BE06508_23.h1
BE06508_03.h1
BE06508_12.h1
BE06508_07.h1
BE06508_06.h2
BE06508_07.h2
BE06508_19.h2
BE06508_02.h1
BE06508_02.h2
BE06508_19.h1
63
BE06508_03.h2
BE06508_14.h2
BE06508_15.h1
BE06508_24.h2
BE06508_01.h2
BE06508_01.h1
BE06508_08.h2
BE06508_11.h1
BE06508_11.h2
100
BE06508_23.h2
BE06508_13.h2
74
96
BE06508_05.h1
BE06508_05.h2
BE06508_04.h1
63
BE06508_18.h1
BE06508_22.h1
BE06508_08.h1
BE06508_14.h1
BE06508_04.h2
BE06508_17.h2
BE06508_17.h1
BE06508_22.h2
0.002

CH05961

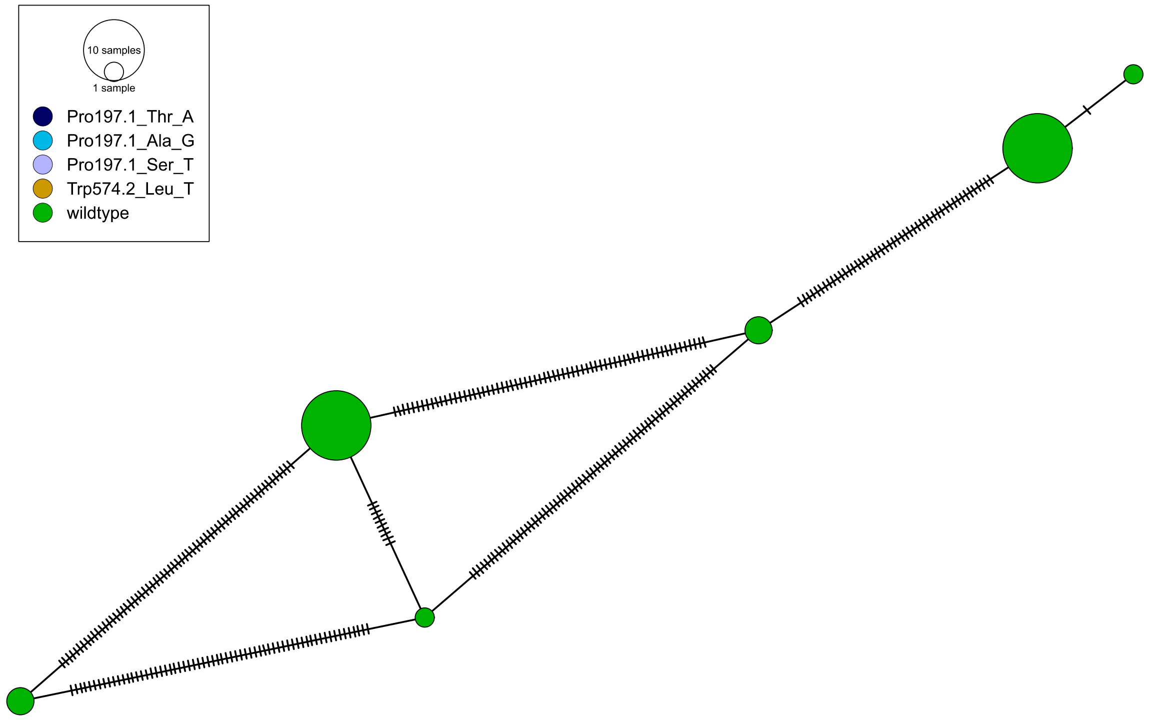

10 samples
1 sample
Pro197.1_Thr_A
Pro197.1_Ala_G
Pro197.1_Ser_T
Trp574.2_Leu_T
wildtype


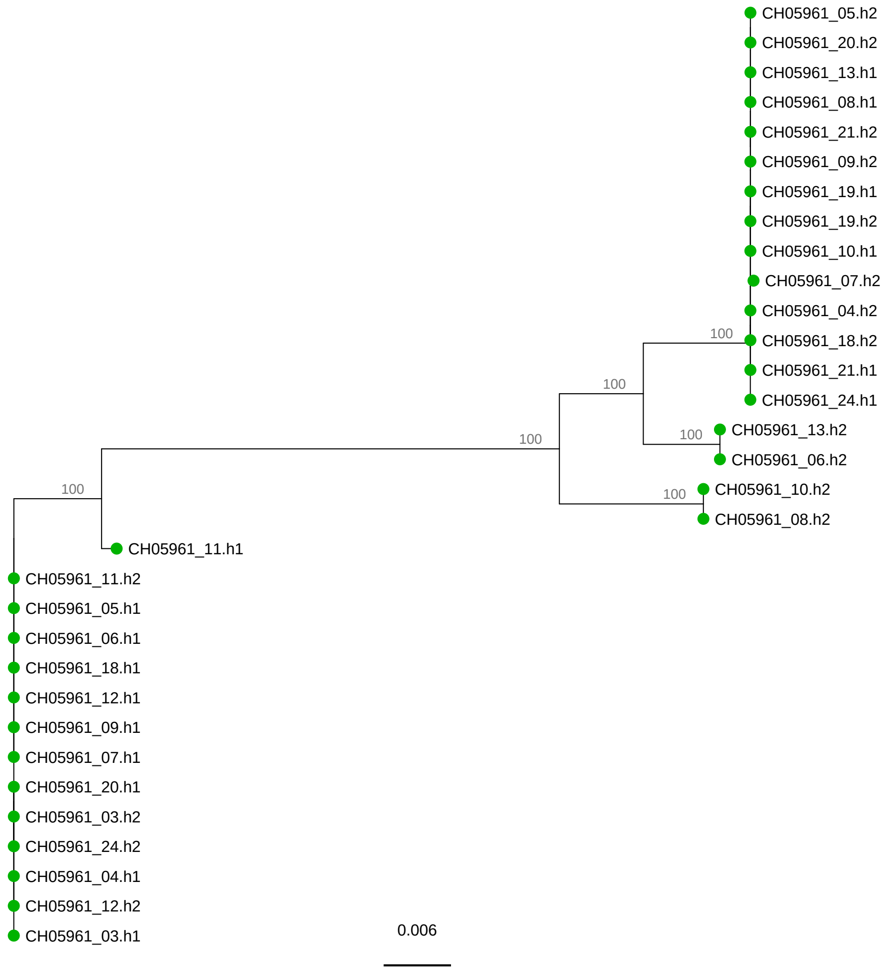

CH05961_05.h2
CH05961_20.h2
CH05961_13.h1
CH05961_08.h1
CH05961_21.h2
CH05961_09.h2
CH05961_19.h1
CH05961_19.h2
CH05961_10.h1
CH05961_07.h2
CH05961_04.h2
CH05961_18.h2
CH05961_21.h1
CH05961_24.h1
CH05961_13.h2
CH05961_06.h2
CH05961_10.h2
CH05961_08.h2
CH05961_11.h1
CH05961_11.h2
CH05961_05.h1
CH05961_06.h1
CH05961_18.h1
CH05961_12.h1
CH05961_09.h1
CH05961_07.h1
CH05961_20.h1
CH05961_03.h2
CH05961_24.h2
CH05961_04.h1
CH05961_12.h2
CH05961_03.h1
100
100
100
100
100
100
0.006

# DE01087

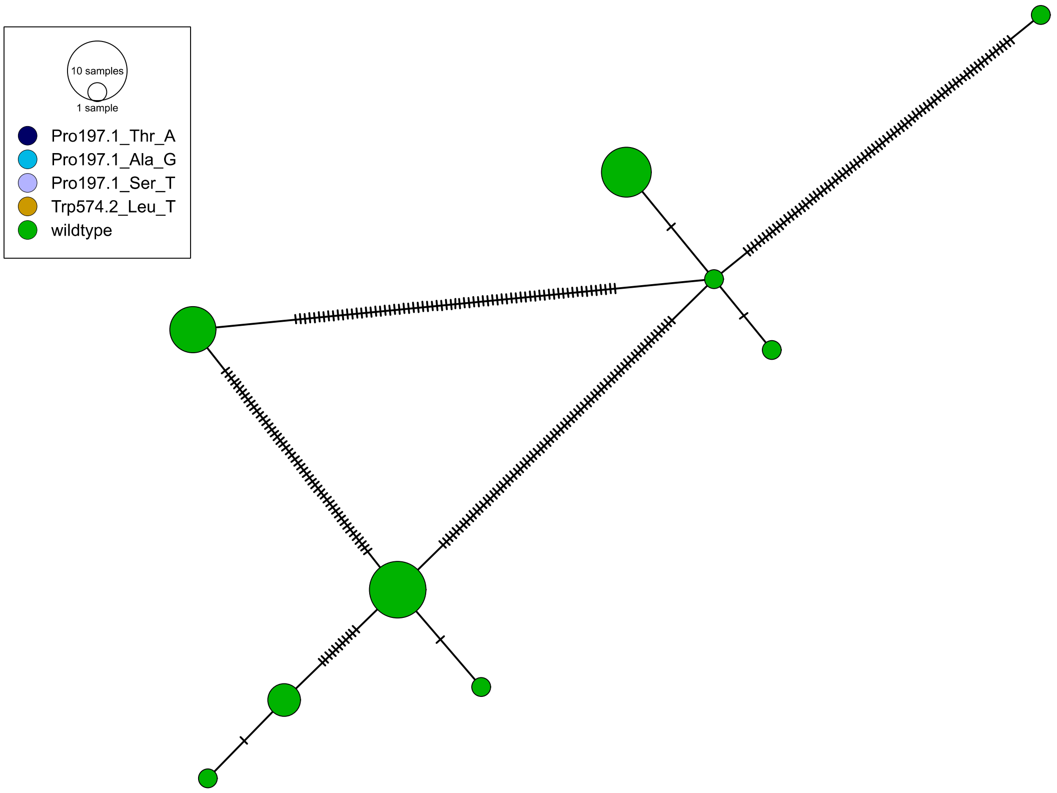

10 samples
1 sample
Pro197.1_Thr_A
Pro197.1_Ala_G
Pro197.1_Ser_T
Trp574.2_Leu_T
wildtype


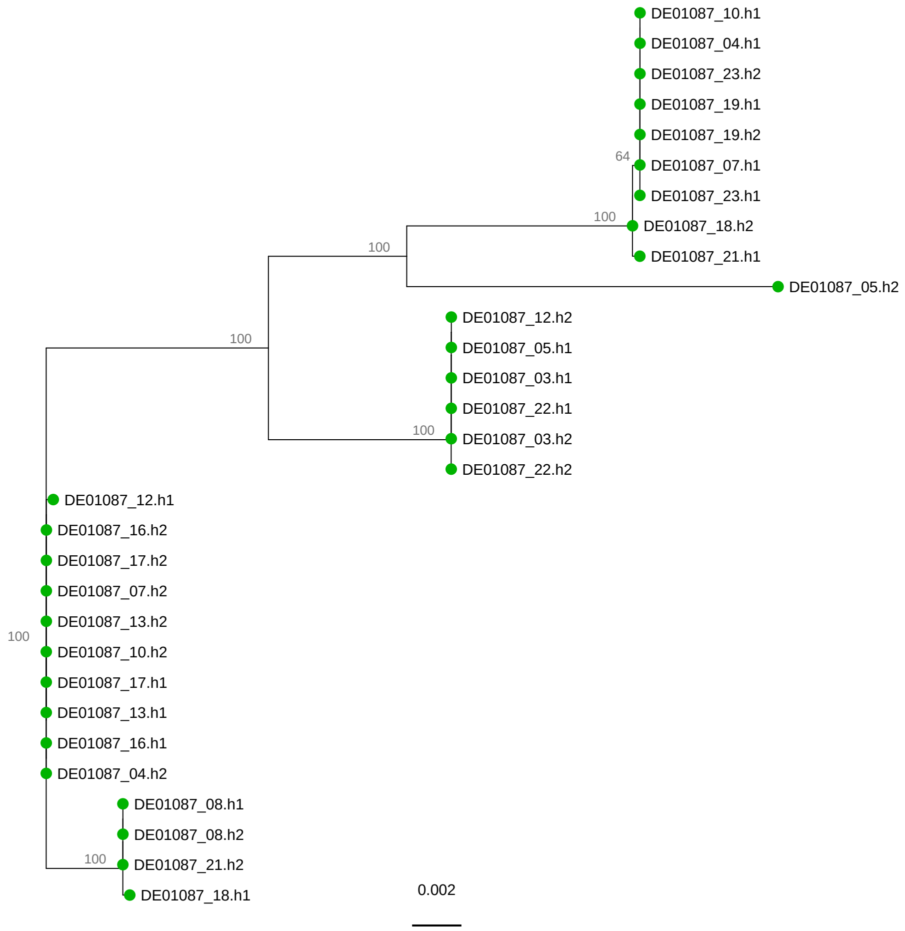

DE01087_10.h1
DE01087_04.h1
DE01087_23.h2
DE01087_19.h1
DE01087_19.h2
64
DE01087_07.h1
DE01087_23.h1
100
DE01087_18.h2
100
DE01087_21.h1
DE01087_05.h2
100
DE01087_12.h2
DE01087_05.h1
DE01087_03.h1
DE01087_22.h1
100
DE01087_03.h2
DE01087_22.h2
DE01087_12.h1
DE01087_16.h2
DE01087_17.h2
DE01087_07.h2
DE01087_13.h2
100
DE01087_10.h2
DE01087_17.h1
DE01087_13.h1
DE01087_16.h1
DE01087_04.h2
DE01087_08.h1
DE01087_08.h2
100
DE01087_21.h2
DE01087_18.h1
0.002

DE01139

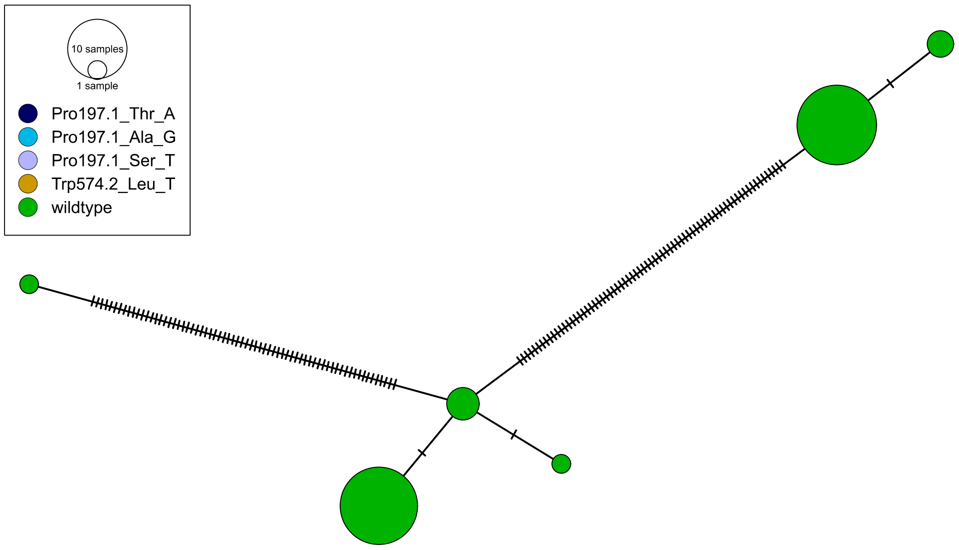
10 samples
1 sample
Pro197.1_Thr_A
Pro197.1_Ala_G
Pro197.1_Ser_T
Trp574.2_Leu_T
wildtype

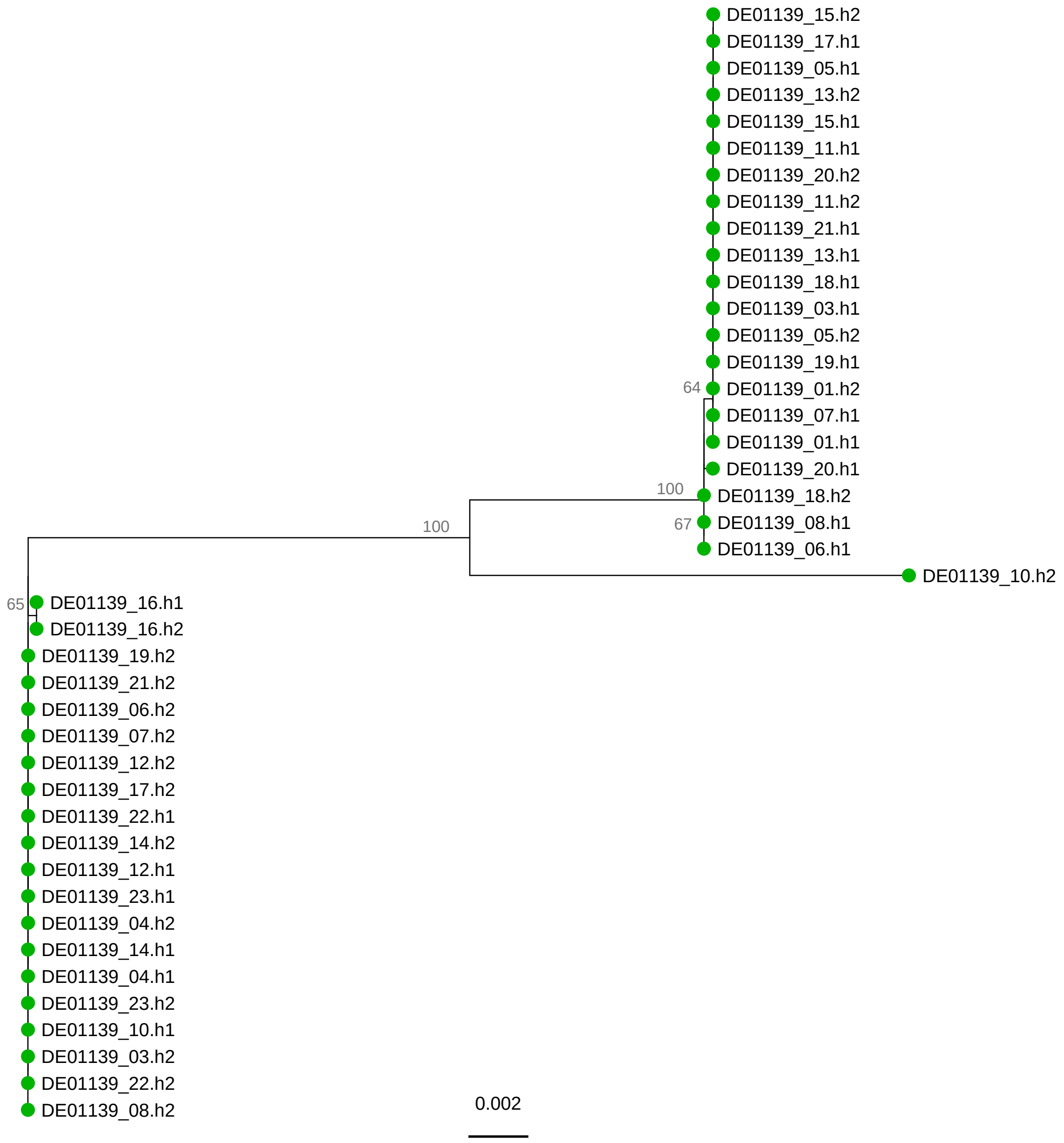
DE01139_15.h2
DE01139_17.h1
DE01139_05.h1
DE01139_13.h2
DE01139_15.h1
DE01139_11.h1
DE01139_20.h2
DE01139_11.h2
DE01139_21.h1
DE01139_13.h1
DE01139_18.h1
DE01139_03.h1
DE01139_05.h2
DE01139_19.h1
64
DE01139_01.h2
DE01139_07.h1
DE01139_01.h1
DE01139_20.h1
100
DE01139_18.h2
67
DE01139_08.h1
DE01139_06.h1
100
DE01139_10.h2
65
DE01139_16.h1
DE01139_16.h2
DE01139_19.h2
DE01139_21.h2
DE01139_06.h2
DE01139_07.h2
DE01139_12.h2
DE01139_17.h2
DE01139_22.h1
DE01139_14.h2
DE01139_12.h1
DE01139_23.h1
DE01139_04.h2
DE01139_14.h1
DE01139_04.h1
DE01139_23.h2
DE01139_10.h1
DE01139_03.h2
DE01139_22.h2
DE01139_08.h2
0.002

# DE01285

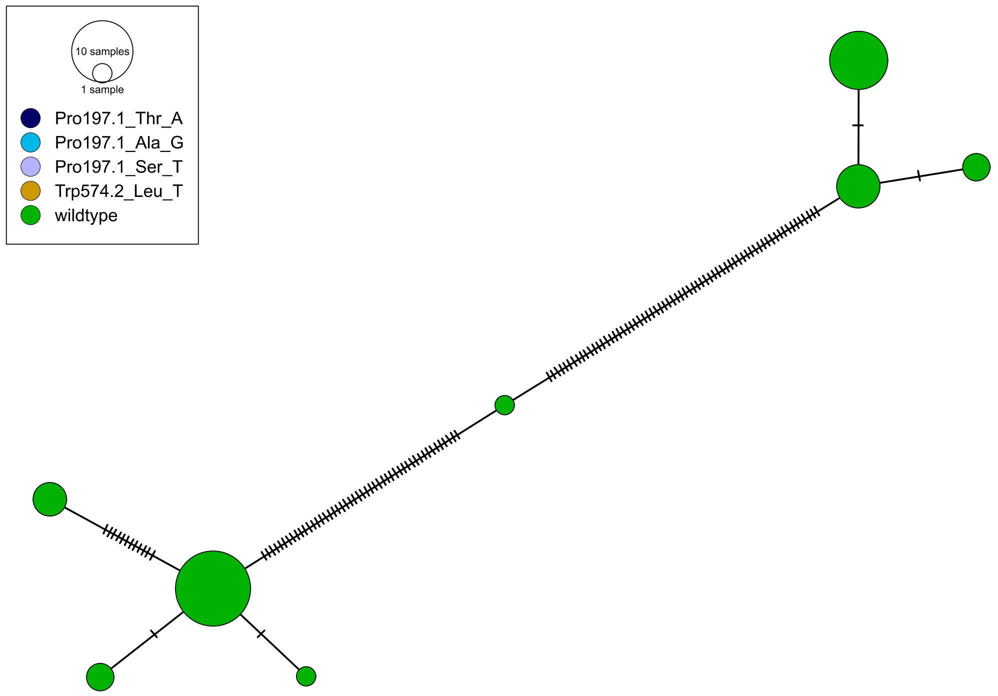
10 samples
1 sample
Pro197.1_Thr_A
Pro197.1_Ala_G
Pro197.1_Ser_T
Trp574.2_Leu_T
wildtype

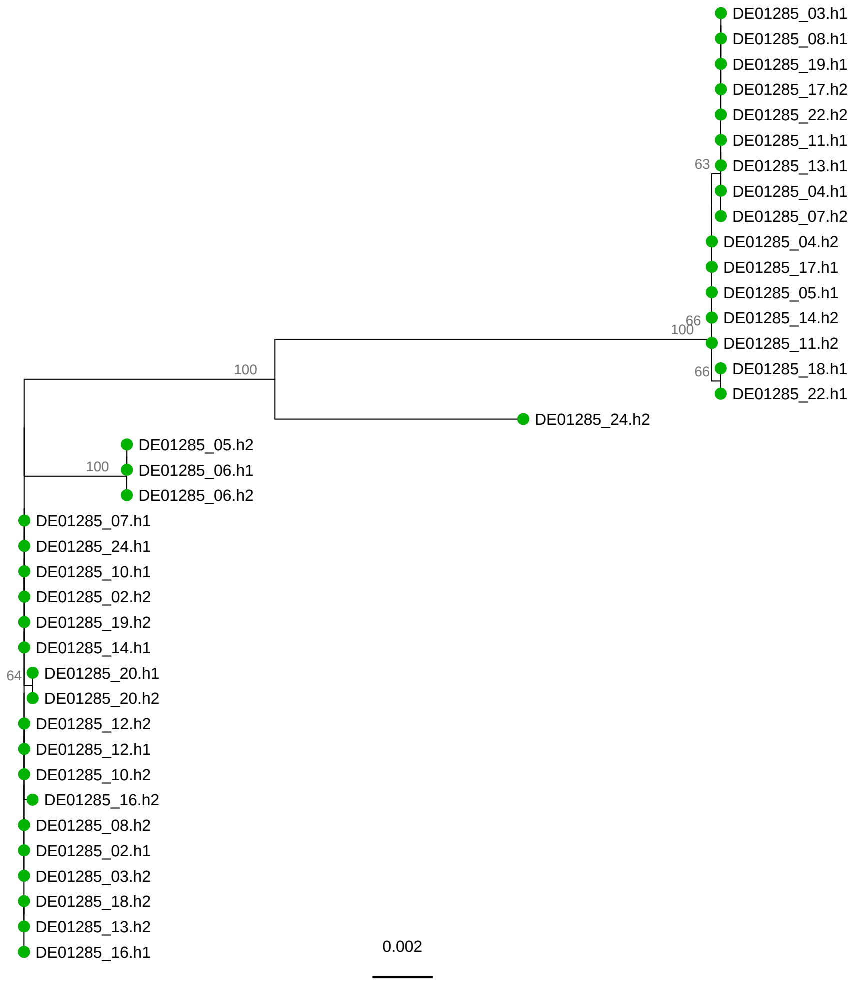
DE01285_03.h1
DE01285_08.h1
DE01285_19.h1
DE01285_17.h2
DE01285_22.h2
DE01285_11.h1
63
DE01285_13.h1
DE01285_04.h1
DE01285_07.h2
DE01285_04.h2
DE01285_17.h1
DE01285_05.h1
66
100
DE01285_14.h2
DE01285_11.h2
66
DE01285_18.h1
DE01285_22.h1
100
DE01285_24.h2
100
DE01285_05.h2
DE01285_06.h1
DE01285_06.h2
DE01285_07.h1
DE01285_24.h1
DE01285_10.h1
DE01285_02.h2
DE01285_19.h2
DE01285_14.h1
64
DE01285_20.h1
DE01285_20.h2
DE01285_12.h2
DE01285_12.h1
DE01285_10.h2
DE01285_16.h2
DE01285_08.h2
DE01285_02.h1
DE01285_03.h2
DE01285_18.h2
DE01285_13.h2
DE01285_16.h1
0.002

DE01321

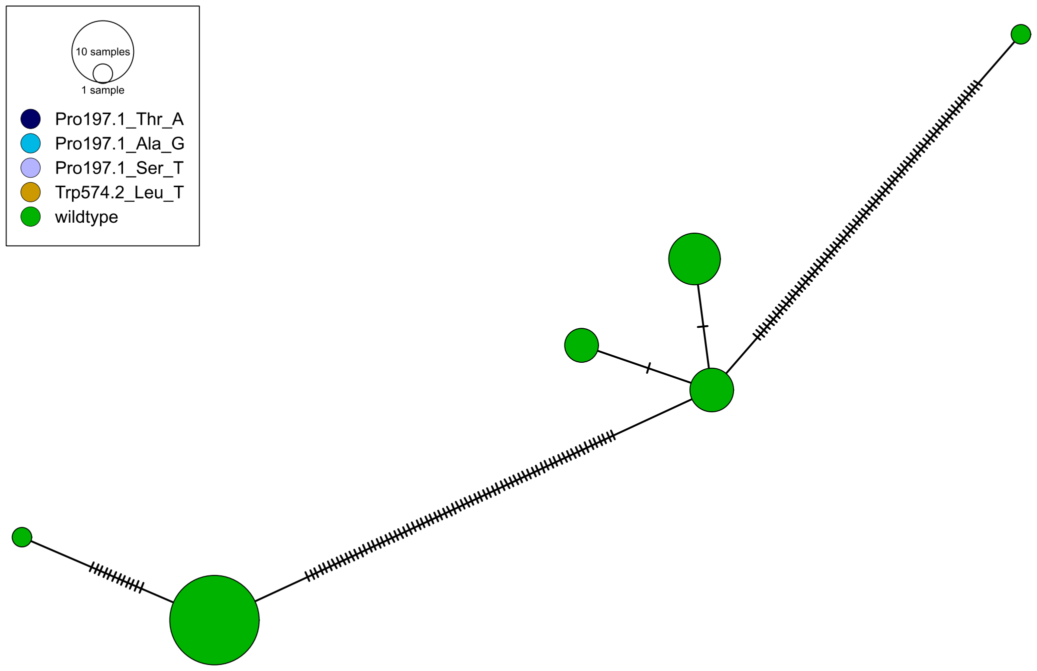
10 samples
1 sample
Pro197.1_Thr_A
Pro197.1_Ala_G
Pro197.1_Ser_T
Trp574.2_Leu_T
wildtype

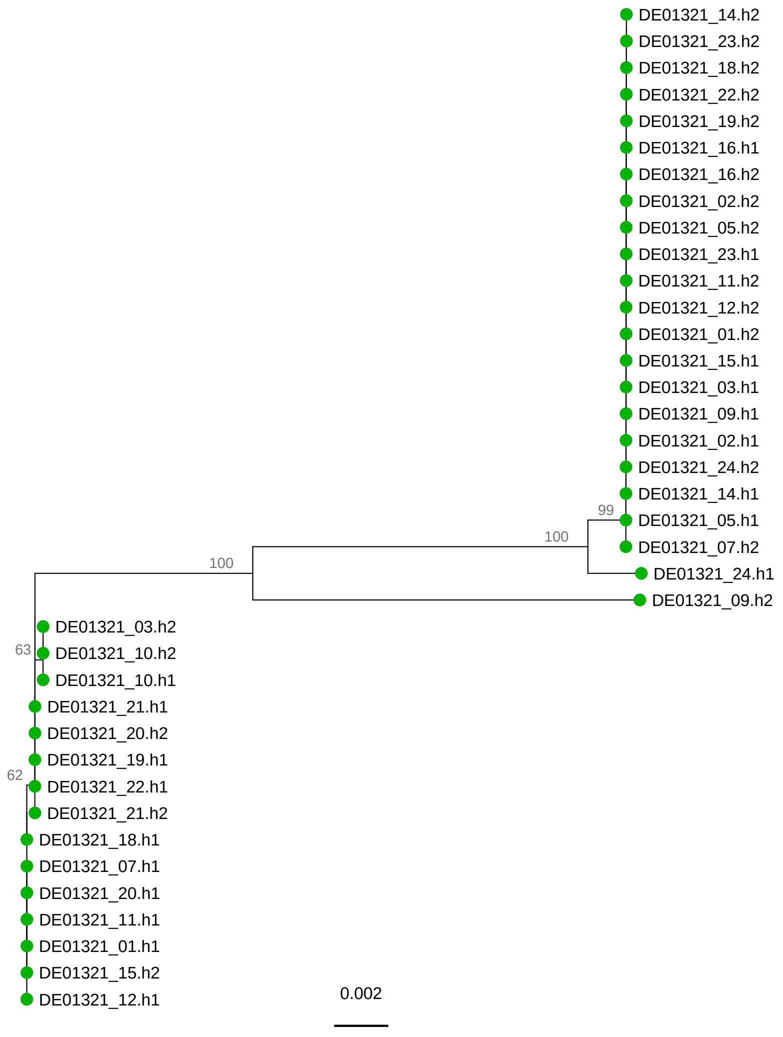
DE01321_14.h2
DE01321_23.h2
DE01321_18.h2
DE01321_22.h2
DE01321_19.h2
DE01321_16.h1
DE01321_16.h2
DE01321_02.h2
DE01321_05.h2
DE01321_23.h1
DE01321_11.h2
DE01321_12.h2
DE01321_01.h2
DE01321_15.h1
DE01321_03.h1
DE01321_09.h1
DE01321_02.h1
DE01321_24.h2
DE01321_14.h1
99
DE01321_05.h1
100
DE01321_07.h2
100
DE01321_24.h1
DE01321_09.h2
DE01321_03.h2
63
DE01321_10.h2
DE01321_10.h1
DE01321_21.h1
DE01321_20.h2
DE01321_19.h1
62
DE01321_22.h1
DE01321_21.h2
DE01321_18.h1
DE01321_07.h1
DE01321_20.h1
DE01321_11.h1
DE01321_01.h1
DE01321_15.h2
DE01321_12.h1
0.002

DE01381

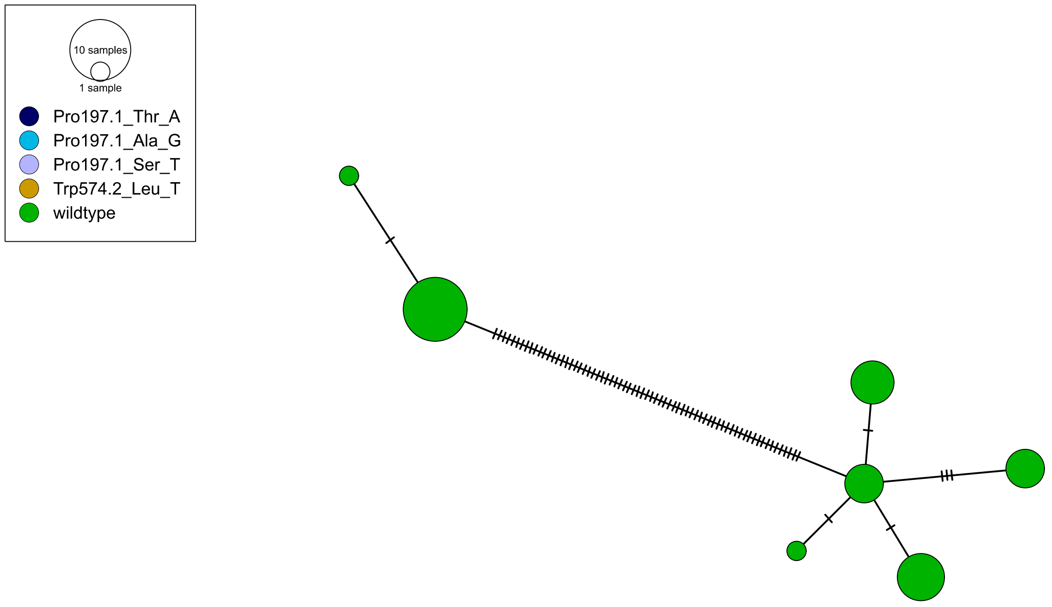
10 samples
1 sample
Pro197.1_Thr_A
Pro197.1_Ala_G
Pro197.1_Ser_T
Trp574.2_Leu_T
wildtype

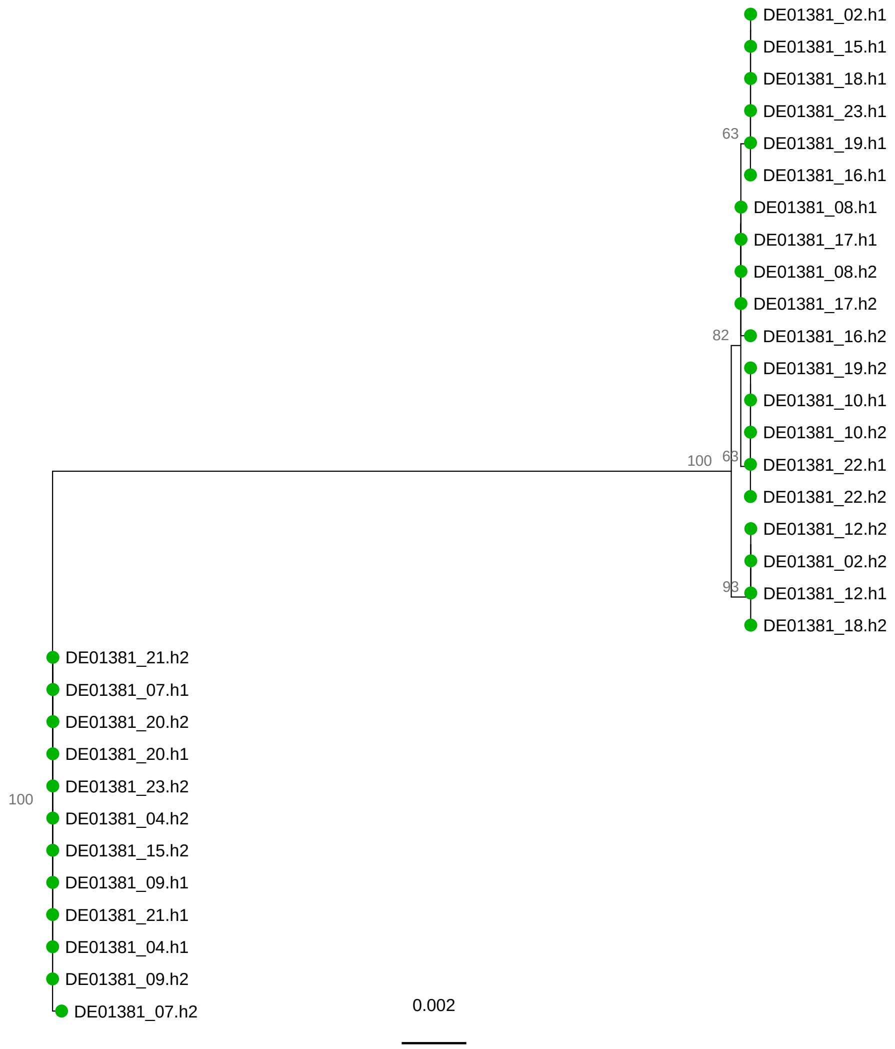
DE01381_02.h1
DE01381_15.h1
DE01381_18.h1
DE01381_23.h1
63
DE01381_19.h1
DE01381_16.h1
DE01381_08.h1
DE01381_17.h1
DE01381_08.h2
DE01381_17.h2
82
DE01381_16.h2
DE01381_19.h2
DE01381_10.h1
DE01381_10.h2
100
63
DE01381_22.h1
DE01381_22.h2
DE01381_12.h2
DE01381_02.h2
93
DE01381_12.h1
DE01381_18.h2
DE01381_21.h2
DE01381_07.h1
DE01381_20.h2
DE01381_20.h1
100
DE01381_23.h2
DE01381_04.h2
DE01381_15.h2
DE01381_09.h1
DE01381_21.h1
DE01381_04.h1
DE01381_09.h2
DE01381_07.h2
0.002

# DE01461

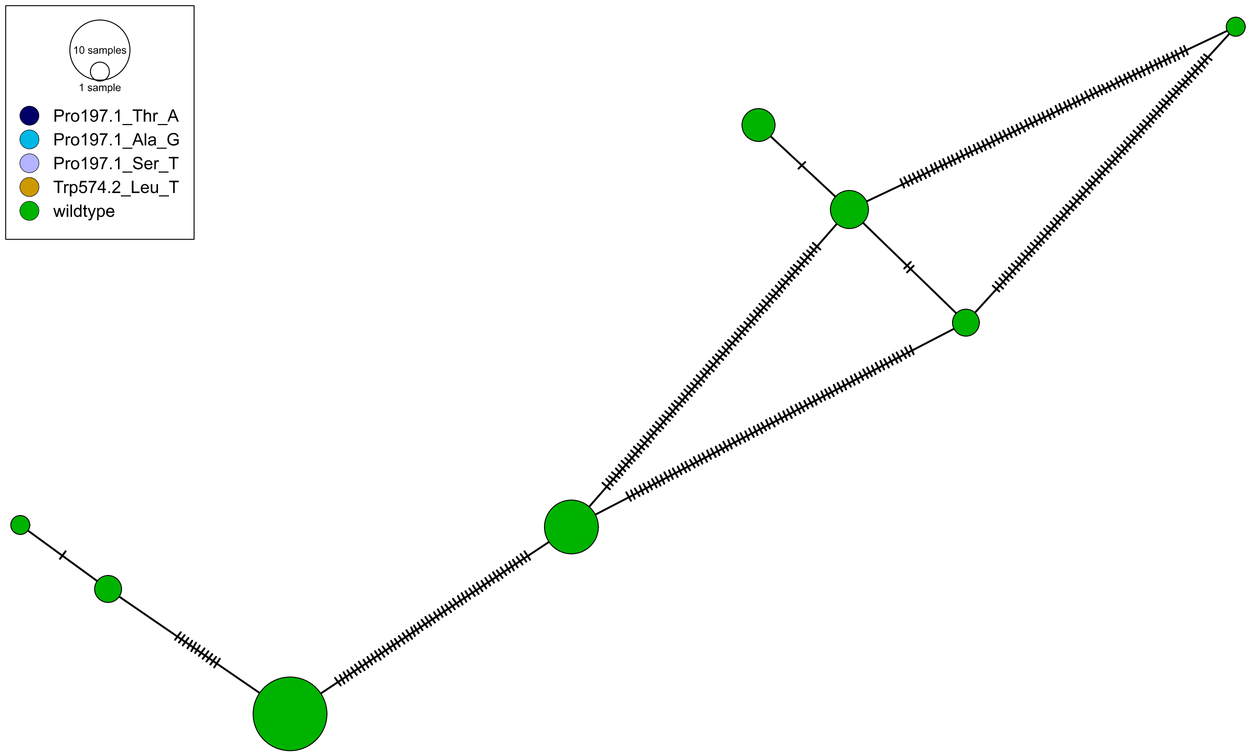
10 samples
1 sample
Pro197.1_Thr_A
Pro197.1_Ala_G
Pro197.1_Ser_T
Trp574.2_Leu_T
wildtype

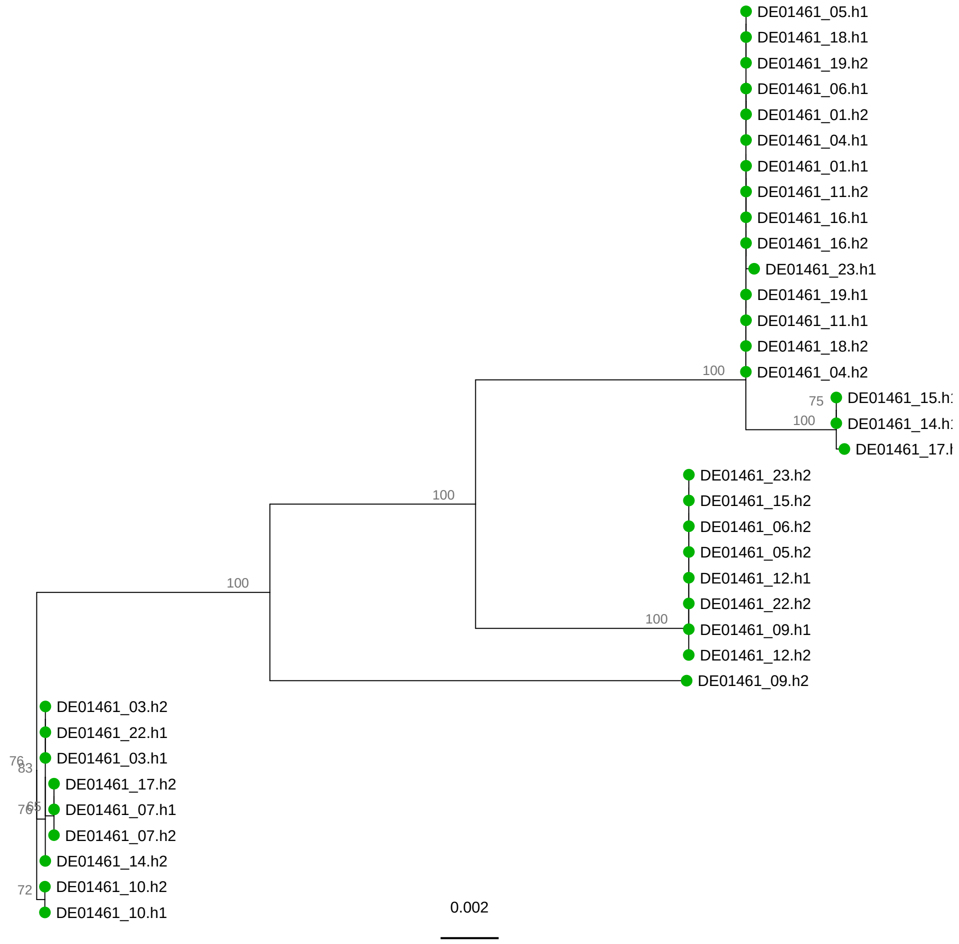
DE01461_05.h1
DE01461_18.h1
DE01461_19.h2
DE01461_06.h1
DE01461_01.h2
DE01461_04.h1
DE01461_01.h1
DE01461_11.h2
DE01461_16.h1
DE01461_16.h2
DE01461_23.h1
DE01461_19.h1
DE01461_11.h1
DE01461_18.h2
DE01461_04.h2
DE01461_15.h
DE01461_14.h
DE01461_17.
DE01461_23.h2
DE01461_15.h2
DE01461_06.h2
DE01461_05.h2
DE01461_12.h1
DE01461_22.h2
DE01461_09.h1
DE01461_12.h2
DE01461_09.h2
DE01461_03.h2
DE01461_22.h1
DE01461_03.h1
DE01461_17.h2
DE01461_07.h1
DE01461_07.h2
DE01461_14.h2
DE01461_10.h2
DE01461_10.h1
100
75
100
100
100
100
76
83
72
0.002

DE01467

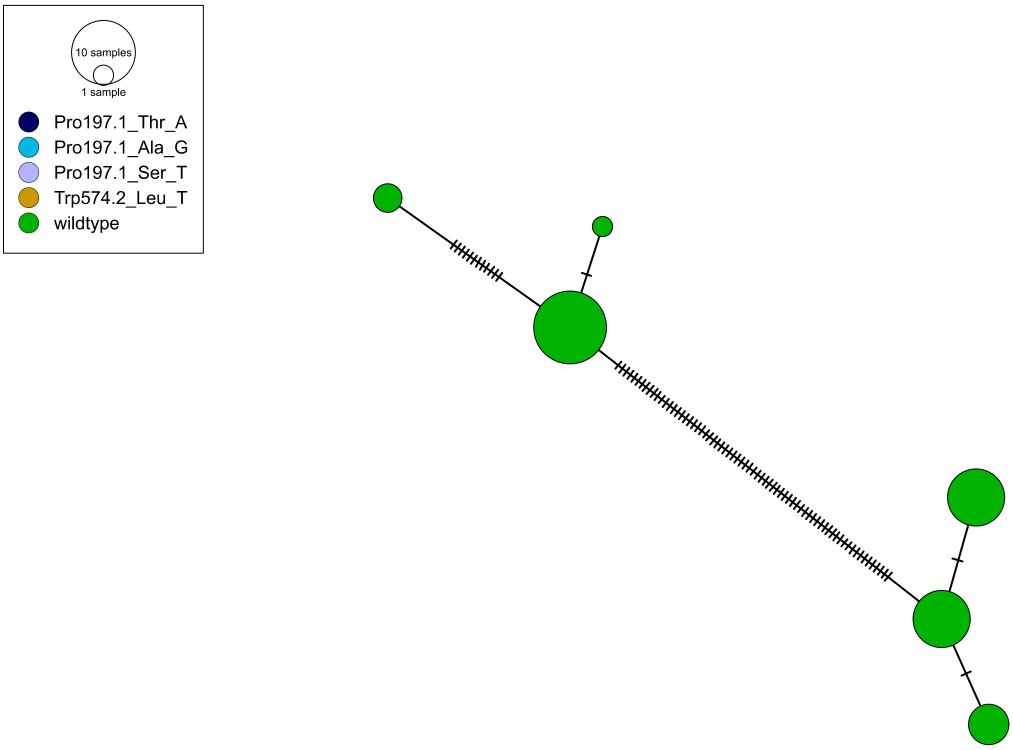
10 samples
1 sample
Pro197.1_Thr_A
Pro197.1_Ala_G
Pro197.1_Ser_T
Trp574.2_Leu_T
wildtype

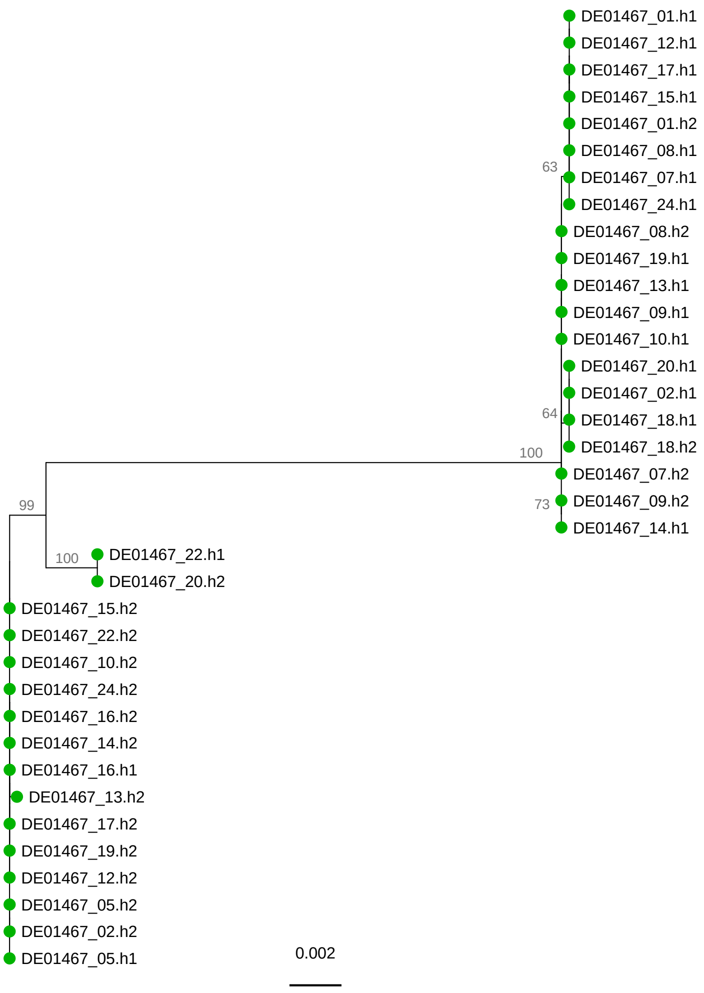
DE01467_01.h1
DE01467_12.h1
DE01467_17.h1
DE01467_15.h1
DE01467_01.h2
DE01467_08.h1
63
DE01467_07.h1
DE01467_24.h1
DE01467_08.h2
DE01467_19.h1
DE01467_13.h1
DE01467_09.h1
DE01467_10.h1
DE01467_20.h1
DE01467_02.h1
64
DE01467_18.h1
DE01467_18.h2
100
DE01467_07.h2
99
73
DE01467_09.h2
DE01467_14.h1
100
DE01467_22.h1
DE01467_20.h2
DE01467_15.h2
DE01467_22.h2
DE01467_10.h2
DE01467_24.h2
DE01467_16.h2
DE01467_14.h2
DE01467_16.h1
DE01467_13.h2
DE01467_17.h2
DE01467_19.h2
DE01467_12.h2
DE01467_05.h2
DE01467_02.h2
DE01467_05.h1
0.002

# DE01580

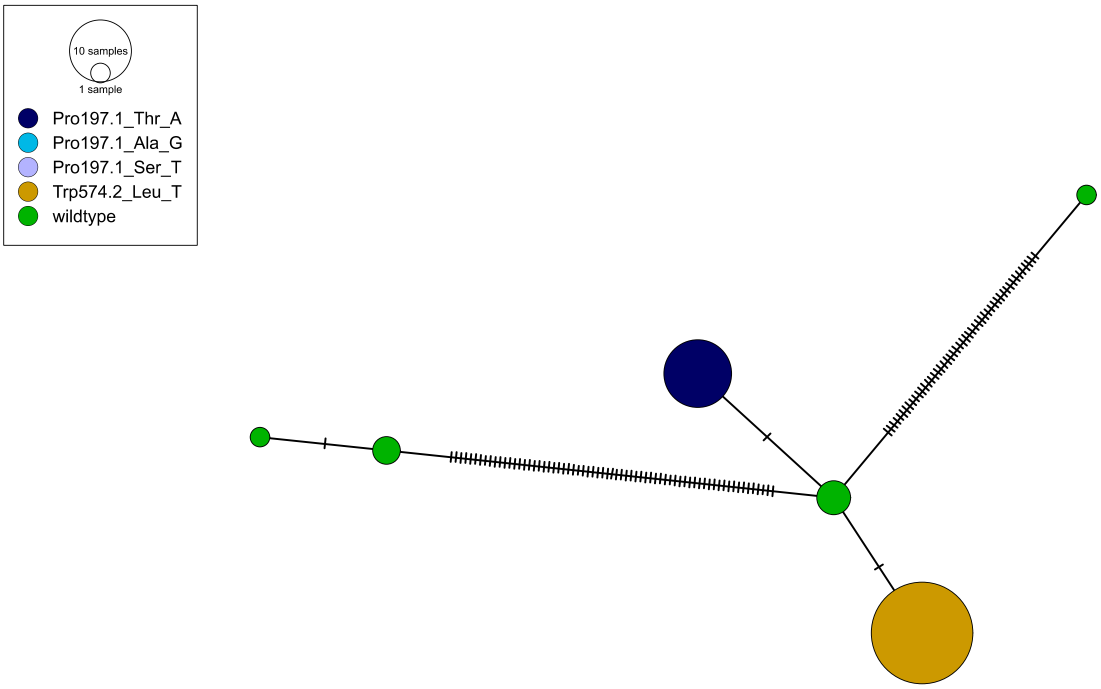
10 samples
1 sample
Pro197.1_Thr_A
Pro197.1_Ala_G
Pro197.1_Ser_T
Trp574.2_Leu_T
wildtype

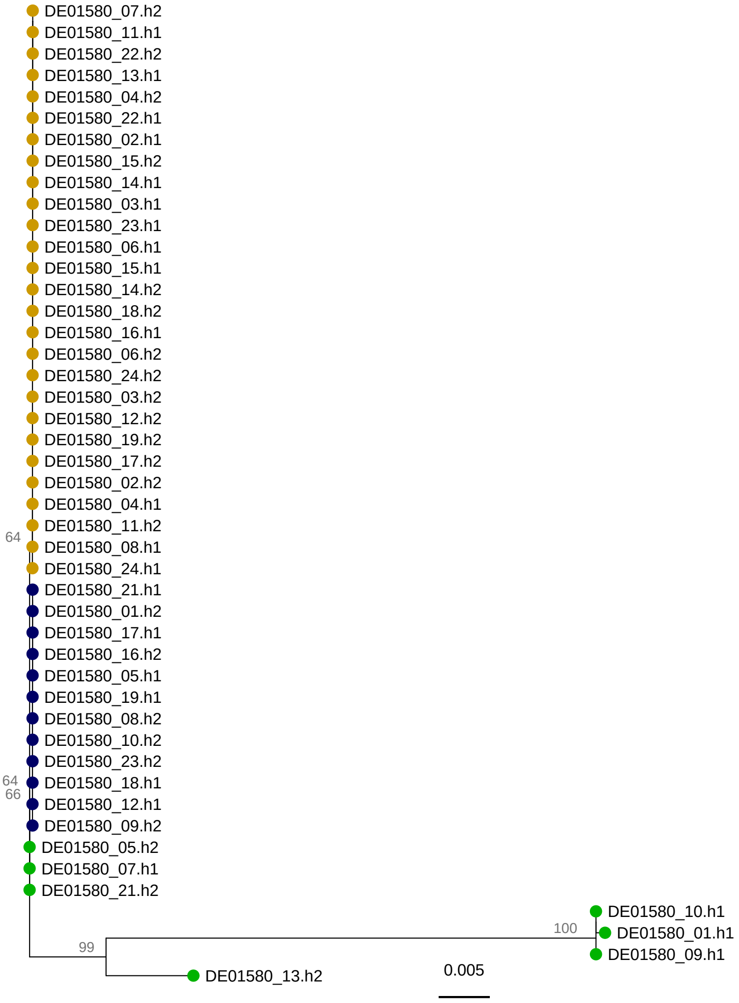
DE01580_07.h2
DE01580_11.h1
DE01580_22.h2
DE01580_13.h1
DE01580_04.h2
DE01580_22.h1
DE01580_02.h1
DE01580_15.h2
DE01580_14.h1
DE01580_03.h1
DE01580_23.h1
DE01580_06.h1
DE01580_15.h1
DE01580_14.h2
DE01580_18.h2
DE01580_16.h1
DE01580_06.h2
DE01580_24.h2
DE01580_03.h2
DE01580_12.h2
DE01580_19.h2
DE01580_17.h2
DE01580_02.h2
DE01580_04.h1
DE01580_11.h2
64
DE01580_08.h1
DE01580_24.h1
DE01580_21.h1
DE01580_01.h2
DE01580_17.h1
DE01580_16.h2
DE01580_05.h1
DE01580_19.h1
DE01580_08.h2
DE01580_10.h2
DE01580_23.h2
64
DE01580_18.h1
66
DE01580_12.h1
DE01580_09.h2
DE01580_05.h2
DE01580_07.h1
DE01580_21.h2
DE01580_10.h1
100
DE01580_01.h1
99
DE01580_09.h1
DE01580_13.h2
0.005

DE01712

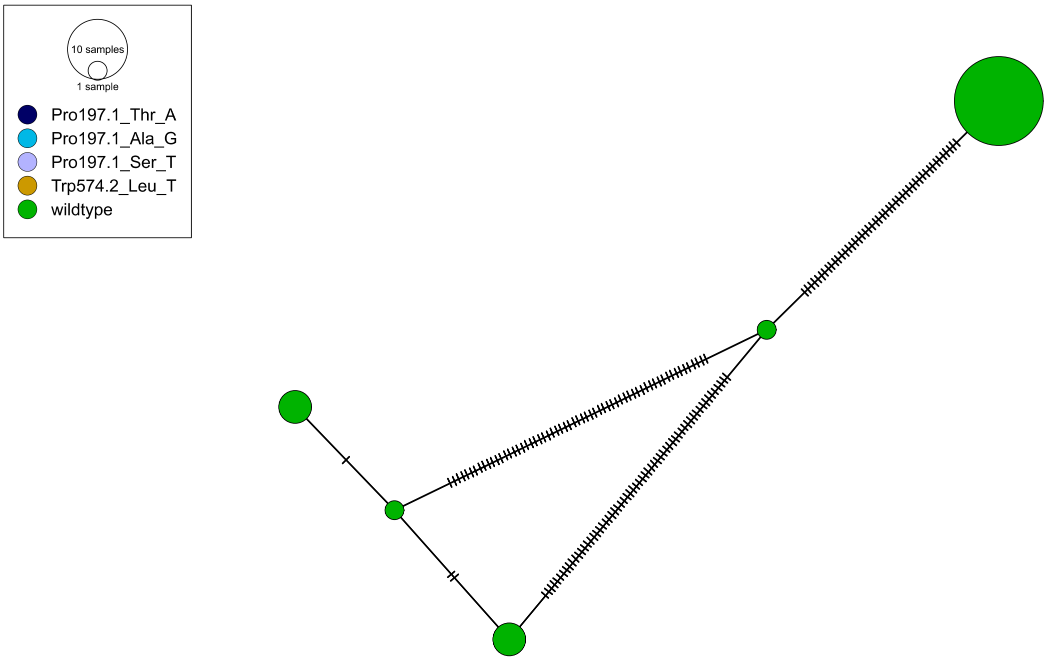
10 samples
1 sample
Pro197.1_Thr_A
Pro197.1_Ala_G
Pro197.1_Ser_T
Trp574.2_Leu_T
wildtype

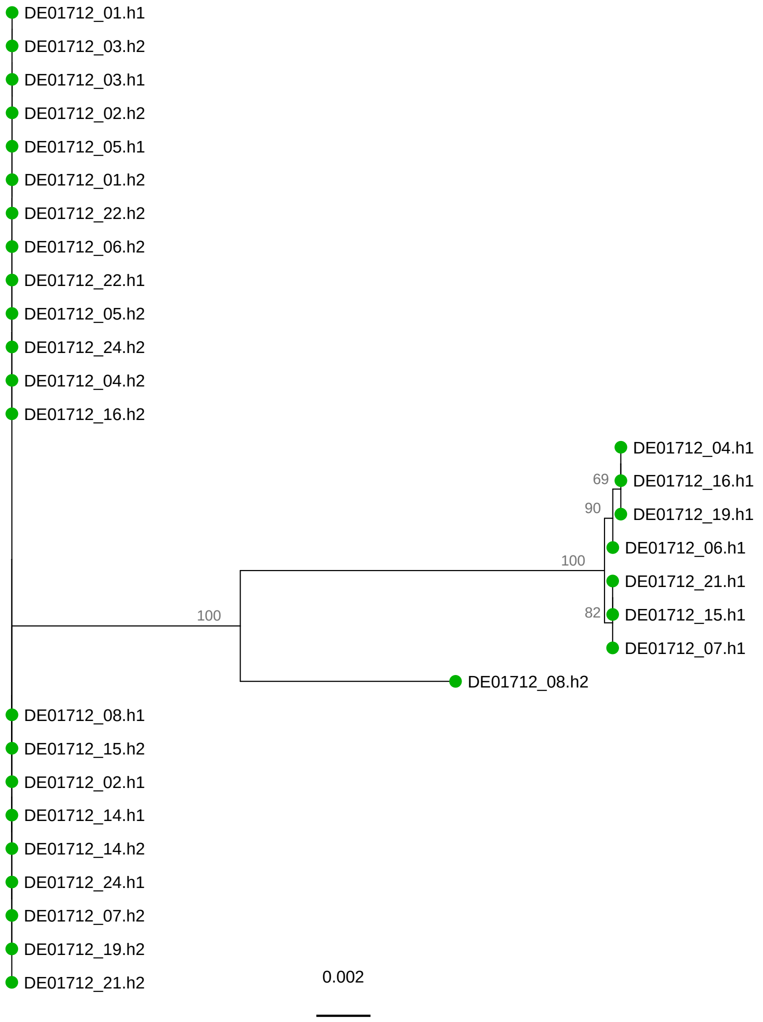
DE01712_01.h1
DE01712_03.h2
DE01712_03.h1
DE01712_02.h2
DE01712_05.h1
DE01712_01.h2
DE01712_22.h2
DE01712_06.h2
DE01712_22.h1
DE01712_05.h2
DE01712_24.h2
DE01712_04.h2
DE01712_16.h2
DE01712_04.h1
69
DE01712_16.h1
90
DE01712_19.h1
DE01712_06.h1
100
DE01712_21.h1
100
82
DE01712_15.h1
DE01712_07.h1
DE01712_08.h2
DE01712_08.h1
DE01712_15.h2
DE01712_02.h1
DE01712_14.h1
DE01712_14.h2
DE01712_24.h1
DE01712_07.h2
DE01712_19.h2
DE01712_21.h2
0.002

## DE06506

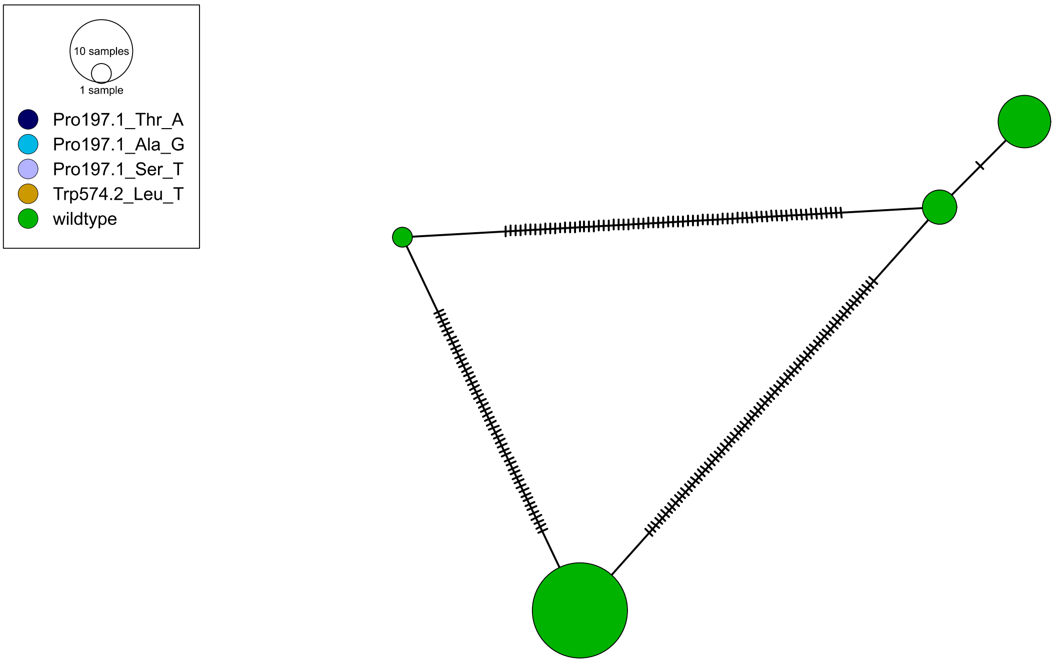

10 samples
1 sample
Pro197.1_Thr_A
Pro197.1_Ala_G
Pro197.1_Ser_T
Trp574.2_Leu_T
wildtype


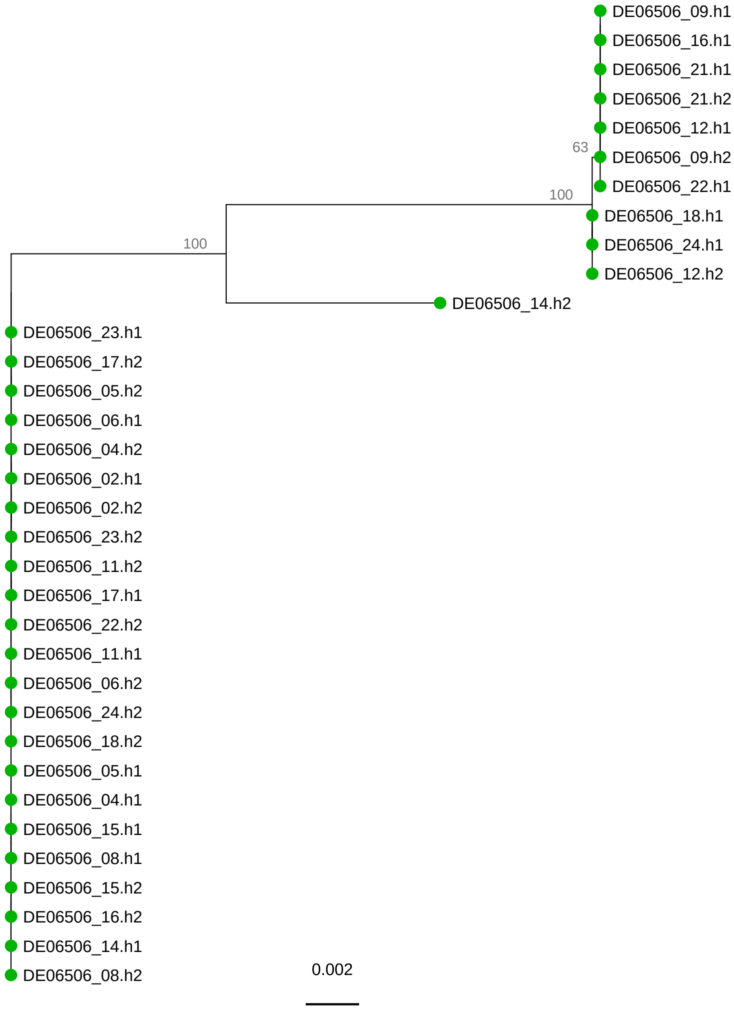

DE06506_09.h1
DE06506_16.h1
DE06506_21.h1
DE06506_21.h2
DE06506_12.h1
63
DE06506_09.h2
DE06506_22.h1
100
DE06506_18.h1
DE06506_24.h1
DE06506_12.h2
100
DE06506_14.h2
DE06506_23.h1
DE06506_17.h2
DE06506_05.h2
DE06506_06.h1
DE06506_04.h2
DE06506_02.h1
DE06506_02.h2
DE06506_23.h2
DE06506_11.h2
DE06506_17.h1
DE06506_22.h2
DE06506_11.h1
DE06506_06.h2
DE06506_24.h2
DE06506_18.h2
DE06506_05.h1
DE06506_04.h1
DE06506_15.h1
DE06506_08.h1
DE06506_15.h2
DE06506_16.h2
DE06506_14.h1
DE06506_08.h2
0.002

# DE07299

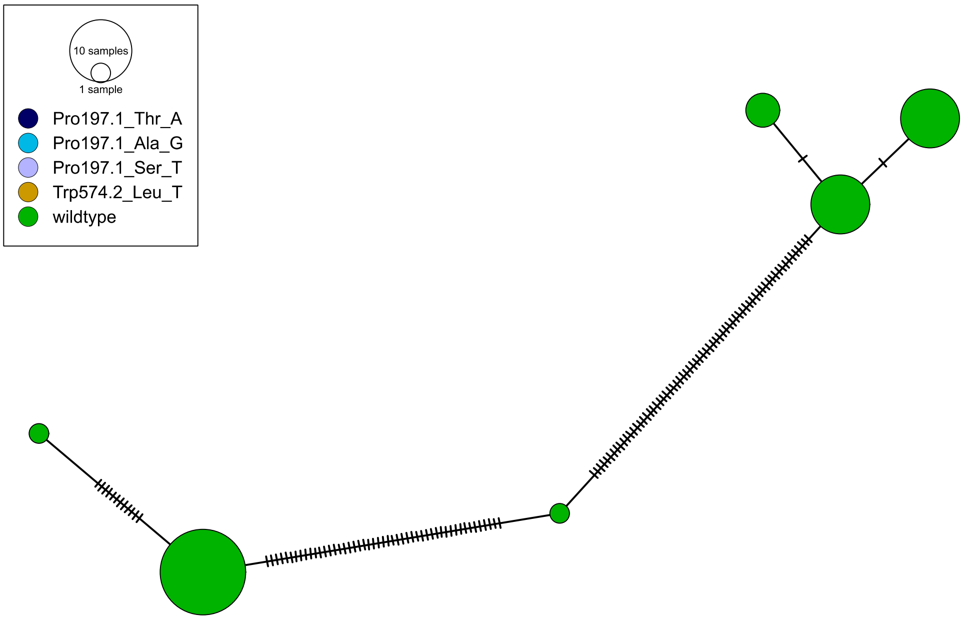

10 samples
1 sample
Pro197.1_Thr_A
Pro197.1_Ala_G
Pro197.1_Ser_T
Trp574.2_Leu_T
wildtype


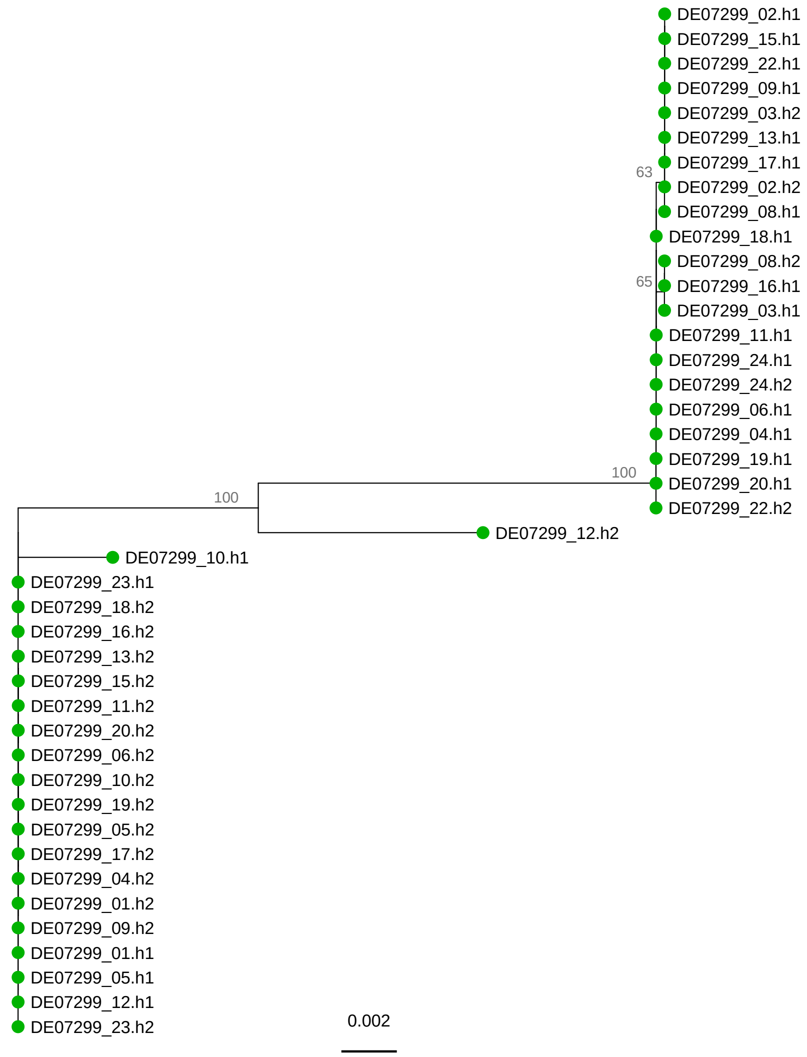

DE07299_02.h1
DE07299_15.h1
DE07299_22.h1
DE07299_09.h1
DE07299_03.h2
DE07299_13.h1
DE07299_17.h1
63
DE07299_02.h2
DE07299_08.h1
DE07299_18.h1
DE07299_08.h2
65
DE07299_16.h1
DE07299_03.h1
DE07299_11.h1
DE07299_24.h1
DE07299_24.h2
DE07299_06.h1
DE07299_04.h1
DE07299_19.h1
100
DE07299_20.h1
DE07299_22.h2
100
DE07299_12.h2
DE07299_10.h1
DE07299_23.h1
DE07299_18.h2
DE07299_16.h2
DE07299_13.h2
DE07299_15.h2
DE07299_11.h2
DE07299_20.h2
DE07299_06.h2
DE07299_10.h2
DE07299_19.h2
DE07299_05.h2
DE07299_17.h2
DE07299_04.h2
DE07299_01.h2
DE07299_09.h2
DE07299_01.h1
DE07299_05.h1
DE07299_12.h1
DE07299_23.h2
0.002

# DE07323

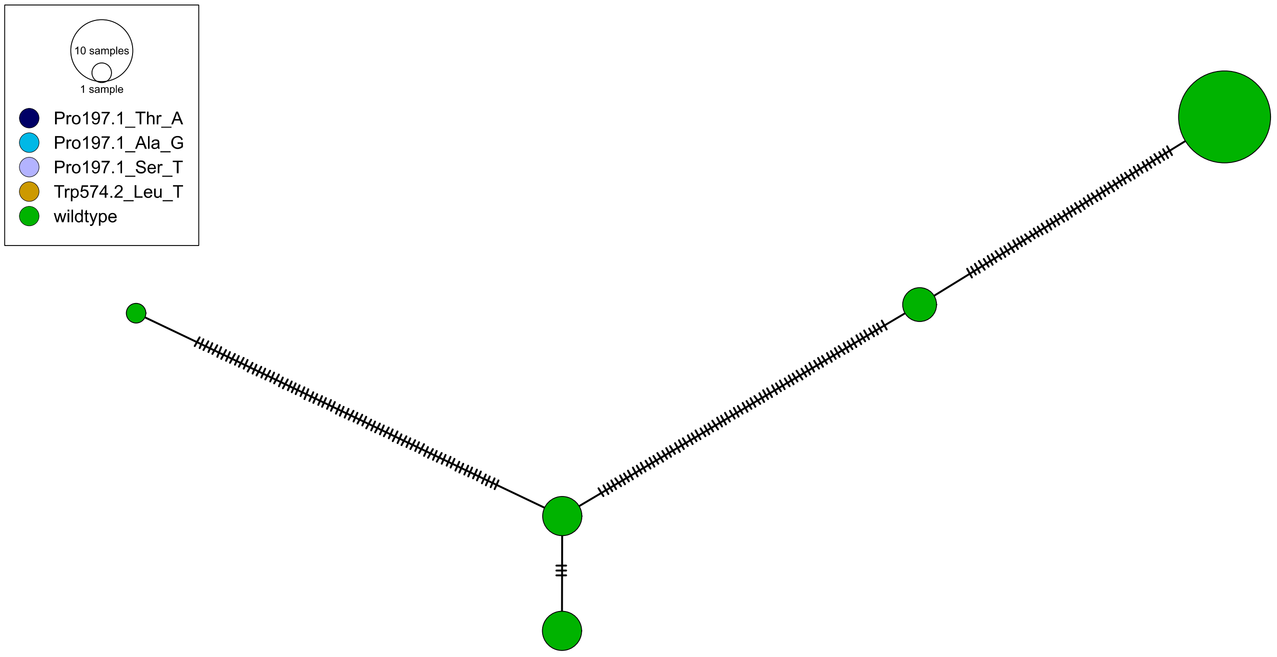
10 samples
1 sample
Pro197.1_Thr_A
Pro197.1_Ala_G
Pro197.1_Ser_T
Trp574.2_Leu_T
wildtype

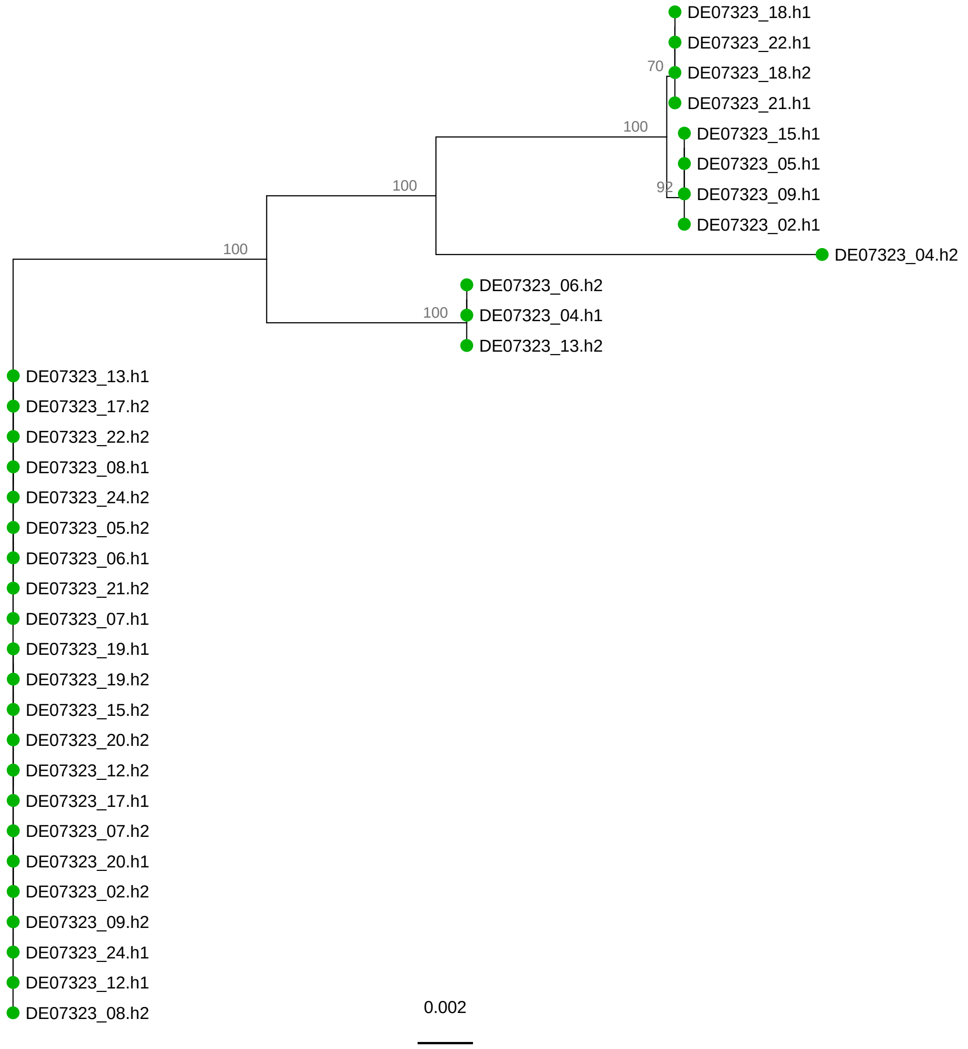
DE07323_18.h1
DE07323_22.h1
70
DE07323_18.h2
DE07323_21.h1
100
DE07323_15.h1
DE07323_05.h1
100
92
DE07323_09.h1
DE07323_02.h1
100
DE07323_04.h2
DE07323_06.h2
100
DE07323_04.h1
DE07323_13.h2
DE07323_13.h1
DE07323_17.h2
DE07323_22.h2
DE07323_08.h1
DE07323_24.h2
DE07323_05.h2
DE07323_06.h1
DE07323_21.h2
DE07323_07.h1
DE07323_19.h1
DE07323_19.h2
DE07323_15.h2
DE07323_20.h2
DE07323_12.h2
DE07323_17.h1
DE07323_07.h2
DE07323_20.h1
DE07323_02.h2
DE07323_09.h2
DE07323_24.h1
DE07323_12.h1
DE07323_08.h2
0.002

# FR01125

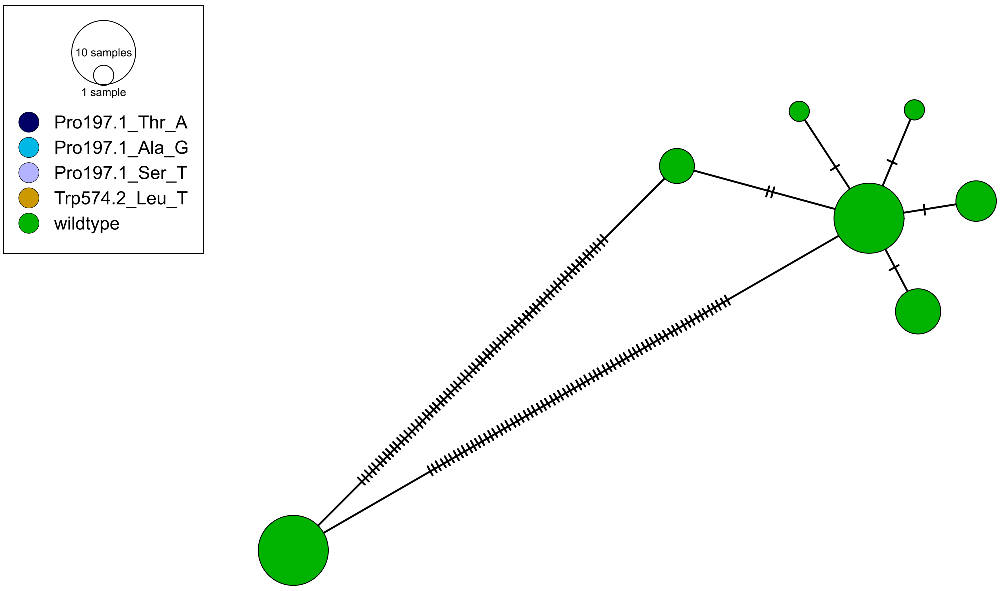
10 samples
1 sample
Pro197.1_Thr_A
Pro197.1_Ala_G
Pro197.1_Ser_T
Trp574.2_Leu_T
wildtype

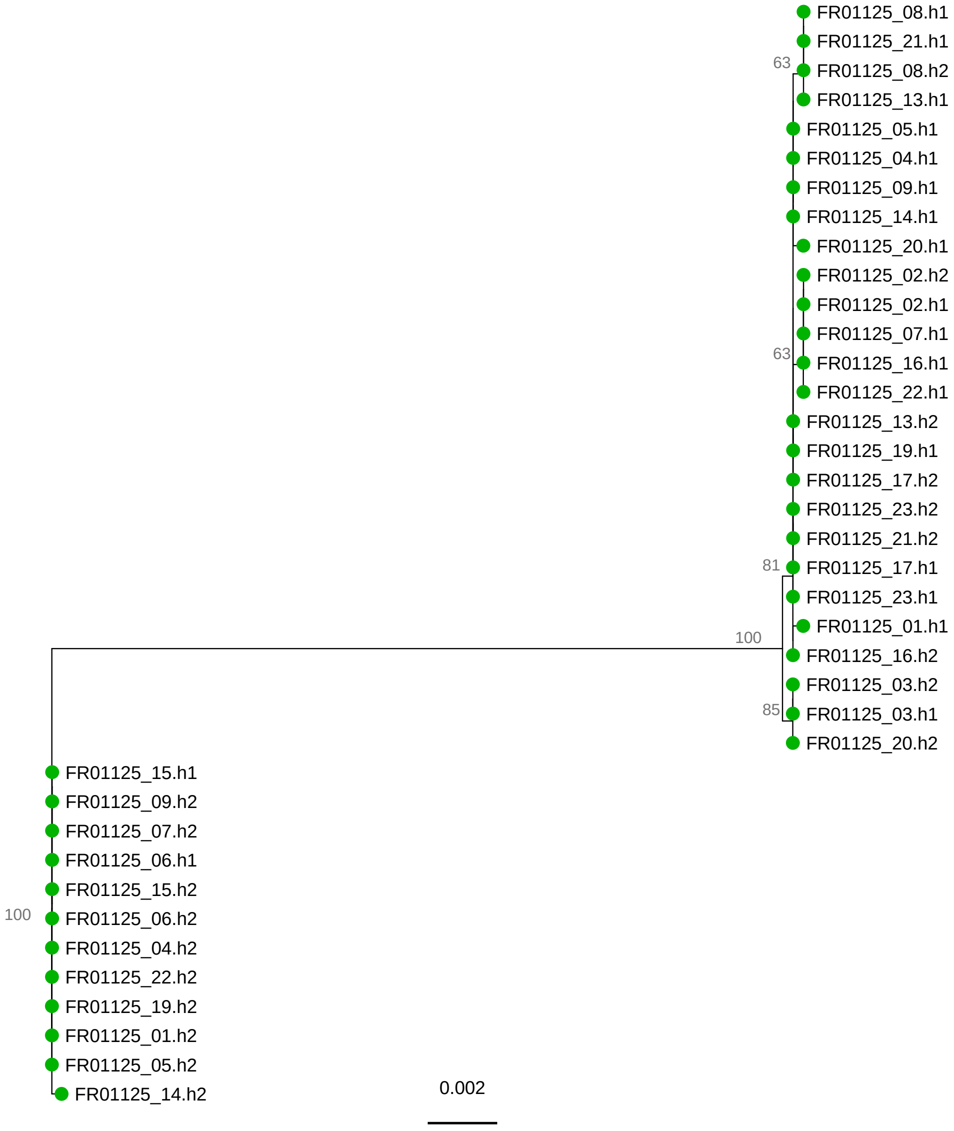
FR01125_08.h1
FR01125_21.h1
63
FR01125_08.h2
FR01125_13.h1
FR01125_05.h1
FR01125_04.h1
FR01125_09.h1
FR01125_14.h1
FR01125_20.h1
FR01125_02.h2
FR01125_02.h1
FR01125_07.h1
63
FR01125_16.h1
FR01125_22.h1
FR01125_13.h2
FR01125_19.h1
FR01125_17.h2
FR01125_23.h2
FR01125_21.h2
81
FR01125_17.h1
FR01125_23.h1
100
FR01125_01.h1
FR01125_16.h2
FR01125_03.h2
85
FR01125_03.h1
FR01125_20.h2
FR01125_15.h1
FR01125_09.h2
FR01125_07.h2
FR01125_06.h1
FR01125_15.h2
100
FR01125_06.h2
FR01125_04.h2
FR01125_22.h2
FR01125_19.h2
FR01125_01.h2
FR01125_05.h2
FR01125_14.h2
0.002

# FR01297

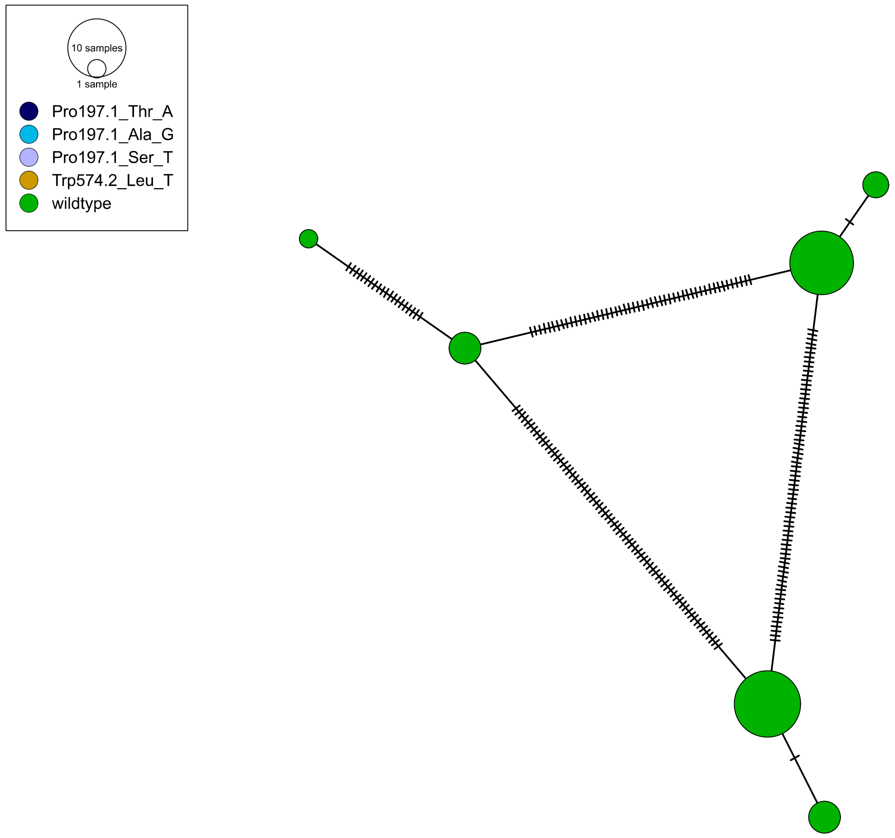
10 samples
1 sample
Pro197.1_Thr_A
Pro197.1_Ala_G
Pro197.1_Ser_T
Trp574.2_Leu_T
wildtype

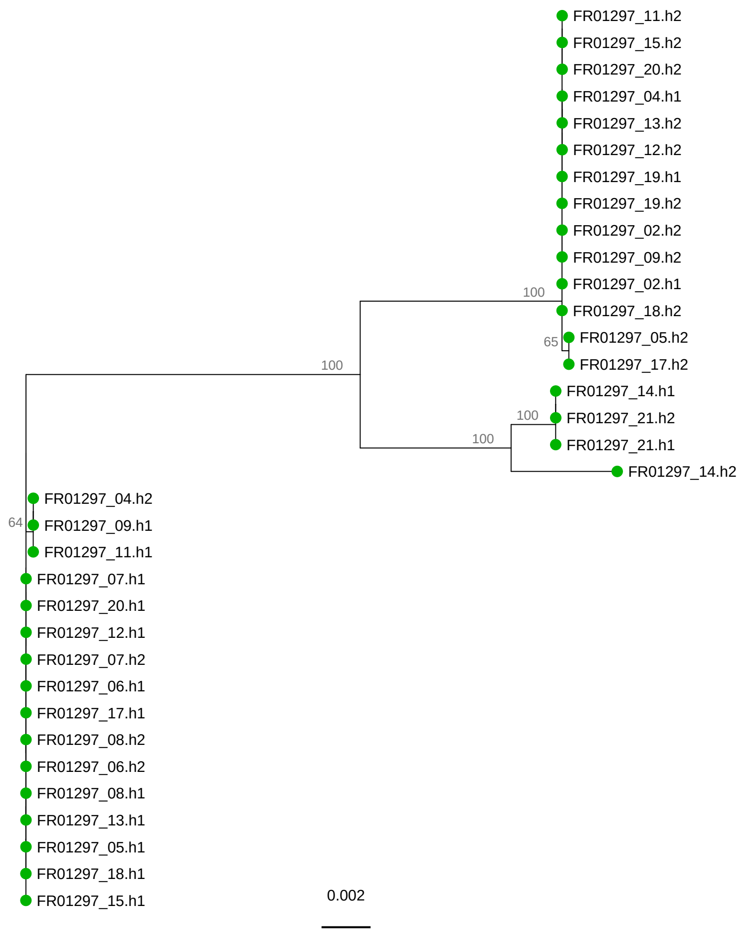
FR01297_11.h2
FR01297_15.h2
FR01297_20.h2
FR01297_04.h1
FR01297_13.h2
FR01297_12.h2
FR01297_19.h1
FR01297_19.h2
FR01297_02.h2
FR01297_09.h2
FR01297_02.h1
100
FR01297_18.h2
65
FR01297_05.h2
100
FR01297_17.h2
FR01297_14.h1
100
FR01297_21.h2
100
FR01297_21.h1
FR01297_14.h2
FR01297_04.h2
64
FR01297_09.h1
FR01297_11.h1
FR01297_07.h1
FR01297_20.h1
FR01297_12.h1
FR01297_07.h2
FR01297_06.h1
FR01297_17.h1
FR01297_08.h2
FR01297_06.h2
FR01297_08.h1
FR01297_13.h1
FR01297_05.h1
FR01297_18.h1
FR01297_15.h1
0.002

FR01331

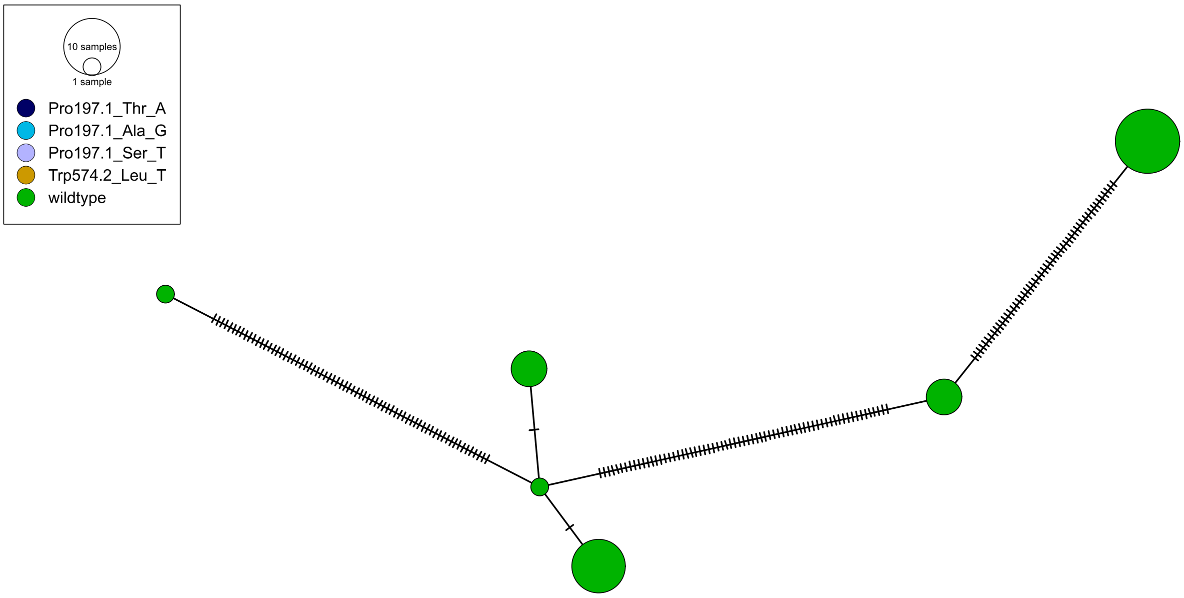
10 samples
1 sample
Pro197.1_Thr_A
Pro197.1_Ala_G
Pro197.1_Ser_T
Trp574.2_Leu_T
wildtype

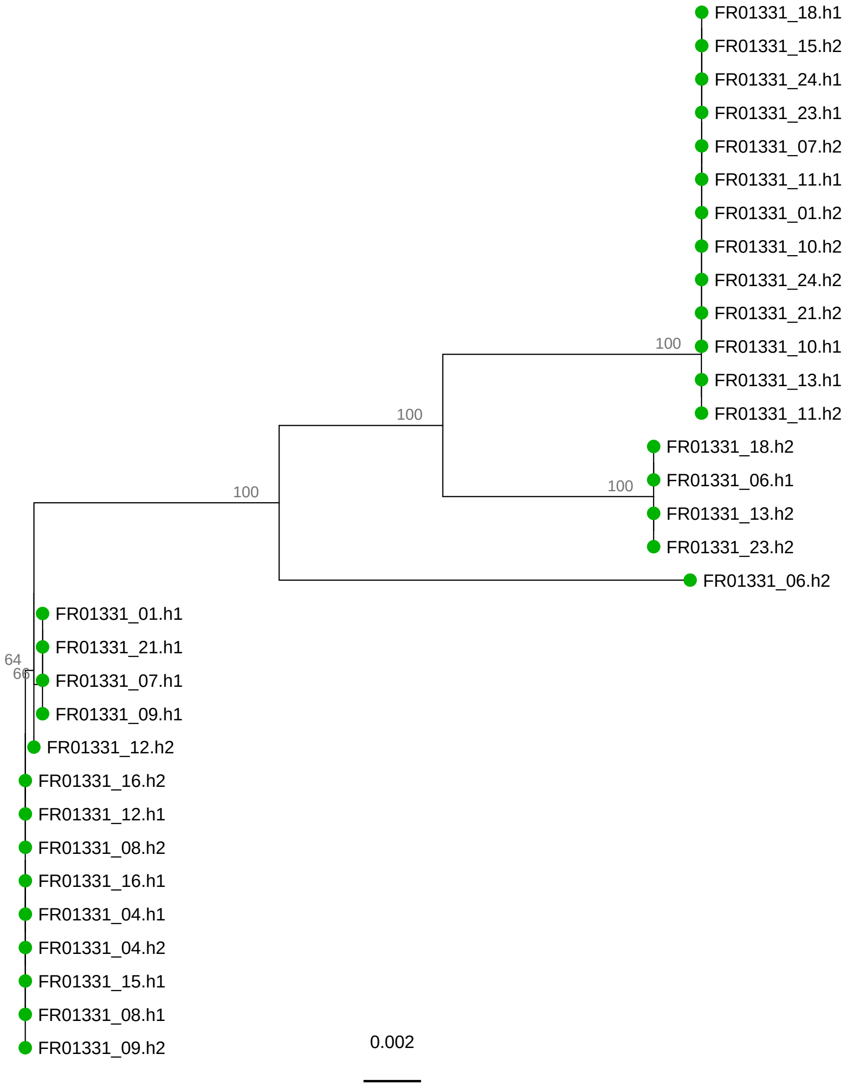
FR01331_18.h1
FR01331_15.h2
FR01331_24.h1
FR01331_23.h1
FR01331_07.h2
FR01331_11.h1
FR01331_01.h2
FR01331_10.h2
FR01331_24.h2
FR01331_21.h2
100
FR01331_10.h1
FR01331_13.h1
100
FR01331_11.h2
FR01331_18.h2
100
FR01331_06.h1
100
FR01331_13.h2
FR01331_23.h2
FR01331_06.h2
FR01331_01.h1
FR01331_21.h1
64
66
FR01331_07.h1
FR01331_09.h1
FR01331_12.h2
FR01331_16.h2
FR01331_12.h1
FR01331_08.h2
FR01331_16.h1
FR01331_04.h1
FR01331_04.h2
FR01331_15.h1
FR01331_08.h1
FR01331_09.h2
0.002

FR01386

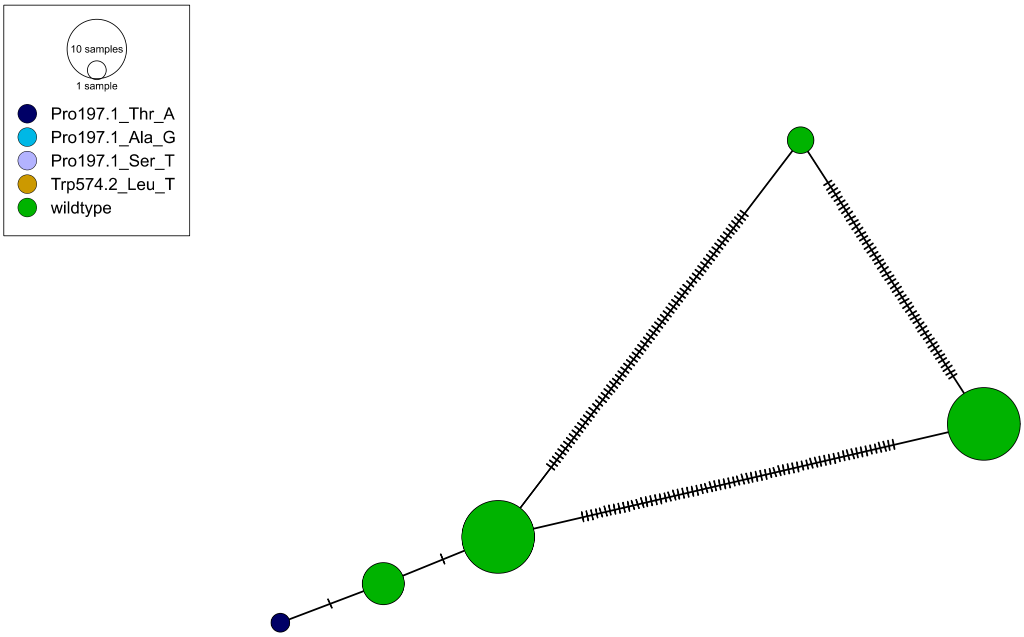
10 samples
1 sample
Pro197.1_Thr_A
Pro197.1_Ala_G
Pro197.1_Ser_T
Trp574.2_Leu_T
wildtype

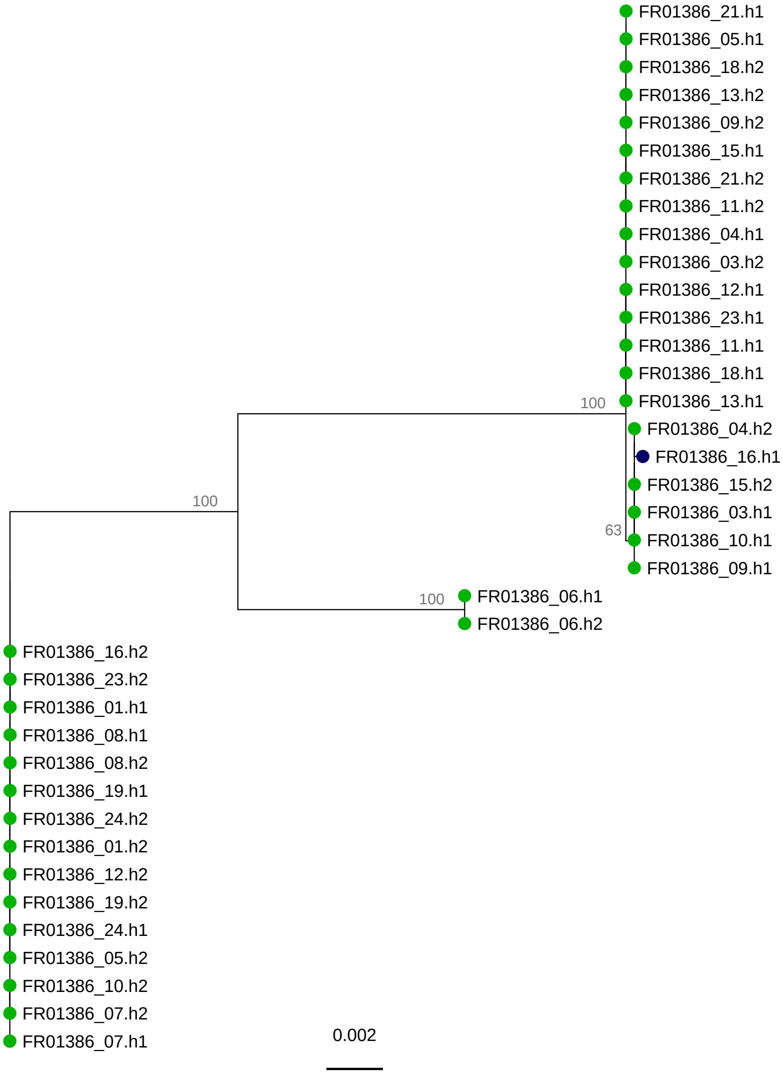
FR01386_21.h1
FR01386_05.h1
FR01386_18.h2
FR01386_13.h2
FR01386_09.h2
FR01386_15.h1
FR01386_21.h2
FR01386_11.h2
FR01386_04.h1
FR01386_03.h2
FR01386_12.h1
FR01386_23.h1
FR01386_11.h1
FR01386_18.h1
FR01386_13.h1
100
FR01386_04.h2
FR01386_16.h1
FR01386_15.h2
FR01386_03.h1
63
FR01386_10.h1
FR01386_09.h1
100
100
FR01386_06.h1
FR01386_06.h2
FR01386_16.h2
FR01386_23.h2
FR01386_01.h1
FR01386_08.h1
FR01386_08.h2
FR01386_19.h1
FR01386_24.h2
FR01386_01.h2
FR01386_12.h2
FR01386_19.h2
FR01386_24.h1
FR01386_05.h2
FR01386_10.h2
FR01386_07.h2
FR01386_07.h1
0.002

FR01434

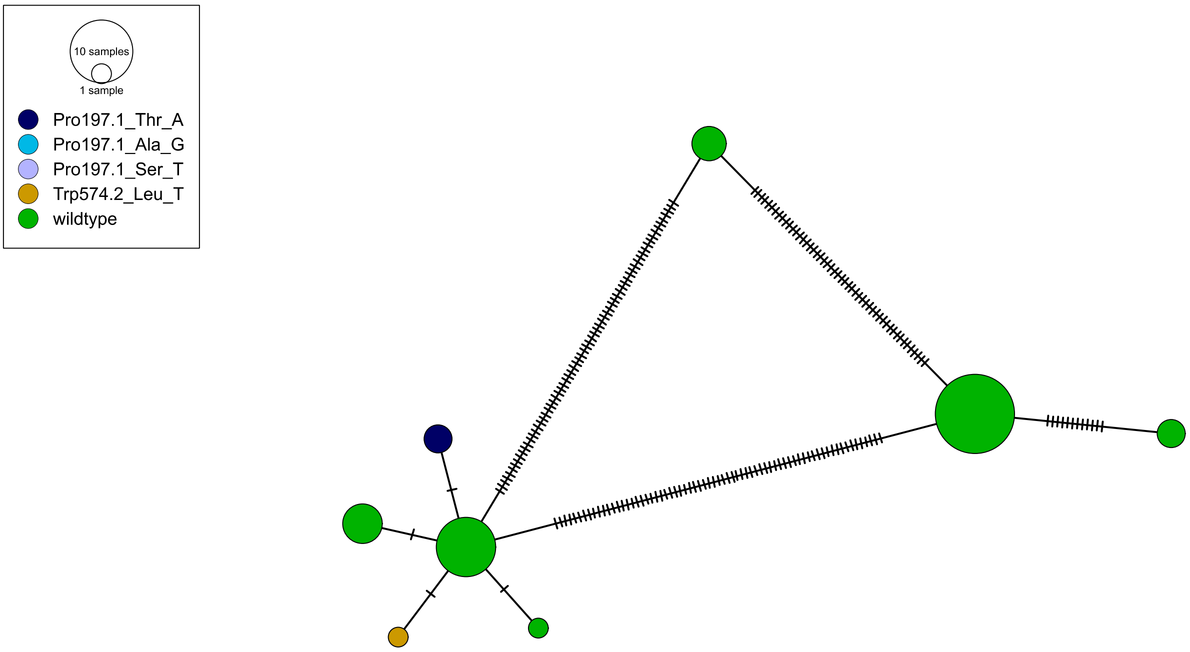

10 samples
1 sample
Pro197.1_Thr_A
Pro197.1_Ala_G
Pro197.1_Ser_T
Trp574.2_Leu_T
wildtype


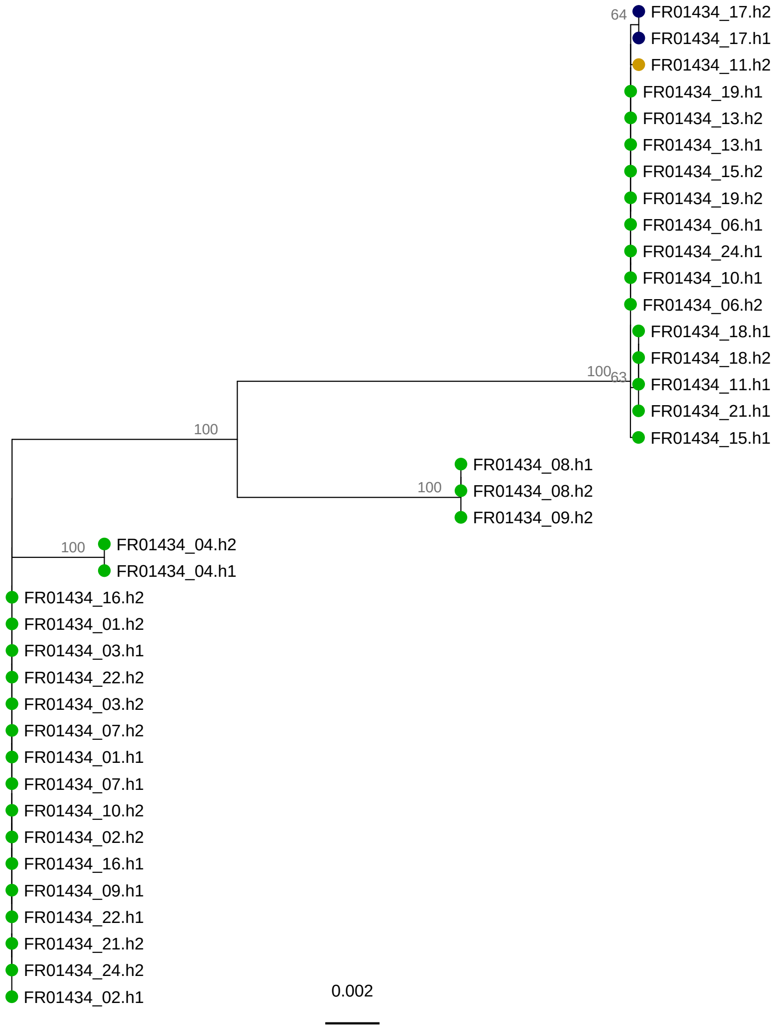

64
FR01434_17.h2
FR01434_17.h1
FR01434_11.h2
FR01434_19.h1
FR01434_13.h2
FR01434_13.h1
FR01434_15.h2
FR01434_19.h2
FR01434_06.h1
FR01434_24.h1
FR01434_10.h1
FR01434_06.h2
FR01434_18.h1
FR01434_18.h2
100
63
FR01434_11.h1
FR01434_21.h1
FR01434_15.h1
100
FR01434_08.h1
100
FR01434_08.h2
FR01434_09.h2
100
FR01434_04.h2
FR01434_04.h1
FR01434_16.h2
FR01434_01.h2
FR01434_03.h1
FR01434_22.h2
FR01434_03.h2
FR01434_07.h2
FR01434_01.h1
FR01434_07.h1
FR01434_10.h2
FR01434_02.h2
FR01434_16.h1
FR01434_09.h1
FR01434_22.h1
FR01434_21.h2
FR01434_24.h2
FR01434_02.h1
0.002

## FR01601

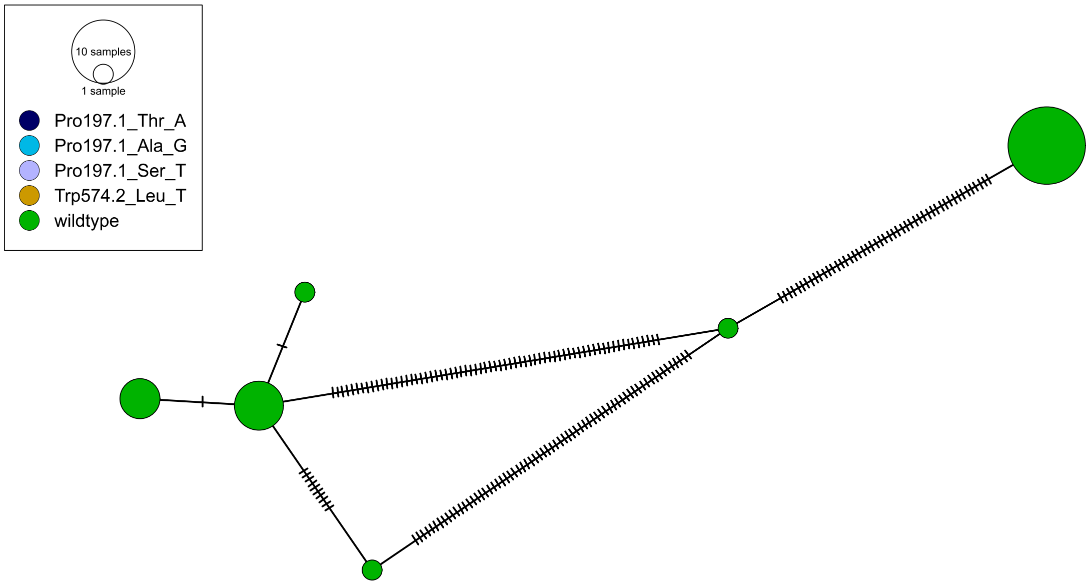
10 samples
1 sample
Pro197.1_Thr_A
Pro197.1_Ala_G
Pro197.1_Ser_T
Trp574.2_Leu_T
wildtype

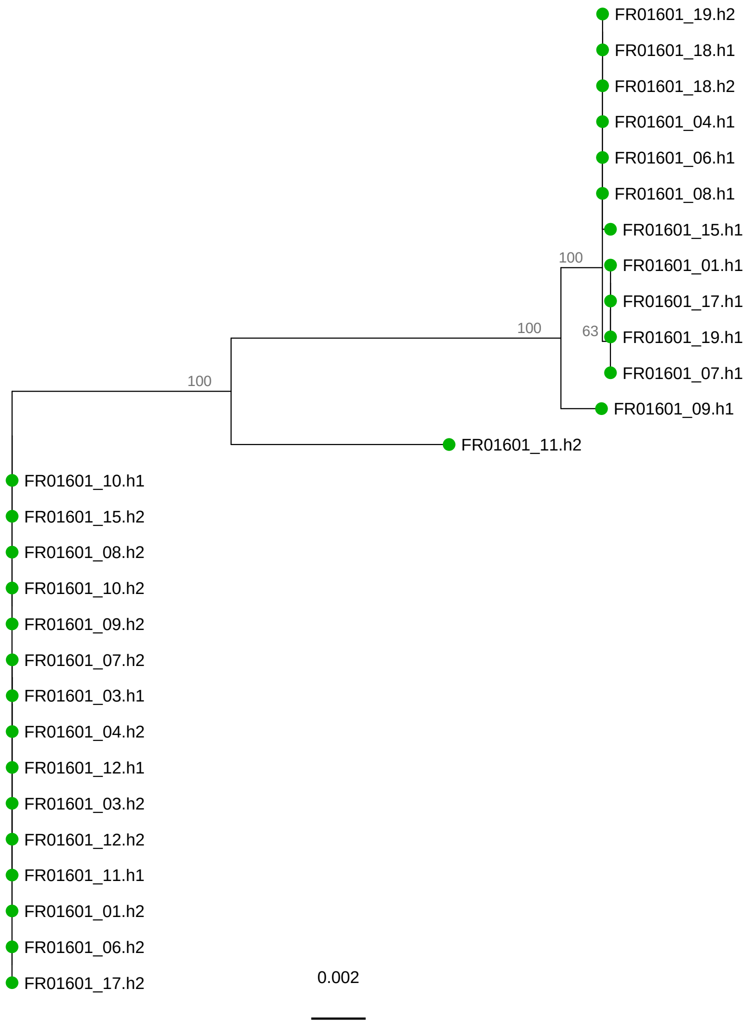
FR01601_19.h2
FR01601_18.h1
FR01601_18.h2
FR01601_04.h1
FR01601_06.h1
FR01601_08.h1
FR01601_15.h1
FR01601_01.h1
FR01601_17.h1
FR01601_19.h1
FR01601_07.h1
FR01601_09.h1
FR01601_11.h2
100
100
100
63
FR01601_10.h1
FR01601_15.h2
FR01601_08.h2
FR01601_10.h2
FR01601_09.h2
FR01601_07.h2
FR01601_03.h1
FR01601_04.h2
FR01601_12.h1
FR01601_03.h2
FR01601_12.h2
FR01601_11.h1
FR01601_01.h2
FR01601_06.h2
FR01601_17.h2
0.002

FR01729

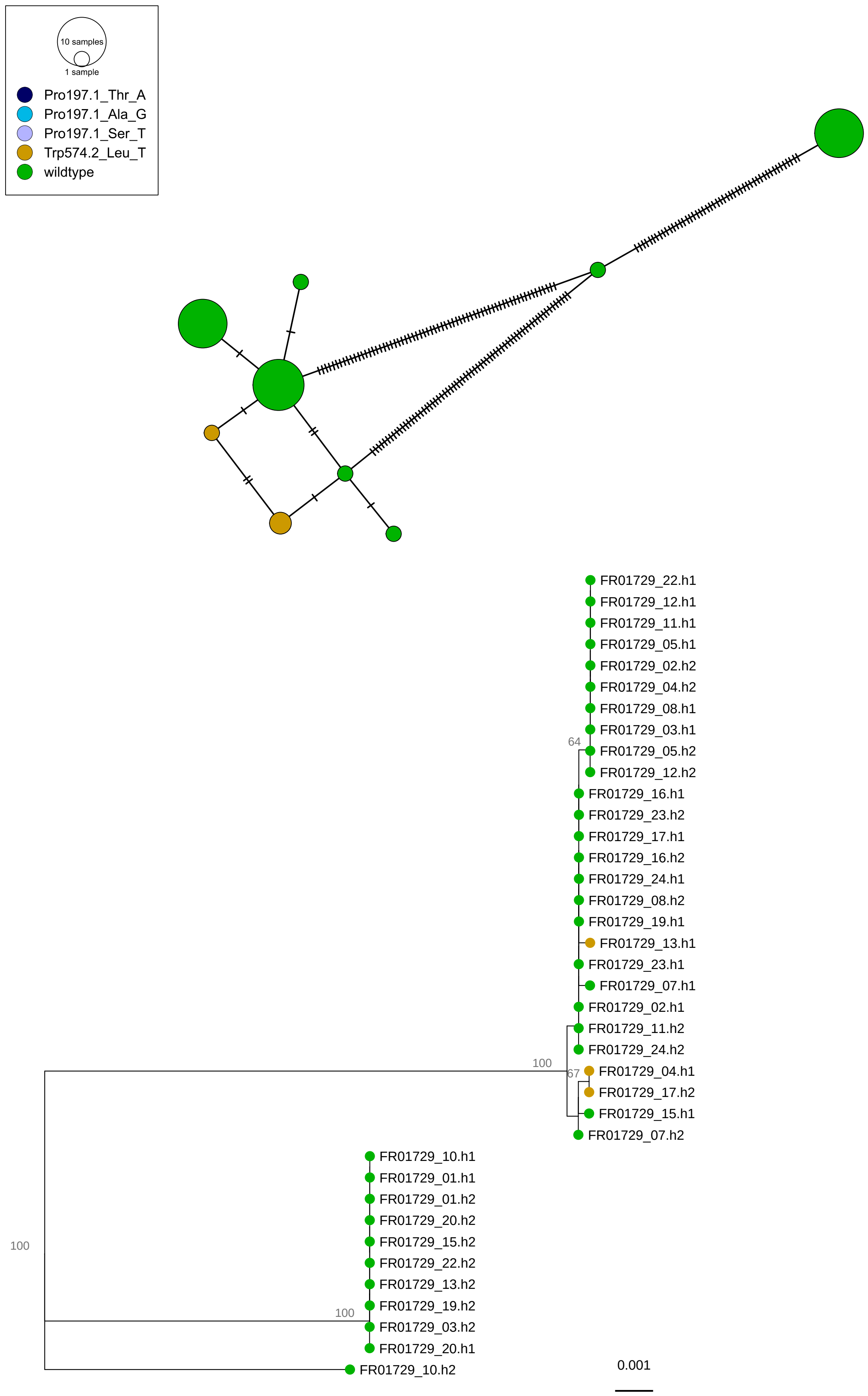
10 samples
1 sample
Pro197.1_Thr_A
Pro197.1_Ala_G
Pro197.1_Ser_T
Trp574.2_Leu_T
wildtype
FR01729_22.h1
FR01729_12.h1
FR01729_11.h1
FR01729_05.h1
FR01729_02.h2
FR01729_04.h2
FR01729_08.h1
FR01729_03.h1
64
FR01729_05.h2
FR01729_12.h2
FR01729_16.h1
FR01729_23.h2
FR01729_17.h1
FR01729_16.h2
FR01729_24.h1
FR01729_08.h2
FR01729_19.h1
FR01729_13.h1
FR01729_23.h1
FR01729_07.h1
FR01729_02.h1
FR01729_11.h2
FR01729_24.h2
100
67
FR01729_04.h1
FR01729_17.h2
FR01729_15.h1
FR01729_07.h2
FR01729_10.h1
FR01729_01.h1
FR01729_01.h2
FR01729_20.h2
FR01729_15.h2
100
FR01729_22.h2
FR01729_13.h2
FR01729_19.h2
100
FR01729_03.h2
FR01729_20.h1
FR01729_10.h2
0.001

## FR03200

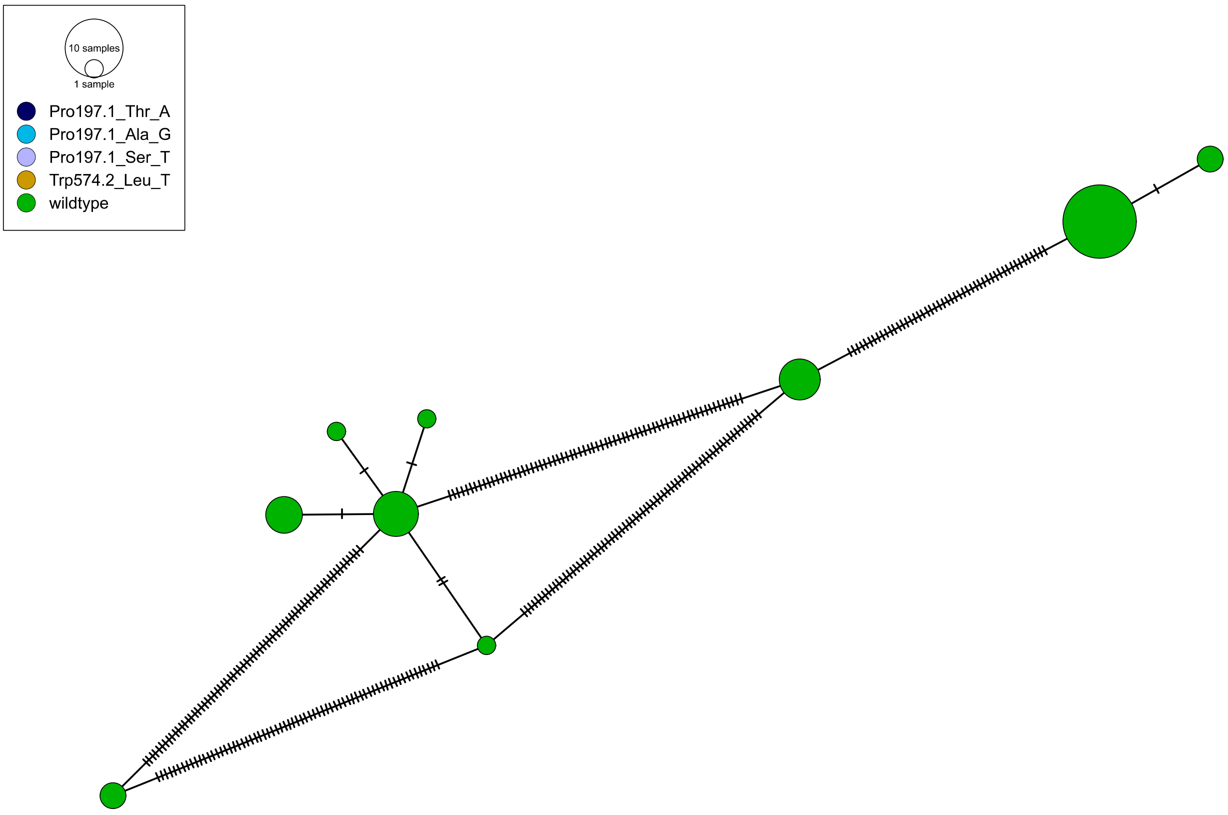
10 samples
1 sample
Pro197.1_Thr_A
Pro197.1_Ala_G
Pro197.1_Ser_T
Trp574.2_Leu_T
wildtype

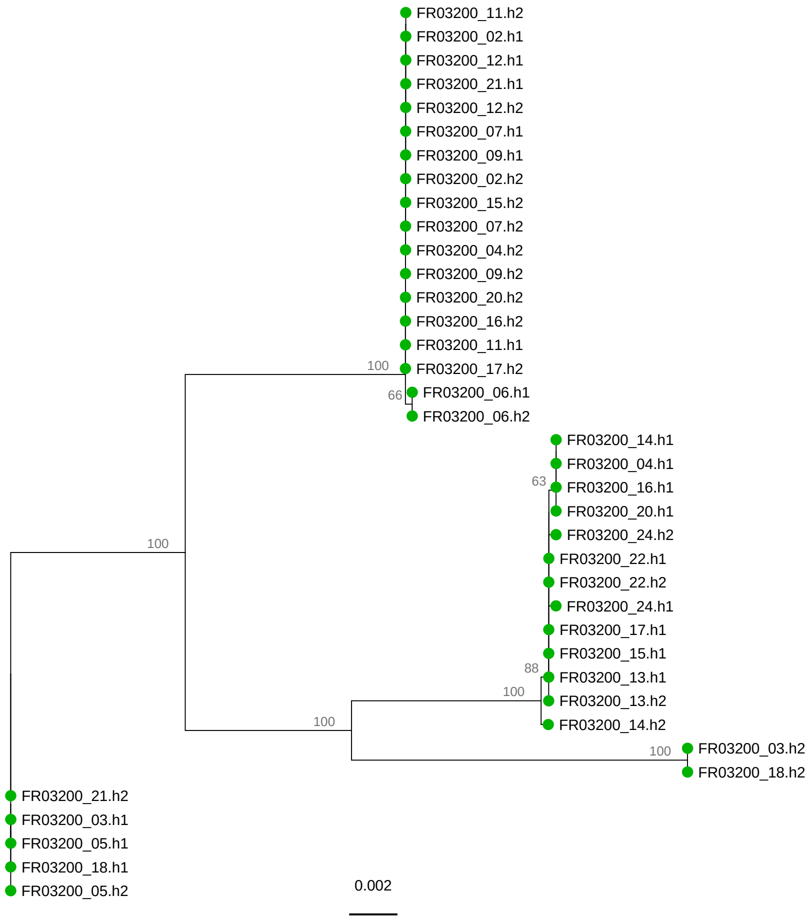
FR03200_11.h2
FR03200_02.h1
FR03200_12.h1
FR03200_21.h1
FR03200_12.h2
FR03200_07.h1
FR03200_09.h1
FR03200_02.h2
FR03200_15.h2
FR03200_07.h2
FR03200_04.h2
FR03200_09.h2
FR03200_20.h2
FR03200_16.h2
FR03200_11.h1
100
FR03200_17.h2
66
FR03200_06.h1
FR03200_06.h2
FR03200_14.h1
FR03200_04.h1
63
FR03200_16.h1
FR03200_20.h1
FR03200_24.h2
100
FR03200_22.h1
FR03200_22.h2
FR03200_24.h1
FR03200_17.h1
FR03200_15.h1
88
FR03200_13.h1
100
FR03200_13.h2
100
FR03200_14.h2
100
FR03200_03.h2
FR03200_18.h2
FR03200_21.h2
FR03200_03.h1
FR03200_05.h1
FR03200_18.h1
FR03200_05.h2
0.002

# FR07250

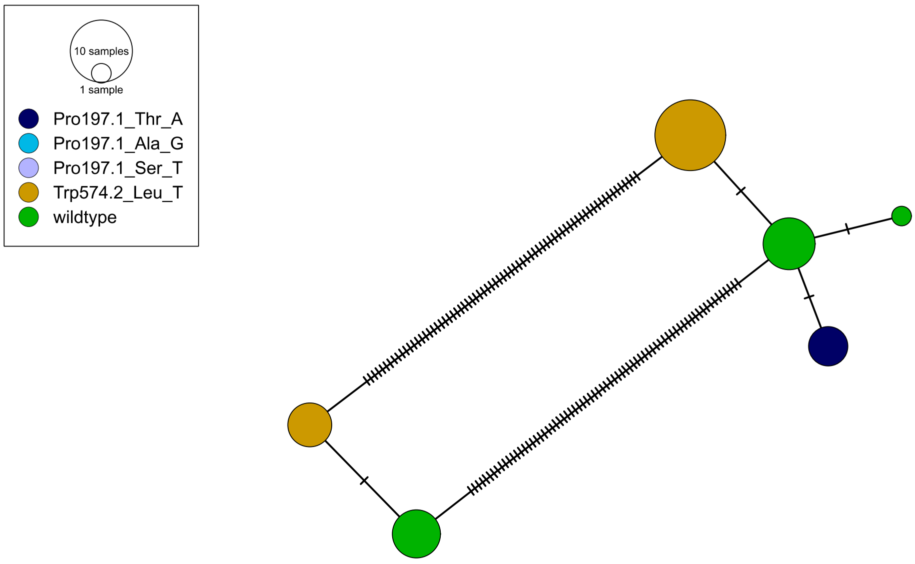
10 samples
1 sample
Pro197.1_Thr_A
Pro197.1_Ala_G
Pro197.1_Ser_T
Trp574.2_Leu_T
wildtype

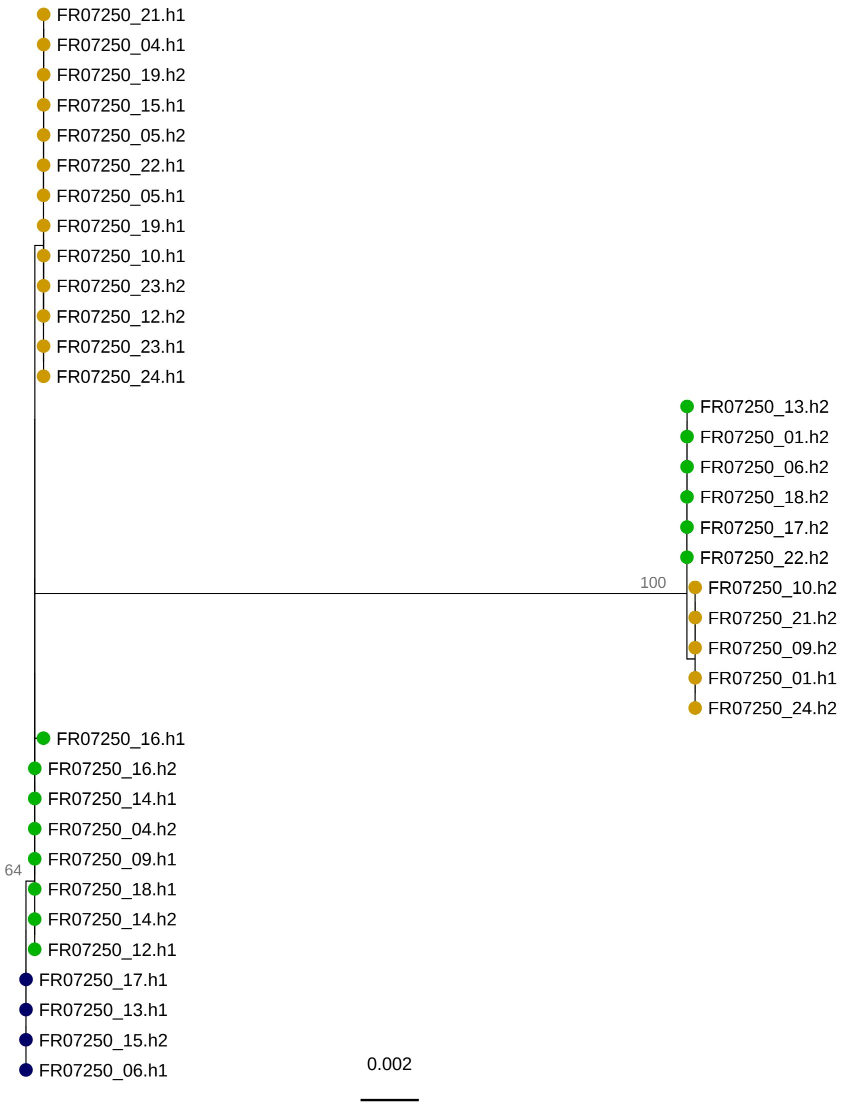
FR07250_21.h1
FR07250_04.h1
FR07250_19.h2
FR07250_15.h1
FR07250_05.h2
FR07250_22.h1
FR07250_05.h1
FR07250_19.h1
FR07250_10.h1
FR07250_23.h2
FR07250_12.h2
FR07250_23.h1
FR07250_24.h1
FR07250_13.h2
FR07250_01.h2
FR07250_06.h2
FR07250_18.h2
FR07250_17.h2
FR07250_22.h2
100
FR07250_10.h2
FR07250_21.h2
FR07250_09.h2
FR07250_01.h1
FR07250_24.h2
FR07250_16.h1
FR07250_16.h2
FR07250_14.h1
FR07250_04.h2
64
FR07250_09.h1
FR07250_18.h1
FR07250_14.h2
FR07250_12.h1
FR07250_17.h1
FR07250_13.h1
FR07250_15.h2
FR07250_06.h1
0.002

LX06513

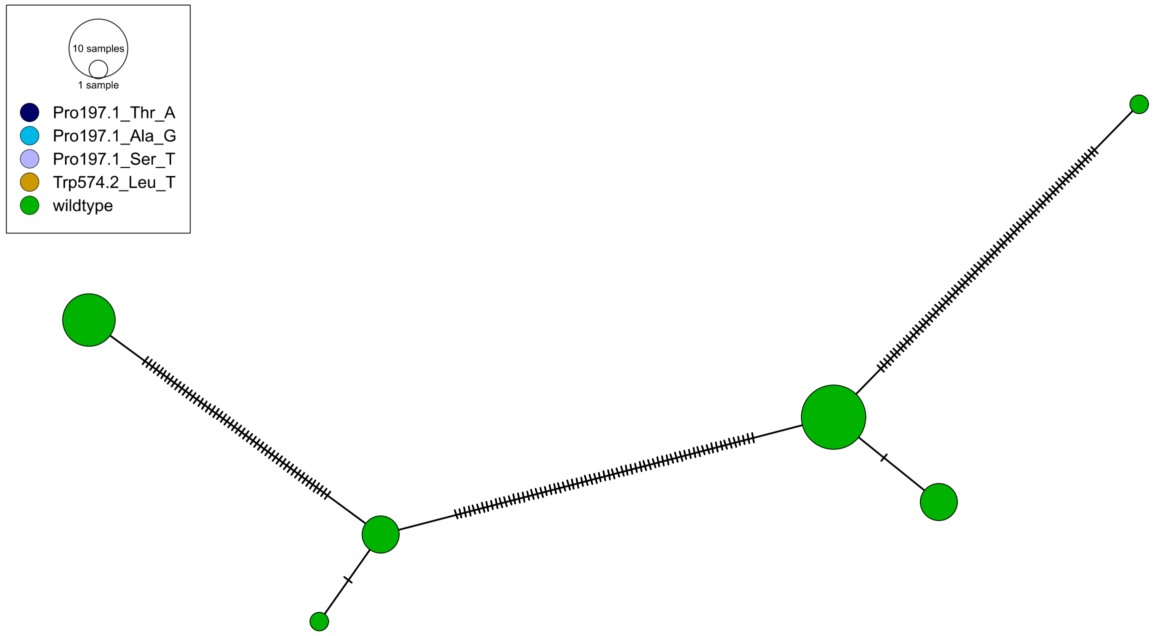
10 samples
1 sample
Pro197.1_Thr_A
Pro197.1_Ala_G
Pro197.1_Ser_T
Trp574.2_Leu_T
wildtype

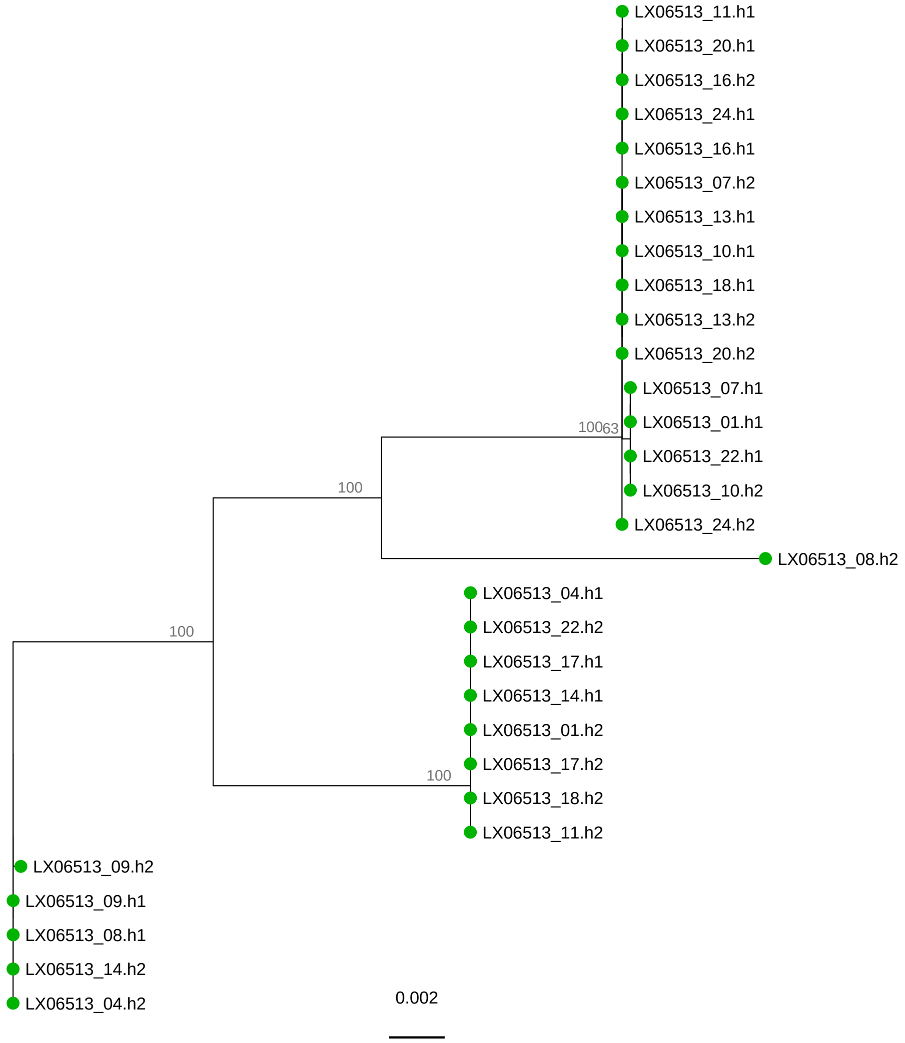
LX06513_11.h1
LX06513_20.h1
LX06513_16.h2
LX06513_24.h1
LX06513_16.h1
LX06513_07.h2
LX06513_13.h1
LX06513_10.h1
LX06513_18.h1
LX06513_13.h2
LX06513_20.h2
LX06513_07.h1
LX06513_01.h1
10063
LX06513_22.h1
100
LX06513_10.h2
LX06513_24.h2
LX06513_08.h2
LX06513_04.h1
LX06513_22.h2
100
LX06513_17.h1
LX06513_14.h1
LX06513_01.h2
100
LX06513_17.h2
LX06513_18.h2
LX06513_11.h2
LX06513_09.h2
LX06513_09.h1
LX06513_08.h1
LX06513_14.h2
LX06513_04.h2
0.002

# NL01505

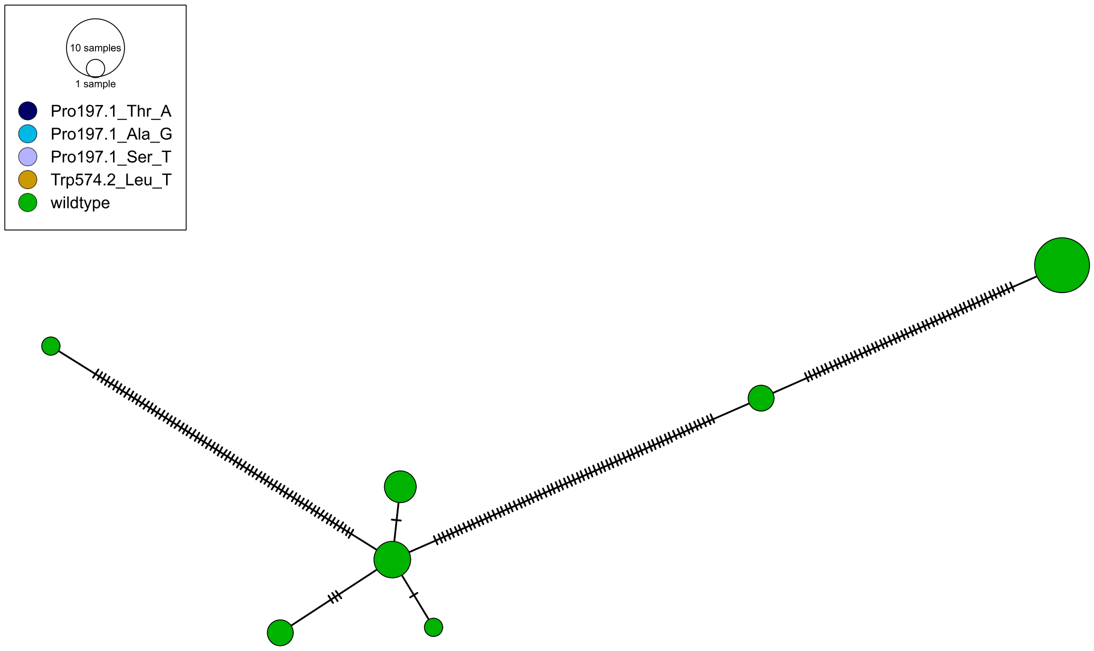
10 samples
1 sample
Pro197.1_Thr_A
Pro197.1_Ala_G
Pro197.1_Ser_T
Trp574.2_Leu_T
wildtype

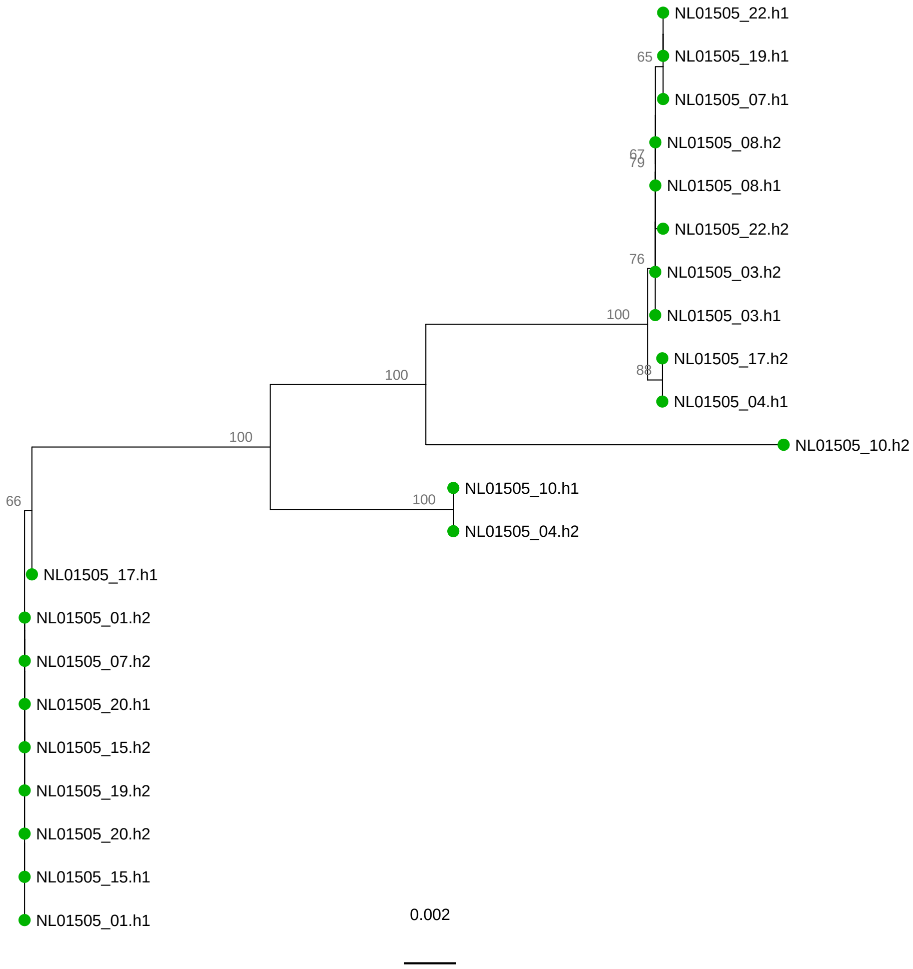
NL01505_22.h1
65
NL01505_19.h1
NL01505_07.h1
NL01505_08.h2
67
79
NL01505_08.h1
NL01505_22.h2
76
NL01505_03.h2
100
NL01505_03.h1
NL01505_17.h2
100
88
NL01505_04.h1
100
NL01505_10.h2
NL01505_10.h1
100
66
NL01505_04.h2
NL01505_17.h1
NL01505_01.h2
NL01505_07.h2
NL01505_20.h1
NL01505_15.h2
NL01505_19.h2
NL01505_20.h2
NL01505_15.h1
NL01505_01.h1
0.002

NL01664

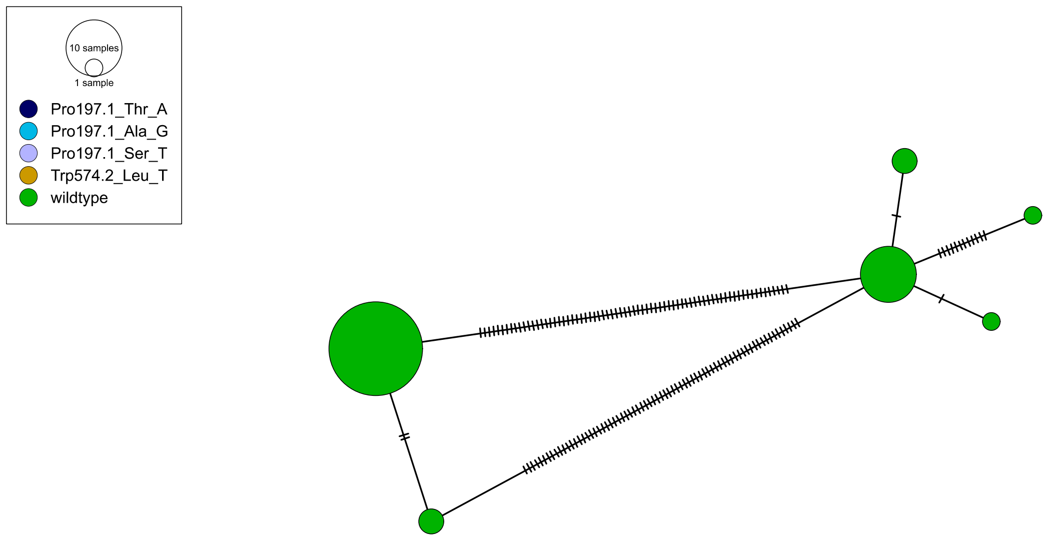
10 samples
1 sample
Pro197.1_Thr_A
Pro197.1_Ala_G
Pro197.1_Ser_T
Trp574.2_Leu_T
wildtype

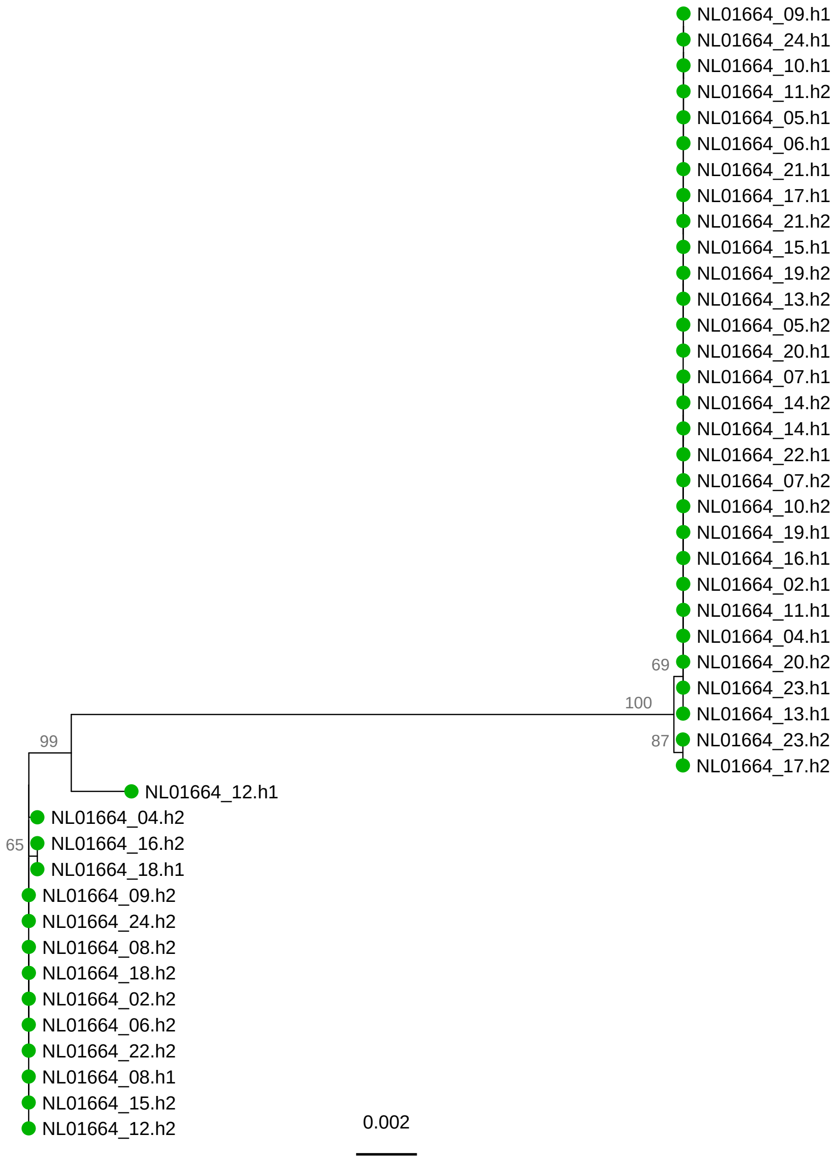
NL01664_09.h1
NL01664_24.h1
NL01664_10.h1
NL01664_11.h2
NL01664_05.h1
NL01664_06.h1
NL01664_21.h1
NL01664_17.h1
NL01664_21.h2
NL01664_15.h1
NL01664_19.h2
NL01664_13.h2
NL01664_05.h2
NL01664_20.h1
NL01664_07.h1
NL01664_14.h2
NL01664_14.h1
NL01664_22.h1
NL01664_07.h2
NL01664_10.h2
NL01664_19.h1
NL01664_16.h1
NL01664_02.h1
NL01664_11.h1
NL01664_04.h1
69
NL01664_20.h2
NL01664_23.h1
100
NL01664_13.h1
87
NL01664_23.h2
NL01664_17.h2
99
NL01664_12.h1
NL01664_04.h2
65
NL01664_16.h2
NL01664_18.h1
NL01664_09.h2
NL01664_24.h2
NL01664_08.h2
NL01664_18.h2
NL01664_02.h2
NL01664_06.h2
NL01664_22.h2
NL01664_08.h1
NL01664_15.h2
NL01664_12.h2
0.002

# NL11330

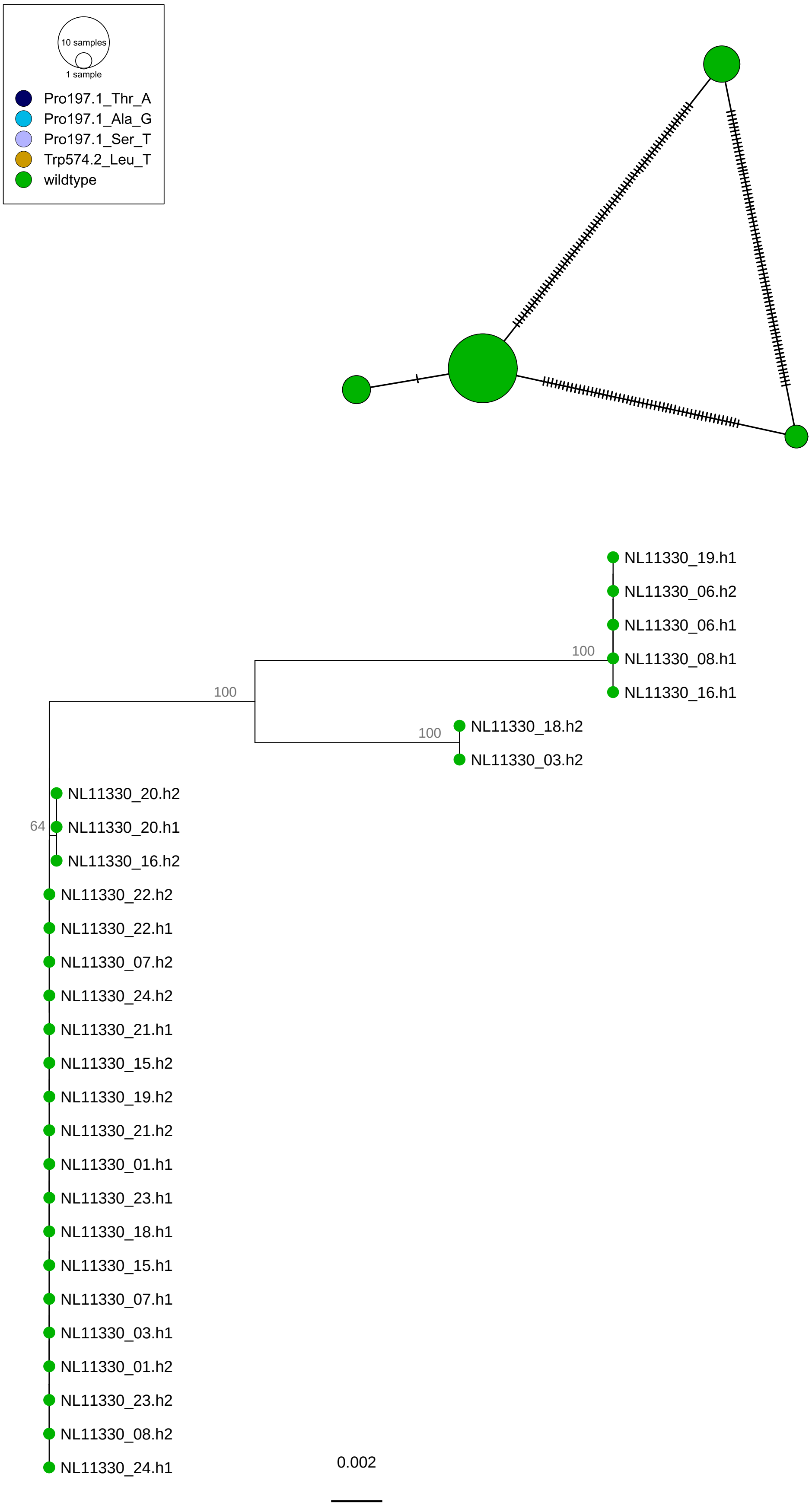
10 samples
1 sample
Pro197.1_Thr_A
Pro197.1_Ala_G
Pro197.1_Ser_T
Trp574.2_Leu_T
wildtype
NL11330_19.h1
NL11330_06.h2
NL11330_06.h1
NL11330_08.h1
NL11330_16.h1
100
100
100
NL11330_18.h2
NL11330_03.h2
64
NL11330_20.h2
NL11330_20.h1
NL11330_16.h2
NL11330_22.h2
NL11330_22.h1
NL11330_07.h2
NL11330_24.h2
NL11330_21.h1
NL11330_15.h2
NL11330_19.h2
NL11330_21.h2
NL11330_01.h1
NL11330_23.h1
NL11330_18.h1
NL11330_15.h1
NL11330_07.h1
NL11330_03.h1
NL11330_01.h2
NL11330_23.h2
NL11330_08.h2
NL11330_24.h1
0.002

PL01515

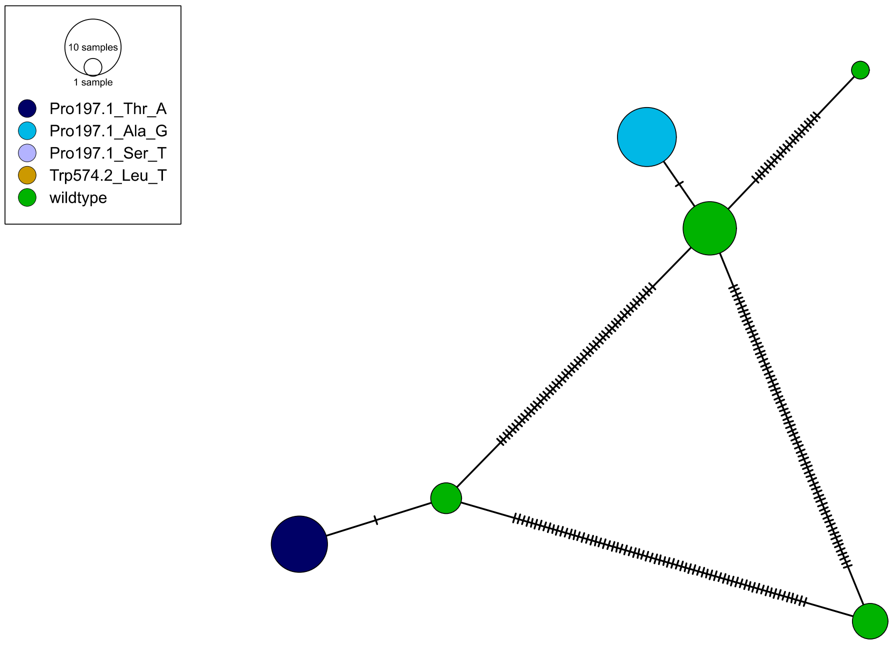
10 samples
1 sample
Pro197.1_Thr_A
Pro197.1_Ala_G
Pro197.1_Ser_T
Trp574.2_Leu_T
wildtype

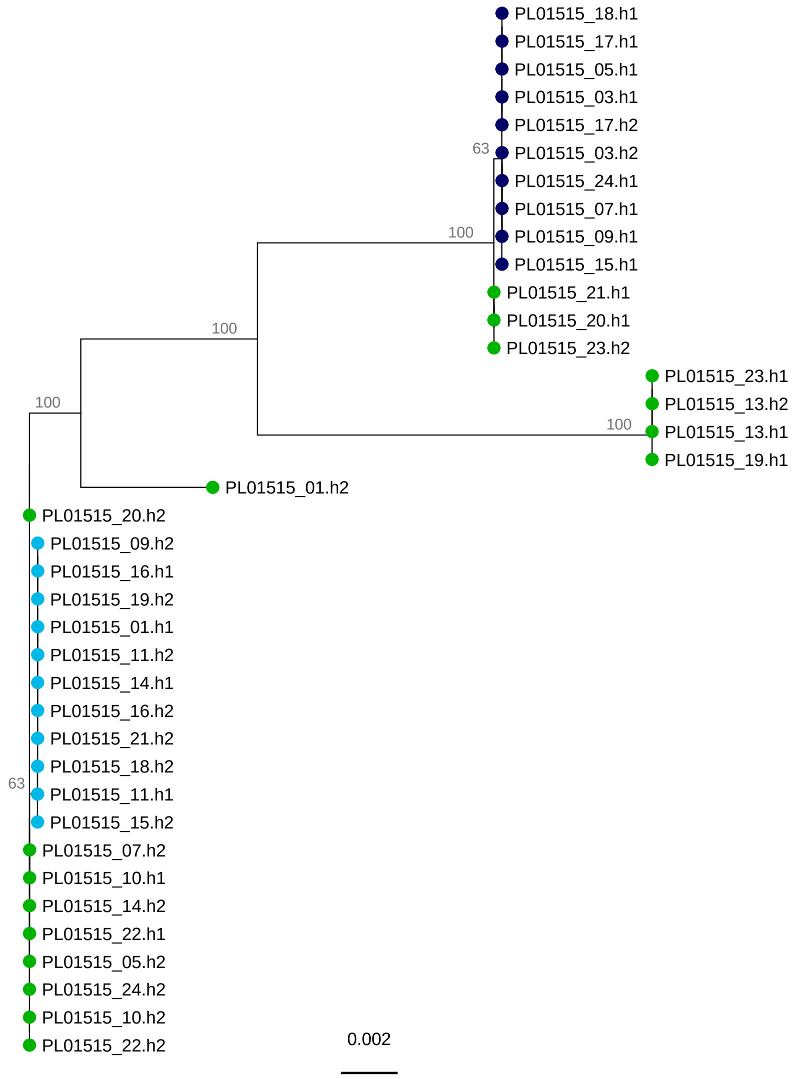
PL01515_18.h1
PL01515_17.h1
PL01515_05.h1
PL01515_03.h1
PL01515_17.h2
63
PL01515_03.h2
PL01515_24.h1
PL01515_07.h1
100
PL01515_09.h1
PL01515_15.h1
PL01515_21.h1
PL01515_20.h1
100
PL01515_23.h2
PL01515_23.h1
100
PL01515_13.h2
100
PL01515_13.h1
PL01515_19.h1
PL01515_01.h2
PL01515_20.h2
PL01515_09.h2
PL01515_16.h1
PL01515_19.h2
PL01515_01.h1
PL01515_11.h2
PL01515_14.h1
PL01515_16.h2
PL01515_21.h2
PL01515_18.h2
63
PL01515_11.h1
PL01515_15.h2
PL01515_07.h2
PL01515_10.h1
PL01515_14.h2
PL01515_22.h1
PL01515_05.h2
PL01515_24.h2
PL01515_10.h2
PL01515_22.h2
0.002

PL01742

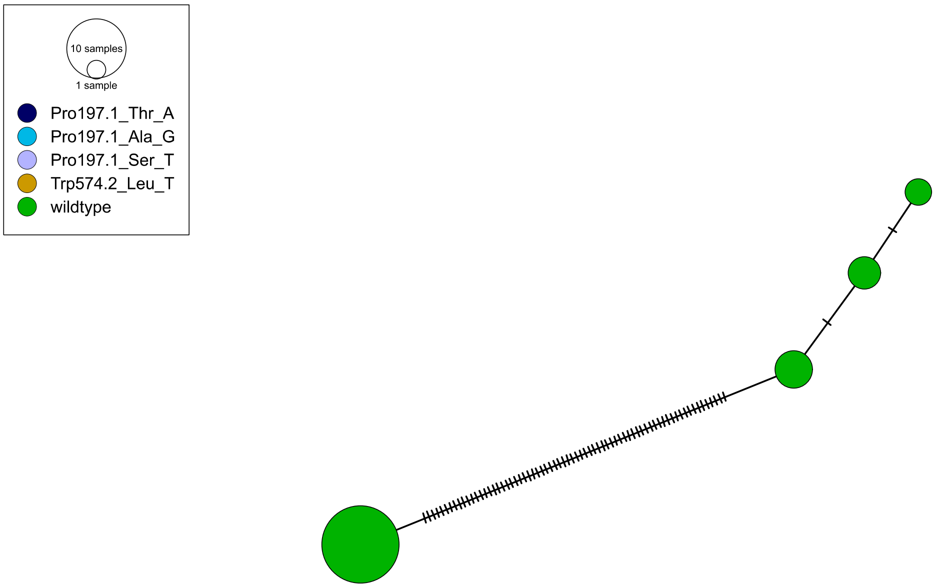
10 samples
1 sample
Pro197.1_Thr_A
Pro197.1_Ala_G
Pro197.1_Ser_T
Trp574.2_Leu_T
wildtype

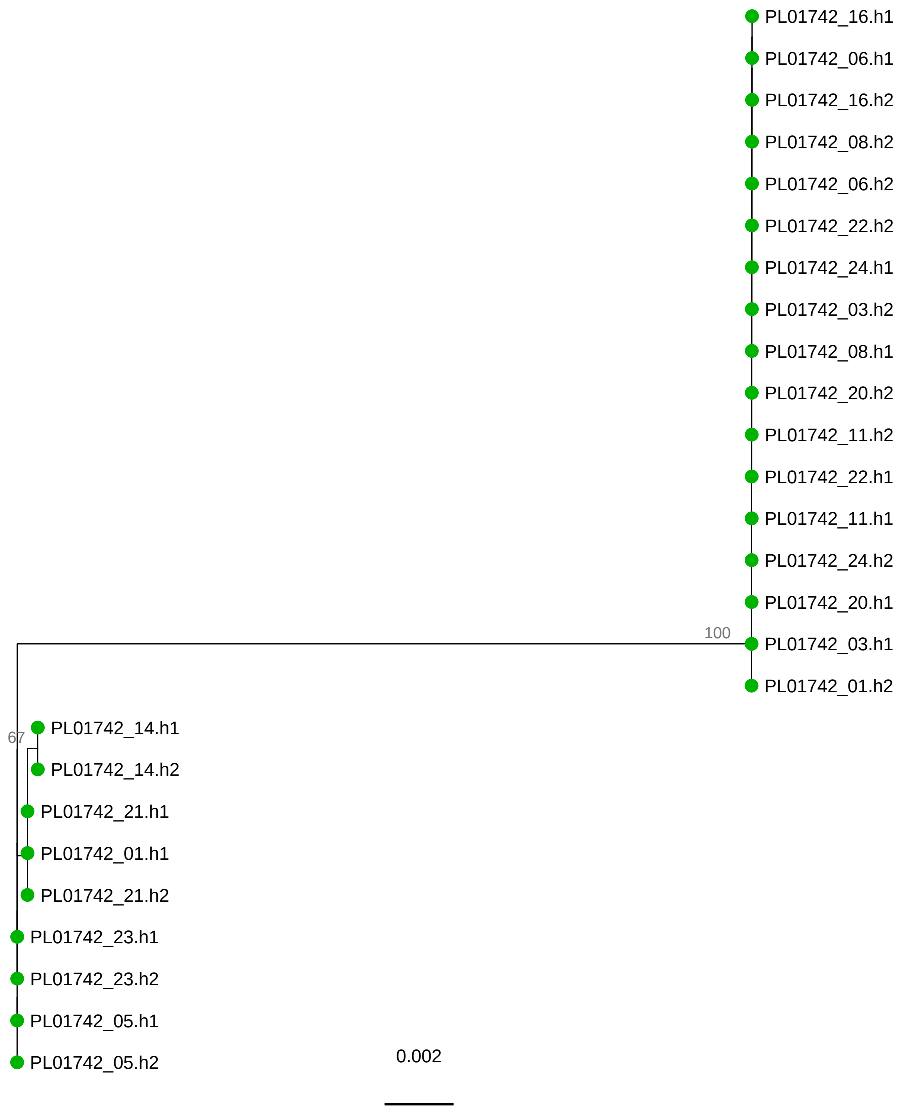
PL01742_16.h1
PL01742_06.h1
PL01742_16.h2
PL01742_08.h2
PL01742_06.h2
PL01742_22.h2
PL01742_24.h1
PL01742_03.h2
PL01742_08.h1
PL01742_20.h2
PL01742_11.h2
PL01742_22.h1
PL01742_11.h1
PL01742_24.h2
PL01742_20.h1
100
PL01742_03.h1
PL01742_01.h2
67
PL01742_14.h1
PL01742_14.h2
PL01742_21.h1
PL01742_01.h1
PL01742_21.h2
PL01742_23.h1
PL01742_23.h2
PL01742_05.h1
PL01742_05.h2
0.002

# UK01109

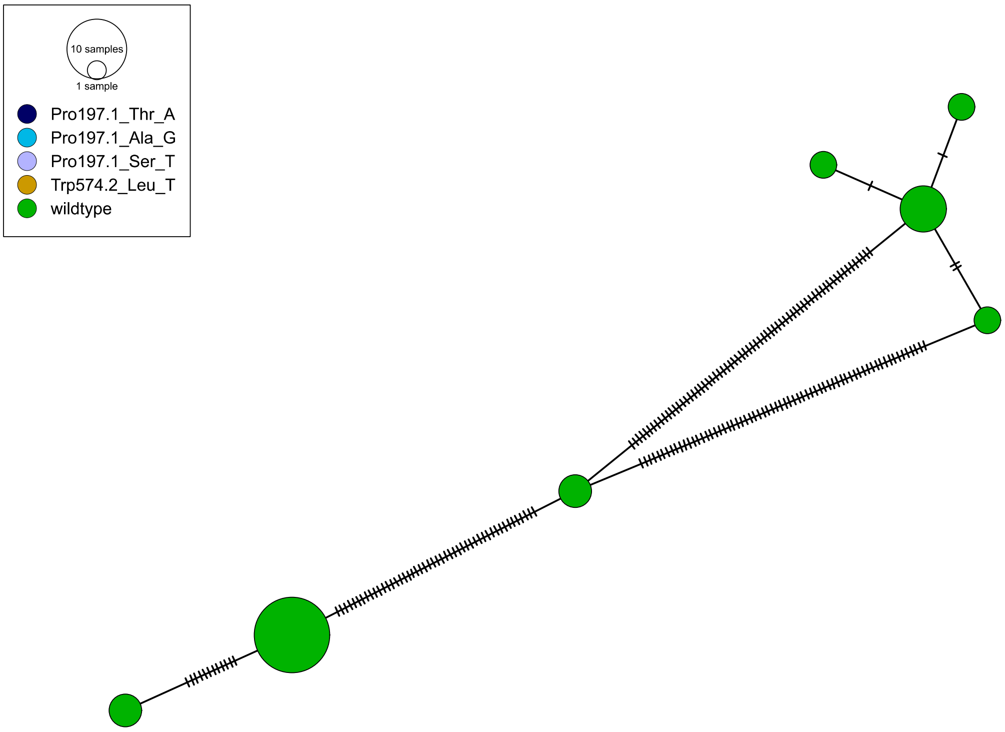
10 samples
1 sample
Pro197.1_Thr_A
Pro197.1_Ala_G
Pro197.1_Ser_T
Trp574.2_Leu_T
wildtype

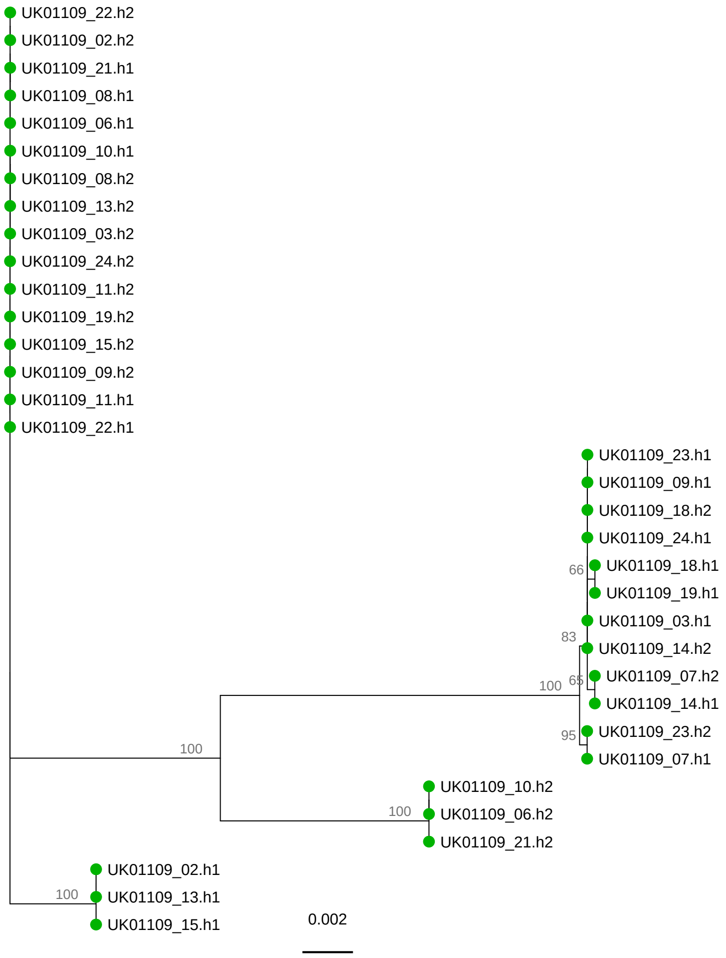
UK01109_22.h2
UK01109_02.h2
UK01109_21.h1
UK01109_08.h1
UK01109_06.h1
UK01109_10.h1
UK01109_08.h2
UK01109_13.h2
UK01109_03.h2
UK01109_24.h2
UK01109_11.h2
UK01109_19.h2
UK01109_15.h2
UK01109_09.h2
UK01109_11.h1
UK01109_22.h1
UK01109_23.h1
UK01109_09.h1
UK01109_18.h2
UK01109_24.h1
66
UK01109_18.h1
UK01109_19.h1
UK01109_03.h1
83
UK01109_14.h2
100
65
UK01109_07.h2
UK01109_14.h1
95
UK01109_23.h2
UK01109_07.h1
100
UK01109_10.h2
100
UK01109_06.h2
UK01109_21.h2
UK01109_02.h1
100
UK01109_13.h1
UK01109_15.h1
0.002

UK01413

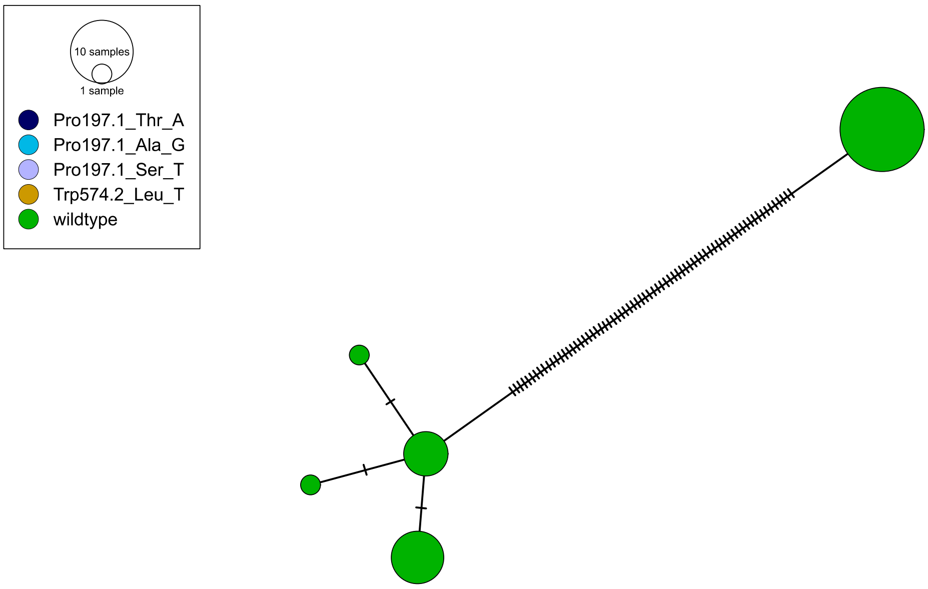
10 samples
1 sample
Pro197.1_Thr_A
Pro197.1_Ala_G
Pro197.1_Ser_T
Trp574.2_Leu_T
wildtype

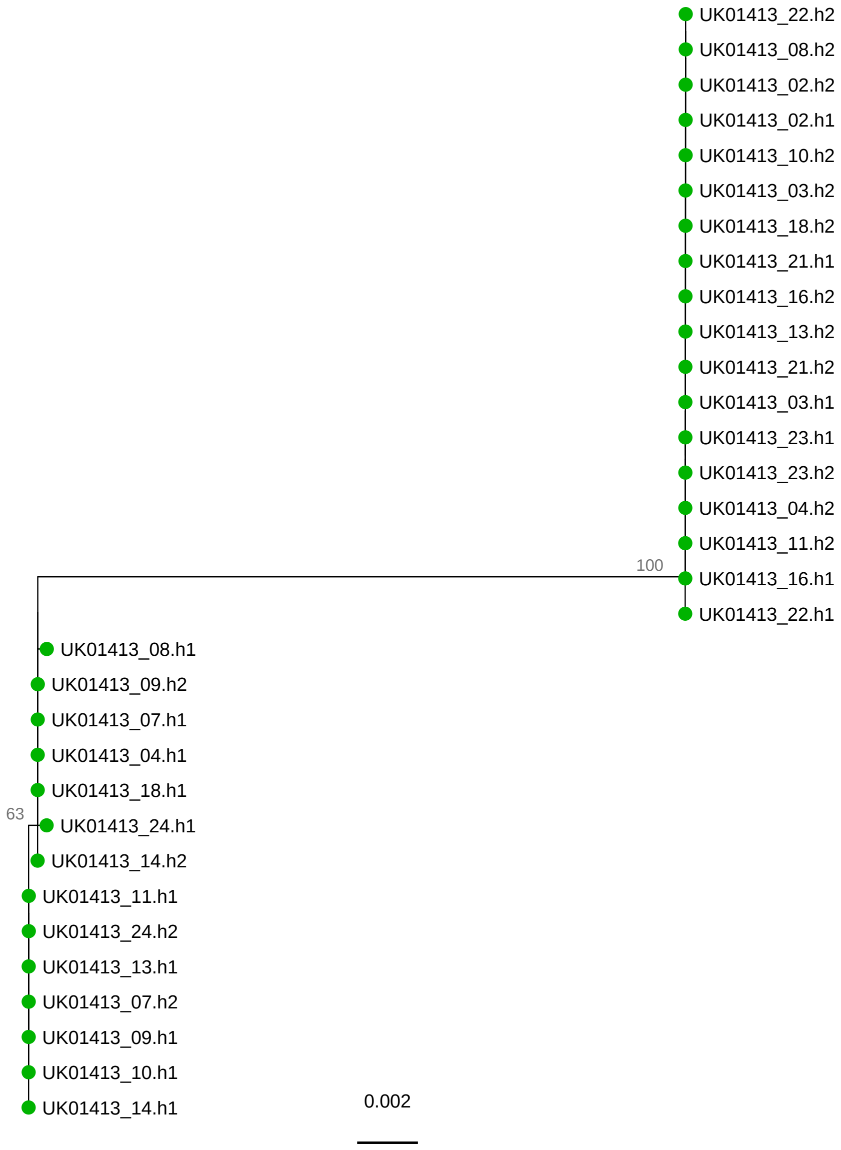
UK01413_22.h2
UK01413_08.h2
UK01413_02.h2
UK01413_02.h1
UK01413_10.h2
UK01413_03.h2
UK01413_18.h2
UK01413_21.h1
UK01413_16.h2
UK01413_13.h2
UK01413_21.h2
UK01413_03.h1
UK01413_23.h1
UK01413_23.h2
UK01413_04.h2
UK01413_11.h2
100
UK01413_16.h1
UK01413_22.h1
UK01413_08.h1
UK01413_09.h2
UK01413_07.h1
UK01413_04.h1
UK01413_18.h1
63
UK01413_24.h1
UK01413_14.h2
UK01413_11.h1
UK01413_24.h2
UK01413_13.h1
UK01413_07.h2
UK01413_09.h1
UK01413_10.h1
UK01413_14.h1
0.002

UK01447

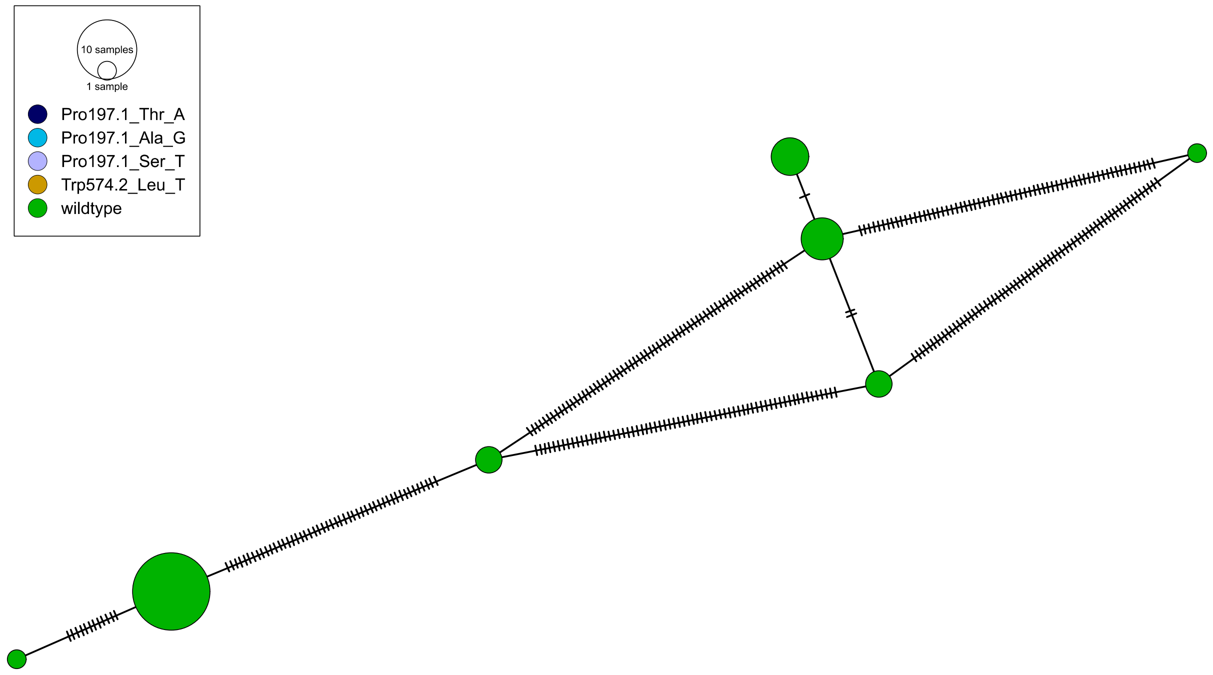
10 samples
1 sample
Pro197.1_Thr_A
Pro197.1_Ala_G
Pro197.1_Ser_T
Trp574.2_Leu_T
wildtype

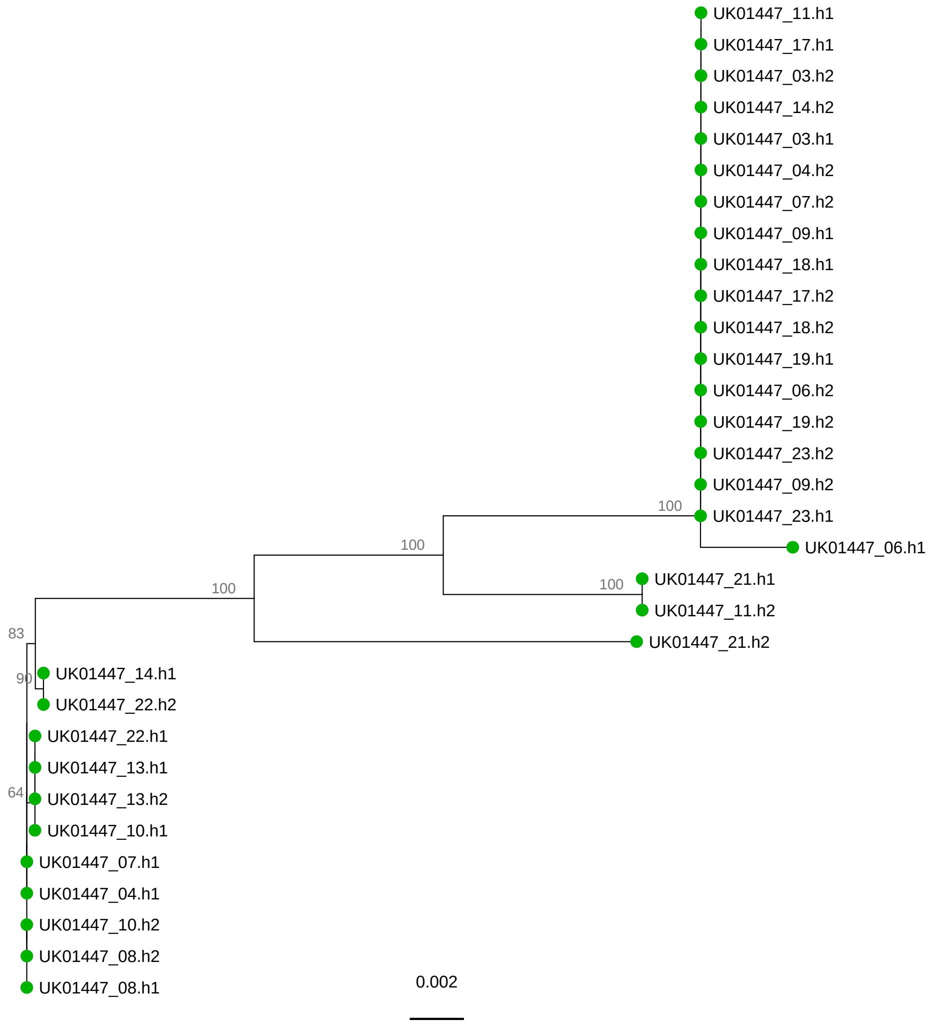
UK01447_11.h1
UK01447_17.h1
UK01447_03.h2
UK01447_14.h2
UK01447_03.h1
UK01447_04.h2
UK01447_07.h2
UK01447_09.h1
UK01447_18.h1
UK01447_17.h2
UK01447_18.h2
UK01447_19.h1
UK01447_06.h2
UK01447_19.h2
UK01447_23.h2
UK01447_09.h2
100
UK01447_23.h1
UK01447_06.h1
100
100
UK01447_21.h1
UK01447_11.h2
100
UK01447_21.h2
83
90
UK01447_14.h1
UK01447_22.h2
UK01447_22.h1
UK01447_13.h1
64
UK01447_13.h2
UK01447_10.h1
UK01447_07.h1
UK01447_04.h1
UK01447_10.h2
UK01447_08.h2
UK01447_08.h1
0.002

UK01630

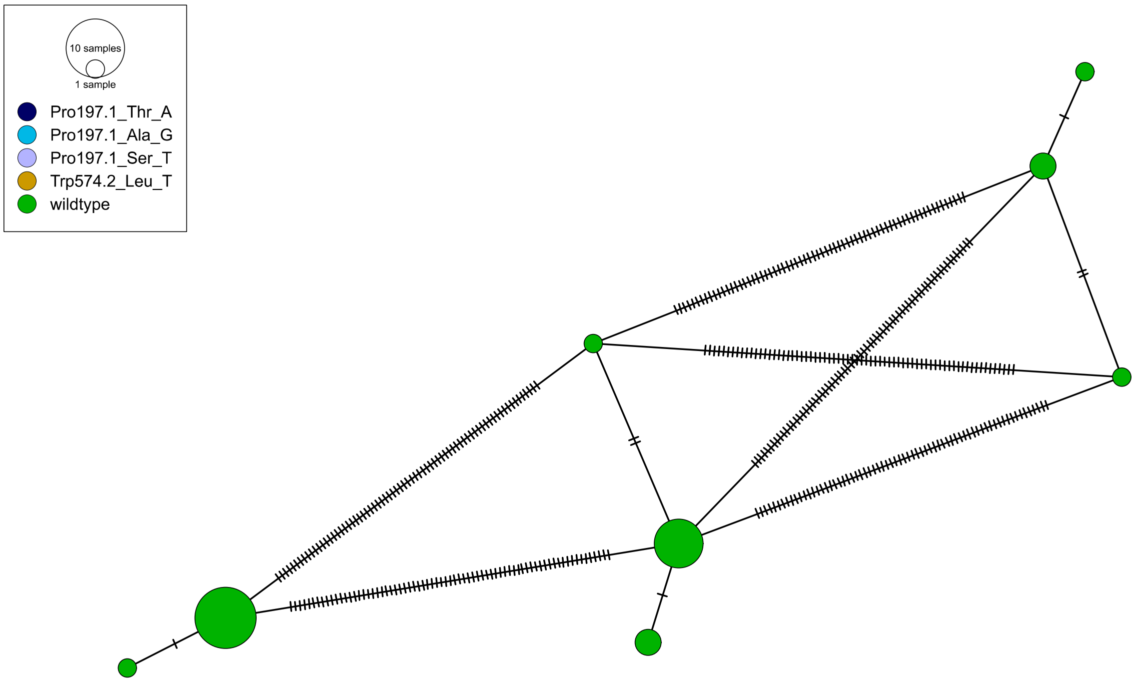
10 samples
1 sample
Pro197.1_Thr_A
Pro197.1_Ala_G
Pro197.1_Ser_T
Trp574.2_Leu_T
wildtype

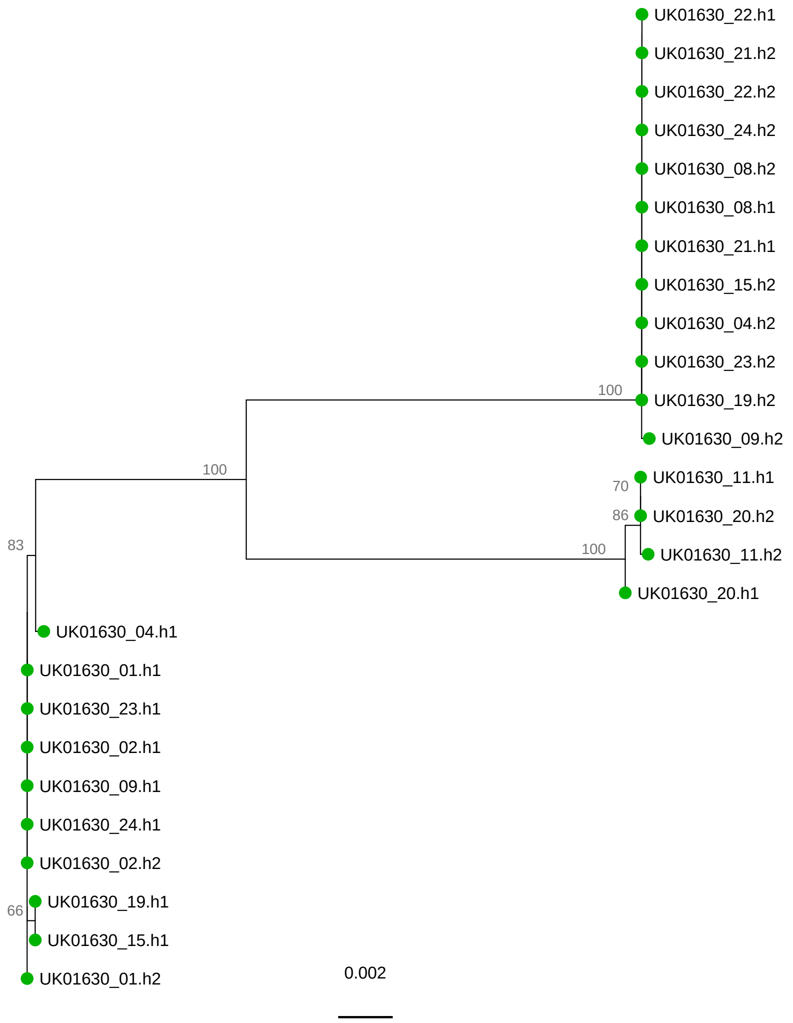
UK01630_22.h1
UK01630_21.h2
UK01630_22.h2
UK01630_24.h2
UK01630_08.h2
UK01630_08.h1
UK01630_21.h1
UK01630_15.h2
UK01630_04.h2
UK01630_23.h2
100
UK01630_19.h2
UK01630_09.h2
100
70
UK01630_11.h1
86
UK01630_20.h2
100
UK01630_11.h2
UK01630_20.h1
83
UK01630_04.h1
UK01630_01.h1
UK01630_23.h1
UK01630_02.h1
UK01630_09.h1
UK01630_24.h1
UK01630_02.h2
66
UK01630_19.h1
UK01630_15.h1
UK01630_01.h2
0.002

UK01726

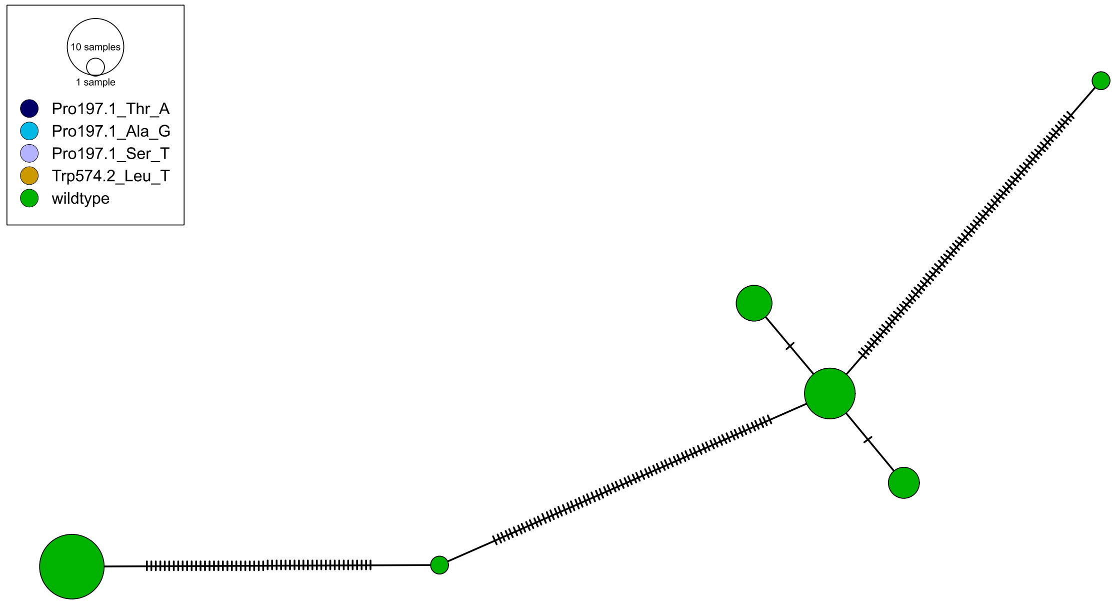
10 samples
1 sample
Pro197.1_Thr_A
Pro197.1_Ala_G
Pro197.1_Ser_T
Trp574.2_Leu_T
wildtype

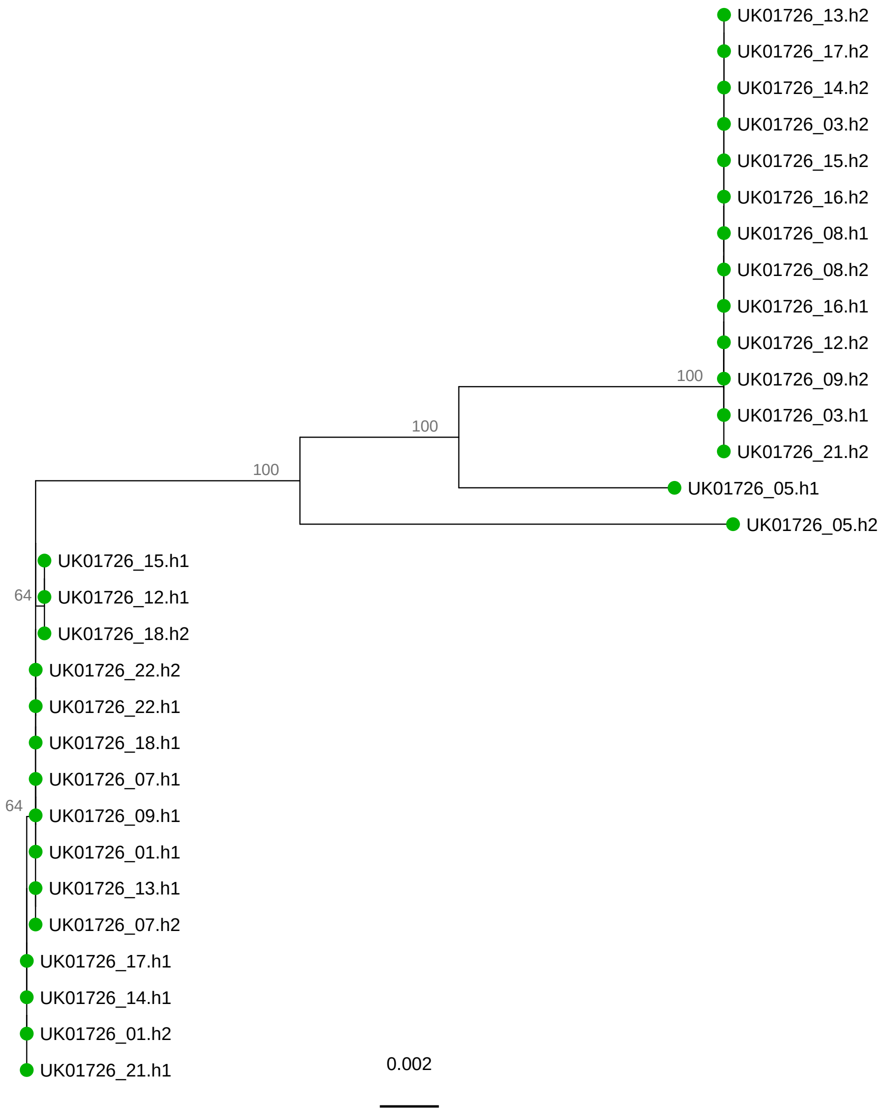
UK01726_13.h2
UK01726_17.h2
UK01726_14.h2
UK01726_03.h2
UK01726_15.h2
UK01726_16.h2
UK01726_08.h1
UK01726_08.h2
UK01726_16.h1
UK01726_12.h2
100
UK01726_09.h2
100
UK01726_03.h1
100
UK01726_21.h2
UK01726_05.h1
UK01726_05.h2
UK01726_15.h1
64
UK01726_12.h1
UK01726_18.h2
UK01726_22.h2
UK01726_22.h1
UK01726_18.h1
UK01726_07.h1
64
UK01726_09.h1
UK01726_01.h1
UK01726_13.h1
UK01726_07.h2
UK01726_17.h1
UK01726_14.h1
UK01726_01.h2
UK01726_21.h1
0.002

UK02013

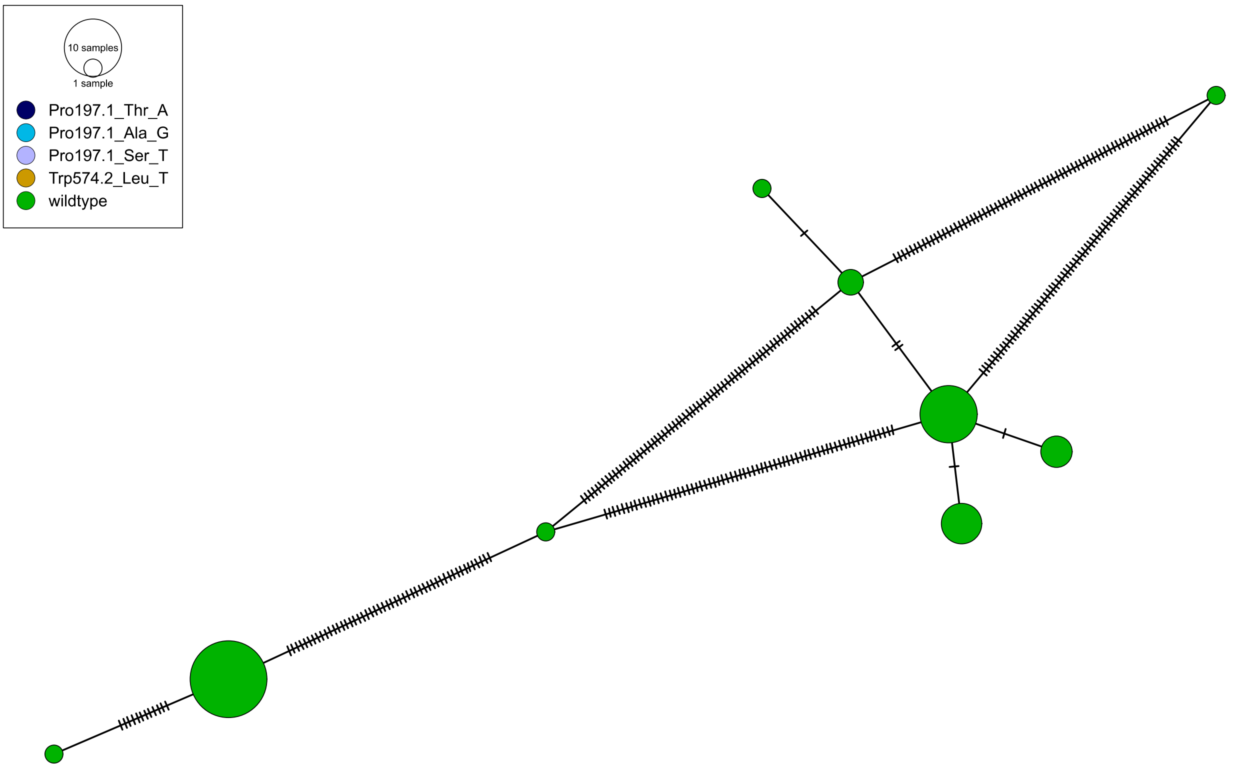
10 samples
1 sample
Pro197.1_Thr_A
Pro197.1_Ala_G
Pro197.1_Ser_T
Trp574.2_Leu_T
wildtype

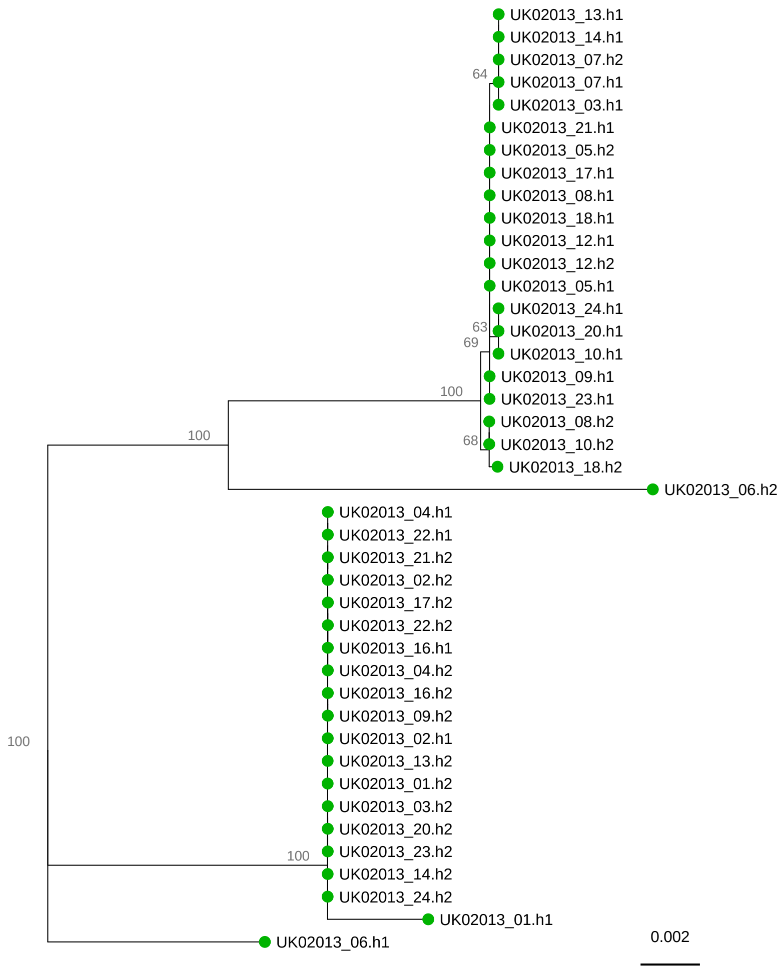
UK02013_13.h1
UK02013_14.h1
UK02013_07.h2
64
UK02013_07.h1
UK02013_03.h1
UK02013_21.h1
UK02013_05.h2
UK02013_17.h1
UK02013_08.h1
UK02013_18.h1
UK02013_12.h1
UK02013_12.h2
UK02013_05.h1
UK02013_24.h1
63
UK02013_20.h1
69
UK02013_10.h1
UK02013_09.h1
100
UK02013_23.h1
UK02013_08.h2
100
68
UK02013_10.h2
UK02013_18.h2
UK02013_06.h2
UK02013_04.h1
UK02013_22.h1
UK02013_21.h2
UK02013_02.h2
UK02013_17.h2
UK02013_22.h2
UK02013_16.h1
UK02013_04.h2
UK02013_16.h2
UK02013_09.h2
100
UK02013_02.h1
UK02013_13.h2
UK02013_01.h2
UK02013_03.h2
UK02013_20.h2
100
UK02013_23.h2
UK02013_14.h2
UK02013_24.h2
UK02013_01.h1
0.002
UK02013_06.h1

# UK03208

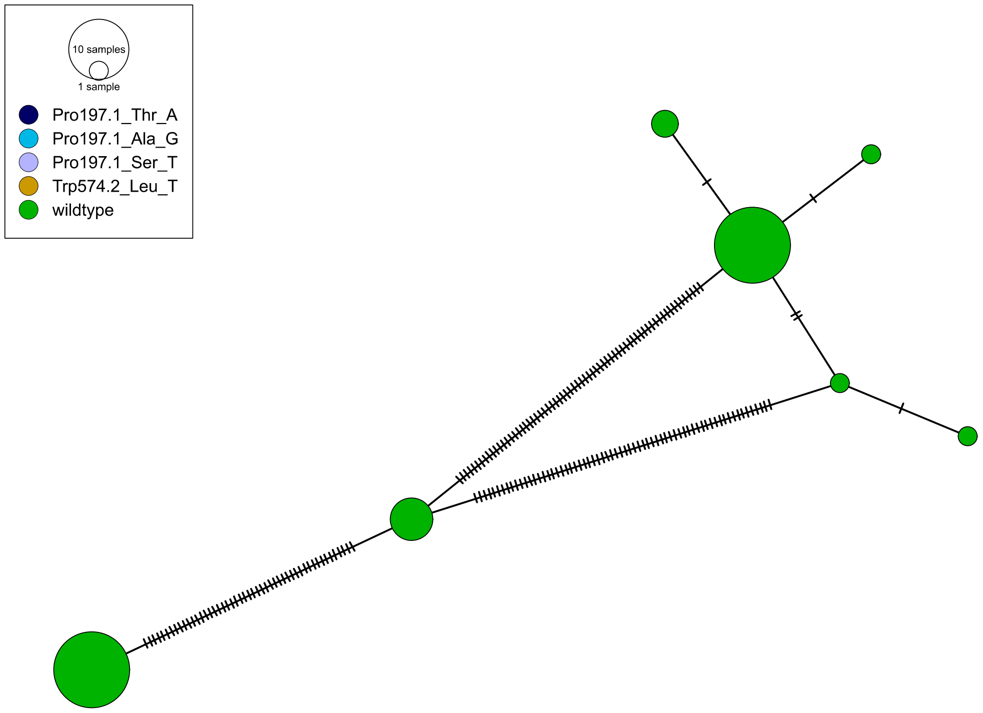
10 samples
1 sample
Pro197.1_Thr_A
Pro197.1_Ala_G
Pro197.1_Ser_T
Trp574.2_Leu_T
wildtype

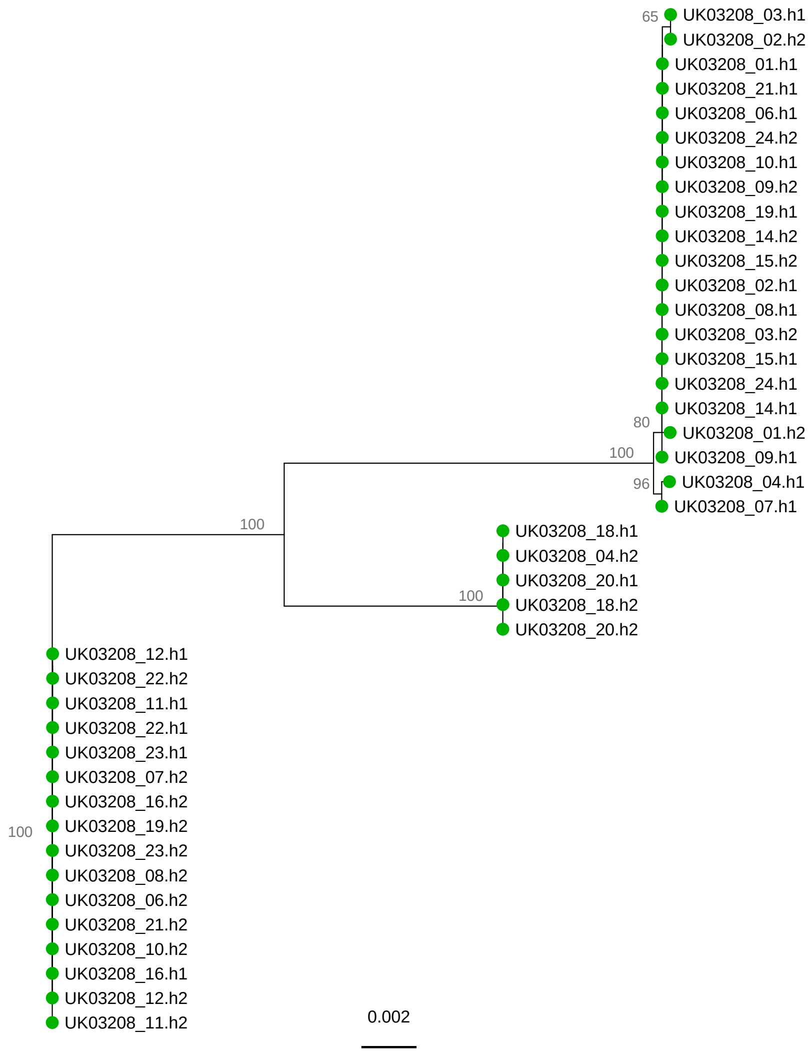
65
UK03208_03.h1
UK03208_02.h2
UK03208_01.h1
UK03208_21.h1
UK03208_06.h1
UK03208_24.h2
UK03208_10.h1
UK03208_09.h2
UK03208_19.h1
UK03208_14.h2
UK03208_15.h2
UK03208_02.h1
UK03208_08.h1
UK03208_03.h2
UK03208_15.h1
UK03208_24.h1
UK03208_14.h1
80
UK03208_01.h2
100
UK03208_09.h1
96
UK03208_04.h1
UK03208_07.h1
100
UK03208_18.h1
UK03208_04.h2
UK03208_20.h1
100
UK03208_18.h2
UK03208_20.h2
UK03208_12.h1
UK03208_22.h2
UK03208_11.h1
UK03208_22.h1
UK03208_23.h1
UK03208_07.h2
UK03208_16.h2
100
UK03208_19.h2
UK03208_23.h2
UK03208_08.h2
UK03208_06.h2
UK03208_21.h2
UK03208_10.h2
UK03208_16.h1
UK03208_12.h2
UK03208_11.h2
0.002

UK06481

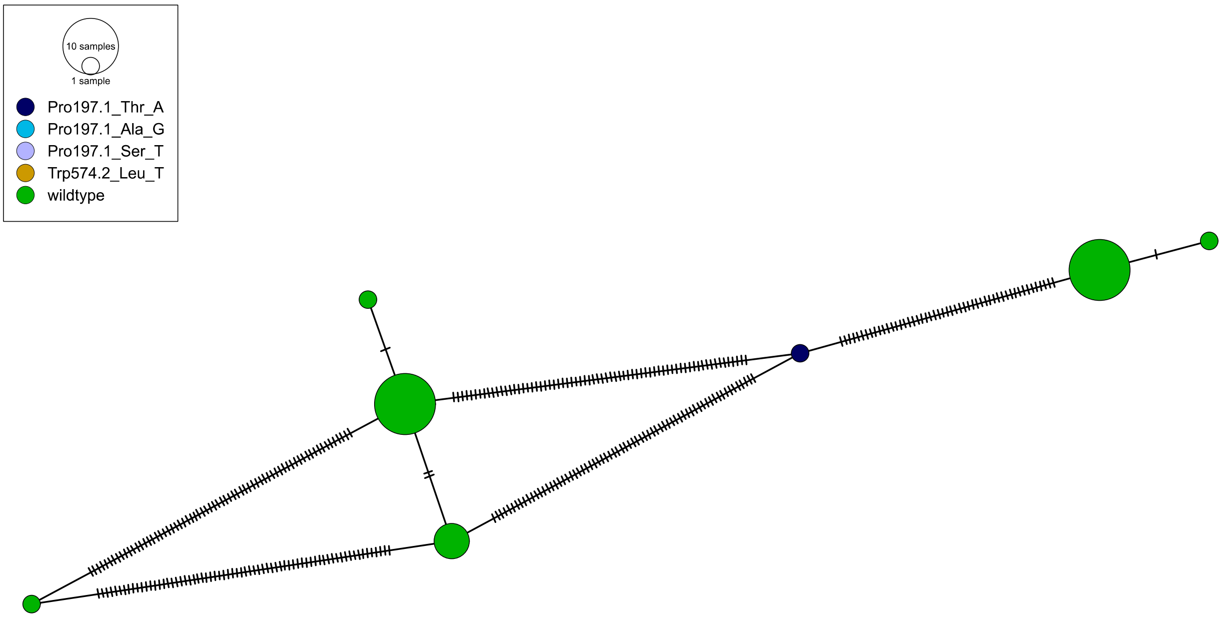
10 samples
1 sample
Pro197.1_Thr_A
Pro197.1_Ala_G
Pro197.1_Ser_T
Trp574.2_Leu_T
wildtype

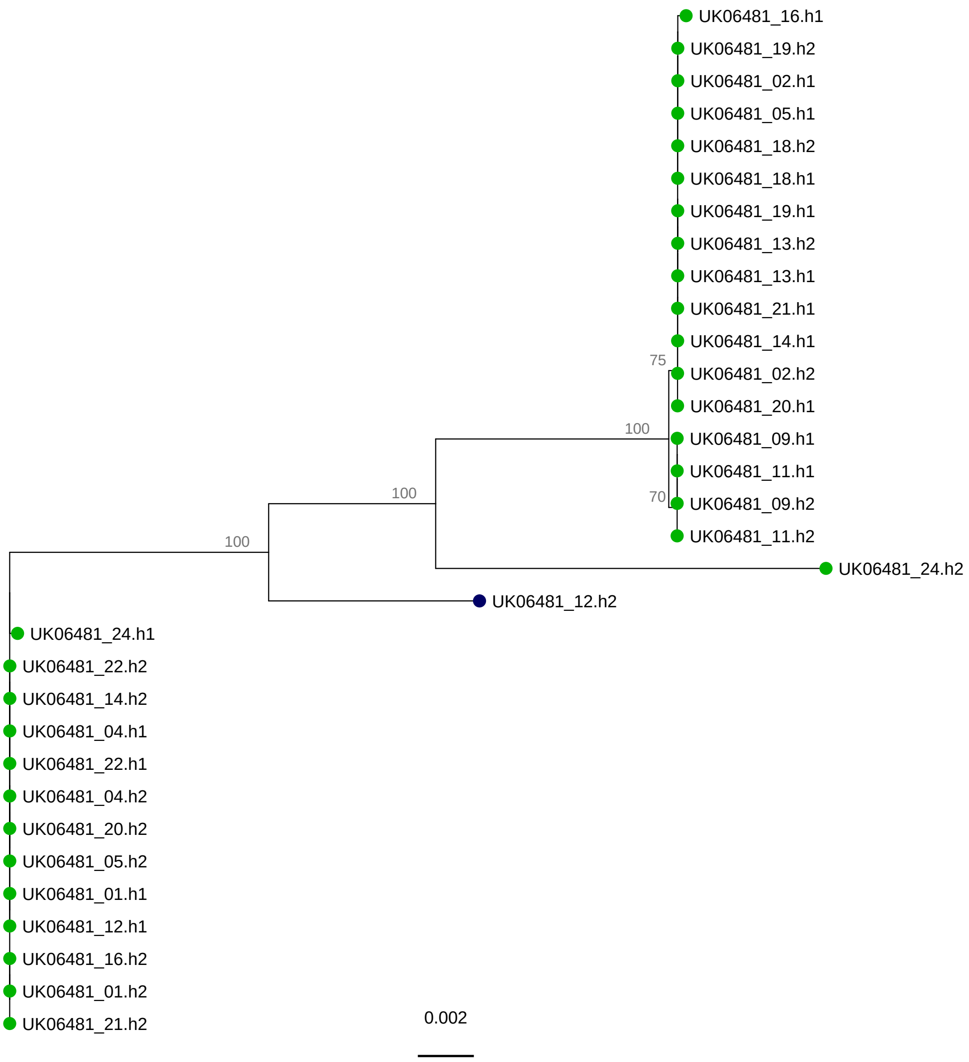
UK06481_16.h1
UK06481_19.h2
UK06481_02.h1
UK06481_05.h1
UK06481_18.h2
UK06481_18.h1
UK06481_19.h1
UK06481_13.h2
UK06481_13.h1
UK06481_21.h1
UK06481_14.h1
75
UK06481_02.h2
UK06481_20.h1
100
UK06481_09.h1
UK06481_11.h1
70
UK06481_09.h2
UK06481_11.h2
100
100
UK06481_24.h2
UK06481_12.h2
UK06481_24.h1
UK06481_22.h2
UK06481_14.h2
UK06481_04.h1
UK06481_22.h1
UK06481_04.h2
UK06481_20.h2
UK06481_05.h2
UK06481_01.h1
UK06481_12.h1
UK06481_16.h2
UK06481_01.h2
UK06481_21.h2
0.002

# UK06500

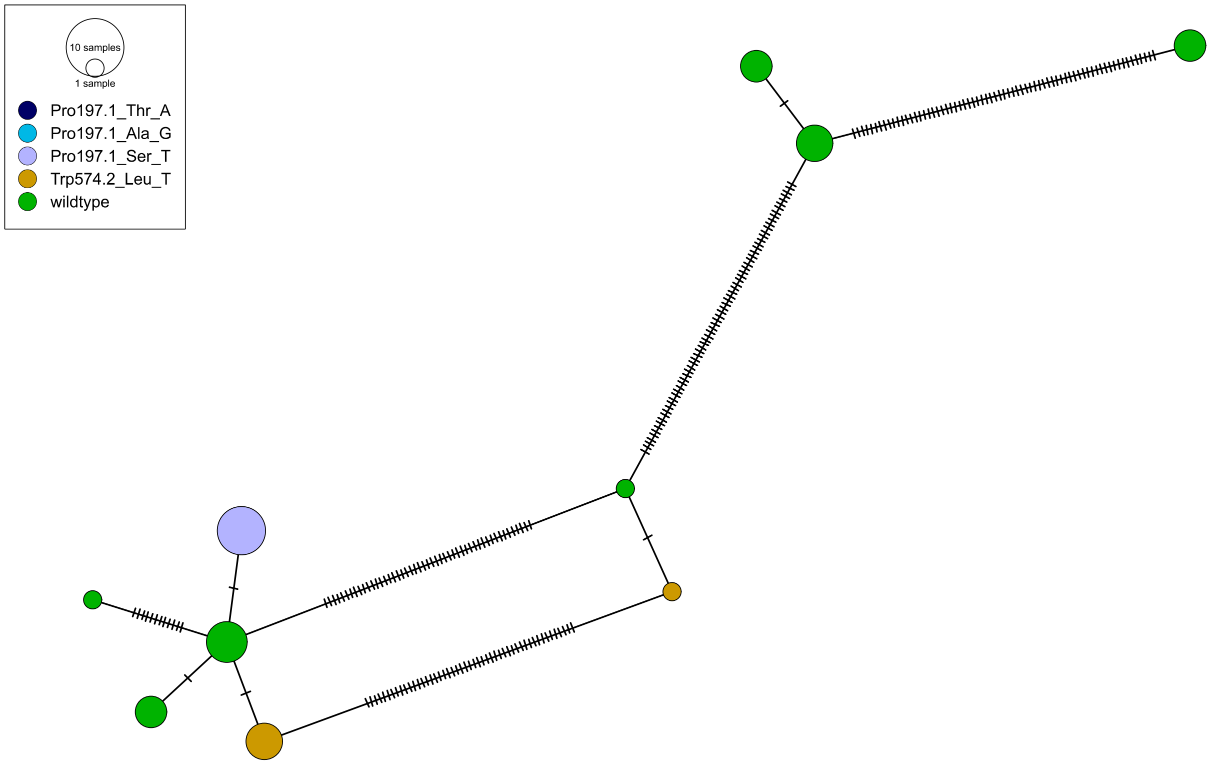

10 samples
1 sample
Pro197.1_Thr_A
Pro197.1_Ala_G
Pro197.1_Ser_T
Trp574.2_Leu_T
wildtype


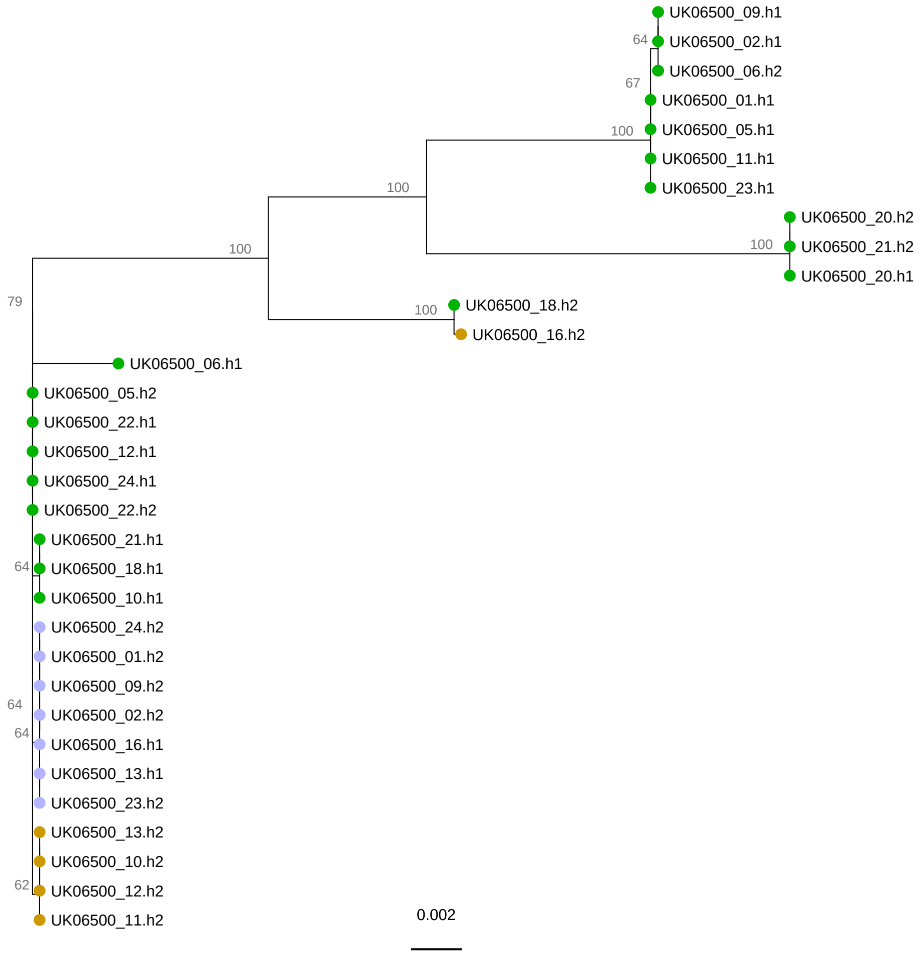

UK06500_09.h1
64
UK06500_02.h1
UK06500_06.h2
67
UK06500_01.h1
100
UK06500_05.h1
UK06500_11.h1
UK06500_23.h1
100
UK06500_20.h2
100
UK06500_21.h2
100
UK06500_20.h1
79
100
UK06500_18.h2
UK06500_16.h2
UK06500_06.h1
UK06500_05.h2
UK06500_22.h1
UK06500_12.h1
UK06500_24.h1
UK06500_22.h2
UK06500_21.h1
64
UK06500_18.h1
UK06500_10.h1
UK06500_24.h2
UK06500_01.h2
UK06500_09.h2
64
UK06500_02.h2
64
UK06500_16.h1
UK06500_13.h1
UK06500_23.h2
UK06500_13.h2
UK06500_10.h2
62
UK06500_12.h2
UK06500_11.h2
0.002

UK06518

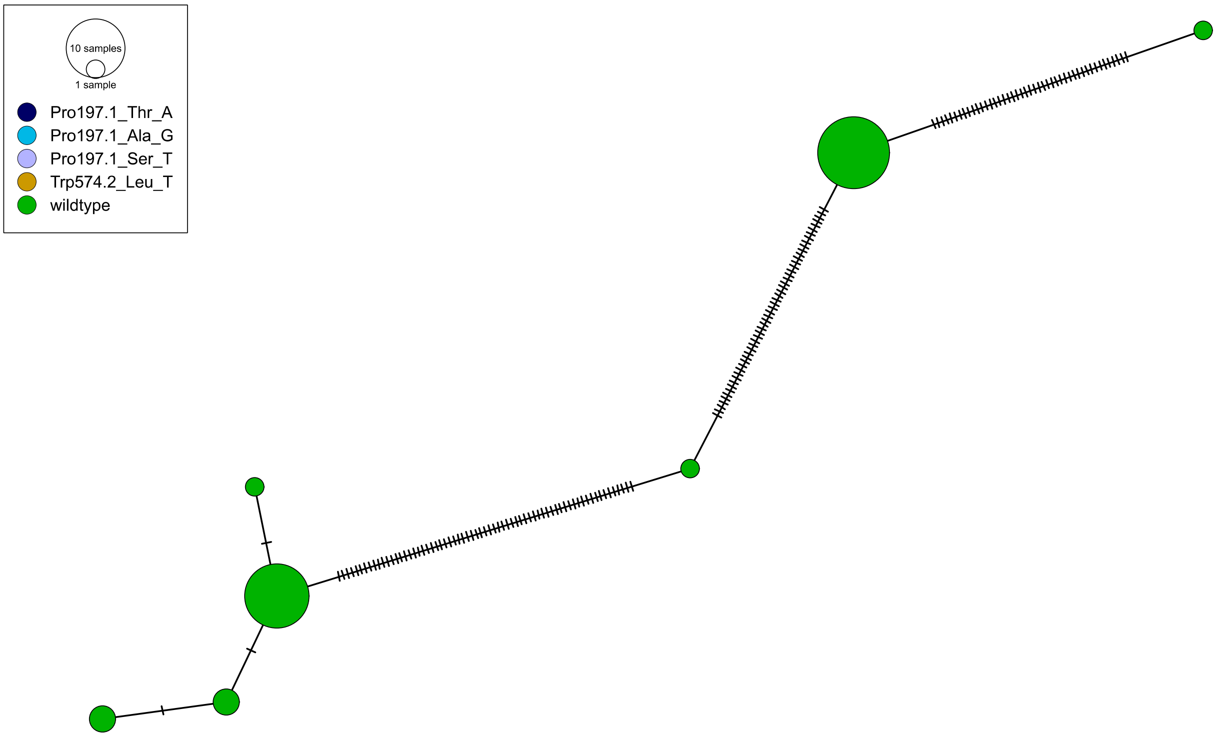
10 samples
1 sample
Pro197.1_Thr_A
Pro197.1_Ala_G
Pro197.1_Ser_T
Trp574.2_Leu_T
wildtype

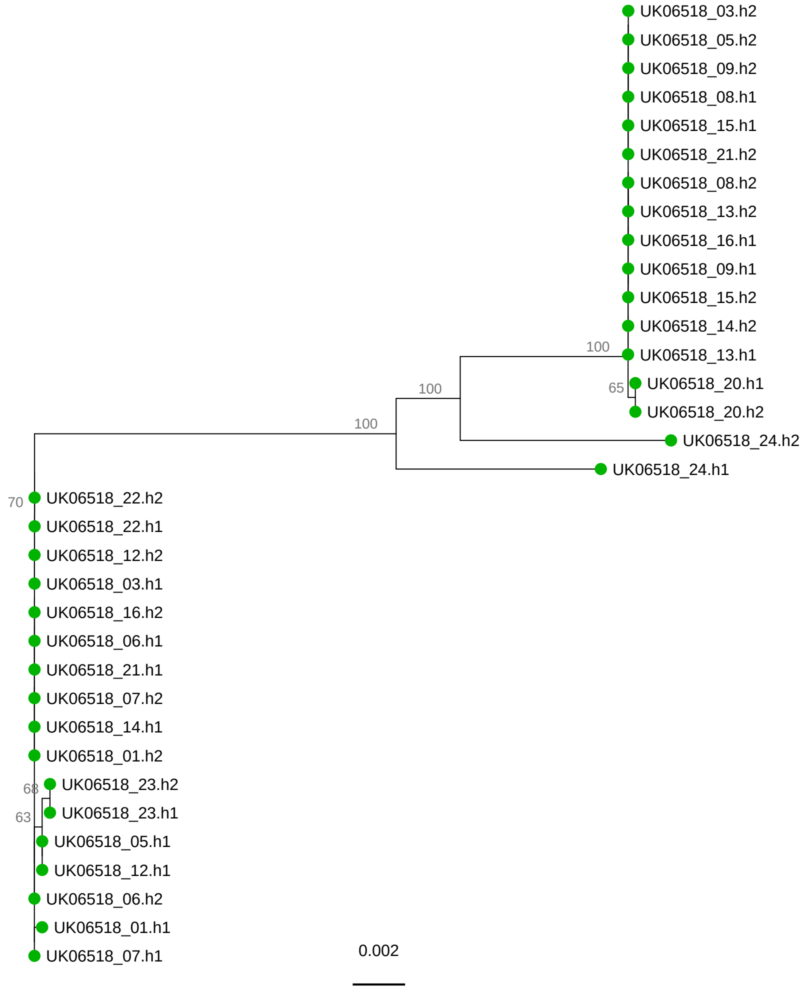
UK06518_03.h2
UK06518_05.h2
UK06518_09.h2
UK06518_08.h1
UK06518_15.h1
UK06518_21.h2
UK06518_08.h2
UK06518_13.h2
UK06518_16.h1
UK06518_09.h1
UK06518_15.h2
UK06518_14.h2
100
UK06518_13.h1
65
UK06518_20.h1
100
UK06518_20.h2
100
UK06518_24.h2
UK06518_24.h1
70
UK06518_22.h2
UK06518_22.h1
UK06518_12.h2
UK06518_03.h1
UK06518_16.h2
UK06518_06.h1
UK06518_21.h1
UK06518_07.h2
UK06518_14.h1
UK06518_01.h2
68
UK06518_23.h2
63
UK06518_23.h1
UK06518_05.h1
UK06518_12.h1
UK06518_06.h2
UK06518_01.h1
UK06518_07.h1
0.002

# UK10545

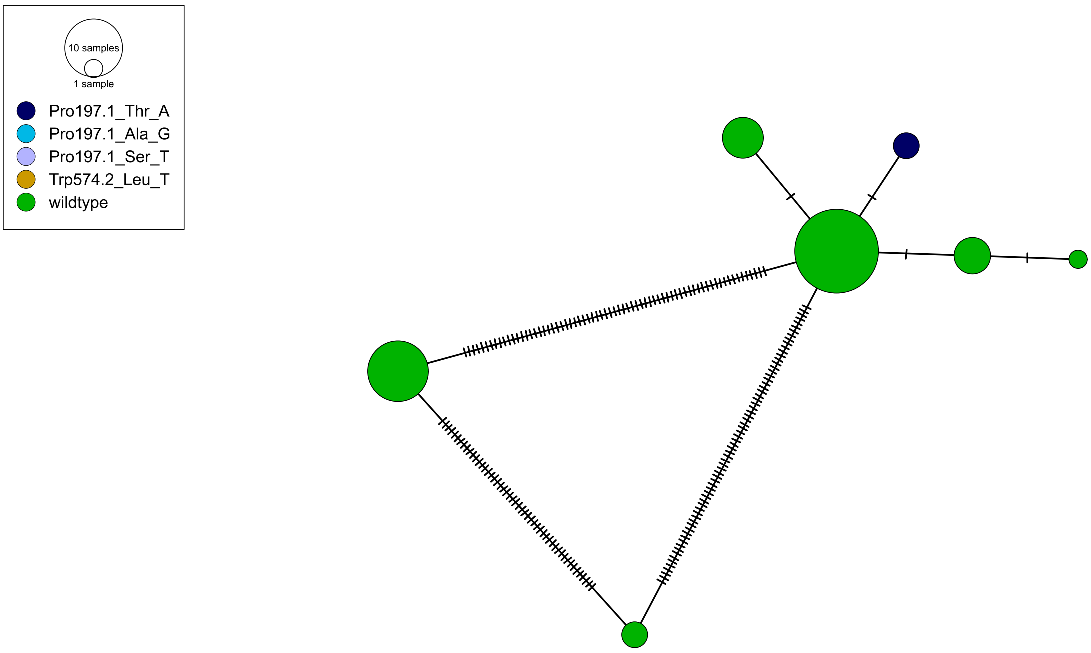

10 samples
1 sample
Pro197.1_Thr_A
Pro197.1_Ala_G
Pro197.1_Ser_T
Trp574.2_Leu_T
wildtype


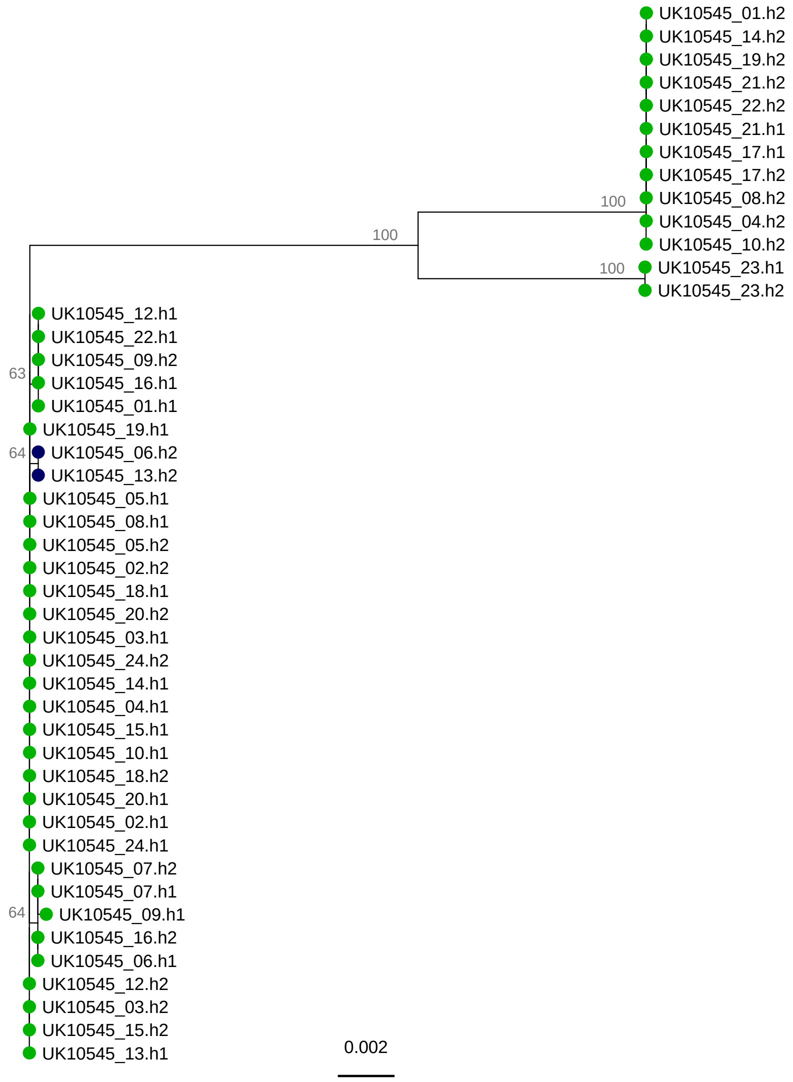

UK10545_01.h2
UK10545_14.h2
UK10545_19.h2
UK10545_21.h2
UK10545_22.h2
UK10545_21.h1
UK10545_17.h1
UK10545_17.h2
100
UK10545_08.h2
UK10545_04.h2
100
UK10545_10.h2
100
UK10545_23.h1
UK10545_23.h2
UK10545_12.h1
UK10545_22.h1
UK10545_09.h2
63
UK10545_16.h1
UK10545_01.h1
UK10545_19.h1
64
UK10545_06.h2
UK10545_13.h2
UK10545_05.h1
UK10545_08.h1
UK10545_05.h2
UK10545_02.h2
UK10545_18.h1
UK10545_20.h2
UK10545_03.h1
UK10545_24.h2
UK10545_14.h1
UK10545_04.h1
UK10545_15.h1
UK10545_10.h1
UK10545_18.h2
UK10545_20.h1
UK10545_02.h1
UK10545_24.h1
UK10545_07.h2
UK10545_07.h1
64
UK10545_09.h1
UK10545_16.h2
UK10545_06.h1
UK10545_12.h2
UK10545_03.h2
UK10545_15.h2
UK10545_13.h1
0.002
